# Identification of potential inhibitors of cutaneous Melanoma and Non-Melanoma skin cancer cells through in-vitro and in-silico screening of a small library of Phenolic compounds

**DOI:** 10.1101/2022.02.28.482167

**Authors:** Samuel T. Boateng, Tithi Roy, Mercy E. Agbo, Sergette Banang-Mbeumi, Roxane-Cherille N. Chamcheu, Marion Bramwell, Long K. Pham, Keith E. Jackson, Ronald A. Hill, Bolni Marius Nagalo, Tatiana Efimova, Jean Fotie, Jean Christopher Chamcheu

## Abstract

Melanoma and non-melanoma skin cancers are the most-lethal and commonest forms of skin cancers, that affecting one-fifth of the US population. With the aim of identifying new lead compounds as starting point for attaining cost-effective therapies, a small library of about 90 molecules was screened *in vitro* against A375, SKMEL-28, A431, SCC-12 skin cancer cell lines. About 35 of them, mainly dihydroquinolines, C–C and C–N linked biphenyls, and substituted methylgallate or aniline derivatives, displayed low-micromolar range activities, primarily against the A431 and SCC-12 squamous carcinoma cell lines, with only a handful of these compounds displaying any activity against the A375 and SKMEL-28 melanoma cell lines. Compounds **11** (A431: IC_50_ = 5.0 µM, SCC-12: IC_50_ = 2.9 µM, SKMEL-28: IC_50_ = 4.9 µM, A375: IC_50_ = 6.7 µM) and **13** (A431: IC_50_ = 5.0 µM, SCC-12: IC_50_ = 3.3 µM, SKMEL-28: IC_50_ = 13.8 µM, A375: IC_50_ = 17.1 µM) were the most active across all these cell lines. Furthermore, many of the hit compounds showed little to no activity against mammalian nontumorigenic immortalized HaCaT cells, with a far better selectivity index than cisplatin (a well-known anticancer agent used as a positive control). Compounds **11** and **13** significantly and dose-dependently induced apoptosis of SCC-12 and SK-MEL-28 cells as evidenced by the downregulation of Bcl-2 and upregulation of Bax protein expression levels, and by cleaved caspase-3, caspase-9 and PARP levels. Both agents also significantly reduced scratch wound healing, colony formation, and activated expression levels of major cancer molecular targets such as RSK/AKT/ERK1/2 and S6K1. To provide a better attribute profile for each of the hit molecules, in-silico target(s) prediction, pharmacokinetic and ADMET studies are also reported, together with some preliminary structure-activity relationship outlines. The SwissTargetPrediction web-based tool identified CDK8, CLK4, nuclear receptor ROR, tyrosine protein-kinase Fyn/LCK, ROCK1/2, and PARP, all of which are dysregulated in skin cancers, as likely targets for these hit compounds. Furthermore, the SwissADME web_tool predicted these compounds to exhibit high GI tract absorption, good skin permeation, and a viable biodegradability profile. To summarize, these data highlight the promising anticancer potential of these small molecules leads, warranting further investigation and/or optimization towards obtaining clinical candidates for combatting both melanoma and non-melanoma skin cancers.

## 1. Introduction

Skin cancer is considered one of the most prevalent cancer types globally, with its prevalence and mortality expected to keep growing at alarming rates.^1, 2^ Non-melanoma and melanoma skin cancers affect approximately 3 million people globally each year.^3^ Studies have shown that the lessened ozone layer protection vs. historical levels, resulting in greater human exposure to UVB rays at the Earth’s surface, is one of the major risk factors for the development of squamous cell carcinoma (SCC), basal cell carcinoma (BCC), and melanoma, with the incidence of these illnesses, predicted to continue to increase.^4–, 6^ One of the most effective therapeutic approaches at the early stages for skin cancers include misplaced surgical tumor excision^1, 2^; however, surgery can result in disfigurement, requiring skin grafts for covering the defects, and such surgeries can be quite costly-prohibitively so for some patients.^7–9^ Arresting proliferation or inducing programmed cell death in proliferating cancer cells is another effective cancer therapy option.^10–13^ Significant advances have been made toward a better understanding of mechanisms through which melanoma and non-melanoma skin cancers are triggered and sustained. Hotspot mutations of the oncogenes BRAF and NRAS are the most common genetic alterations in cutaneous melanoma, and such mutations are known drivers of all melanomas, as mutant BRAF and NRAS can independently activate the downstream MEK1/2-ERK1/2 oncogenic signal transduction pathway.^14–17^ Although inhibitors of BRAF and MEK have shown significant survival benefits in clinical trials, the prognostic significance of BRAF and NRAS mutations outside of clinical trials remains unclear.^14–17^ The overexpression and activation of V-akt murine thymoma viral oncogene homolog (AKT, also known as protein kinase B)^11^ and related signaling pathways are also known to be major contributing factors in many cancers, including some lung cancers, esophageal squamous cell carcinomas and various skin cancers.^18^ Other signaling pathways, including those involving mitogen-activated protein kinase (MAPK),^19–21^ phosphoinositide 3-kinase (PI3K)-Akt,^22–24^ mammalian target of rapamycin (*mTOR*),^22, 25, 26^ nuclear factor κ-light-chain-enhancer of activated B cell (NF-κB),^27^ Janus kinase-signal transducer and activator of transcription (JAK-STAT), transforming growth factor β (TGF-β),^28, 29^ and Notch are shown to play an important role in skin homeostasis, and in melanoma and non-melanoma cancer progression.^18^ Knowledge of these potential drug targets constitutes an important tool towards devising mechanistically novel anti-skin-cancer agents. Despite these remarkable advances in our scientific understanding, current available/validated/approved pharmacological treatments still face severe limitations, in many instances due to the low penetrability of the available drugs into stratum corneum or lesions,^12, 30, 31^ leading to their inability to reach the tumor cells and tissues well enough to achieve robustly efficacious levels.^32^ This low bioavailability at the site of action usually requires the administration of higher doses, resulting in skin irritation or worse, increased severity of side-effects or toxicities.^33–35^ The emergence of new therapies designed to stimulate the immune system, including immune checkpoint inhibitors (ICIs), in order to promote the immunorecognition and the elimination of tumor cells, has provided new hope for the effective treatment of a wide range of malignancies, such as melanoma, renal cell carcinoma, non-small cell lung cancer (NSCLC), and bladder cancers, to name just a few.^36, 37^ Although these agents have dramatically improved the cancer prognosis for many patients; these treatments are unfortunately associated with significant side-effects, due to an uncontrolled activation of the immune system. Furthermore, only about 50% of patients with these cancers currently respond to ICI immunotherapy, the precarity which of course argues for continued search for novel potential targets,^37, 38^ in hopes of devising new, effective and economically sustainable therapies against melanoma and non-melanoma skin cancers.

As part of a continuing effort to introduce new-molecular-scaffold-based entities with low toxicity and enhanced efficacy into the cutaneous cancer treatment pipeline, our research groups have shown that simple phenolic compounds such as a green tea polyphenol, namely (-)-epigallocatechin-3-gallate (EGCG), could modulate human keratinocyte-induced responses and alleviate imiquimod-induced murine psoriasiform dermatitis.^39, 40^ We recently showed that fisetin and its analogs inhibit the PI3K/Akt/mTOR and MAPK and effector pathways in organotypic human inflammatory skin and melanoma models,^41, 42^ as well as c-Kit, CDK2 and mTOR kinases in melanoma and non-melanoma skin cancers.^25^ Herein, we report the *in vitro* screening of a small library of about 90 compounds previously synthesized in our laboratory against melanoma (A375 and SKMEL-28) and non-melanoma (A431 and SCC-12) skin cancer cell lines. *In-silico* studies were carried out, aimed at target(s) prediction and at predicting pharmacokinetic and ADMET (absorption, distribution, metabolism, excretion, and toxicity) attributes for selected hit molecules. Mode-of-action explorations by way of in-vitro cellular and molecular biological methods were conducted for the two most active compounds (**11** and **13**). We report preliminary structure-activity relationship, prospective targets and discuss these in the context of other pertinent pharmacologic and biopharmaceutic aspects of our findings.

## 2. Results and discussion

### 2.1. Chemistry

The focus here is exclusively on the synthesis of 35 compounds from the library that displayed a relevant behavior during either the *in vitro* biological or the *in silico* studies. As such, compounds **1** and **2** are methyl 3,4,5-trimethoxybenzoate derivatives closely related to methyl gallate and were prepared through electrophilic bromination and nitration, respectively, as illustrated in **Scheme 1**.^43^

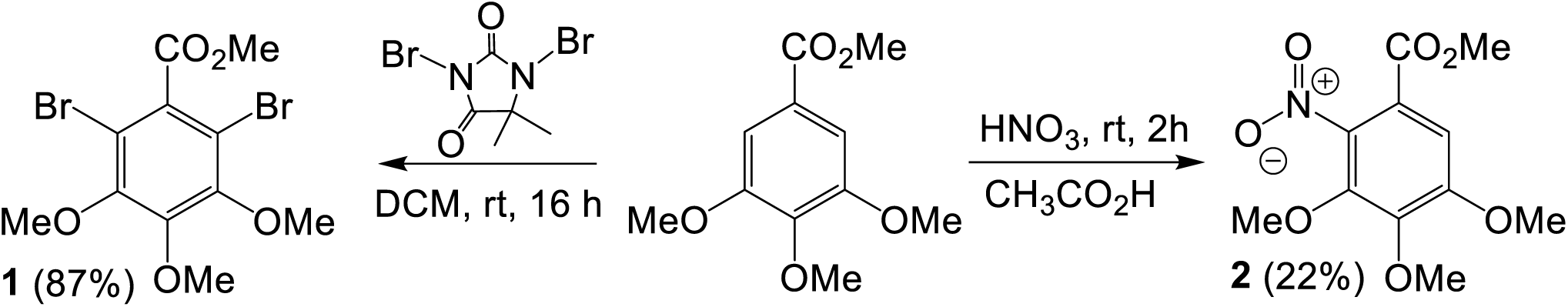

**Scheme 1:** Synthesis of methyl 3,4,5-trimethoxybenzoate derivatives^43^

4-cyclohexenylaniline derivatives (**3** and **4**) were obtained by reacting the corresponding aniline derivative with cyclohexanone in the presence of a catalytic amount of molecular iodine at 160 °C, under neat conditions (Scheme 2).^44^ The same reaction conditions were implemented to prepare of 2,2,4-trimethyl-1,2-dihydroquinoline derivatives (**5** and **6**) by reacting the corresponding aniline derivative with acetone through a Skraup-Doebner-Von-Miller quinoline synthesis mechanism (Scheme 3).^43, 45^ Under identical conditions but using cyclopentanone instead of acetone (Scheme 4), compound **7** was produced.^45^ On the other hand, reacting a 2,2,4-trimethyl-1,2-dihydroquinoline with either cyclohexanone or cyclopentanone produced compounds **8** and **9**, respectively, as illustrated in Scheme 5.^44, 45^

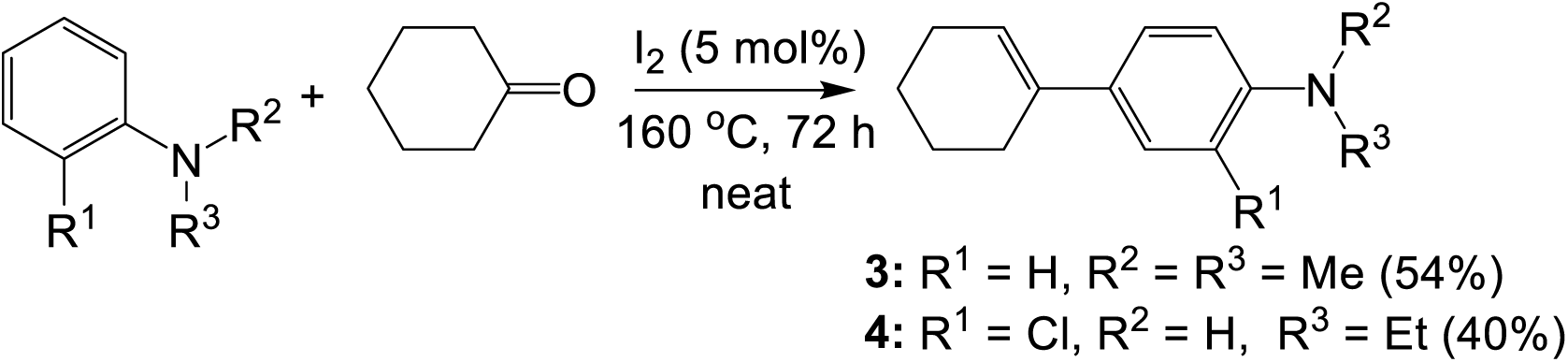

**Scheme 2:** Synthesis of 4-cyclohexenylaniline derivatives^44^

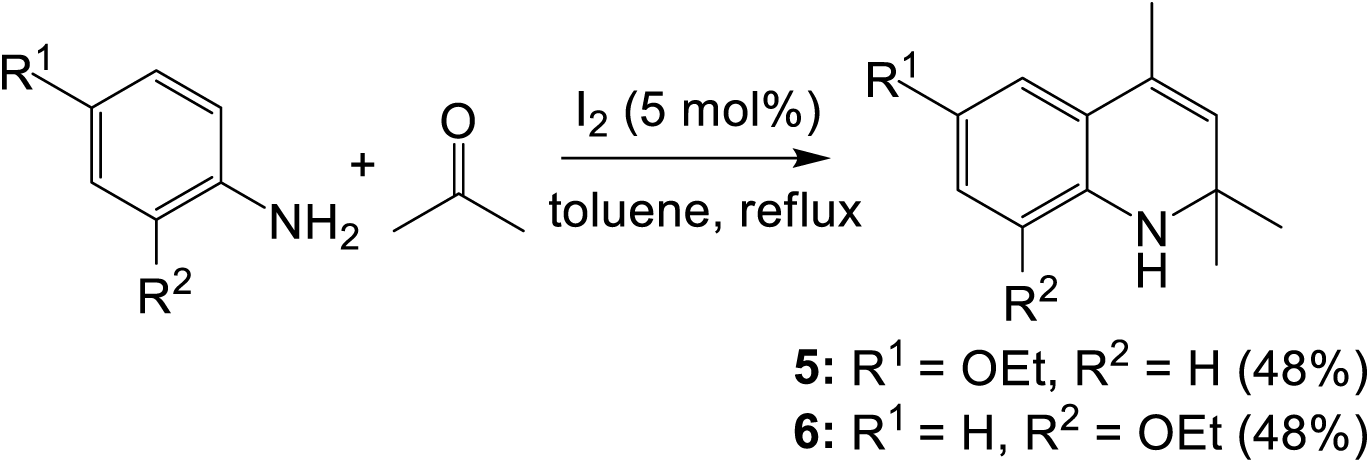

**Scheme 3:** Synthesis of 2,2,4-trimethyl-1,2-dihydroquinoline derivatives^43, 45^

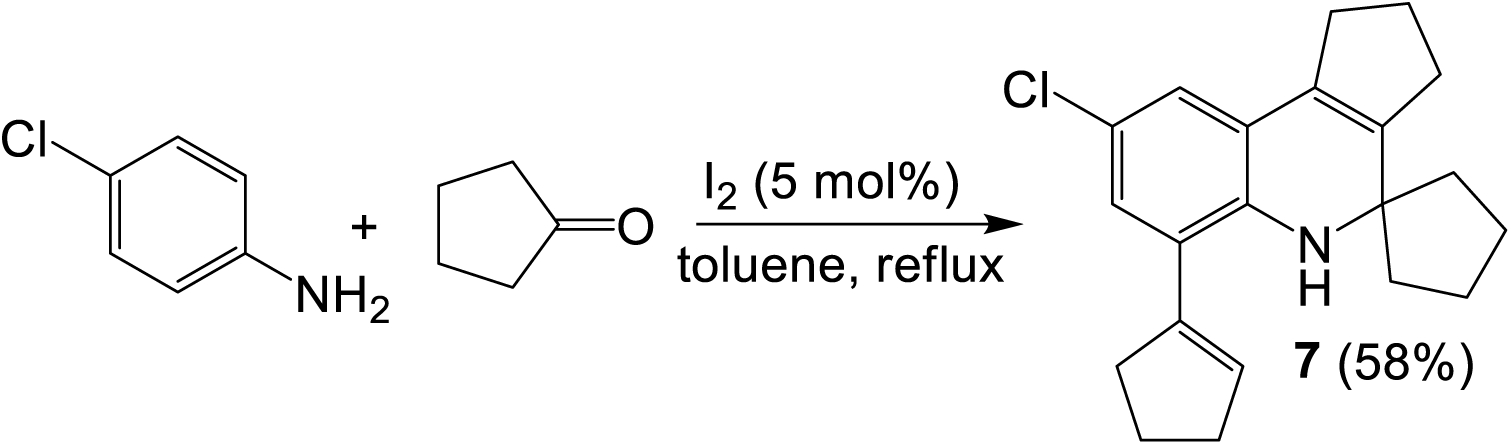

**Scheme 4:** Reaction of 4-ethoxyaniline derivatives with cyclopentanone^45^

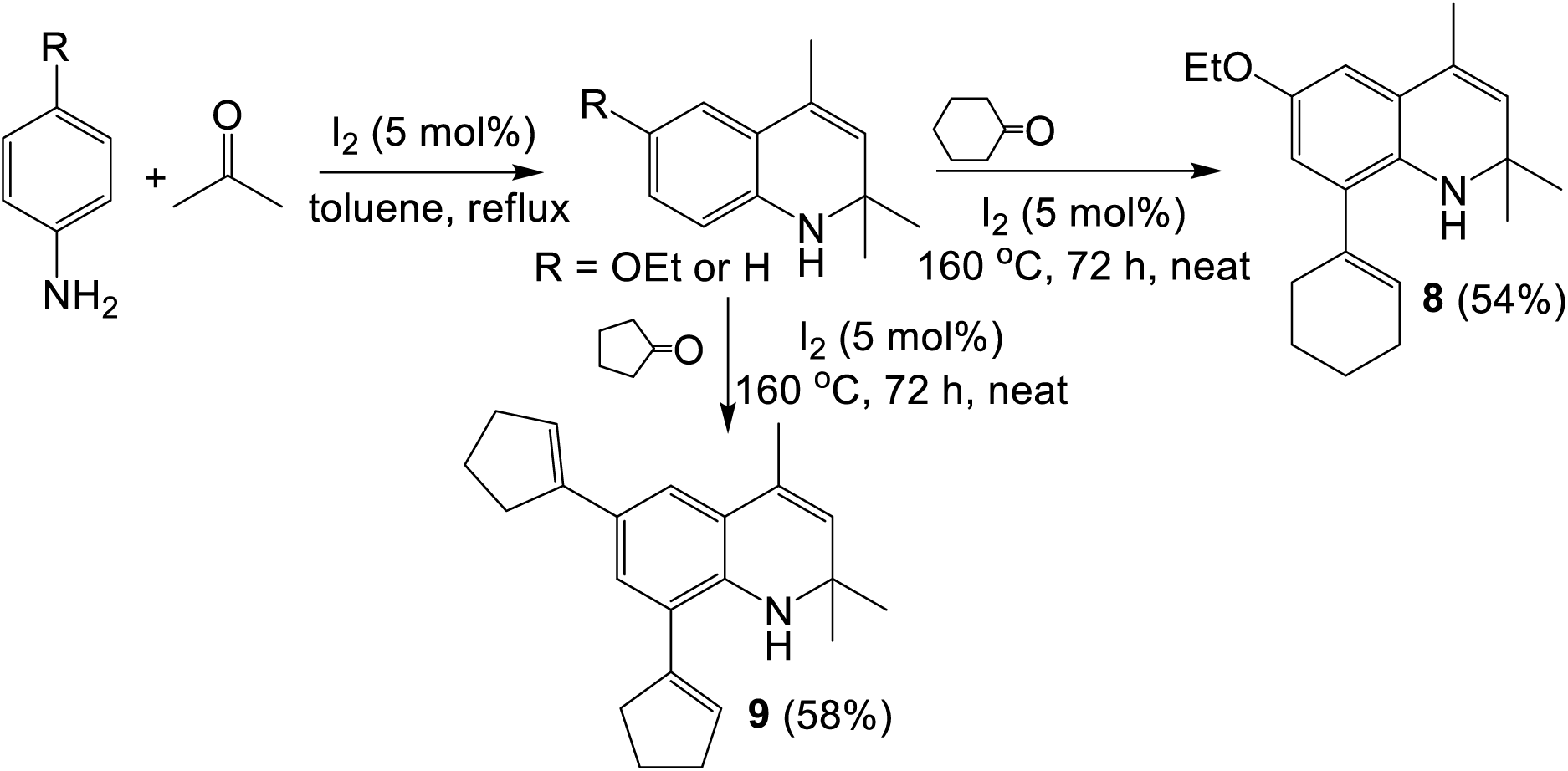

**Scheme 5:** Reaction of 2,2,4-trimethyl-1,2-dihydroquinoline derivative with cyclohexanone or cyclopentanone^44, 45^

C–N linked biphenyls were prepared by dimerizing the corresponding aniline derivative via a silver(I) catalyzed direct C–H functionalization, as illustrated in Scheme 5.^46^ As for 2’-aminobiphenyl-2-ols, the largest family of compounds screened during this study, they were also prepared using silver(I) as a catalyst under reaction conditions very similar to those used in the preparation of C–N linked biphenyls. In this latter case, the aniline derivative was cross-coupled with the appropriate phenol via a regioselective aerobic oxidative cross-coupling, resulting also in a direct C–H functionalization.^47^ However, sodium persulfate (Na_2_S_2_O_8_) used as the oxidizing agent in C–N coupling reactions was replaced in this case by hydrogen peroxide (H_2_O_2_), resulting in a C–C coupling.^47^

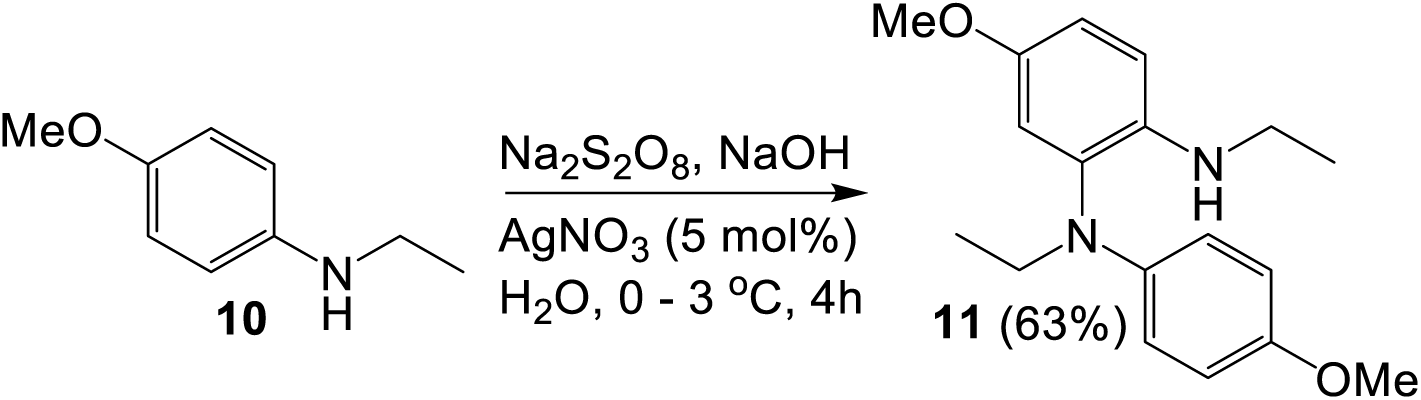

**Scheme 5:** Synthesis of C–N linked biphenyls^46^

As such, 2-naphthol was reacted with a number of aniline derivatives (Scheme 6), and by varying the substituents either on the phenol or aniline starting materials, a number of 2’-aminobiphenyl-2-ol derivatives were prepared, as shown in **Figure 1**. The structure of compounds **12**, **13** and **16** was unambiguously confirmed by single crystal X-ray diffraction, and the molecular structures with thermal ellipsoid representation at 50% probability are provided.

**Figure 1:**
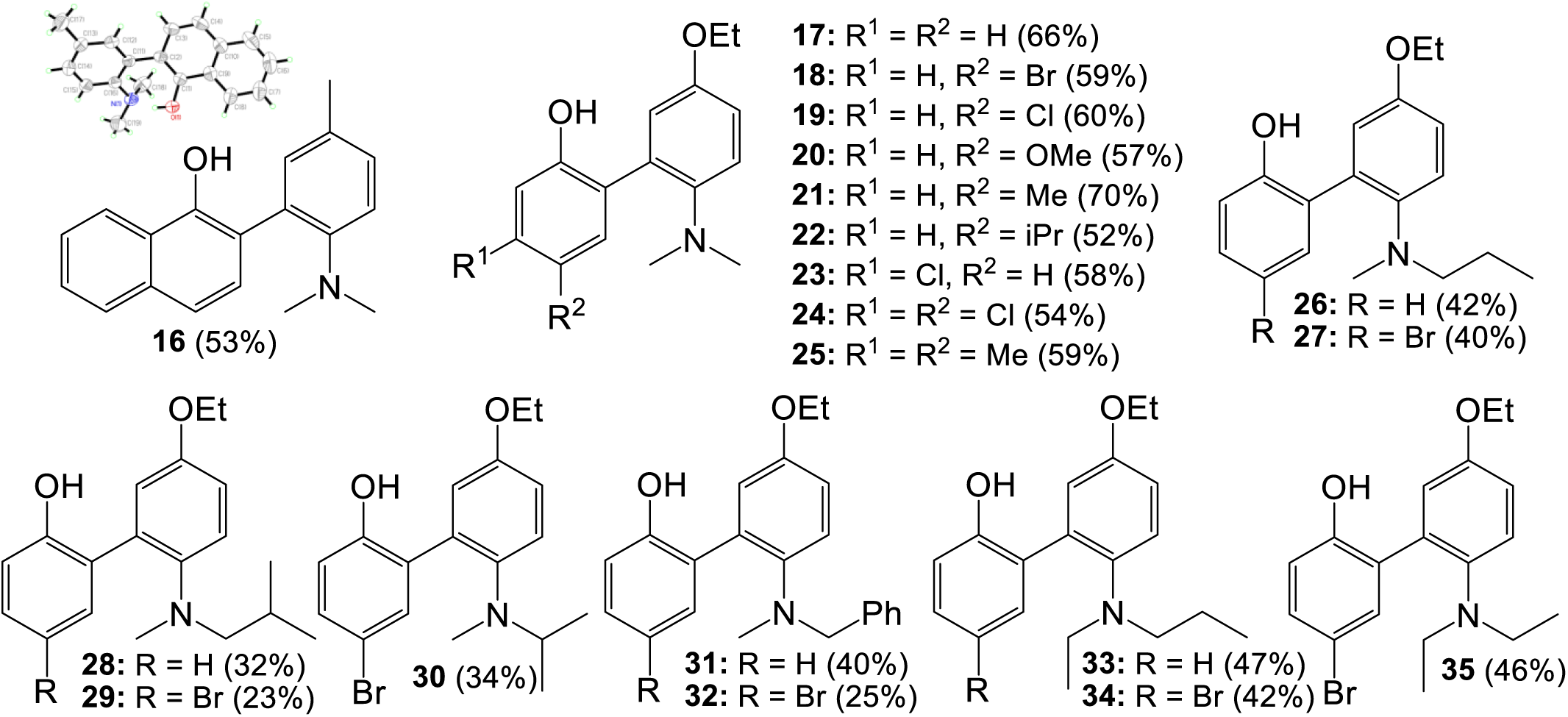
List of other 2’-aminobiphenyl-2-ol derivatives screened in this study. The single crystal X-ray data for **16** (CCDC 1442428)^47^ are deposited in the Cambridge database, and copies of these materials can be obtained free of charge from CCDC, 12 Union Road, Cambridge CB2 1EZ, UK, http://www.ccdc.am.ac.uk.

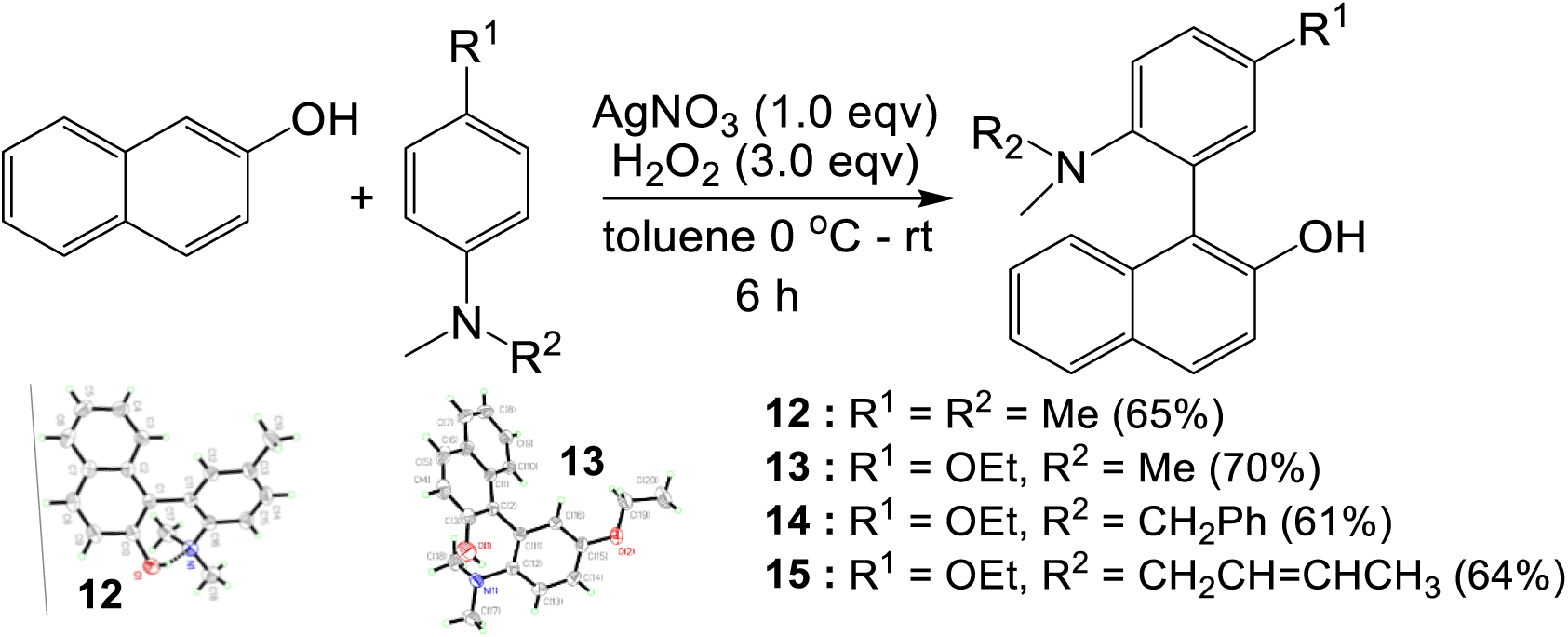

**Scheme 6:** Synthesis of 1-(2-aminophenyl)naphthalen-2-ol derivatives. The single crystal X-ray data for **12** (CCDC 821630)^48^ and **13** (CCDC 1442427)^47^ are deposited in the Cambridge database, and copies of these materials can be obtained free of charge from CCDC, 12 Union Road, Cambridge CB2 1EZ, UK, http://www.ccdc.am.ac.uk.

### 2.2. In vitro anticancer screening

The growth-suppressive and cytotoxic activities of a small library of about 90 compounds containing the above-described molecules were then screened against many of skin cancer cell lines including melanoma (A375 and SKMEL-28) and non-melanoma (A431 and SCC-12), with HaCaT keratinocytes used as a control, using *in vitro* MTT and trypan blue dye exclusion assays. The activity was assessed as previously reported,^25, 41^ and based in each case on the concentration of the sample capable of reducing the measured viability of the cell population to 50% of the maximum for each of the cell lines, and thus is expressed as IC_50_. Hit compounds are identified as any molecule displaying a low micromolar activity against any of the cell lines used in the assay.

Many of the hit compounds appear to be primarily active against the non-melanoma A431 and SCC-12 cell lines at the micromolar range, and only a handful of them display any activity against the melanoma A375 and SKMEL-28 cell lines, as summarized in **Table 1**.

**Table 1:**
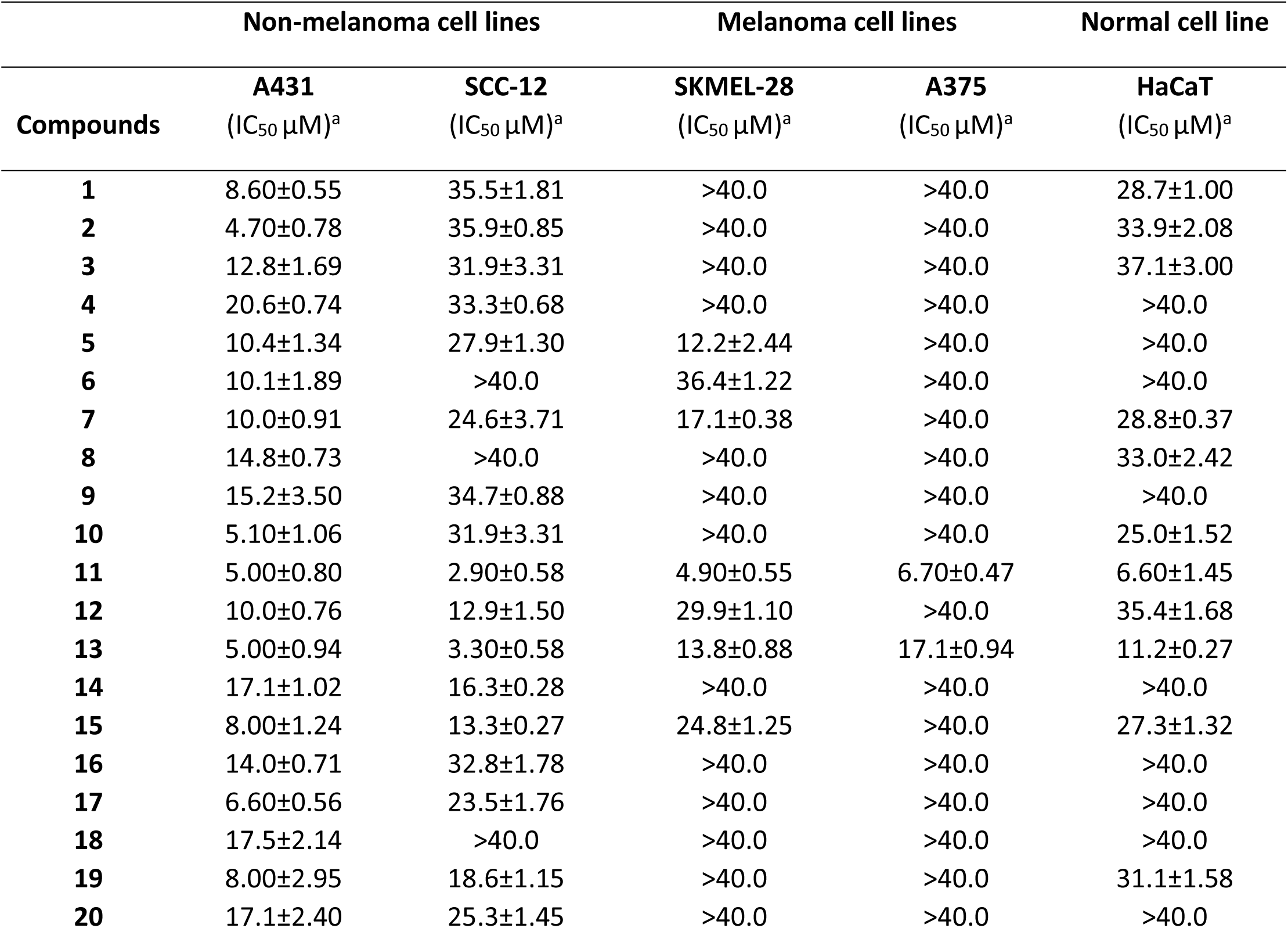
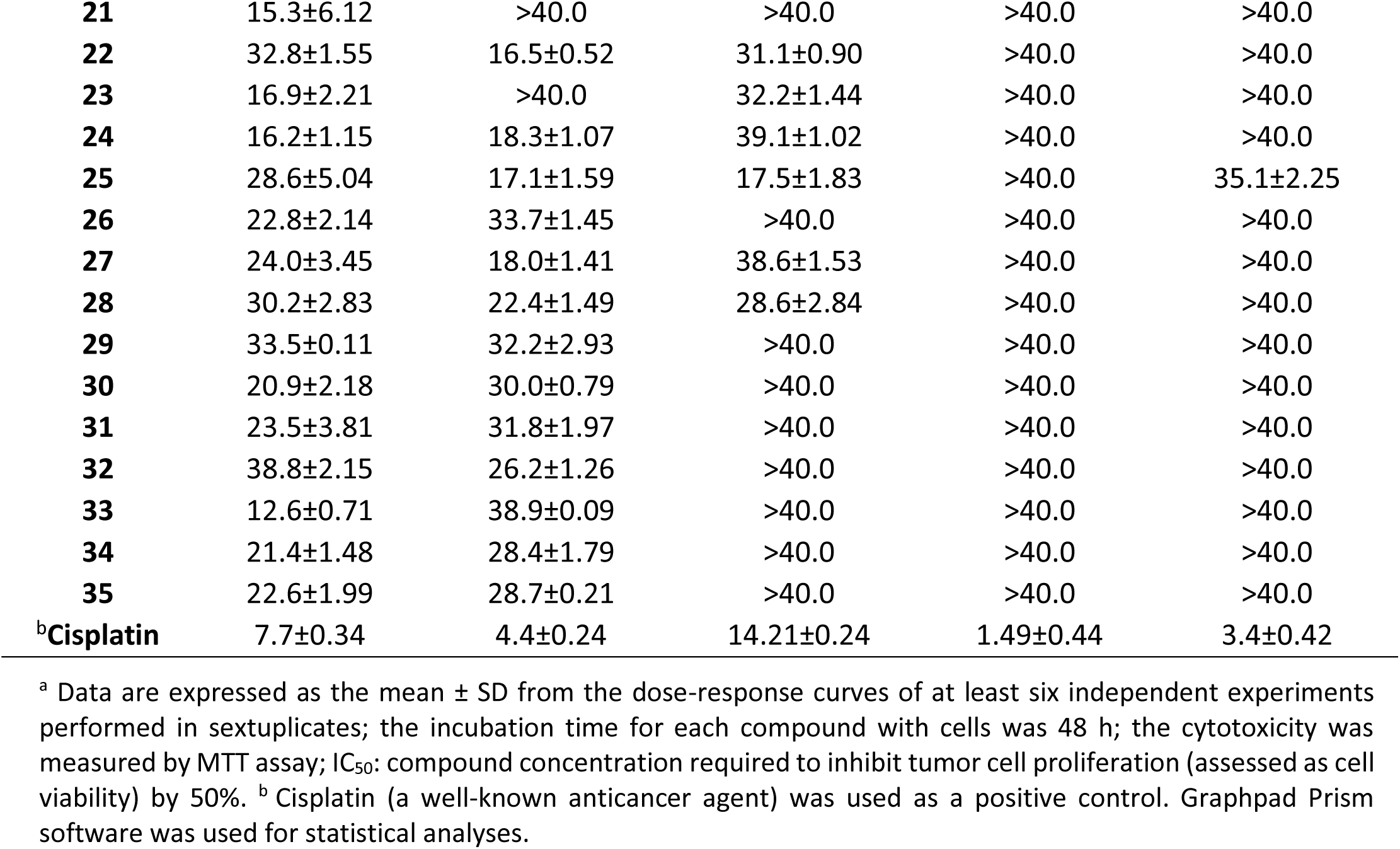
Cytotoxicity (IC_50_; μM) values of the hit compounds against four human skin cancer cell lines; melanoma (A375 and SKMEL-28) and non-melanoma (A431 and SCC-12) cells, with HaCaT keratinocytes as a control

| Compounds | Non-melanoma cell lines |  | Melanoma cell lines |  | Normal cell line |
| --- | --- | --- | --- | --- | --- |
| | A431<br>(IC<sub>50</sub> μM)<sup>a</sup> | SCC-12<br>(IC<sub>50</sub> μM)<sup>a</sup> | SKMEL-28<br>(IC<sub>50</sub> μM)<sup>a</sup> | A375<br>(IC<sub>50</sub> μM)<sup>a</sup> | HaCaT<br>(IC<sub>50</sub> μM)<sup>a</sup> |
| 1 | 8.60±0.55 | 35.5±1.81 | >40.0 | >40.0 | 28.7±1.00 |
| 2 | 4.70±0.78 | 35.9±0.85 | >40.0 | >40.0 | 33.9±2.08 |
| 3 | 12.8±1.69 | 31.9±3.31 | >40.0 | >40.0 | 37.1±3.00 |
| 4 | 20.6±0.74 | 33.3±0.68 | >40.0 | >40.0 | >40.0 |
| 5 | 10.4±1.34 | 27.9±1.30 | 12.2±2.44 | >40.0 | >40.0 |
| 6 | 10.1±1.89 | >40.0 | 36.4±1.22 | >40.0 | >40.0 |
| 7 | 10.0±0.91 | 24.6±3.71 | 17.1±0.38 | >40.0 | 28.8±0.37 |
| 8 | 14.8±0.73 | >40.0 | >40.0 | >40.0 | 33.0±2.42 |
| 9 | 15.2±3.50 | 34.7±0.88 | >40.0 | >40.0 | >40.0 |
| 10 | 5.10±1.06 | 31.9±3.31 | >40.0 | >40.0 | 25.0±1.52 |
| 11 | 5.00±0.80 | 2.90±0.58 | 4.90±0.55 | 6.70±0.47 | 6.60±1.45 |
| 12 | 10.0±0.76 | 12.9±1.50 | 29.9±1.10 | >40.0 | 35.4±1.68 |
| 13 | 5.00±0.94 | 3.30±0.58 | 13.8±0.88 | 17.1±0.94 | 11.2±0.27 |
| 14 | 17.1±1.02 | 16.3±0.28 | >40.0 | >40.0 | >40.0 |
| 15 | 8.00±1.24 | 13.3±0.27 | 24.8±1.25 | >40.0 | 27.3±1.32 |
| 16 | 14.0±0.71 | 32.8±1.78 | >40.0 | >40.0 | >40.0 |
| 17 | 6.60±0.56 | 23.5±1.76 | >40.0 | >40.0 | >40.0 |
| 18 | 17.5±2.14 | >40.0 | >40.0 | >40.0 | >40.0 |
| 19 | 8.00±2.95 | 18.6±1.15 | >40.0 | >40.0 | 31.1±1.58 |
| 20 | 17.1±2.40 | 25.3±1.45 | >40.0 | >40.0 | >40.0 |
| **21** | 15.3±6.12 | >40.0 | >40.0 | >40.0 | >40.0 |
| **22** | 32.8±1.55 | 16.5±0.52 | 31.1±0.90 | >40.0 | >40.0 |
| **23** | 16.9±2.21 | >40.0 | 32.2±1.44 | >40.0 | >40.0 |
| **24** | 16.2±1.15 | 18.3±1.07 | 39.1±1.02 | >40.0 | >40.0 |
| **25** | 28.6±5.04 | 17.1±1.59 | 17.5±1.83 | >40.0 | 35.1±2.25 |
| **26** | 22.8±2.14 | 33.7±1.45 | >40.0 | >40.0 | >40.0 |
| **27** | 24.0±3.45 | 18.0±1.41 | 38.6±1.53 | >40.0 | >40.0 |
| **28** | 30.2±2.83 | 22.4±1.49 | 28.6±2.84 | >40.0 | >40.0 |
| **29** | 33.5±0.11 | 32.2±2.93 | >40.0 | >40.0 | >40.0 |
| **30** | 20.9±2.18 | 30.0±0.79 | >40.0 | >40.0 | >40.0 |
| **31** | 23.5±3.81 | 31.8±1.97 | >40.0 | >40.0 | >40.0 |
| **32** | 38.8±2.15 | 26.2±1.26 | >40.0 | >40.0 | >40.0 |
| **33** | 12.6±0.71 | 38.9±0.09 | >40.0 | >40.0 | >40.0 |
| **34** | 21.4±1.48 | 28.4±1.79 | >40.0 | >40.0 | >40.0 |
| **35** | 22.6±1.99 | 28.7±0.21 | >40.0 | >40.0 | >40.0 |
| <sup>b</sup>**Cisplatin** | 7.7±0.34 | 4.4±0.24 | 14.21±0.24 | 1.49±0.44 | 3.4±0.42 |

<sup>a</sup> Data are expressed as the mean ± SD from the dose-response curves of at least six independent experiments performed in sextuplicates; the incubation time for each compound with cells was 48 h; the cytotoxicity was measured by MTT assay; IC<sub>50</sub>: compound concentration required to inhibit tumor cell proliferation (assessed as cell viability) by 50%. <sup>b</sup> Cisplatin (a well-known anticancer agent) was used as a positive control. Graphpad Prism software was used for statistical analyses.

In fact, the two methylgallate derivatives **1** (IC_50_ 8.60 µM) and **2** (IC_50_ 4.70 µM) and the aniline derivative **10** (IC_50_ 5.10 µM) displayed a low micromolar activity against the A431 cell line, while the other aniline derivatives **3** and **4** were not as active. However, these simple benzene derivatives are all inactive against the other cancer cell lines used in this study. Nevertheless, the dimerization of **10** resulted in compound **11** (A431: IC_50_ = 5.00 µM, SCC-12: IC_50_ = 2.90 µM, SKMEL-28: IC_50_ = 4.90 µM, A375: IC_50_ = 6.70 µM), the most potent of the series across all the four cell lines. While **11** is the most active against all cell lines used in this study, especially in SCC-12 (a non-melanoma cell line) and SKMEL-28 (a melanoma cell line), it exhibited a low selectivity although comparable to that of the control drug cisplatin.

The 1,2-dihydroquinoline derivatives **5** (A431: IC_50_ = 10.4 µM, SKMEL-28: IC_50_ = 12.2 µM), **6** (A431: IC_50_ = 10.1 µM, SKMEL-28: IC_50_ = 36.4 µM) and **7** (A431: IC_50_ = 10.0 µM, SKMEL-28: IC_50_ = 17.1 µM) displayed comparable levels of activity against the non-melanoma cell A431, but were more active against the melanoma cell line SKMEL-28 than they were against SCC-12 (the other non-melanoma cell line), while completely inactive against A375. They also displayed a better selectivity index than the simple benzene derivatives. As for the C–C-linked biphenyl group **12 – 35**, many of them displayed some activity against A431, including **12** (A431: IC_50_ = 10.0 µM), **15** (A431: IC_50_ = 8.00 µM), **17** (A431: IC_50_ = 6.60 µM), **19** (A431: IC_50_ = 8.00 µM), and **33** (A431: IC_50_ = 12.60 µM), while being only marginally active or completely inactive against the other cell lines, including SCC-12. Compound **13** (A431: IC_50_ = 5.00 µM, SCC-12: IC_50_ = 3.30 µM, SKMEL-28: IC_50_ = 13.8 µM, A375: IC_50_ = 17.1 µM) was the most active of the group across all the cell lines, with a better selectivity index than **11** or cisplatin. The diversity of substituents on the biphenyl rings appears to be very limited, and as such, no evident electronic or steric trend could be observed. For example, the only difference between **12** and **13** is that the para-methyl group on the aniline derivative in **12** is replaced at the same position by an ethoxy group in **13**; although both groups are electron-donating in nature, these two compounds displayed a wide difference in activity. Surprisingly, compound **14**, carrying an ethoxy group at the same position as **13**, but a benzyl rather than a methyl group on the nitrogen atom, displayed a similar activity as **12** that carries a methyl group at both positions. Furthermore, the presence of 4-Br (**18**), 4-Cl (**19**), 4-OMe (**20**), 4-Me (**21**), 4-iPr (**22**), 3,4-diCl (**24**), or 3,4-diMe (**25**) on the phenol ring did not have a significant impact on the activity of these compounds as compared to **17**, which carries no substituent on the phenol ring. The variation of the substituents on the nitrogen of the aniline also does not appear to significantly impact on the activity of the resulting compounds, although some differences in activity were observed, primarily with the A431 cell line. As such, it is still unclear why compounds **11** and **13** appear to have significantly greater activity than the remaining hit compounds across the four cell lines used in the study. Furthermore, cisplatin, a well-known anticancer agent used as a positive control, appeared to be less cytotoxic than some of the hit compounds towards these cancer cell lines and displayed a much lower selectivity index than many of these small molecules, as indicated by its activity against mammalian nontumorigenic immortalized HaCaT cells. In fact, some compounds such as **2** (A431: IC_50_ = 4.70 µM, HaCaT: IC_50_ = 33.9 µM), **10** (A431: IC_50_ = 5.00 µM, HaCaT: IC_50_ = 25.0 µM), or **19** (A431: IC_50_ = 8.00 µM, HaCaT: IC_50_ = 31.1 µM) for example, exhibited a similar or better activity than cisplatin against A431 cell line, and almost not toxicity against HaCaT, even if the activity was not maintained across all cell lines. This is a good indication that the activity and selectivity of this family of compounds can be improved by designing and preparing more appropriate analogs. As such, further study with more diversely substituted biphenyl analogs is needed to be able to develop a more complete structure-activity profile, and in the process hopefully improve their anticancer activity. More importantly, since these compounds bear some similarity to ellagic acid, a well-established anticancer molecule,^49–52^ further studies on the way in our lab will establish whether the planarity of ellagic acid or the atropisomerism of the hit compounds described in this report have any effects on the anticancer property of the respective molecules. The culmination will be a better understanding of the anticancer profile of the biphenyl family of compounds. Towards this objective, a preliminary target validation was undertaken using a combination of molecular docking and ADMET, western blotting, scratch wound healing, apoptosis, and colony formation assays. Beyond the scope of the current report, further investigations of target proteins are underway in our laboratory.

### 2.3. In silico target(s) prediction, pharmacokinetic and ADMET studies

Bio- and chemo-informatics approaches have progressively established themselves as important tools to estimate the most probable targets of small molecules.^53–55^ As such, computer-aided ligand-based target prediction techniques, also known as “target fishing,” have proven to be fast and exceedingly accurate in predicting the correct protein target(s) in drug discovery.^56, 57^ SwissTargetPrediction, the tool used in this study, quantifies similarities between compounds by computing a pair-wise comparison of 1D vectors describing molecular structures,^58^ determines the Tanimoto index between path-based binary fingerprints (FP2) through 2D measurements,^58^ and evaluates the Manhattan distance similarity between Electroshape 5D (ES5D) float vectors through a 3D assessment,^58–60^ to intuitively validate the “molecular similarity hypothesis,” based on the credence that similar molecules target common proteins.^58–61^ Based on these parameters, a “CombinedScore” of 0.5 or higher is an indication that the molecules are likely to share a common protein target.^58^ It should be clarified that these data are only suggestive of the probability for a molecule to interact with a given protein as a target, and not an indication of the compound being bioactive. Nevertheless, SwissTargetPrediction has documented and stored about 2,579 protein targets, 1,768 of which are human’s, 280,381 active compounds and more than 440,534 interactions.^58^

As such, in order to gain more insights into the behavioral profile of the hit compounds identified through whole-cell *in vitro* assays, their binding efficiency was compared *in silico*, to that of a natural and a commercial ligand, through a reverse screening study. These compounds appear to have a favorable binding affinity with only a selected number of enzymes, phosphodiesterases, kinases, proteases, oxidoreductases, erasers, and ligand-gated ion channels, as summarized in **Table 2 and 3**. Compounds **1 – 4** and **10 – 11**, which are primarily substituted benzene derivatives, displayed considerable interactions only with two kinases (RAF1 and dual-specificity protein kinase CLK4), one protease (MMP9), one oxidoreductase (monoamine oxidase B), and one eraser (deacetylase NAD-dependent Sirtuin 2). They also showed some strong binding with the cannabinoid receptor 2, and with the nuclear receptor ROR-gamma. Compounds **5 – 9**, which are dihydroquinoline derivatives, were less selective, exhibiting a strong binding with six enzymes (anandamide amidohydrolase, arachidonate 5 lipoxygenase, arachidonate 15 lipoxygenase, nicotinamide phosphoribosyltransferase, PARP and PI3-kinase p110-alpha subunit), three phosphodiesterases (4B, 5A and 10A), eight kinases (RAF1, ROCK1, ROCK2, serine-threonine protein kinase Chk1, tyrosine-protein kinase FYN, tyrosine protein kinase LCK, CDK8, and dual specificity protein kinase CLK4), one protease (MMP9), two oxidoreductase (COX-2 and monoamine oxidase B), and one eraser (deacetylase NAD dependent Sirtuin 2). These compounds (**5 – 9**) also displayed good interaction with other potential targets including the cholesterol ester transfer protein, the cannabinoid receptor 2, a primary active transporter (P-glycoprotein 1), and several of nuclear proteins and receptors (peroxisome-proliferator activated receptor-gamma, orphan nuclear receptor LRH-1, nuclear receptor ROR-gamma, and MDM2).

**Table 2:** Binding affinity of the hit compounds toward enzymes and phosphodiesterases as compared to natural and commercial ligands

| Ligands | Enzymes |  |  |  |  |  | Phosphodiesterases |  |  |
| --- | --- | --- | --- | --- | --- | --- | --- | --- | --- |
|  | Anandamide<br>amidohydrolase | Arachidonate 5<br>lipoxygenase | Arachidonate 15<br>lipoxygenase | Nicotinamide<br>phosphoribosyltra<br>nsferase | PARP | PI3-kinase<br>p110-alpha<br>subunit | Phosphodi<br>esterase 4B | Phosphodi<br>esterase 5A | Phosphodi<br>esterase 10A |
| **Natural** | **- 8.6** | **- 6.6** | **- 5.3** | **- 11.9** | **- 11.1** | **- 9.1** | **- 9.1** | **- 10.7** | **- 10.5** |
| **CL** | MK4409<br>**- 8.1** | α-boswellic acid<br>**- 8.5** | Hydroxyoctade<br>cadiomic acid<br>**- 7.3** | Daporinad<br>**- 9.7** | Olaparib<br>**- 11.1** | Copanlisib<br>**- 8.3** | Apremilast<br>**- 7.7** | Sildenafil<br>**- 8.8** | pubchem<br>629793<br>**- 8.2** |
| **1** | - 5.9 | - 4.6 | - 5.1 | - 5.6 | - 5.2 | - 5.2 | - 6.1 | - 5.8 | - 5.0 |
| **2** | - 6.3 | - 4.7 | - 5.1 | - 6.1 | - 6.8 | - 5.4 | - 6.2 | - 5.7 | - 5.9 |
| **3** | - 7.7 | - 6.8 | - 6.8 | - 7.8 | - 7.9 | - 7.9 | - 7.2 | - 7.3 | - 7.4 |
| **4** | **- 8.3** | - 6.9 | - 7.6 | - 7.9 | - 7.8 | - 7.7 | - 7.8 | - 7.2 | - 7.4 |
| **5** | - 7.9 | - 6.2 | - 6.8 | - 7.5 | - 7.7 | - 6.5 | - 6.9 | - 7.4 | - 7.5 |
| **6** | - 7.6 | - 6.1 | - 6.5 | - 7.2 | - 7.4 | - 6.3 | - 7.0 | - 7.3 | - 7.3 |
| **7** | **- 10.3** | **- 8.2** | **- 8.7** | **- 9.8** | - 7.8 | - 8.0 | **- 9.2** | **- 9.8** | **- 9.3** |
| **8** | **- 9.6** | **- 8.0** | **- 8.4** | **- 9.1** | - 7.7 | - 7.6 | **- 8.3** | **- 9.4** | - 7.7 |
| **9** | **- 9.1** | **- 8.6** | **- 9.1** | **- 10.5** | **- 9.6** | **- 9.0** | **- 9.3** | **- 10.7** | **- 8.7** |
| **10** | - 5.7 | - 4.7 | - 5.6 | - 5.6 | - 5.8 | - 5.5 | - 5.5 | - 5.3 | - 5.5 |
| **11** | - 7.5 | - 6.0 | - 6.9 | - 7.5 | - 7.8 | - 6.5 | - 6.5 | - 7.2 | - 6.3 |
| **12** | - 7.8 | **- 8.0** | - 7.3 | - 7.7 | - 7.8 | - 6.9 | - 7.6 | - 7.3 | **- 8.0** |
| **13** | **- 8.2** | - 6.1 | - 7.4 | - 7.7 | **- 8.3** | - 6.5 | - 7.8 | - 7.2 | - 7.4 |
| **14** | **- 8.4** | - 7.7 | - 8.0 | **- 8.9** | - 7.2 | - 7.2 | **- 8.4** | **- 8.3** | - 7.8 |
| **15** | - 7.8 | - 7.6 | - 7.6 | **- 8.2** | **- 8.4** | - 6.6 | - 7.0 | - 7.6 | - 7.6 |
| **16** | **- 8.3** | **- 8.0** | **- 8.2** | **- 8.6** | **- 9.0** | **- 9.3** | **- 8.0** | **- 8.2** | **- 8.5** |
| **17** | - 7.3 | - 6.2 | - 6.8 | - 7.2 | - 6.6 | - 7.0 | - 6.5 | **- 8.1** | - 6.7 |
| **18** | - 7.0 | - 6.4 | - 6.4 | - 7.0 | - 6.7 | - 6.3 | - 6.8 | - 7.7 | - 6.4 |
| **19** | - 7.4 | - 6.3 | - 6.6 | - 6.9 | - 6.8 | - 6.2 | - 6.8 | - 7.6 | - 6.4 |
| **20** | - 7.8 | - 6.0 | - 6.4 | - 6.9 | - 6.7 | - 6.2 | - 6.8 | **- 8.1** | - 7.1 |
| **21** | - 7.4 | - 6.5 | - 6.7 | - 7.1 | - 7.0 | - 6.4 | - 6.9 | - 7.8 | - 6.5 |
| **22** | **- 8.1** | - 7.0 | - 6.9 | - 7.4 | - 7.2 | - 8.0 | - 7.1 | - 7.2 | - 7.1 |
| **23** | - 7.9 | - 6.5 | - 6.7 | - 7.0 | - 6.8 | - 6.4 | - 6.6 | - 7.8 | - 6.9 |
| **24** | - 7.3 | - 6.4 | - 6.9 | - 7.1 | - 6.9 | - 5.7 | - 6.3 | - 6.3 | - 6.4 |
| **25** | - 7.6 | - 6.8 | - 7.0 | - 7.2 | - 7.1 | - 6.5 | - 6.6 | - 7.1 | - 6.7 |
| **26** | - 7.3 | - 6.1 | - 6.7 | - 7.0 | - 7.1 | - 6.3 | - 6.4 | - 8.0 | - 6.2 |
| **27** | **- 8.0** | - 6.0 | - 6.7 | - 7.1 | - 7.2 | - 6.7 | - 6.7 | - 7.5 | - 6.1 |
| **28** | - 7.7 | - 6.3 | - 7.3 | - 7.2 | - 6.9 | - 6.5 | - 6.6 | - 7.8 | - 6.5 |
| **29** | **- 8.4** | - 6.5 | - 6.9 | - 7.3 | - 7.0 | - 6.9 | - 6.7 | - 7.9 | - 6.6 |
| **30** | **- 8.2** | - 6.3 | - 6.7 | - 7.7 | - 7.4 | - 6.6 | - 7.0 | - 8.0 | - 6.4 |
| **31** | **- 8.0** | - 6.7 | - 7.8 | **- 8.2** | - 7.2 | - 7.0 | - 7.5 | - 7.1 | - 7.0 |
| **32** | - 7.6 | - 7.0 | - 7.5 | **- 8.2** | - 6.8 | - 7.9 | - 7.1 | - 7.5 | - 7.4 |
| **33** | - 7.0 | - 6.1 | - 6.7 | - 7.1 | - 7.2 | - 6.3 | - 6.4 | - 7.9 | - 6.3 |
| **34** | - 7.8 | - 6.2 | - 6.5 | - 7.2 | - 7.3 | - 6.5 | - 6.6 | - 7.5 | - 6.5 |
| **35** | - 7.6 | - 6.3 | - 6.4 | - 7.2 | - 7.2 | - 6.4 | - 6.8 | - 7.7 | - 6.1 |

**Table 3:** Binding affinity of the hit compounds toward kinases, proteases, oxidoreductases and erasers as compared to natural and commercial ligands

| Ligands | Kinases |  |  |  |  |  |  |  | Protease | Oxidoreductase |  | Eraser |
| --- | --- | --- | --- | --- | --- | --- | --- | --- | --- | --- | --- | --- |
|  | RAF1 | ROCK1 | ROCK2 | Serine-threonine protein kinase Chk1 | Tyrosine protein kinase FYN | Tyrosine protein kinase LCK | CDK8 | Dual Specificity protein kinase CLK4 | MMP9 | COX-2 | Monoamine oxidase B | deacetylase NAD dependent Sirtuin 2 |
| **Natural** | **- 10.2** | **- 8.2** | **- 10.7** | **- 10.1** | **- 11.2** | **- 12.8** | **- 8.2** | **- 10.3** | **- 11.4** | **- 10.3** | **- 6.9** | **- 12.9** |
| **CL** | Sorafinib<br>**- 10.6** | Y27632<br>**- 7.2** | fasudil<br>**- 8.3** | Rabusertib<br>**- 8.4** | Saracatinib<br>**- 8.9** | Masitinib<br>**- 9.0** | Cortistatin - A<br>**- 9.9** | CID 46926514<br>**- 10.6** | Apigenin<br>**- 10.2** | Celecoxib<br>**- 10.8** | Benmoxin<br>**- 8.9** | 3'-Phenethyl Oxy benzamide<br>**- 13.2** |
| **1** | - 4.6 | - 4.8 | - 5.9 | - 5.7 | - 5.8 | - 4.8 | - 6.3 | - 6.2 | - 6.2 | - 6.1 | - 5.6 | - 5.5 |
| **2** | - 5.8 | - 5.9 | - 6.0 | - 5.6 | - 6.1 | - 5.5 | - 6.1 | - 6.2 | - 6.3 | - 6.2 | - 6.2 | - 6.3 |
| **3** | **- 8.4** | - 6.9 | - 7.3 | - 7.2 | - 7.3 | - 8.0 | - 7.0 | **- 8.2** | **- 9.1** | - 7.4 | **- 8.2** | **- 10.1** |
| **4** | **- 8.1** | - 6.7 | - 7.3 | - 7.4 | - 7.6 | - 7.9 | - 7.5 | **- 8.4** | **- 9.1** | - 7.6 | **- 8.2** | **- 10.6** |
| **5** | - 6.4 | - 7.2 | - 7.2 | - 7.2 | - 7.2 | - 6.9 | - 7.6 | **- 9.2** | - 7.6 | - 7.5 | **- 8.2** | **- 8.1** |
| **6** | - 6.4 | - 6.7 | - 7.4 | - 6.9 | - 7.0 | - 6.7 | - 7.4 | **- 9.1** | - 7.0 | - 7.2 | - 7.7 | **- 8.1** |
| **7** | **- 8.4** | - 7.2 | - 9.0 | **- 10.0** | **- 10.0** | **- 8.6** | **- 9.5** | **- 10.4** | **- 8.3** | **- 9.8** | **- 9.2** | - 5.8 |
| **8** | - 7.4 | - 7.8 | **- 9.0** | **- 8.3** | **- 9.4** | **- 8.3** | **- 8.8** | **- 10.6** | **- 8.2** | **- 9.4** | **- 8.6** | **- 8.5** |
| **9** | **- 9.1** | **- 8.4** | **- 9.4** | **- 9.8** | **- 9.4** | **- 9.8** | **- 9.1** | **- 10.9** | **- 9.5** | **- 9.7** | **- 10.3** | **- 9.5** |
| **10** | - 5.7 | - 5.3 | - 5.5 | - 5.4 | - 5.1 | - 5.3 | - 5.4 | - 6.2 | - 6.4 | - 5.9 | - 5.9 | - 7.0 |
| **11** | - 6.8 | - 6.5 | - 7.0 | - 7.0 | - 6.8 | - 6.7 | - 7.2 | - 7.0 | - 7.7 | - 7.6 | - 7.7 | - 5.2 |
| **12** | - 5.7 | - 7.2 | **- 8.2** | **- 8.0** | **- 8.5** | **- 8.9** | **- 8.4** | **- 8.2** | **- 8.4** | **- 8.3** | **- 8.6** | **- 8.1** |
| **13** | - 5.9 | - 7.1 | **- 8.6** | **- 8.0** | **- 8.3** | **- 8.8** | **- 8.2** | **- 8.8** | **- 8.0** | **- 8.4** | **- 8.4** | **- 8.4** |
| **14** | - 5.9 | - 7.8 | **- 9.7** | **- 8.1** | - 7.5 | **- 8.0** | - 7.1 | **- 9.7** | **- 8.3** | **- 9.3** | **- 8.5** | - 4.5 |
| **15** | - 5.8 | - 7.3 | **- 8.7** | - 7.6 | - 7.7 | **- 8.1** | **- 8.7** | **- 8.9** | **- 8.0** | **- 8.6** | **- 8.3** | - 7.2 |
| **16** | **- 8.6** | **- 8.0** | **- 8.3** | **- 9.4** | **- 9.1** | **- 10.0** | **- 8.4** | **- 9.1** | **- 9.1** | **- 9.3** | **- 9.4** | **- 10.3** |
| **17** | - 6.6 | - 6.3 | - 6.7 | - 7.0 | - 7.2 | - 7.8 | - 7.1 | - 7.7 | - 7.6 | **- 8.0** | **- 8.1** | **- 8.1** |
| **18** | - 7.0 | - 6.5 | - 7.2 | - 7.3 | - 7.3 | **- 8.0** | - 7.1 | - 7.4 | - 7.6 | - 7.9 | **- 8.6** | - 7.9 |
| **19** | - 7.0 | - 6.7 | - 7.1 | - 7.4 | - 7.4 | **- 8.0** | - 7.0 | - 7.4 | - 7.8 | - 7.9 | **- 8.5** | **- 8.5** |
| **20** | - 6.9 | - 6.9 | - 7.0 | - 6.9 | - 7.2 | - 7.8 | - 7.2 | - 7.4 | - 7.9 | - 7.8 | **- 8.3** | **- 8.0** |
| **21** | - 7.3 | - 6.9 | - 7.2 | - 7.5 | - 7.7 | **- 8.2** | - 7.2 | - 7.6 | - 7.9 | **- 8.1** | **- 8.6** | **- 8.8** |
| **22** | - 6.5 | - 7.0 | - 7.0 | **- 8.1** | - 7.5 | **- 8.0** | - 7.9 | - 7.7 | **- 8.4** | **- 8.1** | **- 9.0** | **- 8.6** |
| **23** | - 6.4 | - 6.5 | - 7.0 | - 7.2 | - 6.9 | **- 8.0** | - 7.3 | **- 8.1** | - 7.5 | **- 8.0** | **- 8.5** | **- 8.7** |
| **24** | - 6.5 | - 6.3 | - 7.3 | - 6.8 | - 6.9 | - 7.0 | - 7.3 | - 7.5 | - 6.6 | - 7.3 | - 7.5 | - 6.8 |
| **25** | - 6.7 | - 6.4 | - 7.6 | - 7.3 | - 6.9 | **- 8.1** | - 7.3 | - 7.8 | - 6.8 | - 7.5 | - 7.8 | - 7.4 |
| **26** | - 6.5 | - 6.1 | - 6.8 | - 6.8 | - 7.6 | - 6.7 | - 7.1 | - 7.4 | - 7.3 | **- 8.1** | - 7.6 | - 7.5 |
| **27** | - 7.2 | - 6.3 | - 7.3 | - 7.2 | - 7.6 | - 7.0 | - 7.0 | **- 8.2** | - 7.3 | - 7.9 | **- 8.3** | - 7.4 |
| **28** | - 7.0 | - 6.6 | - 7.2 | - 6.9 | - 7.4 | - 7.4 | - 7.1 | **- 8.3** | - 7.5 | **- 8.1** | - 7.7 | **- 8.2** |
| **29** | - 7.3 | - 6.3 | - 7.7 | - 7.0 | **- 8.3** | - 7.1 | - 7.4 | - 7.6 | - 7.6 | - 7.9 | **- 8.0** | - 5.7 |
| **30** | - 5.5 | - 6.6 | - 7.8 | - 6.5 | **- 8.4** | - 6.3 | - 7.2 | - 7.5 | - 7.7 | - 8.0 | - 8.1 | - 7.9 |
| **31** | - 7.1 | - 6.1 | - 7.7 | **- 8.3** | **- 8.7** | - 7.3 | **- 8.3** | **- 8.5** | **- 8.3** | **- 8.9** | **- 8.8** | **- 9.0** |
| **32** | - 7.9 | - 7.4 | **- 8.4** | - 7.8 | **- 8.5** | - 7.6 | **- 8.3** | **- 8.5** | **- 8.3** | **- 8.6** | **- 8.7** | - 5.8 |
| **33** | - 6.4 | - 6.2 | - 6.5 | - 6.7 | - 7.2 | - 6.8 | - 6.9 | - 7.4 | - 7.1 | - 7.9 | **- 8.0** | - 6.2 |
| **34** | - 5.2 | - 6.2 | - 7.0 | - 7.2 | - 7.2 | - 6.0 | - 7.1 | - 7.0 | - 7.2 | - 7.5 | - 7.8 | - 6.7 |
| **35** | - 7.9 | - 7.4 | **- 8.4** | - 7.8 | **- 8.5** | - 7.6 | **- 8.3** | **- 8.5** | **- 8.3** | **- 8.6** | **- 8.7** | - 5.8 |

The most prominent family of compounds among the identified hits, made of C – C-linked biphenyls (**12 – 35**), displayed interaction with almost the same array of potential targets as dihydroquinoline derivatives. They showed similar interaction with six enzymes (anandamide amidohydrolase, arachidonate 5 lipoxygenase, arachidonate 15 lipoxygenase, nicotinamide phosphoribosyltransferase, PARP and PI3-kinase p110-alpha subunit), three phosphodiesterases (4B, 5A and 10A), eight kinases (RAF1, ROCK1, ROCK2, serine-threonine protein kinase Chk1, tyrosine-protein kinase FYN, tyrosine protein kinase LCK, CDK8, and dual-specificity protein kinase CLK4), one protease (MMP9), two oxidoreductases (COX-2 and monoamine oxidase B), and one eraser (deacetylase NAD dependent Sirtuin 2). Like dihydroquinoline derivatives, compounds (**12 – 35**) consistently showed a favorable binding coefficient with the nuclear receptor ROR-gamma. These compounds also exhibited some interactions with the cholesterol ester transfer protein, the cannabinoid receptor 2, a primary active transporter (P-glycoprotein 1), the orphan nuclear receptor LRH-1, and the nuclear receptor ROR-gamma. However, unlike the dihydroquinoline derivatives, they showed no interactions with the peroxisome-proliferator activated receptor-gamma or with MDM2. It should be mentioned that many of these enzymes, kinases, proteases, transporters, and receptors have been shown to play essential roles in skin carcinogenesis, and cancer progression. A decrease in anandamide signaling was shown as a potential contributor to the maintenance of cutaneous mechanical hyperalgesia in a model of bone cancer pain.^62–64^ Taken together with the fact that pharmacological activation of cannabinoid receptors appeared to induce the apoptotic death of tumorigenic epidermal cells without affecting the viability of non-transformed epidermal cells,^65^ these studies suggest that the manipulation of peripheral endocannabinoid signaling can be a promising strategy for the management of some types of cancers.

Moreover, lipoxygenases are well known to catalyze the formation of corresponding hydroperoxides from polyunsaturated fatty acids such as linoleic acid and arachidonic acid, and they are expressed in immune, epithelial, and tumor cells associated with other physiological conditions, such as skin inflammation, and tumorigenesis.^66, 67^ Many kinases, including RAF1,^68, 69^ ROCK1 and 2,^70, 71^ serine-threonine protein kinase (CHK1),^72, 73^ tyrosine-protein kinase (FYN),^74, 75^ tyrosine protein kinase (LCK),^76^ and dual specificity protein kinase (CLK4) are also well known to play critical roles in skin tumorigenesis as well as in other cancer types. In fact, SKMEL-28 (one of a series of melanoma cell lines established from patient-derived tumor samples) and A375, both used in this study, are well-known to harbor BRAF V600E driver mutation and wildtype N-RAS.^77^ On the other hand, nicotinamide adenine dinucleotide (NAD)-biosynthetic enzyme (nicotinamide phosphoribosyltransferase also known as NAMPT) was shown to be a driving factor in resistance associated with serine-threonine protein kinase B-RAF (BRAF)-mutated metastatic melanoma, one of the highly aggressive types of skin cancer.^78, 79^

Further studies revealed that NAMPT over-expression is necessary and sufficient to recapitulate the BRAFi-resistant phenotype plasticity.^78^ It should also be mentioned that PI3K/RAC alpha serine/threonine protein kinase (AKT)/mTOR signaling pathway has been shown to be over-activated in some types of skin cancers,^80^ with the anti-proliferative activity of some phenolic compounds against SK-MEL-2 cancer cells attributed to the downregulation of the PI3K/AKT/mTOR signaling pathway, which induces a mitochondrial-dependent apoptosis.^81^

Other potential targets revealed during the *in silico* “target fishing” reverse screening, including Poly(ADP-ribose)polymerase (PARP),^82, 83^ phosphodiesterase type 5,^84, 85^ orphan nuclear receptor LRH-1,^86, 87^ nuclear receptor ROR-gamma,^88, 89^ matrix metallopeptidase 9 (MMP9),^90^ NAD-dependent deacetylase Sirtuin 2,^91^ and cholesterol ester transfer protein,^92^ all believed to play important roles in skin carcinogenesis. There is no doubt that the wealth of information gathered here will be useful in further studies towards the optimizing the hit molecules’ activity and determining their putative drug target and potential mechanism(s) of action.

Finally, the drug-likeness properties of these compounds were evaluated *in silico*, using the SwissADME web tool.^58, 93^ **Table 4** displays some physicochemical properties, ADME parameters, and violations of drug-likeness rules for these compounds. These properties were computed exploiting various filters widely used to evaluate the drug-likeness of a wide range of compounds. This includes the Lipinski (Pfizer) filter (MW ≤ 500; MLOGP ≤ 4.15; HBA ≤ 10; HBD ≤ 5),^94^ the Ghose filter (160 ≤ MW ≤ 480; -0.4 ≤ WLOGP ≤ 5.6; 40 ≤ MR ≤130; 20 ≤ atoms ≤ 70),^95^ Veber (GSK) filter (RB ≤ 10; TPSA ≤ 140),^96^ Egan (Pharmacia) filter (WLOGP ≤ 5.88; TPSA ≤ 131.6),^97^ and Muegge (Bayer) filter (200 ≤ MW ≤ 600; -2 ≤ XLOGP ≤ 5; TPSA ≤ 157; HBA ≤ 10; HBD ≤ 5; RB ≤ 15; number of rings ≤ 7; number of carbons > 4; number of heteroatoms >1).^98^

**Table 4:** Physicochemical and pharmacokinetic properties of the hit compounds

| Comp. | MW<br>(g/mol) | Lipophilicity |  |  |  |  |  | Drug-likeness<br>(number of violations) |  |  |  |  | Water Solubility |  | Pharmacokinetics |  |
| --- | --- | --- | --- | --- | --- | --- | --- | --- | --- | --- | --- | --- | --- | --- | --- | --- |
| | | ilogP | XlogP3 | WlogP | MlogP | Silicos-IT logP | Consensus<br>logP | Lipinski | Ghose | Veber | Egan | Muegge | ESOL | Class | Log $K_p$<br>(cm/s) | F |
| **1** | 384.018 | 3.18 | 3.07 | 3.02 | 2.17 | 3.23 | 2.93 | 0 | 0 | 0 | 0 | 0 | -4.07 | moderately | -6.46 | 0.55 |
| **2** | 271.223 | 2.23 | 1.52 | 1.41 | -0.11 | -0.26 | 0.96 | 0 | 0 | 0 | 0 | 0 | -2.32 | easily | -6.88 | 0.55 |
| **3** | 201.307 | 3.00 | 4.65 | 3.71 | 3.38 | 3.23 | 3.60 | 0 | 0 | 0 | 0 | 1 | -4.18 | moderately | -4.23 | 0.55 |
| **4** | 235.752 | 3.21 | 4.62 | 4.54 | 3.91 | 4.29 | 4.11 | 0 | 0 | 0 | 0 | 1 | -4.29 | moderately | -4.46 | 0.55 |
| **5** | 217.307 | 3.01 | 3.11 | 3.12 | 2.73 | 3.32 | 3.06 | 0 | 0 | 0 | 0 | 0 | -3.29 | easily | -5.42 | 0.55 |
| **6** | 203.280 | 2.72 | 2.74 | 2.73 | 2.46 | 2.96 | 2.72 | 0 | 0 | 0 | 0 | 0 | -3.06 | easily | -5.59 | 0.55 |
| **7** | 335.482 | 4.20 | 4.78 | 5.76 | 4.74 | 5.82 | 5.06 | 1 | 1 | 0 | 0 | 0 | -4.91 | moderately | -4.95 | 0.55 |
| **8** | 297.434 | 4.04 | 4.83 | 5.08 | 4.09 | 5.09 | 4.62 | 0 | 0 | 0 | 0 | 0 | -4.73 | moderately | -4.69 | 0.55 |
| **9** | 305.456 | 4.21 | 5.13 | 5.86 | 5.10 | 5.89 | 5.24 | 1 | 1 | 0 | 0 | 2 | -5.03 | moderately | -4.52 | 0.55 |
| **10** | 151.206 | 2.24 | 1.60 | 1.94 | 1.80 | 1.83 | 1.88 | 0 | 1 | 0 | 0 | 1 | -1.99 | very | -6.09 | 0.55 |
| **11** | 300.395 | 3.56 | 4.26 | 4.10 | 3.01 | 3.17 | 3.62 | 0 | 0 | 0 | 0 | 0 | -4.33 | moderately | -5.11 | 0.55 |
| **12** | 277.360 | 3.18 | 4.93 | 4.59 | 3.97 | 4.20 | 4.17 | 0 | 0 | 0 | 0 | 0 | -5.10 | moderately | -4.49 | 0.55 |
| **13** | 307.386 | 3.43 | 4.90 | 4.68 | 3.59 | 4.11 | 4.14 | 0 | 0 | 0 | 0 | 0 | -5.08 | moderately | -4.70 | 0.55 |
| **14** | 383.482 | 4.16 | 6.40 | 6.10 | 4.65 | 5.59 | 5.38 | 1 | 1 | 0 | 1 | 1 | -6.41 | poorly | -4.10 | 0.55 |
| **15** | 347.450 | 4.14 | 5.78 | 5.62 | 4.17 | 5.11 | 4.96 | 1 | 1 | 0 | 0 | 1 | -5.69 | moderately | -4.32 | 0.55 |
| **16** | 277.360 | 3.27 | 4.93 | 4.59 | 3.97 | 4.20 | 4.19 | 0 | 0 | 0 | 0 | 0 | -5.10 | moderately | -4.49 | 0.55 |
| **17** | 257.328 | 3.17 | 3.65 | 3.52 | 2.84 | 3.03 | 3.24 | 0 | 0 | 0 | 0 | 0 | -3.94 | easily | -5.28 | 0.55 |
| **18** | 336.224 | 3.62 | 4.34 | 4.29 | 3.47 | 3.70 | 3.88 | 0 | 0 | 0 | 0 | 0 | -4.84 | moderately | -5.27 | 0.55 |
| **19** | 291.773 | 3.51 | 4.28 | 4.18 | 3.35 | 3.66 | 3.88 | 0 | 0 | 0 | 0 | 0 | -4.53 | moderately | -5.04 | 0.55 |
| **20** | 287.354 | 3.68 | 3.63 | 3.53 | 2.50 | 3.08 | 3.28 | 0 | 0 | 0 | 0 | 0 | -4.00 | moderately | -5.48 | 0.55 |
| **21** | 271.354 | 3.35 | 4.02 | 3.83 | 3.09 | 3.54 | 3.57 | 0 | 0 | 0 | 0 | 0 | -4.23 | moderately | -5.10 | 0.55 |
| **22** | 299.407 | 3.70 | 4.78 | 4.65 | 3.56 | 4.15 | 4.17 | 0 | 0 | 0 | 0 | 0 | -4.78 | moderately | -4.73 | 0.55 |
| **23** | 291.773 | 3.53 | 4.28 | 4.18 | 3.35 | 3.66 | 3.80 | 0 | 0 | 0 | 0 | 0 | -4.53 | moderately | -5.04 | 0.55 |
| **24** | 326.218 | 3.30 | 4.77 | 4.83 | 3.86 | 4.31 | 4.21 | 0 | 0 | 0 | 0 | 0 | -5.03 | moderately | -4.90 | 0.55 |
| **25** | 285.381 | 3.43 | 4.38 | 4.14 | 3.32 | 4.06 | 3.87 | 0 | 0 | 0 | 0 | 0 | -4.53 | moderately | -4.93 | 0.55 |
| **26** | 285.381 | 3.75 | 4.55 | 4.30 | 3.32 | 3.80 | 3.95 | 0 | 0 | 0 | 0 | 0 | -4.50 | moderately | -4.81 | 0.55 |
| **27** | 364.277 | 3.41 | 5.24 | 5.07 | 3.94 | 4.48 | 4.43 | 0 | 0 | 0 | 0 | 1 | -5.41 | moderately | -4.80 | 0.55 |
| **28** | 299.407 | 3.56 | 4.98 | 4.55 | 3.56 | 4.02 | 4.13 | 0 | 0 | 0 | 0 | 0 | -4.84 | moderately | -4.59 | 0.55 |
| **29** | 378.303 | 4.16 | 5.68 | 5.31 | 4.16 | 4.70 | 4.80 | 0 | 0 | 0 | 0 | 1 | -5.75 | moderately | -4.57 | 0.55 |
| **30** | 364.277 | 3.84 | 5.14 | 5.07 | 3.94 | 4.31 | 4.46 | 0 | 0 | 0 | 0 | 1 | -5.41 | moderately | -4.87 | 0.55 |
| **31** | 333.424 | 3.70 | 5.15 | 4.94 | 3.99 | 4.54 | 4.46 | 0 | 0 | 0 | 0 | 1 | -5.29 | moderately | -4.68 | 0.55 |
| **32** | 412.320 | 3.92 | 5.84 | 5.70 | 4.58 | 5.22 | 5.05 | 1 | 1 | 0 | 0 | 1 | -6.19 | poorly | -4.67 | 0.55 |
| **33** | 299.407 | 3.99 | 4.91 | 4.69 | 3.56 | 4.19 | 4.27 | 0 | 0 | 0 | 0 | 0 | -4.73 | moderately | -4.64 | 0.55 |
| **34** | 378.303 | 4.08 | 5.61 | 5.46 | 4.16 | 4.87 | 4.84 | 0 | 0 | 0 | 0 | 1 | -5.64 | moderately | -4.62 | 0.55 |
| **35** | 364.277 | 3.75 | 5.08 | 5.07 | 3.94 | 4.48 | 4.46 | 0 | 0 | 0 | 0 | 1 | -5.31 | moderately | -4.92 | 0.55 |

Molecular weight (MW), LogP values (an indicator of lipophilicity), aqueous solubility parameters (ESOL), an indicator of skin permeation (Log $K_p$ ), and bioavailability score (F)

The obtained results indicate that the drug-likeness of all these hit compounds are well within the defined parameters for a good drug candidate. More importantly, the bioavailability score of about 0.55 obtained for all these compounds satisfies Lipinski’s rule for a neutral compound for an optimal absorption through oral ingestion,^94^ suggesting that these compounds could be good oral drug candidates, in case they were to display useable anticancer properties. The synthetic accessibility scores for these compounds were also very low, indicating ease to readily prepare derivatives for further structure-activity relationship studies. Furthermore, as indicated in the chemistry section, all these compounds were synthesized by our research group through simple and straightforward methods that can be easily adapted for the preparation of the more potent analogs.

### 2.4. Preliminary investigation of the potential antiproliferative mechanism of action

In order to explore the potential antiproliferative mechanism of action of these molecules, the effects of the two most active compounds (**11** and **13**) on wound healing, colony formation and apoptosis-related properties were further investigated. A scratch wound assay was used to scrutinize the ability of these compounds to modulate the migration of SCC-12 and SKMEL-28 cells into cell-free areas produced by scratch-wounding after 48 h of incubation. Both compounds significantly decreased the scratched wound areas of cultured SCC-12 and SKMEL-28 cells after a 48 h treatment relative to the control untreated cells, in a dose-dependent manner, as illustrated in **Figure 2**. The long-term impacts of different concentrations (0, ½IC_50_, IC_50,_ and 1½IC_50_) of compounds **11** and **13** on colony formation potentials for SCC-12 and SKMEL-28 cells compared to controls were then explored. As depicted in **Figure 3A-B**, these compounds reduced the number of colonies formed in both SCC-12 and SKMEL-28 cells in a concentration-dependent manner. Further analysis revealed that these dose-dependent reductions in clonogenicity were significant in both cell lines, when compared to untreated control groups (**Figure 3C-F**).

**Figure 2.**
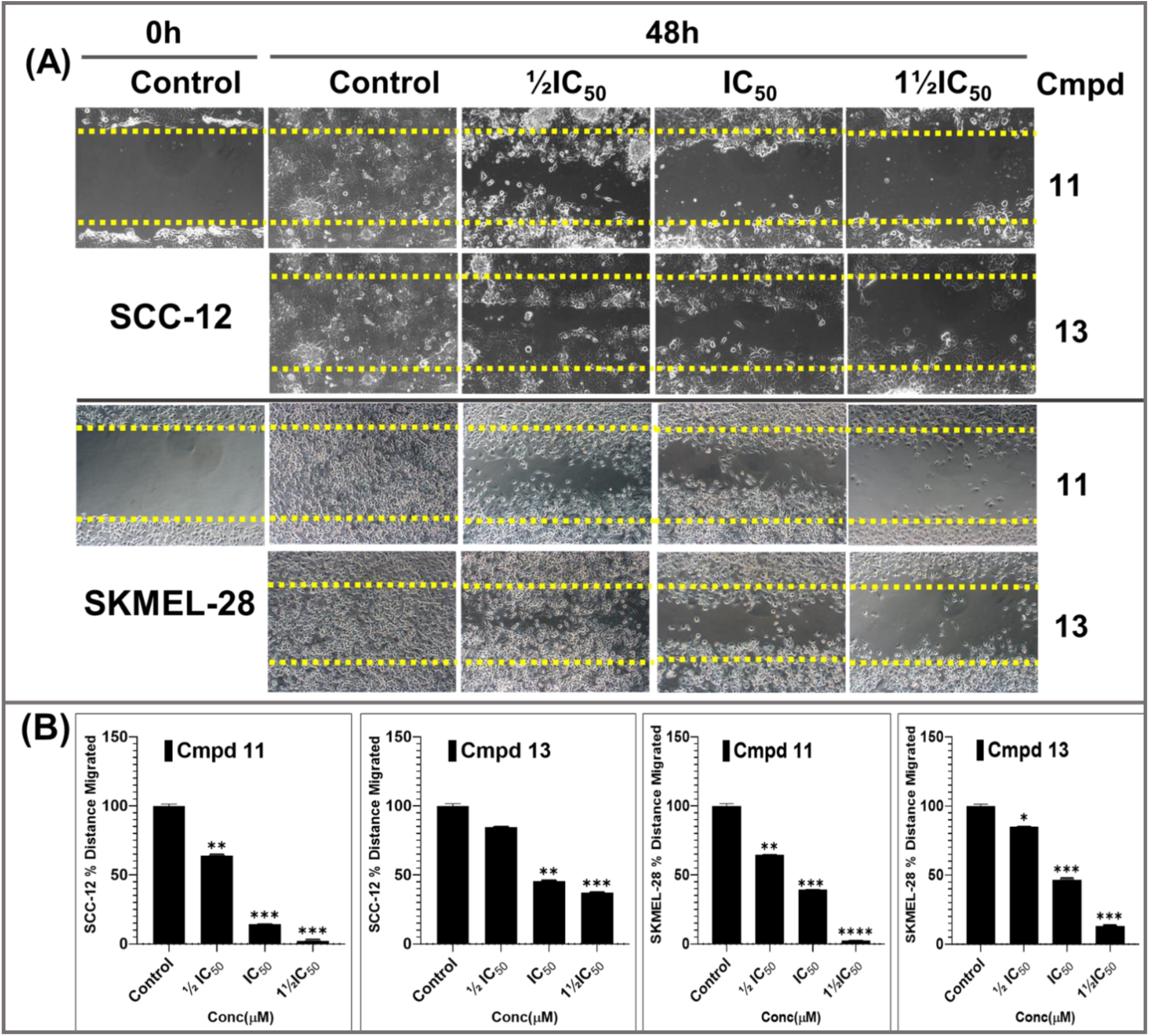
**The potent hit compounds 11 and 13, dose-dependently (0, ½IC_50_, IC_50_ and 1½IC_50_; µM), inhibited the *in-vitro* migration of SCC-12 (A; top panel) and SKMEL-28 (A; bottom panel) cells into the cell-free wounded areas of a confluent cell monolayer scratched wound.** Compounds **11** and **13** induced suppressions of wound closure of SCC-12 and SKMEL-28 cells were significant (**B**). Data show the evidential effect of test compounds in limiting the migration of cells in the inflicted *in-vitro* wounds healing. The bar graphs (**B**) represent the mean ± SD of scratched-wound area values after 48 h and are expressed as a percentage at 0 h untreated control from three independent experiments conducted in triplicates. Statistical significance was assessed using one-way ANOVA and Dunn’s multiple comparison tests, p < 0.05 (*), p < 0.01 (**), p < 0.001 (***) and p < 0.0001 (****), were considered significant.

**Figure 3.**
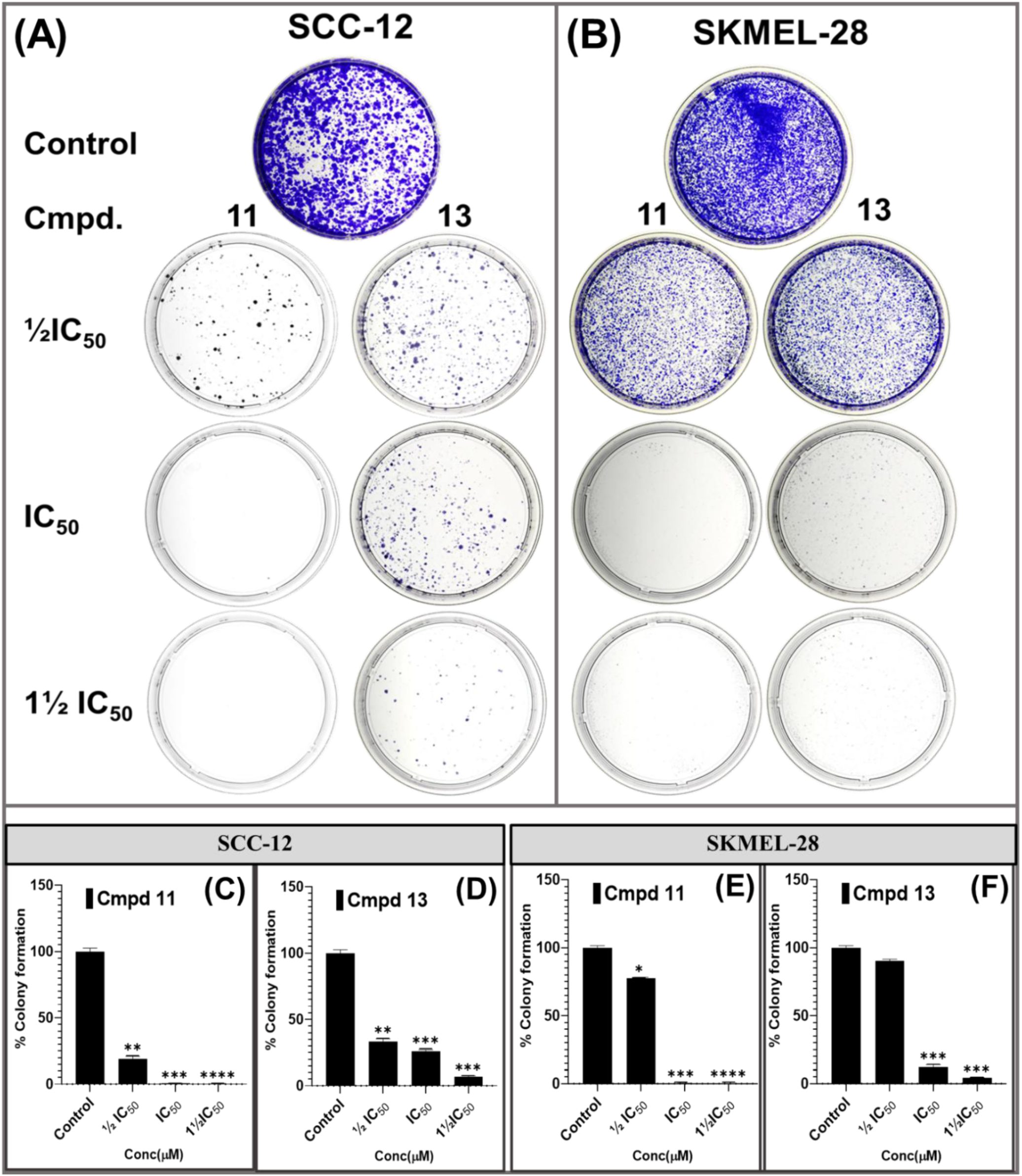
**The potent phenolic compounds 11 and 13 significantly inhibit colony formation potentials in SCC-12 (A) and SKMEL-28 (B) cells cultured in monolayers after 14 days when control cells became confluent.** The percentage decrease in colony formation was in a dose-dependent (0, ½IC_50_, IC_50_ and 1½IC_50_; µM), and were comparable in SCC-12 (C-D) and SK-MEL-28 (E-F) cells. The data expressed in the bar graphs represent the mean ± SD of of colonies analyzed in compounds **11** and **13** treated groups expressed as a percentage of the untreated control group. Data are from three independent experiments, each performed in quadruplicate. Statistical significance was assessed using one-way ANOVA and Dunn’s multiple comparison tests, p < 0.05 (*), p < 0.01 (**), p < 0.001 (***) and p < 0.0001 (****), were considered significant.

Since **11** and **13** have proven to induce significant decreases in cell viability, it was worthwhile investigating the molecular mechanisms by which these compounds induce cytotoxicity by assessing their potential to modulate apoptosis in SCC-12 and SKMEL-28 cancer cell lines selectively. As such, the effects of the test compounds on cellular cytoskeleton (actin staining) and nuclear morphology (DAPI staining) **Figure 4A-B** (control; c and ca, and drug-treated; g-k and ga-ka), as well as on the activation of proapoptotic caspase-3 were assessed by comparing the apoptotic cells and caspase-3 cleavage to untreated groups via an immunofluorescent microscopy analysis. It was observed that the nuclei of SCC-12 and SKMEL-28 cells treated with compounds **11** and **13** exhibited the morphologies characteristic of fragmented chromatin with punctate apoptotic nuclei in a dose-dependent manner (data not shown). Compared to untreated SCC-12 cells (**Figure 4A**(a) and **4B**(aa)), which presented a well-organized actin filament cytoskeleton radiating across the cell cytoskeleton and extending to the lamilipods, the drug-treated cells showed disoriented and diminished actin filament cytoskeleton. These effects were dose-dependent for both compound **11** (**Figure 4A**(e-i), red) and compound **13** (**Figure 4B**(ea-ia), red) in the SCC-12 cell line. As shown in **Figure 5**, similar potent effects were observed for **11** (**Figure 5A**(e-i), red) and **13** (**Figure 5B**(ea-ia), red) in the SK-MEL-28 cell line. With regards to the activation of proapoptotic caspase-3, untreated SCC-12 cells showed a regular staining pattern of procaspase-3 (**Figure 4A**(b) and **4B**(ba), green). However, treatment with the indicated hit drugs dose-dependently activated caspase-3 and induced its translocation into the nucleus, as evidenced by the punctated active caspase-3 immunofluorescence staining detected in the nuclei as can be observed in cells treated with compounds **11** (**Figure 4A**(f and j) and **13** (**Figure 4B**(fa and ja), the punctated caspase-3 staining was colocalized with the nuclear actin staining, as illustrated in **Figure 4A** (h and i) for **11** and **Figure 4B** (ha-ia), cyan) for **13**. Compounds **11** (**Figure 5A**(f and j)) and **13** (**Figure 5B**(fa and ja)) also exhibited similar dose-dependent effects on the actin cytoskeleton as well as the caspase-3 activation status, which also served for the localization (cyan yellow) of the actin (red), nucleus (DAPI) and pro and active caspase-3 immunostaining (green) (see **Figure 4 and 5**).

**Figure 4.**
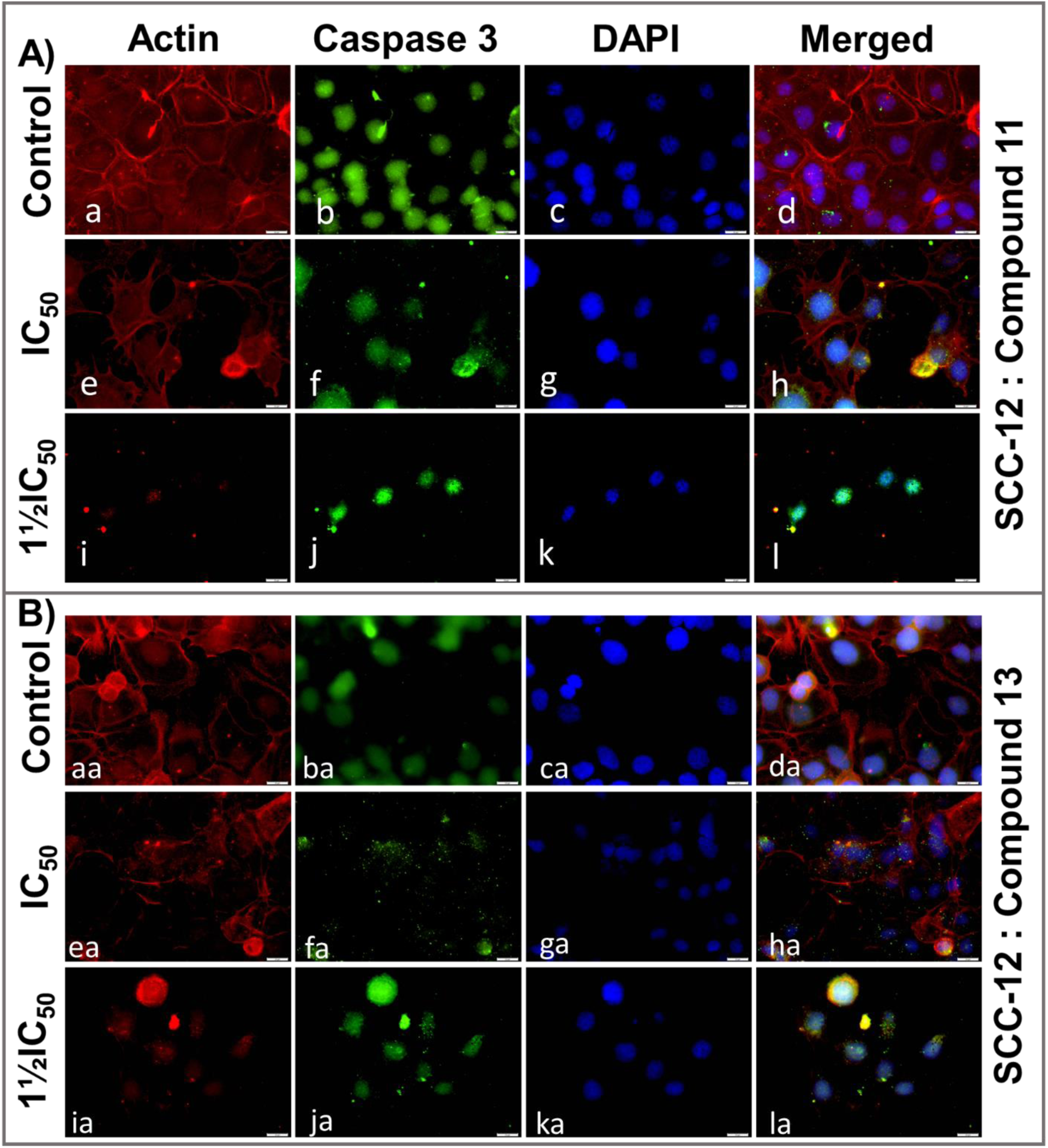
**Compounds 11 and 13 dose-dependently alter the actin cytoskeleton cell morphology and activate pro-apoptotic caspase-3 expression into the nucleus of the SCC-12 non-melanoma skin cancer cell line.** Immunofluorescent micrographs analyses indicate that compounds **11** (A; e-l) and **13** (B; ea-la) compared to untreated controls; for **11** (a-d) and for **13** (aa-da) morphologically distorted the actin cytoskeleton as well as decreased the actin expression levels (red) in monolayer cultures of SCC-12 in a dose-dependent (0, ½IC_50_, IC_50_ and 1½IC_50_; µM), as shown (red; a, e, i and aa, ae, ai). Similarly, the effect of **11**(A) and **13**(B) in activation (green; b,f,j and ba,fa,ja) and translocation of pro-Caspase 3(green) into the nucleus (DAPI, blue) after treatment (colocalization, cyan) as shown (c, g, k and ca, ga, ka). Photomicrographs delineate the colocalization of actin and active caspase-3 in the nucleus (see h, I and ha, la) upon compounds **11** and **13** treatments respectively. Images were obtained after 48 h of incubation SCC-12 cells with or without test compound and analyzed as in the materials and methods section.

**Figure 5.**
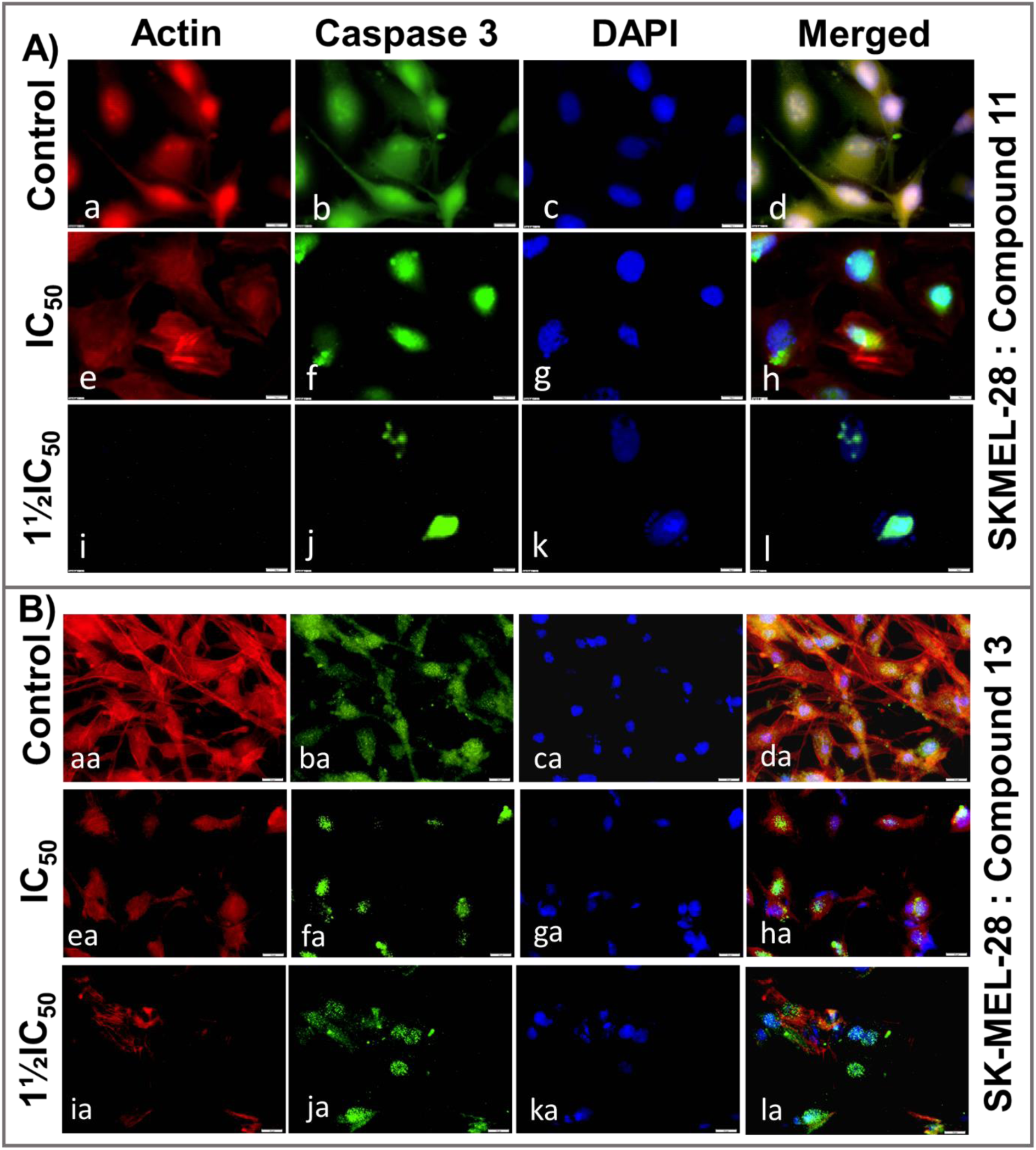
**Compounds 11 and 13 dose-dependently alter the actin cytoskeleton cell morphology and activate pro-apoptotic caspase-3 expression into the nucleus of SKMEL-28 cutaneous melanoma cancer cell line.** Immunofluorescent micrographs analyses indicate that compound **11** (A; e-l) and **13** (B; ea-la) compared to untreated controls; for **11** (a-d) and for **13** (aa-da) morphologically distorted the actin cytoskeleton as well as decreased the actin expression levels (red) in monolayer cultures of SKMEL-28 in a dose-dependent manner (0, ½IC_50_, IC_50_ and 1½IC_50_; µM), as shown (red; a, e, i and aa, ae, ai). Similarly, the effect of **11**(A) and **13**(B) in activation (green; b, f, j and ba, fa, ja) and translocation of pro-caspase 3 into the nucleus (DAPI, blue) after treatment (colocalization, cyan yellow) as shown (c, g, k and ca, ga, ka). Photomicrographs delineate the colocalization of actin and active caspase-3 (green) in the nucleus (see h, I and ha, la) upon compounds **11** and **13** treatments, respectively. Images were obtained after 48 h of incubation of SKMEL-28 cells with or without test compounds and analyzed as described in the materials and methods section.

The cysteinyl aspartate-specific protease family of apoptotic caspases, classified as initiators (caspases 8, 9, and 10) and executioners (caspases 3, 6, and 7), are often targeted for anticancer treatment, including skin tumor treatment.^99, 100^ Using both the immunofluorescence microscopy (**Figure 4 and 5**) and Western blot analyses (**Figure 6 and 7**) of SCC-12 and SKMEL-28 skin cancer cell lines, the modulatory effects of compounds **11** and **13** treatments on the intrinsic and extrinsic apoptosis pathways were investigated by analyzing the protein expression levels of several key apoptosis-related markers. The assessed markers included an executioner pro- and cleaved (activated) forms of caspase -3 (by IF see above **Figure 4 and 5**; green), and an initiator pro- and cleaved caspase-9 as well as activation of PARP cleavage, evaluated by western blot analysis. In addition to the elevated protein expression levels of cleaved caspases-3 observed by immunofluorescence (**Figure 4 and 5**), the treatment of SCC-12 and SKMEL-28 cells with **11** and **13** resulted in significant activation of apoptosis, as evidenced by the increased activation of caspase-9, and PARP, as assessed by western blotting (**Figure 6**). The expression levels of cleaved caspase-9 and cleaved PARP were increased in a concentration-dependent manner compared to untreated controls.

**Figure 6.**
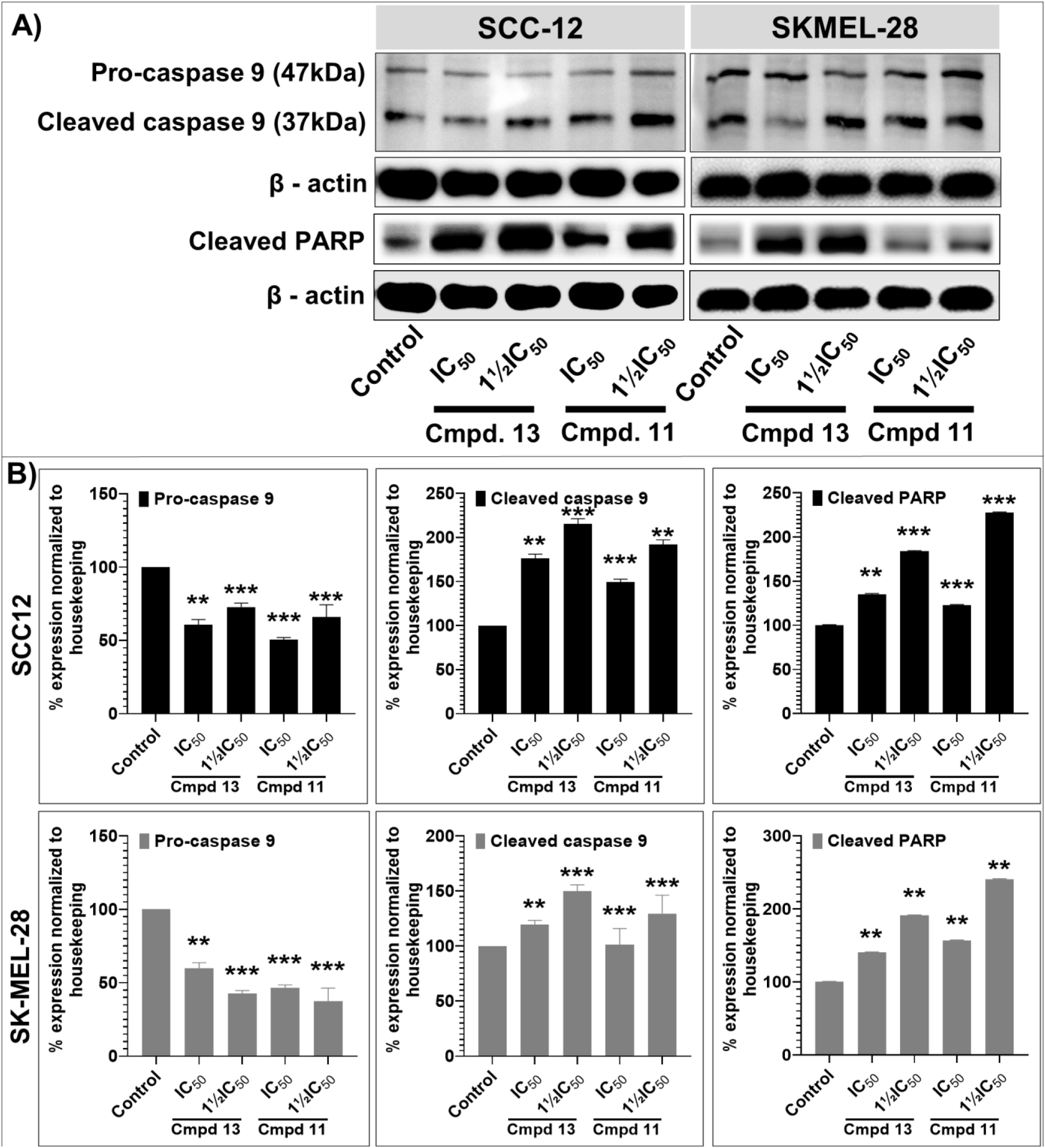
**Compounds 11 and 13 induce apoptosis by activating the extrinsic and intrinsic apoptotic pathway caspases in cutaneous melanoma (SKMel-28) and non-melanoma cancer cells (SCC-12). A)** The blots show a dose-dependent effect (0, ½IC_50_, IC_50_ and 1½IC_50_; µM), of protein expression levels of markers of apoptosis, including pro- and-cleaved caspase 9, and PARP after 48 h of treatment. **B)** The data shown are representative immunoblots from three independent experiments with similar results. β-actin was used as a loading control to confirm the loading uniformity. The actual protein levels were normalized and expressed as a percentage of the loading control (mean ± SD of relative quantitative density values are plotted in the bar graphs). Statistical significance was assessed using one-way ANOVA and Bonnferoni’s multiple comparison tests, *p < 0.01* (**), *p < 0.001* (***) and *p < 0.0001* (****), were considered significant.

**Figure 7.**
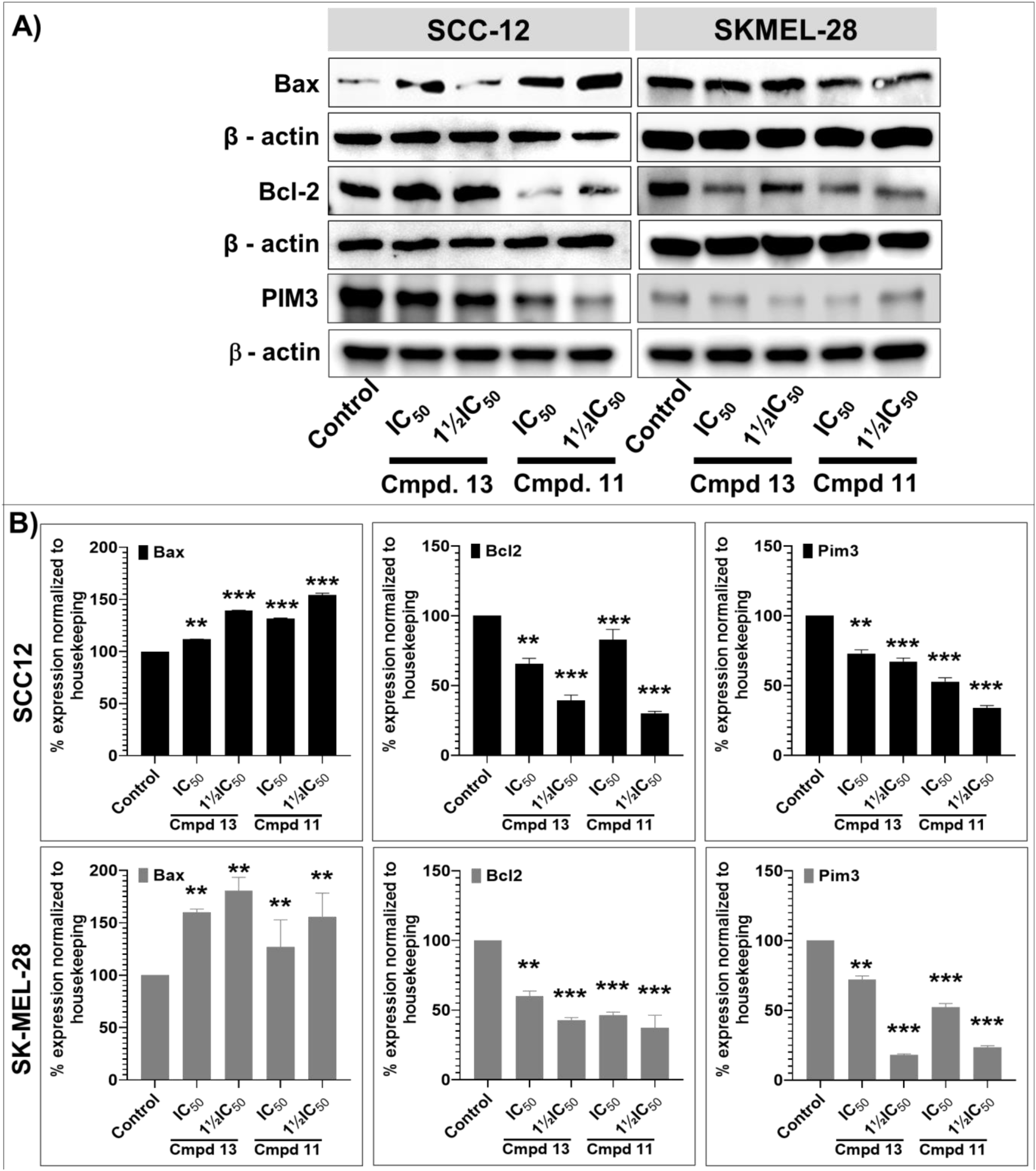
**Compounds 11 and 13 induce apoptosis by activating the extrinsic and intrinsic apoptotic pathway in cutaneous melanoma (SKMel-28) and non-melanoma cancer cells (SCC-12). A)** The blots show a dose-dependent effect (0, ½IC_50_, IC_50_ and 1½IC_50_; µM), of protein expression levels of markers of apoptosis, including Bax, Bcl-2 and PIM3 after 48 h of treatment. **B)** The data shown are representative immunoblots from three independent experiments with similar results. β-actin was used as a loading control to confirm the loading uniformity. The actual protein levels were normalized and expressed as a percentage of the loading control (mean ± SD of relative quantitative density values are plotted in the bar graphs). Statistical significance was assessed using one-way ANOVA and Bonnferoni’s multiple comparison tests, *p < 0.01 (**) and p < 0.001 (***),* were considered significant.

Furthermore, the expression levels of important Bcl-2-family of proteins that regulate the induction of mitochondrial pathway of apoptosis, specifically Bax (pro-apoptotic), and Bcl-2 (anti-apoptotic),^101^ and PIM3, another apoptosis-related marker, were additionally studied in compounds **11** and **13** treated SCC-12 and SKMEL-28 skin cancer cell lines and compared with untreated controls.

As shown in **Figure 7**, the treatment of both SCC-12 and SKMEL-28 cells with **11** and **13** significantly and dose-dependently increased the protein expression level of Bax while decreasing the expression levels of Bcl-2 and PIM3 compared to the untreated controls. Treatment with each of the two compounds resulted in a concentration-dependent increase in the Bax/Bcl-2 ratio (see **Figure 7**), supporting that **11** and **13** can induce the apoptosis of skin cancer cells, as observed herein in SCC-12 and SKMEL-28 skin cancer cell lines, through the intrinsic mitochondrial apoptotic pathway, leading to the activation of PARP.

As described above, uncontrolled cell proliferation and aberrant differentiation, which are the hallmarks of cancers^102^ including melanoma and non-melanoma skin cancers^103, 104^ are often associated with dysregulated signaling pathways, such as the MAPK/ERK and PI3K/AKT pathways,^103, 104^ which have been therapeutically targeted in several advanced melanoma clinical trials using a wide range of inhibitors.^103^ In this study, the treatment of SCC-12 and SKMEL-28 with escalating concentrations (0, IC_50_ and 1½IC_50_) of **11** and **13** for 48 h resulted in a significant reduction in the protein levels of activated phosphorylated AKT, MAPK and mTOR targets when compared to the control group as indicated by the Western blot analysis. The protein expression levels of phosphorylated p90RSK (Ser^380^), Akt (Ser^473^), p44/42-MAPK (ERK1/2; Thr^202^/Try^20^ and ribosomal protein S6(Ser^235/236^) pathways targets were significantly suppressed in SCC-12 and SKMEL-28 cells treated with compound **11** and **13**, compared to the untreated controls as observed by Western blotting (**Figure 8**).

**Figure 8.**
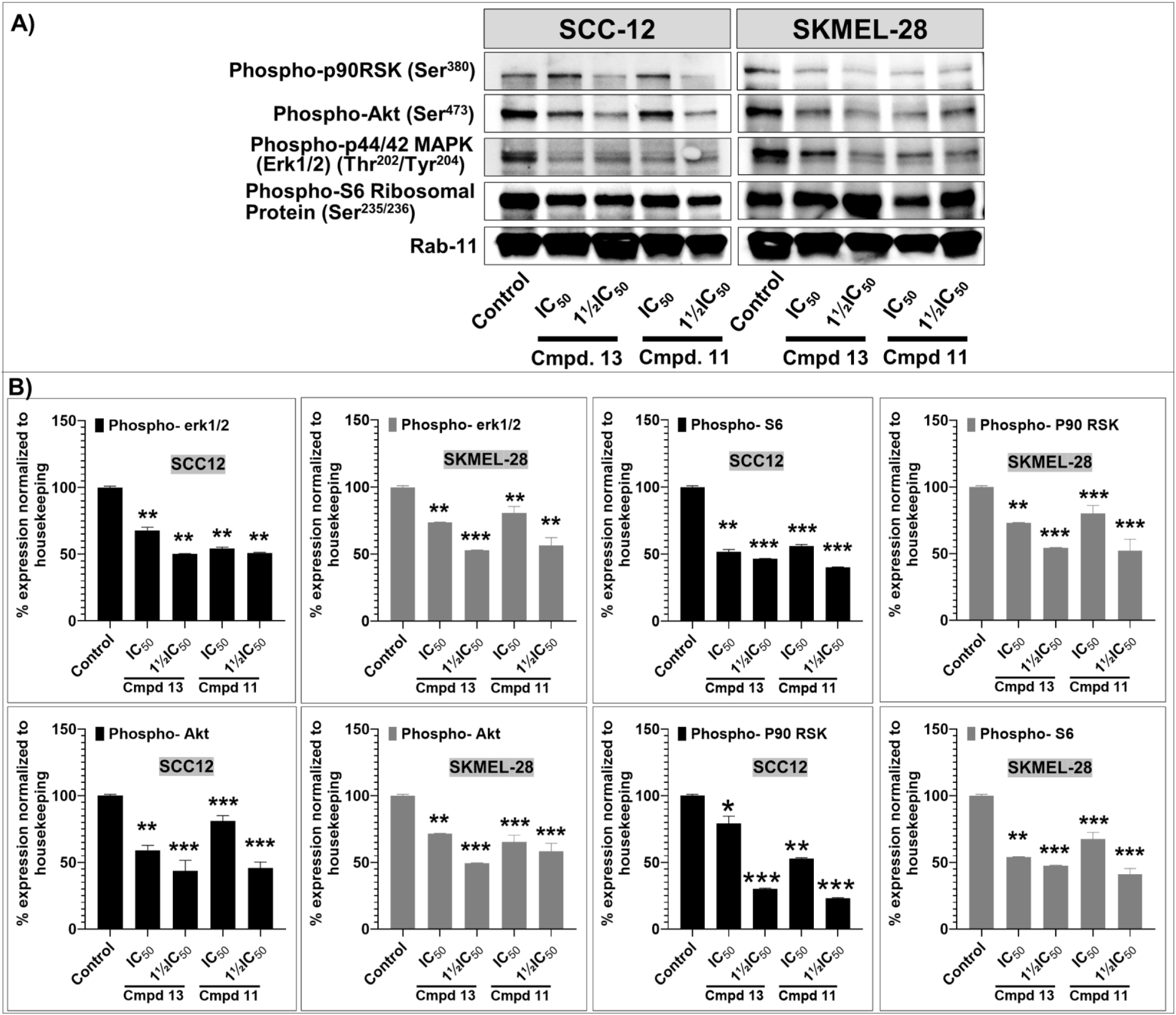
**Compounds 11 and 13 inhibits the expression of markers of the AKT/ mTOR and MAPK pathway, including p90RSK, phosphorylated AKT, Phospho-p44/42 MAPK (Erk1/2) (Thr^202^/Tyr^204^), and Phospho-S6 Ribosomal Protein (Ser^235/236^) in cutaneous melanoma (SKMel-28) and non-melanoma skin cancer cells (SCC-12). A)** The blots show a dose-dependent effect (0, IC_50_ and 1½IC_50_; µM) of protein expression levels after 48 h of treatment. **B)** The data shown are representative immunoblots from three independent experiments with similar results. Rab-11 was used as loading control to confirm the loading uniformity. To obtain the bar graphs, the actual protein levels were normalized and expressed as a percentage of the loading control to obtain the bar graphs (mean ± SD of relative quantitative density values are plotted). Statistical significance was assessed using one-way ANOVA and Dunn’s multiple comparison tests, *p < 0.01 (**) and p < 0.001 (***),* were considered significant.

## 3. Conclusion

Skin cancer is one of the most prevalent types of cancer in the United States, with approximately 9,500 people diagnosed with skin cancer every day.^105–107^ It is no secret that currently available treatments are hampered by drug resistance, side effects, low bioavailability, and high cost. Toward a goal of introducing new small molecule scaffolds into the skin cancer treatment pipeline, our screening efforts of a small library of over 90 compounds based on seven different skeletal frameworks have enabled the identification of 35 lead compounds with varying degrees of activities against melanoma (A375 and SKMEL-28) and non-melanoma (squamous carcinoma A431 and SCC-12) skin cancer cell lines. Many of these hit compounds displayed a useable anticancer activity, primarily against the squamous carcinoma A431 and SCC-12 cell lines, while exhibiting a superior safety profile than cisplatin (a well-known anticancer agent), used as a positive control, as indicated by their lack of activity against mammalian nontumorigenic immortalized HaCaT cells. The most promising compounds (**11** and **13**) were remarkably effective in inhibiting cell proliferation, inducing apoptosis, and suppressing cell migration and invasion *in vitro* in SK-MEL-28 and SCC-12 skin cancer cell lines. Data obtained from anti-proliferative/cytotoxic, scratch-wound healing, and colony formation assays, together with altered levels of aggregation of protein markers of disease, indicate that compounds **11** and **13** inhibit the growth, migration and proliferation of cutaneous carcinoma cells. The results presented in this initial report suggest that compounds **11** and **13** modulate molecular signaling pathway targets involved in apoptosis and cancer progression. Furthermore, since many of the lead compounds uncovered during this study exhibited satisfactory pharmacokinetic and ADMET properties, this work provides a good framework toward a target-based design and the systematic synthesis of safer and more effective analogs for *in vivo* preclinical validation and mechanistic studies, including the analysis of differentially expressed proteins, activity-based protein profiling (ABPP) experiment, western blotting analysis and IHC analysis. There is no doubt that the wealth of information gathered here will be helpful in further studies towards the optimization of the hit molecules’ activity and the determination of their putative drug target(s) and potential mechanism(s) of action.

## 4. Experimental section

### 4. 1. Synthesis

NMR data were collected on a Bruker Ascend^TM^ 400 spectrometer operating at 400 MHz for ^1^H and 100 MHz for ^13^C. The concentration of all samples was about 20 to 50 mg in 0.5 mL of CDCl_3_. The NMR data were recorded at 300 K, with chemical shifts (δ) reported in parts per million (ppm) relative to TMS (δH 0.00 and δC 0.0) used as the internal standard, or residual chloroform (δH 7.28 and δC 77.2), and coupling constants (*J*) in hertz. The multiplicity listed as follow: s = singlet, d = doublet, t = triplet, q = quartet, q* = quintet, m = multiplet. Structures were confirmed based on 1D and 2D NMR data including proton, broadband carbon 13, COSY, NOESY, HSQC and HMBC. Reaction mixtures were monitored by TLC using silica gel 60 F254 plates or by a 200-MS Varian GC/MS ion trap mass spectrometer on which all the GC-MS spectra were recorded. The product(s) from each reaction was purified on a Teledyne ISCO RF200 CombiFlash, using a 40 g RediSepRF silica column, or by gravity and flash column chromatography using type 60A silica gel (60-230 mesh). Single crystals of various samples suitable for X-ray structure determination were mounted on the tip of a glass fiber using silicon grease, and intensity data were recorded at 90 K on an STOE IPDS-2T diffractometer using graphite-monochromated MoKα radiation (λ = 0.71073 Å). Intensity data for all crystals were readily indexed and the structures were solved by direct method and refined by full-matrix least-squares techniques in the SHELTXL package of programs. The structure solutions revealed the positions of all non-hydrogen atoms within the crystal with some of the hydrogen positions. In subsequent refinement steps, the positions of the remaining hydrogen atoms were deduced from different Fourier syntheses. The software Diamond was utilized to create the graphic representations of the crystal structures with ellipsoid representations (50% probability level) for all non-hydrogen atoms.

All chemicals and solvents were purchased from major chemical suppliers and were used without further purification unless stated otherwise.

#### 4.1.1. Methyl 2,6-dibromo-3,4,5-trimethoxybenzoate (**1**)^43^

A mixture of methyl 3,4,5-trimethoxybenzoate (5 g, 22.1 mmol) and 1,3-dibromo-5,5-dimethylhydantoin also known as DBDMH (9.5 g, 33.2 mmol) in dichloromethane was allowed to stir at room temperature for 16 h. The reaction mixture was then filtered and concentrated under reduced pressure. The residue was purified on a silica gel column using a mixture of hexanes-dichloromethane (7:3) to yield the product as yellowish needles (7.4 g, 87%). mp 67.2 – 67.3 °C. ^1^H NMR (CDCl_3_, 400 MHz): δ 3.91 (6H, s), 3.95 (3H, s), 3.98 (3H, s). ^13^CNMR (CDCl_3_, 100 MHz): δ 52.9, 61.0, 61.1, 109.5, 133.1, 148.3, 150.8, 165.9. GC-MS: *m/z* (%) 385 [M+H]^+^ (43), 384 [M]^+^ (100), 353 (25), 326 (15). The structure of this compound was unambiguously determined by single crystal X-ray diffraction. Crystal data: C_11_H_12_Br_2_O_5_, *M*_r_ = 384.03 g/mol, Crystal system monoclinic, Space group *Cc*, *a* = 17.287(4) Å*, b* = 8.9326(2) Å, *c* = 9.0071(2) Å, α = 90°, β = 105.90(3)°, γ = 90°, *V* = 1337.64(5) Å^3^, T = 100(2) K, *Z* = 4, *D*_x_ = 1.91 g/cm^3^, µ = 6.070 mm^-1^, F(000) = 752, λ = 0.71073 Å. Full-matrix least-squares refinement led to a final R = 0.028, wR = 0.069 and GOF = 1.049 (see supplemental data for the molecular representation at 50% probability level.^43^

#### 4.1.2. Methyl 3,4,5-trimethoxy-2-nitrobenzoate (**2**)^43^

This compound was prepared as followed: to a solution of methyl 3,4,5-trimethoxybenzoate (5 g, 22.1 mmol) in glacial acetic acid (25 mL) in an ice bath, 1.1 mL of fuming nitric acid (1.55 g, 24.6 mmol) was carefully added (dropwise) over a period of 30 min. The resulting mixture was allowed to stir at room temperature for 2 h, and was then poured into icy water and neutralized with a saturated solution of sodium bicarbonate. It was then extracted with ethyl acetate, concentrated under reduced pressure and purified on a silica gel column using a mixture of hexane – ethyl acetate (8:2) to yield the product as colorless needles (1.32 g, 22%). mp 62.3 – 63.5 °C. ^1^H NMR (CDCl_3_, 400 MHz): δ 3.87 (3H, s), 3.95 (9H, s), 7.27 (1H, s). ^13^CNMR (CDCl_3_, 100 MHz): δ 52.7, 56.3, 61.0, 62.4, 108.2, 117.2, 139.9, 145.3, 146.0, 153.8, 162.8. GC-MS: *m/z* (%) 272 [M+H]^+^ (15), 271 [M]^+^ (100), 256 (5), 241 (12), 210 (14), 195 (16). The structure of this compound was unambiguously determined by single crystal X-ray diffraction. Crystal data: C_11_H_13_NO_7_, *M*_r_ = 271.22 g/mol, Crystal system monoclinic, Space group *C2/c*, *a* = 20.520(4) Å*, b* = 9.7438(2 Å, *c* = 24.434(5) Å, 〈 = 90°, ® = 99.46(3)°, © = 90°, *V* = 4818.96(2) Å^3^, T = 100(2) K, *Z* = 14, *D*_x_ = 1.31 g/cm^3^, µ = 0.111 mm^-1^, F(000) = 1988, λ = 0.71073 Å. Full-matrix least-squares refinement led to a final R = 0.032, wR = 0.099 and GOF = 1.033.^43^

#### 4.1.3. 4-Cyclohexenyl-N,N-dimethylaniline (**3**)^44^

This compound was prepared by reacting *N,N*-dimethylaniline (5g, 41.3 mmol) with cyclohexanone (20.2 g, 206 mmol) in the presence of a catalytic amount of iodine (524 mg, 2.06 mmol). After purification on a silica gel column using a mixture of hexanes-ethyl acetate (97.5:2.5), compound **3** was obtained as a yellowish oil (4.5 g, 54 %). ^1^H NMR (CDCl_3_, 600 MHz): δ 1.68 (2H, m), 1.80 (2H, m), 2.23 (2H, m), 2.42 (2H, m), 2.96 (6H, s), 6.04 (1H, m), 6.74 (2H, d, *J* = 8.9 Hz), 7.33 (2H, d, *J* = 8.9 Hz). ^13^C NMR (CDCl_3_, 100 MHz): δ 22.5, 23.4, 26.0, 27.5, 40.9, 112.4, 121.3, 125.5, 131.2, 136.0, 149.6. HRESI-MS: [M + H]^+^ Calcd for C_14_H_20_N 202.159026; Found 202.1621.

#### 4.1.4. 2-Chloro-4-cyclohexenyl-N-ethylaniline (**4**)^44^

This compound was obtained by reacting 2-chloro-*N*-ethylaniline (3.55 g, 22.8 mmol.) and cyclohexanone (11.2 g, 114 mmol) in the presence of a catalytic amount of iodine (290 mg, 1.14 mmol). After purification on a silica gel column using hexanes-ethyl acetate (95: 5), compound **4** was obtained as a brownish oil (2.15 g, 40%). ^1^H NMR (CDCl_3_, 400 MHz): δ. 1.29 (3H, t, *J* = 8.0 Hz), 1.63 (2H, m), 1.76 (2H, m), 2.15 (2H, m), 2.32 (2H, m), 3.19 (2H, q, *J* = 8.0 Hz), 4.12 (1H, brs, NH), 5.99 (1H, m), 6.59 (1H, d, *J* = 8.0 Hz), 7.18 (1H, dd, *J* = 4.0 and 8.0 Hz), 7.30 (1H, d, *J* = 4.0 Hz). ^13^C NMR (CDCl_3_, 100 MHz): δ 14.7, 22.2, 23.1, 25.8, 27.3, 38.3, 110.8, 118.9, 122.3, 124.2, 125.6, 132.0, 135.2, 142.7. HRESI-MS: [M]^+^ Calcd for C_14_H_18_ClN 235.1123; Found 235.1121.

#### 4.1.5. 6-Ethoxy-2,2,4-trimethyl-1,2-dihydroquinoline (**5**)^43,45^

*p*-Phenetidine (0.5 g, 3.6 mmol) and an excess of acetone (5 mL) were refluxed in toluene (10 mL) in the presence of a catalytic amount of iodine (92 mg, 0.36 mmol, 10 mol%) for 18 h. The solvent was then removed under reduced pressure on an rota-evaporator, and after purification on a silica gel column using hexanes-ethyl acetate (9: 1), compound **5** was obtained as brownish oil (380 mg, 48 %). ^1^H NMR (CDCl_3_, 400 MHz): δ 1.28 (6H, s), 1.41 (3H, t, *J* = 7.2 Hz), 2.01 (3H, s), 3.42 (1H, brs, NH), 3.99 (2H, q, *J* = 7.2 Hz), 5.39 (1H, s), 6.42 (1H, d, *J* = 7.2Hz), 6.41 (1H, dd, *J* = 7.2 and 4.0 Hz), 6.74 (1H, s). ^13^CNMR (CDCl_3_, 100 MHz): δ 15.1, 18.6, 30.4, 51.7, 64.2, 111.1, 113.7, 114.4, 123.0, 128.6, 129.7, 137.5, 151.3. GC-MS: [M]^+^ *m/z* (%): 217 (13), 216 (24), 201 (100), 173 (22), 144 (6).

#### 4.1.6. 8-Ethoxy-2,2,4-trimethyl-1,2-dihydroquinoline (**6**)^43,45^

*o*-Phenetidine (0.5 g, 3.6 mmol) and an excess of acetone (5 mL) were refluxed in toluene (10 mL) in the presence of a catalytic amount of iodine (92 mg, 0.36 mmol, 10 mol%) for 18 h. The solvent was then removed under reduced pressure on an rota-evaporator, and after purification on a silica gel column using hexanes-ethyl acetate (9: 1), compound **6** was obtained as brownish oil (380 mg, 48 %). ^1^H NMR (CDCl_3_, 400 MHz): δ 1.33 (6H, s), 1.47 (3H, t, *J* = 7.2 Hz), 2.03 (3H, s), 4.01 (2H, q, *J* = 7.2 Hz), 4.29 (1H, brs, NH), 5.35 (1H, s), 6.60 (1H, d, *J* = 6.0 Hz), 6.71 (1H, t, *J* = 6.0 Hz), 6.80 (1H, d, *J* = 6.0 Hz). ^13^CNMR (CDCl_3_, 100 MHz): δ 15.1, 18.8, 31.1, 51.3, 64.1, 110.8, 115.6, 116.2, 121.6, 128.2, 128.7, 133.2, 144. GC-MS: [M]^+^ *m/z* (%): 217 (6), 216 (14), 201 (100), 173 (20), 144 (6).

#### 4.1.7. 8-Chloro-6-cyclopentenyl-2,3,4,5-tetrahydro-4,4-tetramethylene-1H-cyclopenta[c]quinoline (**7**)^45^

This compound was obtained by reacting 4-chloroaniline (5 g, 39 mmol) and cyclopentanone (16.5 g, 196 mmol) in the presence of a catalytic amount of iodine (498 mg, 1.96 mmol). After purification on a silica gel column using hexanes-ethyl acetate (9:1), compound **7** was obtained as yellowish oil (7. 4 g, 58 %). ^1^H NMR (CDCl_3_, 400 MHz): δ 1.67 – 1.70 (6H, m), 1.83 – 1.85 (2H, m), 1.95 – 2.03 (4H, m), 2.49 – 2.52 (2H, m), 2.54 – 2.56 (2H, m), 2.60 – 2.65 (4H, m), 4.64 (1H, brs, NH), 5.90 (1H, m), 6.73 (1H, d, *J* = 2.0 Hz), 6.84 (1H, d *J* = 2.0 Hz). ^13^CNMR (CDCl_3_, 100 MHz): δ 22.4, 23.3, 23.8, 31.4, 32.2, 33.9, 36.7, 39.6, 65.1, 121.1, 121.7, 122.0, 122.8, 125.9, 128.4, 132.0, 138.3, 140.1, 140.7. The *spiro*-quaternary carbon (C-4) appears at 65.1 ppm, while the g-HSQC and g-HMBC sequences enable to find the key correlation between the protons at δ 5.90 ppm (the proton on the cyclopentenyl double bond carried by the carbon at 125.9 ppm) and the quaternary aromatic carbon (C-6) at δ 122.8 ppm. HRESI-MS: [M + H]^+^ Calcd for C_21_H_25_NCl 326.167004; Found 326.1669.

#### 4.1.8. 8-Cyclohexenyl-6-ethoxy-1,2-dihydro-2,2,4-trimethylquinoline (**8**)^44,45^

This compound was obtained by reacting ethoxyquin (1.5 g, 6.9 mmol) and cyclohexanone (3.4 g, 34.5 mmol) in the presence of a catalytic amount of iodine (88 mg, 0.35 mmol). After purification on a silica gel column using hexanes-ethyl acetate (9: 1), compound **8** was obtained as yellowish oil (1.11 g, 54 %). ^1^H NMR (CDCl_3_, 400 MHz): δ 1.22 (6H, s), 1.36 (3H, t, *J* = 6.9 Hz), 1.68 – 1.69 (2H, m), 1.75 – 1.76 (2H, m), 1.98 (3H, s), 2.16 – 2.18 (4H, m), 3.42 (1H, brs, NH), 3.96 (2H, q, *J* = 6.9 Hz), 5.37 (1H, s), 5.71 (1H, m), 6.46 (1H, d, *J* = 2.8 Hz), 6.62 (1H, d, *J* = 2.8 Hz). ^13^CNMR (CDCl_3_, 100 MHz): δ 15.2, 18.9, 22.2, 23.3, 25.5, 29.8, 30.0, 50.8, 51.5, 64.2, 109.5, 114.1, 123.4, 127.2, 129.1, 129.9, 134.0, 136.0, 150.7. The g-HSQC and g-HMBC sequences enable us to find the key correlation between the proton at δ 5.71 ppm (the proton on the cyclohexenyl double bond carried by the carbon at 127.2 ppm) and the quaternary aromatic carbon (C-6) at δ 129.1 ppm. HRESI-MS: [M + H]^+^ Calcd for C_20_H_28_NO 298.216541; Found 298.2158.

#### 4.1.9. 6,8-Dicyclopentenyl-1,2-dihydro-2,2,4-trimethylquinoline (**9**)^44,45^

This compound was obtained by reacting 2,2,4-trimethyl-1,2-dihydroquinoline (2.5 g, 14.4 mmol.) and cyclopentanone (6.07 g, 72.2 mmol) in the presence of a catalytic amount of iodine (183 mg, 0.722 mmol). After purification on a silica gel column using hexanes-ethyl acetate (97.5: 2.5), compound **9** was obtained as a yellowish oil (2.56 g, 58 %). ^1^H NMR (CDCl_3_, 400 MHz): δ 1.27 (6H, s), 1.51-1.55 (2H, m), 1.57-1.59 (2H, m), 1.63-1.68 (2H, m), 2.01-2.05 (2H, m), 2.02 (3H, s), 3.54-3.60 (2H, m), 2.65-2.70 (2H, m), 4.39 (1H, brs, NH), 5.33 (1H, s), 5.86-5.89 (2H, m), 6.84 (1H, d, *J* = 2.0 Hz), 6.87 (1H, d, *J* = 2.0 Hz). ^13^C NMR (CDCl_3_, 100 MHz): δ 19.1, 23.4, 25.6, 31.1, 34.0, 34.9, 36.9, 45.7, 51.7, 121.3, 121.6, 122.1, 126.1, 127.9, 128.5, 129.2, 133.8, 138.5, 141.7. HRESI-MS: [M]^+^ Calcd for C_22_H_27_N 305.2144; Found 305.2152.

#### 4.1.10. N1,N2-diethyl-4-methoxy-N2-(4-methoxyphenyl)benzene-1,2-diamine (**11**)^46^

This compound was prepared by allowing *N*-ethyl-4-methoxyaniline (**10**) (1.3 g, 8.60 mmol.), obtained from Sigma-Aldrich, to stir for 20 minutes in an ice bath in the presence of sodium hydroxide (688 mg, 17.2 mmol, 2 equiv.), silver nitrate (73 mg, 0.43 mmol., 5 mol%.) and water (15 mL). Then a 25% solution of sodium persulfate (1 equiv.) in water was added to the reaction, and the reaction was allowed to stir in an ice bath for 4 hours, with the temperature kept between 0 – 3 °C. The reaction mixture was then extracted with ethyl acetate (2 x 50 mL), dried with sodium sulfate, gravity filtered, and solvent was removed under reduced pressure. The residue was then purified on a silica gel column using hexanes-ethyl acetate (97.5: 2.5) to yield compound **11** as a yellowish oil (1.63 g, 63%). ^1^H NMR (400 MHz, CDCl_3_) 1.19 (6H, t, *J* = 8.0 Hz), 3.13 (2H, q, *J* = 7.9 Hz), 3.57 (2H, q, *J* = 7.8 Hz), 3.74 (3H, s), 3.77 (3H, s), 3.91 (1H, brs, NH), 6.66 (2H, d, *J* = 6.8 Hz), 6.72 (1H, d, *J* = 3.4 Hz), 6.79 (2H, d, *J* = 6.8 Hz), 7.29 (2H, d, *J* = 6.7 Hz). ^13^C NMR (100 MHz, CDCl_3_): δ 12.8, 15.0, 39.1, 45.3, 55.8, 55.9, 112.3, 112.6, 114.7, 114.9, 115.5, 134.0, 140.6, 142.7, 151.7, 152.2. HRESI-MS: [M + H]^+^ Calcd for C_18_H_24_N_2_O_2_ 301.19105; Found 301.19224.

Compounds **12 – 35** were all prepared following a similar protocol.^47^ The aniline derivative (1 eqv) and the phenol coupling partner (1 eqv) were dissolved in toluene, and 1 eqv of AgNO_3_ was added. The mixture was then cooled in an ice bath and 3 eqv of H_2_O_2_ (30% in H_2_O) was added dropwise with a Pasteur pipette. The reaction was then stirred for 6 h under atmospheric air during which the temperature was allowed to rise to room temperature. The reaction mixture was then extracted from water with 2 portions of 25 ml of ethyl acetate, and the combined extracts were dried over Na_2_SO_4_ and purified on a silica gel column using a mixture of hexanes-ethyl acetate to yield the expected product.

#### 4.1.11. 1-(2-Amino-N,N,5-trimethylphenyl)naphthalen-2-ol **(12)**^47^

*N*,*N*,4-trimethylaniline (500 mg, 3.70 mmol) and 2-naphthol (533 mg, 3.70 mmol) were reacted in toluene (5 mL) in the presence of AgNO_3_ (628 mg, 3.70 mmol) and H_2_O_2_ (1.26 g, 11.1 mmol). The product was isolated by chromatography on a silica gel column using hexanes-ethyl acetate (9:1), and obtained as clear colorless crystals (670 mg, 65 %). mp 171.8 – 172.2 °C. ^1^H NMR (400 MHz, CDCl_3_): δ (ppm) 2.32 (3H, s), 2.66 (6H, s), 7.15 – 7.28 (4H, m), 7.31 – 7.39 (2H, m), 7.76 – 7.83 (3H, m), 10.53 (1H, brs). ^13^C NMR (100 MHz, CDCl_3_): δ (ppm) 20.7, 43.9, 118.2, 120.7, 121.0, 123.1, 125.4, 126.2, 128.2, 129.1, 129.4, 130.0, 130.6, 132.8, 133.5, 135.8, 147.4, 152.2. HRMS (ESI/Q-TOF): [M + H]^+^ Calcd for C_19_H_20_NO 278.1539; Found 278.1538. The structure of this compound was unambiguously determined by single crystal X-ray diffraction. Crystal data: C_19_H_19_NO, *M*_r_ = 277.35 g/mol, Crystal system monoclinic, Space group P 2_1_/c, *a* = 9.302(5) Å*, b* = 8.916(4) Å, *c* = 19.097(9) Å, α = 90°, β = 104.096(1)°, γ = 90°, *V* = 1536.2(13) Å^3^, T = 294(2) K, *Z* = 4, *D*_x_ = 1.199 g/cm^3^, µ = 0.074 mm^-1^, F(000) = 592, λ = 0.71073 Å. Full-matrix least-squares refinement led to a final R = 0.0443, wR = 0.1231 and GOF = 1.033. The crystal structure data are deposited in the Cambridge database (CCDC 821630).^48^

#### 4.1.12. 1-(2-Amino-5-ethoxy-N,N-dimethylphenyl)naphthalen-2-ol **(13)**^47^

4-Ethoxy-*N*,*N*-dimethylaniline (500 mg, 3.03 mmol) and 2-naphthol (436 mg, 3.03 mmol) were reacted in toluene (5 mL) in the presence of AgNO_3_ (514 mg, 3.03 mmol) and H_2_O_2_ (1.03 g, 9.09 mmol). The product was isolated by chromatography on a silica gel column using hexanes-ethyl acetate (9:1), and obtained as clear colorless crystals (653 mg, 70 %). mp 130.8 – 131.1 °C. ^1^H NMR (400 MHz, CDCl_3_): δ (ppm) 1.34 (3H, t, *J* = 6.8 Hz), 2.63 (6H, s), 3.94 (2H, q, *J* = 6.8 Hz), 6.95 (2H, m), 7.16 (1H, d, *J* = 8.8 Hz), 7.27 (1H, d, *J* = 8.8 Hz), 7.31 – 7.38 (2H, m), 7.76 – 7.81 (2H, m), 7.85 (1H, d, *J* = 8.0 Hz), 10.61 (1H, brs). ^13^C NMR (100 MHz, CDCl_3_): δ (ppm) 14.8, 43.9, 63.8, 114.9, 119.0, 120.5, 120.8, 120.9, 123.0, 125.3, 126.1, 128.1, 129.4, 129.8, 132.1, 133.4, 143.3, 152.3, 154.4. HRMS (ESI/Q-TOF): [M + H]^+^ Calcd for C_20_H_22_NO_2_ 308.1645; Found 308.1655. The structure of this compound was unambiguously determined by single crystal X-ray diffraction. Crystal data: C_20_H_21_NO_2_, *M*_r_ = 307.38 g/mol, Crystal system monoclinic, Space group P 2_1_/c, *a* = 9.267(3) Å*, b* = 7.093(2) Å, *c* = 25.135(8) Å, α = 90°, β = 91.104(4)°, γ = 90°, *V* = 1651.8(9) Å^3^, T = 546(2) K, *Z* = 4, *D*_x_ = 1.236 g/cm^3^, *θ*_max_ = 28.96°(MoKα), *R* = 0.0374 for 4084 data and 208 refined parameters. The crystal structure data are deposited in the Cambridge database (CCDC 1442427).^47^

#### 4.1.13. 1-(2-Amino-N-benzyl-5-ethoxy-N-methylphenyl)naphthalen-2-ol (**14**)^47^

*N*-benzyl-4-ethoxy-*N*-methylaniline (500 mg, 2.07 mmol) and 2-naphthol (299 mg, 2.07 mmol) were reacted in toluene (5 mL) in the presence of AgNO_3_ (352 mg, 2.07 mmol) and H_2_O_2_ (705 mg, 6.22 mmol). The product was isolated by chromatography on a silica gel column using hexanes-ethyl acetate (9:1), and obtained as brownish viscous oil (481 mg, 61 %). ^1^H NMR (400 MHz, CDCl_3_): δ (ppm) 1.36 (3H, t, *J* = 6.8 Hz), 2.59 (3H, s), 3.75 (1H, d, *J* = 13.2 Hz), 3.85 (1H, d, *J* = 13.2 Hz), 3.96 (2H, q, *J* = 6.8 Hz), 6.95 (3H, m), 7.00 (1H, d, *J* = 2.7 Hz), 7.12 (1H, d, *J* = 8.8 Hz), 7.19 (3H, m), 7.31 – 7.37 (3H, m), 7.80 (2H, d, *J* = 8.6 Hz), 7.88 (1H, d, *J* = 8.4Hz), 10.01 (1H, brs). ^13^C NMR (100 MHz, CDCl_3_): δ (ppm) 14.8, 39.0, 62.1, 63.8, 115.0, 120.4, 120.6, 120.7, 120.9, 123.1, 125.1, 126.3, 127.5, 128.3, 129.4, 129.6, 130.0, 132.3, 133.4, 135.9, 143.0, 152.0, 154.7. HRMS (ESI/Q-TOF): [M + H]^+^ Calcd for C_26_H_26_NO_2_ 384.1958; Found 384.1973.

#### 4.1.14. 1-[2-Amino-N-((2E)but-2-enyl)-5-ethoxy-N-methylphenyl]naphthalen-2-ol (**15**)^47^

(E)-*N*-(but-2-enyl)-4-ethoxy-*N*-methylaniline (500 mg, 2.44 mmol) and 2-naphthol (351 mg, 2.44 mmol) were reacted in toluene (5 mL) in the presence of AgNO_3_ (414 mg, 2.44 mmol) and H_2_O_2_ (829 mg, 7.31 mmol). The product was isolated by chromatography on a silica gel column using hexanes-ethyl acetate (9:1), and obtained as yellowish oil (542 mg, 64 %). ^1^H NMR (400 MHz, CDCl_3_): δ (ppm) 1.35 (3H, t, *J* = 7.2 Hz), 1.57 (3H, d, *J* = 6.0 Hz), 2.68 (3H, s), 3.19 – 3.30 (2H, m), 3.95 (2H, q, *J* = 7.2 Hz), 5.20 – 5.27 (1H, m), 5.34 – 5.39 (1H, m), 6.92 – 6.97 (2H, m), 7.15 (1H, d, *J* = 8.4 Hz), 7.24 – 7.36 (3H, m), 7.78 – 7.85 (3H, m), 10.45 (1H, s). ^13^C NMR (100 MHz, CDCl_3_): δ (ppm) 14.8, 17.7, 39.2, 53.4, 59.7, 64.3, 114.8, 120.1, 120.5, 120.9, 123.0, 125.3, 125.9, 126.1, 128.2, 129.5, 129.9, 130.7, 132.4, 133.4, 142.8, 152.1, 154.4. HRMS (ESI/Q-TOF): [M + H]^+^ Calcd for C_23_H_26_NO_2_ 348.1958; Found 348.1962.

#### 4.1.15. 2-(2-Amino-5,N,N-trimethylphenyl)naphthalen-1-ol (**16**)^47^

*N*,*N*,4-trimethylaniline (500 mg, 3.70 mmol) and 1-naphthol (533 mg, 3.70 mmol) were reacted in toluene (5 mL) in the presence of AgNO_3_ (628 mg, 3.70 mmol) and H_2_O_2_ (1.26 g, 11.1 mmol). The product was isolated by chromatography on a silica gel column using hexanes-ethyl acetate (9:1), and obtained as brownish crystals (540 mg, 53 %). mp 124.5 – 125.0 °C. ^1^H NMR (400 MHz, CDCl_3_): δ (ppm). 2.36 (3H, s), 2.66 (6H, s), 7.10 – 7.17 (2H, m), 7.31 (1H, d, *J* = 1.6 Hz), 7.43 – 7.53 (4H, m), 7.77 – 7.80 (1H, m), 8.46 – 8.48 (1H, m), 11.92 (1H, brs).^13^C NMR (100 MHz, CDCl_3_): δ (ppm) 20.8, 43.6, 118.1, 119.6, 120.7, 123.6, 125.0, 126.3, 126.9, 127.0, 128.4, 128.6, 134.1, 134.2, 134.4, 134.9, 145.8, 151.5. HRMS (ESI/Q-TOF): [M + H]^+^ Calcd for C_19_H_20_NO 278.1539; Found 278.1544. The structure of this compound was unambiguously determined by single crystal X-ray diffraction. Crystal data: C_19_H_19_NO, *M*_r_ = 277.35 g/mol, Crystal system orthorhombic, Space group Pbca, *a* = 7.633(2) Å*, b* = 15.037(4) Å, *c* = 27.693(8) Å, α = 90°, β = 90°, γ = 90°, *V* = 3178.5(16) Å^3^, T = 293(2) K, *Z* = 8, *D*_x_ = 1.159 g/cm^3^, *θ*_max_ = 23.29°(MoKα), *R* = 0.0690 for 2284 data and 190 refined parameters. The crystal structure data are deposited in the Cambridge database (CCDC 1442428).^47^

#### 4.1.16. 2’-(Dimethylamino)-5’-ethoxybiphenyl-2-ol (**17**)^47^

4-Ethoxy-*N*,*N*-dimethylaniline (500 mg, 3.03 mmol) and phenol (285 mg, 3.03 mmol) were reacted in toluene (5 mL) in the presence of AgNO_3_ (514 mg, 3.03 mmol) and H_2_O_2_ (1.03 g, 9.08 mmol). The product was isolated by chromatography on a silica gel column using hexanes-ethyl acetate (9:1), and obtained as yellowish oil (512 mg, 66 %). ^1^H NMR (400 MHz, CDCl_3_): δ (ppm) 1.33 (3H, t, *J* = 7.2 Hz), 2.75 (6H, s), 3.89 (2H, q, *J* = 7.2 Hz), 6.45 (1H, d, *J* = 2.8 Hz), 6.60 (1H, dd, *J* = 2.8 and 8.8 Hz), 6.95 (1H, d, *J* = 8.8 Hz), 7.00 (1H, d, *J* = 7.6 Hz), 7.01 (1H, brs), 7.06 (1H, dd, *J* = 7.2 and 7.6 Hz), 7.29 (1H, d, *J* = 7.6 Hz), 7.31 (1H, d, *J* = 7.6 Hz).^13^C NMR (100 MHz, CDCl_3_): δ (ppm) 14.8, 43.7, 63.7, 107.3, 109.1, 118.3, 119.4, 122.9, 129.6, 138.4, 150.1, 155,3, 158.5. HRMS (ESI/Q-TOF): [M + H]^+^ Calcd for C_16_H_20_NO_2_ 258.1498; Found 258.1489.

#### 4.1.17. 5-Bromo-2’-(dimethylamino)-5’-ethoxybiphenyl-2-ol (**18**)^47^

4-Ethoxy-*N*,*N*-dimethylaniline (500 mg, 3.03 mmol) and 4-bromophenol (524 mg, 3.03 mmol) were reacted in toluene (5 mL) in the presence of AgNO_3_ (514 mg, 3.03 mmol) and H_2_O_2_ (1.03 g, 9.08 mmol). The product was isolated by chromatography on a silica gel column using hexanes-ethyl acetate (9:1), and obtained as yellowish oil (602 mg, 59 %). ^1^H NMR (400 MHz, CDCl_3_): δ (ppm) 1.34 (3H, t, *J* = 7.2 Hz), 2.71 (6H, s), 3.91 (2H, q, *J* = 7.2 Hz), 6.46 (1H, d, *J* = 2.8 Hz), 6.64 (1H, dd, *J* = 2.8 and 8.8 Hz), 6.85 (1H, d, *J* = 8.8 Hz), 6.86 (1H, brs), 6.95 (1H, d, *J* = 8.8 Hz), 7.38 (2H, d, *J* = 8.8 Hz). ^13^C NMR (100 MHz, CDCl_3_): δ (ppm) 14.8, 43.6, 63.9, 107.7, 109.9, 115.1, 119.5, 119.6, 132.4, 138.5, 149.3, 154.6, 156.5. HRMS (ESI/Q-TOF): [M]^+^ Calcd for C_16_H_18_BrNO_2_ 336.0594; Found 336.0615.

#### 4.1.18. 5-Chloro-2’-(dimethylamino)-5’-ethoxybiphenyl-2-ol (**19**)^47^

4-ethoxy-*N*,*N*-dimethylaniline (500 mg, 3.03 mmol) and 4-chlorophenol (389 mg, 3.03 mmol) were reacted in toluene (5 mL) in the presence of AgNO_3_ (514 mg, 3.03 mmol) and H_2_O_2_ (1.03 g, 9.08 mmol). The product was isolated by chromatography on a silica gel column using hexanes-ethyl acetate (9:1), and obtained as yellowish oil (532 mg, 60 %). ^1^H NMR (400 MHz, CDCl_3_): δ (ppm) 1.32 (3H, t, *J* = 7.2 Hz), 2.70 (6H, s), 3.88 (2H, q, *J* = 7.2 Hz), 6.47 (1H, d, *J* = 2.8 Hz), 6.63 (1H, dd, *J* = 2.8 and 8.8 Hz), 6.89 (1H, d, *J* = 8.8 Hz), 6.90 (1H, brs), 6.93 (1H, d, *J* = 8.8 Hz), 7.21 (2H, d, *J* = 9.2 Hz). ^13^C NMR (100 MHz, CDCl_3_): δ (ppm) 14.8, 43.5, 63.7, 107.7, 109.8, 119.0 119.5, 127.6, 129.4, 138.4, 149.4, 154.6, 156.0. HRMS (ESI/Q-TOF): [M + H]^+^ Calcd for C_16_H_19_ClNO_2_ 292.1099; Found 292.1116.

#### 4.1.19. 2’-(Dimethylamino)-5’-ethoxy-5-methoxybiphenyl-2-ol (**20**)^47^

4-Ethoxy-*N*,*N*-dimethylaniline (500 mg, 3.03 mmol) and 4-methoxyphenol (376 mg, 3.03 mmol) were reacted in toluene (5 mL) in the presence of AgNO_3_ (514 mg, 3.03 mmol) and H_2_O_2_ (1.03 g, 9.08 mmol). The product was isolated by chromatography on a silica gel column using hexanes-ethyl acetate (9:1), and obtained as yellowish oil (493 mg, 57 %). ^1^H NMR (400 MHz, CDCl_3_): δ (ppm) 1.31 (3H, t, *J* = 7.2 Hz), 2.77 (6H, s), 3.76 (3H, s), 3.87 (2H, q, *J* = 7.2 Hz), 6.34 (1H, d, *J* = 2.8 Hz), 6.53 (1H, dd, *J* = 2.8 and 8.8 Hz), 6.85 (1H, d, *J* = 8.8 Hz), 6.87 (1H, brs), 6.93 (1H, d, *J* = 8.8 Hz), 6.98 (2H, d, *J* = 9.2 Hz). ^13^C NMR (100 MHz, CDCl_3_): δ (ppm) 14.7, 43.7, 55.5, 63.6, 105.7, 107.7, 114.7, 119.1, 120.2, 137.6, 150.1, 151.7, 154.7, 155.7. HRMS (ESI/Q-TOF): [M + H]^+^ Calcd for C_17_H_22_NO_3_ 288.1594; Found 288.1596.

#### 4.1.20. 2’-(Dimethylamino)-5’-ethoxy-5-methylbiphenyl-2-ol (**21**)^47^

4-Ethoxy-*N*,*N*-dimethylaniline (500 mg, 3.03 mmol) and 4-chlorophenol (327 mg, 3.03 mmol) were reacted in toluene (5 mL) in the presence of AgNO_3_ (514 mg, 3.03 mmol) and H_2_O_2_ (1.03 g, 9.08 mmol). The product was isolated by chromatography on a silica gel column using hexanes-ethyl acetate (9:1), and obtained as colorless oil (573 mg, 70 %). ^1^H NMR (400 MHz, CDCl_3_): δ (ppm) 1.31 (3H, t, *J* = 7.2 Hz), 2.31 (3H, s), 2.75 (6H, s), 3.88 (2H, q, *J* = 7.2 Hz), 6.40 (1H, d, *J* = 2.8 Hz), 6.57 (1H, dd, *J* = 2.8 and 8.8 Hz), 6.91 (1H, d, *J* = 7.2 Hz), 6.93 (1H, brs), 6.93 (1H, d, *J* = 7.6 Hz), 7.10 (2H, d, *J* = 8.4 Hz). ^13^C NMR (100 MHz, CDCl_3_): δ (ppm) 14.8, 20.7, 43.8, 63.7, 106.6, 108.5, 118.6, 119.2, 130.1, 132.5, 138.0, 151.6, 154.6. HRMS (ESI/Q-TOF): [M + H]^+^ Calcd for C_17_H_22_NO_2_ 272.1645; Found 272.1656.

#### 4.1.21. 2’-(Dimethylamino)-5’-ethoxy-5-isopropylbiphenyl-2-ol (**22**)^47^

4-Ethoxy-*N*,*N*-dimethylaniline (500 mg, 3.03 mmol) and 4-isopropylphenol (412 mg, 3.03 mmol) were reacted in toluene (5 mL) in the presence of AgNO_3_ (514 mg, 3.03 mmol) and H_2_O_2_ (1.03 g, 9.08 mmol). The product was isolated by chromatography on a silica gel column using hexanes-ethyl acetate (9:1), and obtained as yellowish oil (474 mg, 52 %). ^1^H NMR (400 MHz, CDCl_3_): δ (ppm) 1.24 (6H, d, *J* = 7.2 Hz), 1.32 (3H, t, *J* = 7.2 Hz), 2.76 (6H, s), 2.88 (1H, m, *J* = 7.2 Hz), 3.89 (2H, q, *J* = 7.2 Hz), 6.42 (1H, d, *J* = 2.8 Hz), 6.57 (1H, dd, *J* = 2.8 and 8.8 Hz), 6.93 (1H, d, *J* = 6.8 Hz), 6.96 (1H, d, *J* = 6.8 Hz), 6.98 (1H, brs), 7.16 (2H, d, *J* = 8.4 Hz). ^13^C NMR (100 MHz, CDCl_3_): δ (ppm) 14.8, 24.2, 33.4, 43.8, 63.5, 106.7, 108.2, 118.6, 119.2, 127.5, 138.0, 143.6, 150.9, 154.6, 154.7. HRMS (ESI/Q-TOF): [M + H]^+^ Calcd for C_19_H_26_NO_2_ 300.1958; Found 300.1954.

#### 4.1.22. 4-Chloro-2’-(dimethylamino)-5’-ethoxybiphenyl-2-ol (**23**)^47^

4-Ethoxy-*N*,*N*-dimethylaniline (500 mg, 3.03 mmol) and 3-chlorophenol (389 mg, 3.03 mmol) were reacted in toluene (5 mL) in the presence of AgNO_3_ (514 mg, 3.03 mmol) and H_2_O_2_ (1.03 g, 9.08 mmol). The product was isolated by chromatography on a silica gel column using hexanes-ethyl acetate (9:1), and obtained as yellowish oil (509 mg, 58 %). ^1^H NMR (400 MHz, CDCl_3_): δ (ppm) 1.35 (3H, t, *J* = 7.2 Hz), 2.71 (6H, s), 3.92 (2H, q, *J* = 7.2 Hz), 6.49 (1H, d, *J* = 2.8 Hz), 6.66 (1H, dd, *J* = 2.4 and 8.8 Hz), 6.86 (1H, dd, *J* = 1.6 and 8.4 Hz), 6.95 (2H, m), 7.01 (1H, d, *J* = 8.0 Hz), 7.19 (1H, t, *J* = 8.0 Hz). ^13^C NMR (100 MHz, CDCl_3_): δ (ppm) 14.8, 43.6, 63.8, 108.1, 110.2, 115.9, 117.9, 119.6, 122.7, 130.3, 134.9, 138.6, 148.9, 154.6, 158.2. HRMS (ESI/Q-TOF): [M + H]^+^ Calcd for C_16_H_19_ClNO_2_ 292.1099; Found 292.1107.

#### 4.1.23. 4,5-Dichloro-2’-(dimethylamino)-5’-ethoxybiphenyl-2-ol (**24**)^47^

4-Ethoxy-*N*,*N*-dimethylaniline (500 mg, 3.03 mmol) and 3,4-dichlorophenol (493 mg, 3.03 mmol) were reacted in toluene (5 mL) in the presence of AgNO_3_ (514 mg, 3.03 mmol) and H_2_O_2_ (1.03 g, 9.08 mmol). The product was isolated by chromatography on a silica gel column using hexanes-ethyl acetate (9:1), and obtained as yellowish oil (531 mg, 54 %). ^1^H NMR (400 MHz, CDCl_3_): δ (ppm) 1.36 (3H, t, *J* = 7.2 Hz), 2.69 (6H, s), 3.93 (2H, q, *J* = 7.2 Hz), 6.50 (1H, d, *J* = 2.4 Hz), 6.68 (1H, dd, *J* = 2.4 and 8.8 Hz), 6.80 (1H, dd, *J* = 2.8 and 8.8 Hz), 6.96 (1H, d, *J* = 8.8 Hz), 7.04 (1H, d, *J* = 2.8 Hz), 7.32 (1H, d, *J* = 8.8 Hz). ^13^C NMR (100 MHz, CDCl_3_): δ (ppm) 14.8, 43.5, 63.8, 108.2, 110.5, 116.9, 119.1, 119.7, 125.7, 130.7, 132.9, 138.7, 148.4, 154.6, 156.6. HRMS (ESI/Q-TOF): [M]^+^ Calcd for C_16_H_17_Cl_2_NO_2_ 326.0709; Found 326.0715.

#### 4.1.24. 2’-(Dimethylamino)-5’-ethoxy-4,5-dimethylbiphenyl-2-ol (**25**)^47^

4-Ethoxy-*N*,*N*-dimethylaniline (500 mg, 3.03 mmol) and 3,4-dimethylphenol (370 mg, 3.03 mmol) were reacted in toluene (5 mL) in the presence of AgNO_3_ (514 mg, 3.03 mmol) and H_2_O_2_ (1.03 g, 9.08 mmol). The product was isolated by chromatography on a silica gel column using hexanes-ethyl acetate (9:1), and obtained as colorless oil (513 mg, 59 %). ^1^H NMR (400 MHz, CDCl_3_): δ (ppm) 1.31 (3H, t, *J* = 7.2 Hz), 2.21 (6H, s), 2.76 (6H, s), 3.87 (2H, q, *J* = 7.2 Hz), 6.40 (1H, d, *J* = 2.8 Hz), 6.56 (1H, dd, *J* = 2.8 and 8.8 Hz), 6.78 (1H, dd, *J* = 2.8 and 8.4 Hz), 6.84 (1H, d, *J* = 2.4 Hz), 6.93 (1H, d, *J* = 8.8 Hz), 7.05 (1H, d, *J* = 8.4 Hz). ^13^C NMR (100 MHz, CDCl_3_): δ (ppm) 14.8, 19.0, 19.9, 43.8, 63.6, 106.5, 108.1, 116.0, 119.1, 120.1, 130.5, 131.2, 137.9, 138.0, 151.0, 154.6, 154.7. HRMS (ESI/Q-TOF): [M + H]^+^ Calcd for C_18_H_24_NO_2_ 286.1802; Found 286.1800.

#### 4.1.25. 5’-Ethoxy-2’-(methyl(propyl)amino)biphenyl-2-ol (**26**)^47^

4-Ethoxy-*N*-methyl-*N*-propylaniline (500 mg, 2.59 mmol) and phenol (243 mg, 2.59 mmol) were reacted in toluene (5 mL) in the presence of AgNO_3_ (439 mg, 2.59 mmol) and H_2_O_2_ (880 mg, 7.76 mmol). The product was isolated by chromatography on a silica gel column using hexanes-ethyl acetate (9:1), and obtained as yellowish oil (308 mg, 42 %). ^1^H NMR (400 MHz, CDCl_3_): δ (ppm) 0.78 (3H, t, *J* = 7.2 Hz), 1.34 (3H, t, *J* = 7.2 Hz), 1.47 (2H, q, *J* = 7.6 Hz), 2.73 (3H, s), 2.94 (2H, t, *J* = 7.6 Hz), 3.91 (2H, q, *J* = 7.2 Hz), 6.48 (1H, d, *J* = 2.8 Hz), 6.62 (1H, dd, *J* = 2.8 and 8.8 Hz), 6.97 (2H, d, *J* = 8.4 Hz), 6.99 (1H, brs), 7.04 (1H, d, *J* = 7.2 Hz), 7.29 (2H, t, *J* = 8.0 Hz). ^13^C NMR (100 MHz, CDCl_3_): δ (ppm) 11.5, 14.8, 20.3, 40.8, 57.7, 63.8, 107.5, 109.3, 117.9, 121.0, 122.5, 129.5, 137.8, 150.3, 154.6, 157.4. HRMS (ESI/Q-TOF): [M + H]^+^ Calcd for C_18_H_24_NO_2_ 286.1802; Found 286.1801.

#### 4.1.26. 5-Bromo-5’-ethoxy-2’-(methyl(propyl)amino)biphenyl-2-ol (**27**)^47^

4-Ethoxy-*N*-methyl-*N*-propylaniline (500 mg, 2.59 mmol) and 4-bromophenol (448 mg, 2.59 mmol) were reacted in toluene (5 mL) in the presence of AgNO_3_ (439 mg, 2.59 mmol) and H_2_O_2_ (880 mg, 7.76 mmol). The product was isolated by chromatography on a silica gel column using hexanes-ethyl acetate (9:1), and obtained as yellowish oil (381 mg, 40 %). ^1^H NMR (400 MHz, CDCl_3_): δ (ppm) 0.75 (3H, t, *J* = 7.6 Hz), 1.35 (3H, t, *J* = 7.2 Hz), 1.42 (2H, q, *J* = 7.6 Hz), 2.68 (3H, s), 2.88 (2H, t, *J* = 7.6 Hz), 3.92 (2H. q, *J* = 7.2 Hz), 6.48 (1H, d, *J* = 2.4 Hz), 6.65 (1H, dd, *J* = 2.8 and 8.8 Hz), 6.81 (2H, d, *J* = 8.8 Hz), 6.96 (1H, d, *J* = 8.8 Hz), 7.37 (2H, d, *J* = 8.8 Hz). ^13^C NMR (100 MHz, CDCl_3_): δ (ppm) 11.5, 14.9, 20.2, 40.7, 57.6, 63.8, 108.0, 110.1, 114.7, 119.2, 121.2, 132.3, 138.0, 149.5, 154.6, 156.9. HRMS (ESI/Q-TOF): [M]^+^ Calcd for C_18_H_22_BrNO_2_ 364.0907; Found 364.0927.

#### 4.1.27. 5’-Ethoxy-2’-(isobutyl(methyl)amino)biphenyl-2-ol (**28**)^47^

4-Ethoxy-*N*-isobutyl-*N*-methylaniline (500 mg, 2.41 mmol) and phenol (227 mg, 2.41 mmol) were reacted in toluene (5 mL) in the presence of AgNO_3_ (410 mg, 2.41 mmol) and H_2_O_2_ (820 mg, 7.24 mmol). The product was isolated by chromatography on a silica gel column using hexanes-ethyl acetate (9:1), and obtained as colorless oil (230 mg, 32 %). ^1^H NMR (400 MHz, CDCl_3_): δ (ppm) 0.75 (6H, d, *J* = 6.8 Hz), 1.35 (3H, t, *J* = 6.8 Hz), 1.77 (1H, m), 2.70 (5H, m), 3.92 (2H, q, *J* = 7.2 Hz), 6.50 (1H, d, *J* = 2.8 Hz), 6.63 (1H, dd, *J* = 2.8 and 8.8 Hz), 6.91 (2H, d, *J* = 8.0 Hz), 6.98 (1H, d, *J* = 8.8 Hz), 7.02 (1H, d, *J* = 7.2 Hz), 7.25 – 7.29 (2H, m). ^13^C NMR (100 MHz, CDCl_3_): δ (ppm) 14.8, 20.5, 26.5, 41.5, 63.6, 108.0, 109.8, 117.5, 121.3, 122.3, 129.4, 138.7, 150.1, 154.6, 157.7. EI-MS: [M]^+^ *m/z* 299.

#### 4.1.28. 5-Bromo-5’-ethoxy-2’-(isobutyl(methyl)amino)biphenyl-2-ol **(29)**^47^

4-Ethoxy-*N*-isobutyl-*N*-methylaniline (500 mg, 2.41 mmol) and 4-bromophenol (417 mg, 2.41 mmol) were reacted in toluene (5 mL) in the presence of AgNO_3_ (410 mg, 2.41 mmol) and H_2_O_2_ (820 mg, 7.24 mmol). The product was isolated by chromatography on a silica gel column using hexanes-ethyl acetate (9:1), and obtained as yellowish oil (208 mg, 23 %). ^1^H NMR (400 MHz, CDCl_3_): δ (ppm) 0.73 (6H, d, *J* = 6.8 Hz), 1.36 (3H, t, *J* = 7.2 Hz), 1.75 (1H, m), 2.65 – 2.68 (5H, m), 3.93 (2H, t, *J* = 7.2 Hz), 6.50 (1H, d, *J* = 2.8 Hz), 6.66 (1H, dd, *J* = 2.8 and 8.8 Hz), 6.78 (2H, d, *J* = 9.2 Hz), 6.98 (1H, d, *J* = 8.8 Hz), 7.35 (2H, d, *J* = 9.2 Hz). ^13^C NMR (100 MHz, CDCl_3_): δ (ppm) 14.8, 20.5, 26.4, 29.7, 41.3, 63.5, 63.8, 108.3, 110.5, 114.3, 118.8, 121.4, 132.2, 138.9, 149.3, 154.6, 157.1. HRMS (ESI/Q-TOF): [M]^+^ Calcd for C_19_H_24_BrNO_2_ 378.1063; Found 378.1087.

#### 4.1.29. 5-Bromo-5’-ethoxy-2’-(isopropyl(methyl)amino)biphenyl-2-ol (**30**)^47^

4-Ethoxy-*N*-isopropyl-*N*-methylaniline (500 mg, 2.59 mmol) and 4-bromophenol (448 mg, 2.59 mmol) were reacted in toluene (5 mL) in the presence of AgNO_3_ (439 mg, 2.59 mmol) and H_2_O_2_ (880 mg, 7.76 mmol). The product was isolated by chromatography on a silica gel column using hexanes-ethyl acetate (9:1), and obtained as yellowish oil (323 mg, 34 %). ^1^H NMR (400 MHz, CDCl_3_): δ (ppm) 0.97 (6H, d, *J* = 6.4 Hz), 1.36 (3H, t, *J* = 7.2 Hz), 2.55 (3H, s), 3.52 (1H, m), 3.93 (2H, q, *J* = 7.2 Hz), 6.49 (1H, d, *J* = 2.4 Hz), 6.65 (1H, dd, *J* = 2.4 and 8.8 Hz), 6.80 (2H, d, *J* = 8.8 Hz), 6.97 (1H, d, *J* = 8.8 Hz), 7.35 (2H, d, *J* = 8.8 Hz). ^13^C NMR (100 MHz, CDCl_3_): δ (ppm) 14.9, 18.8, 32.5, 52.7, 64.0, 108.0, 110.0, 114.6, 119.0, 122.8, 132.2, 138.1, 149.9, 154.9, 157.0. HRMS (ESI/Q-TOF): [M]^+^ Calcd for C_18_H_22_BrNO_2_ 364.0907; Found 364.0909.

#### 4.1.30. 2’-(Benzyl(methyl)amino)-5’-ethoxybiphenyl-2-ol (**31**)^47^

*N*-Benzyl-4-ethoxy-*N*-methylaniline (500 mg, 2.07 mmol) and phenol (195 mg, 2.07 mmol) were reacted in toluene (5 mL) in the presence of AgNO_3_ (352 mg, 2.07 mmol) and H_2_O_2_ (705 mg, 6.22 mmol). The product was isolated by chromatography on a silica gel column using hexanes-ethyl acetate (9:1), and obtained as yellowish oil (274 mg, 40 %). ^1^H NMR (400 MHz, CDCl_3_): δ (ppm) 1.34 (3H, t, *J* = 7.2 Hz), 2.59 (3H, s), 3.91 (2H, q, *J* = 7.2 Hz), 4.12 (2H, s), 6.55 (1H, d, *J* = 2.4 Hz), 6.62 (1H, dd, *J* = 2.4 and 8.8 Hz), 6.93 (1H, d, *J* = 8.8 Hz), 6.97 (2H, d, *J* = 8.0 Hz), 7.05 – 7.11 (3H, m), 7.18 – 7.19 (3H, m), 7.29 (2H, dd, *J* = 7.6 and 8.4 Hz). ^13^C NMR (100 MHz, CDCl_3_): δ (ppm) 14.8, 39.8, 60.3, 63.9, 107.9, 109.6, 117.6, 121.1, 122.5, 126.8, 128.0, 128.7, 129.5, 137.9, 139.1, 149.7, 154.9, 157.7. HRMS (ESI/Q-TOF): [M + H]^+^ Calcd for C_22_H_24_NO_2_ 334.1802; Found 334.1814.

#### 4.1.31. 2’-(Benzyl(methyl)amino)-5-bromo-5’-ethoxybiphenyl-2-ol (**32**)^47^

*N*-Benzyl-4-ethoxy-*N*-methylaniline (500 mg, 2.07 mmol) and 4-bromophenol (358 mg, 2.07 mmol) were reacted in toluene (5 mL) in the presence of AgNO_3_ (352 mg, 2.07 mmol) and H_2_O_2_ (705 mg, 6.22 mmol). The product was isolated by chromatography on a silica gel column using hexanes-ethyl acetate (9:1), and obtained as yellowish oil (216 mg, 25 %). ^1^H NMR (400 MHz, CDCl_3_): δ (ppm) 1.36 (3H, t, *J* = 7.2 Hz), 2.56 (3H, s), 3.93 (2H, q, *J* = 7.2 Hz), 4.08 (2H, s), 6.61 (1H, d, *J* = 2.8 Hz), 6.65 (1H, dd, *J* = 2.8 and 8.8 Hz), 6.81 (2H, d, *J* = 8.8 Hz), 6.95 (1H, d, *J* = 8.8 Hz), 7.04 (2H, d, *J* = 7.6 Hz), 7.19 – 7.23 (3H, m), 7.38 (2H, d, *J* = 8.8 Hz). ^13^C NMR (100 MHz, CDCl_3_): δ (ppm) 14.8, 39.7, 60.1, 63.9, 108.3, 110.3, 114.6, 118.8, 121.2, 126.8, 128.0, 128.4, 132.3, 137.9, 138.5, 149.2, 154.8, 157.1. HRMS (ESI/Q-TOF): [M]^+^ Calcd for C_22_H_22_BrNO_2_ 412.0907; Found 412.0908.

#### 4.1.32. 5’-Ethoxy-2’-(ethyl(propyl)amino)biphenyl-2-ol (**33**)^47^

4-Ethoxy-*N*-ethyl-*N*-propylaniline (500 mg, 2.41 mmol) and phenol (227 mg, 2.41 mmol) were reacted in toluene (5 mL) in the presence of AgNO_3_ (410 mg, 2.41 mmol) and H_2_O_2_ (820 mg, 7.24 mmol). The product was isolated by chromatography on a silica gel column using hexanes-ethyl acetate (9:1), and obtained as yellowish oil (279 mg, 47 %). ^1^H NMR (400 MHz, CDCl_3_): δ (ppm) 0.77 (3H, t, *J* = 7.6 Hz), 0.94 (3H, t, *J* = 7.2 Hz), 1.35 (3H, t, *J* = 7.6 Hz), 1.39 (2H, m, *J* = 7.2 Hz), 2.93 (2H, m), 3.05 (2H, q, *J* = 7.2 Hz), 3.91 (2H, d, *J* = 7.2 Hz), 6.45 (1H, d, *J* = 2.8 Hz), 6.62 (1H, dd, *J* = 2.8 and 8.4 Hz), 6.92 (2H, d, *J* = 8.8 Hz), 6.99 (1H, d, *J* = 8.4 Hz), 7.04 (1H. d, *J* = 8.8 Hz), 7.27 (2H, t, *J* = 7.6 Hz). ^13^C NMR (100 MHz, CDCl_3_): δ (ppm) 11.7, 12.3, 14.8, 20.4, 47.5, 54.9, 63.6, 107.6, 109.5, 117.8, 122.3, 124.0, 129.4, 135.4, 152.0, 155.1, 157.7. HRMS (ESI/Q-TOF): [M + H]^+^ Calcd for C_19_H_26_NO_2_ 300.1958; Found 300.1976.

#### 4.1.33. 5-Bromo-5’-ethoxy-2’-(ethyl(propyl)amino)biphenyl-2-ol (**34**)^47^

4-Ethoxy-*N*-ethyl-*N*-propylaniline (500 mg, 2.41 mmol) and 4-bromophenol (417 mg, 2.41 mmol) were reacted in toluene (5 mL) in the presence of AgNO_3_ (410 mg, 2.41 mmol) and H_2_O_2_ (820 mg, 7.24 mmol). The product was isolated by chromatography on a silica gel column using hexanes-ethyl acetate (9:1), and obtained as yellowish oil (325 mg, 42 %). ^1^H NMR (400 MHz, CDCl_3_): δ (ppm) 0.75 (3H, t, *J* = 7.6 Hz), 0.91 (3H, t, *J* = 7.2 Hz), 1.33 – 1.38 (5H, m), 2.88 – 2.92 (2H, m), 3.00 (2H, q, *J* = 7.2 Hz), 3.93 (2H, q, *J* = 7.2 Hz), 6.49 (1H, d, *J* = 2.8 Hz), 6.65 (1H, dd, *J* = 2.8 and 8.8 Hz), 6.78 (2H, d, *J* = 9.2 Hz), 6.99 (1H, d, *J* = 8.8 Hz), 7.35 (2H, d, *J* = 9.2 Hz). ^13^C NMR (100 MHz, CDCl_3_): δ (ppm) 11.6, 12.2, 14.8, 20.5, 47.5, 54.8, 63.8, 108.0, 110.2, 114.5, 119.1, 123.9, 132.3, 135.6, 150.8, 155.1, 157.3. HRMS (ESI/Q-TOF): [M]^+^ Calcd for C_19_H_24_BrNO_2_ 378.1063; Found 378.1069.

#### 4.1.34. 5-Bromo-2’-(diethylamino)-5’-ethoxybiphenyl-2-ol (**35**)^47^

4- Ethoxy-*N*,*N*-diethylaniline (500 mg, 2.59 mmol) and 4-bromophenol (448 mg, 2.59 mmol) were reacted in toluene (5 mL) in the presence of AgNO_3_ (439 mg, 2.59 mmol) and H_2_O_2_ (880 mg, 7.76 mmol). The product was isolated by chromatography on a silica gel column using hexanes-ethyl acetate (9:1), and obtained as yellowish oil (432 mg, 46 %). ^1^H NMR (400 MHz, CDCl_3_): δ (ppm) 0.92 (6H, t, *J* = 7.2 Hz), 1.35 (3H, t, *J* = 7.2 Hz), 3.02 (4H, q, *J* = 7.2 Hz), 3.92 (2H, q, *J* = 7.2 Hz), 6.50 (1H, d, *J* = 2.8 Hz), 6.65 (1H, dd, *J* = 2.8 and 8.8 Hz), 6.80 (2H, d, *J* = 8.8 Hz), 7.00 (1H, d, *J* = 8.8 Hz), 7.35 (2H, d, *J* = 9.2 Hz). ^13^C NMR (100 MHz, CDCl_3_): δ (ppm) 12.3, 14.8, 46.9, 63.7, 107.9, 110.0, 114.5, 119.2, 124.0, 132.2, 134.9, 151.2, 155.2, 156.9. HRMS (ESI/Q-TOF): [M]^+^ Calcd for C_18_H_22_BrNO_2_ 364.0907; Found 364.0910.

### 4.2. In vitro biological evaluation

#### 4.2.1. Antibodies, chemicals and reagents

3-(4,5-Dimethylthiazol-2-yl)-2,5-diphenyltetrazolium bromide (MTT), dimethyl sulfoxide (DMSO) and 4′,6-diamidino-2-phenylindole (DAPI-#DUO82040), *in situ* mounting media and bovine serum albumin (BSA) were purchased from Sigma–Aldrich Chemical Company (St. Louis, MO, USA) and MP Biomedicals (Irvine, California, USA), respectively. Antibodies for immunoblotting and immunocytochemistry, including caspase 3 (9662S), caspases 9 (#9502), Bcl-2 (2876S), Bax (2772S), β-Actin (13E5) Rabbit mAb #4970, pS6 (Ser235/236) #4858, a pan antibody mix of Phospho-Akt (Ser473)(D9E) XP® Rabbit mAb #4060, Phospho-p90RSK, Phospho-p44/42 MAPK(Erk 1/2) (Thr202/Tyr204), PARP Antibody #9542, Phospho-S6 Ribosomal Protein Detection Cocktail I (5301S), and horseradish peroxidase-conjugated (HRP) anti-mouse and anti-rabbit secondary antibodies were all obtained from Cell Signaling Technologies (Beverly, MA, USA). Mini-protean precast Tris-Glycine Gels (TGX) were from Bio-Rad (Bio-Rad Laboratories Inc., Hercules, California, USA). Radioimmunoprecipitation assay (RIPA) buffer and Pierce Bicinchoninic acid (BCA™) protein assay kit were purchased from ThermoFisher Scientific (Rockford, Illinois, USA). The SuperSignal™ West Pico PLUS Chemiluminescent substrate detection system was from ThermoFisher Scientific (Waltham, Massachusetts, USA). A 2% (w/v) Aqueous Solution of Gentian Violet and crystal violet were from Ricca Chemical Company (Arlington, TX, USA). Dulbecco’s modified Eagle’s medium (DMEM) and Roswell Park Memorial Institute Medium (RPMI 1640) were from Corning (Corning, Manassas, VA, USA). Epi-Life® Growth Medium with 60 µM calcium and Cascade Biologist human keratinocyte growth supplement (HKGS) 100X (S-001–5) were from Life Technologies (Grand Island, NY, USA). The Dulbecco’s phosphate-buffered saline (DPBS), phosphate buffer saline (PBS) 1X, defined trypsin inhibitor (DTI) 1X (R007-100), trypsin EDTA 0.25%, 1X (R25200-072), Trypsin neutralizer (TN) 1X (R-002–100), penicillin-streptomycin (Pen Strep,15140–122) (PEST) 100X were purchased from Gibco, ThermoFisher Scientific (Rockford, IL, USA), and human keratinocyte growth supplement (HKGS) 100X (S-001–5) from Cascade Biologist were procured from VWR Corporation (Missouri, Texas, USA). USDA-approved Origin Fetal Bovine Serum (FBS) was procured from VWR Seradigm Life Science (Missouri, Texas, USA). The organic solvents including, ethanol (6183– 10) and methanol (BDH 1135-4LP), were acquired from Macron Chemicals (Radnor, Pennsylvania, USA) and VWR (Missouri, Texas, USA), respectively.

#### 4.2.2. Cell lines, cell cultures, cytotoxicity and viability evaluation assay

Human-derived GFP-expressing melanoma A375, and epidermoid carcinoma A431 cell lines were purchased from Angio-Proteomie (Boston, MA, USA). Human melanoma carcinoma SK-Mel-28 and Human immortalized keratinocytes (HaCaT) cell lines were acquired from American Type Culture Collection (ATCC; Manassas, VA, USA). Human cutaneous SCC cell line SCC-12 was generously provided by Dr. James G. Rheinwald^108^ to Dr. Tatiana Efimova. Except SK-MEL-28, which was cultured in an RPMI-1640 medium supplemented with 5% FBS, all other human immortalized cell lines were grown and maintained in DMEM containing varying percentages of FBS; 5% FBS for A431, SCC-12 and HaCaT lines, while A375 was grown in 10% FBS. All culture media were routinely supplemented with 1% PEST 100X; 100U/ml penicillin, 100 μg/ml streptomycin, and cells were cultured in incubators maintained at 37 °C under an atmosphere of 95% humidity, 20% O_2_ and 5% CO_2_. The growth media of incubated cells were changed every 2-3 days until they attained 70-80% confluence, after which they were sub-cultured for experiments and/or re-passaging. DMSO was used as vehicle to prepare a 10 mM stock solution of the test compounds. Control cells were treated with the vehicle (DMSO) at concentrations (0.01-0.2%), not affecting cell viability.

Stock solutions (10 mM) were prepared by dissolving each compound to be screened in DMSO. From these 10 mM stock solutions, escalating concentrations were prepared using the corresponding cell growth media as a diluent, and the cells (65-75% confluent) were treated with (0-40 µM) or without the drug in sex-to -octuplicate and incubated for 48 h. All treatment protocols and controls were prepared as previously described.^25^

The MTT assay method was performed to assess the potency and cytotoxic influence of the test compounds against two cutaneous melanoma (A375 and SK-MEL-28) and two non-melanoma (A431 and SCC-12) skin cancer cell-lines in addition to a control immortalized normal human epidermal keratinocytes (HaCaT) cell line. Briefly, exponentially growing parent passage cells at 70-85% confluency were harvested, washed, counted using a hemocytometer, and individually seeded at a density of 2-5×10^3^ Cells/well in a 96 well plate in 200 µL of their respective growth media, incubated and monitored until reaching their log growth phase at over 70% confluent prior to treatment. All experimental treatment concentrations were repeated at least in 8-10 wells, and the experiment was repeated 6-8 times. Subsequently, the cells were harvested following the MTT protocol essentially as previously described,^41^ with absorbance measured at 562 nm wavelengths using a SynergyLX, multimode microplate reader (BioTek Instruments Inc., Winooski, VT, USA). The results from escalating concentration of each test and control (cisplatin) agent were analyzed using the zero drug control groups as the relative comparator and are expressed as a percentile following the equation: [(absorbance in treatment group/ absorbance in control) x 100%]. GraphPad Prism software version 9.3 (GraphPad Software, Inc., La Jolla, CA, USA) was then used to compute the various IC_50_ values as earlier described.^25^

#### 4.2.3. Colony formation assay

This assay was performed to demonstrate the clonogenic potential of the active compounds on SCC-12 and SKMeL-28 cell lines, following a previously described protocol.^25^ Briefly, 3000 viable cells treated with compounds **11** or **13** at varying concentrations (0, ½IC_50_, IC_50_ and 1½IC_50_) for 48 h in a T-25 flask were harvested and seeded into a 10 cm^2^ plate in quadruplicates in respective drug free growth medium. This was monitored with routine media change for 10-14 days until control cells reached about 90% confluency or cell plates formed about 50-150 colonies. The cell plates were then harvested by washing twice in 1XPBS, followed by fixing and staining in a submerged solution containing 0.5% gentian violet staining agent in 64% (v/v) methanol and 4% paraformaldehyde (4% PFA) for 45 to 60 minutes at room temperature. Stained cells were gently washed in tap water until the solution became clear and were air-dried and imaged with a Nikon D7500 camera. Colonies were counted and analyzed using the count and Plot Histograms of Colony Size (countPHICS) analysis tool^99^ and GraphPad Prism software version 9.1 (GraphPad Software, Inc., La Jolla, CA, USA), respectively. Statistical analyses data showed the mean ± SD of three independent experiments, comparing each treatment group to the DMSO group: * P < 0.05, ** P < 0.01, *** P < 0.001.

#### 4.2.4. Immunofluorescence Staining and immunocytochemical analysis of cells morphology by Actin and apoptosis by Caspase-3 activation

Immunofluorescence analysis was employed to detect morphological and biochemical changes associated with the expression of apoptosis marker (Caspase 3) and the cell actin cytoskeleton reorganization following cells treatment. Briefly, cells pre-seeded into an 8-well chamber slide were treated with escalating concentrations (0, ½IC_50_, IC_50_ and 1½IC_50_) of the test compounds **11** or **13** for 48 h. The cells were washed and fixated using 4% PFA in PBS pH 7.4 for 20 to 30 min at room temperature, followed by permeabilization in 0.1% Triton X-100 stabilization buffer for 5 minutes, washed three times with cold 1xPBS, and non-specific binding was reduced using blocking buffer with 1%BSA for 30 min - 1 h. The cells were added to the respective primary antibodies [β-Actin or Caspase-3 at (1:1000 dilution)] for fluorescence staining, and cell slides were incubated overnight at 4 °C. Slides were subsequently washed thrice, and incubated with Texas Red@-X Goat Anti-mouse (dilution 1:600) and Alexa fluor Goat Anti-rabbit (dilution 1:600) for 2 h at room temperature. Slides were re-washed, and mounted using in situ mounting media containing phalloidin and 4′,6-diamidino-2-phenylindole (DAPI). The mounted slides were visualized and imaged using the inverted fluorescence microscope Olympus IX71 equipped with a DP71 digital camera at 10x and 1.6×10x magnification. The images captured were processed using the Cellsens 6 software and each experiment was performed three times.^41^

#### 4.2.5. In vitro wound healing assay

The scratch wound healing assay was performed to evaluate compounds **11** or **13**’s ability to limit the migration and wound closure using a previously described protocol.^25^ Briefly, a 24 well plate seeded with cells was incubated until a fully confluent monolayer of cells was scratched a straight line across the center using a sterile 200 µL Gilson pipette tip. Cells were washed gently to remove debris and detached cells and treated with **11** or **13** at escalating concentrations (0, ½IC_50_, IC_50,_ and 1½IC_50_). At times 0 h, 24 h, and 48 h, the wounded and re-epithelialization areas were imaged using an inverted phase-contrast microscope Olympus IX51 (Olympus America Inc., Center Valley, PA, USA) attached to an infinity 2 digital camera processed on an infinity analyze software version 6.5.6 (Teledyne Lumenera, Ottawa, Canada) and analyzed with Adobe Photoshop. The ImageJ software (NIH, Bethesda, MD) was then used to measure the migratory area of each treatment group at 24 h and 48 h, and are expressed as a percentage of wound surface at time 0 h.^41^

#### 4.2.6. Preparation of cell lysates, Western blot assay and analysis

Cell lysate preparation and immunoblotting were performed. Briefly, lysates were prepared after subjecting 3×10 cm^2^ plates seeded with about 3000 cells each and treated with varying concentrations (0, ½IC_50_, IC_50_ and 1½IC_50_) of compounds for 48 h, following the reference protocol.^25^ After treatment with compound **11** or **13**, the cell growth media were aspirated, and the cells were harvested by washing in cold 1xPBS (pH 7.4), and whole-cell, cytosolic and nuclear lysates were prepared as described earlier.^25^ Cells were lysed by incubating in cell lysis RIPA buffer supplemented with protease and phosphatase inhibitors; 1 mM PMSF (phenylmethylsulfonyl fluoride), 10 mM RIPA buffer, 1 mM protease inhibitor cocktail (1 mM EDTA, 20 mM NaF, 0.5% NP-40, 1% Triton X-100) (Roche) and 0.5% Na_3_VO_4_ on ice. The homogenate was then centrifuged at 14,000× g for 25 min at 4 °C, the supernatant was collected and the cleared lysate protein concentration was quantified using the BCA protein assay kit (Thermo Scientific, Pierce Rockford, IL) according to the manufacturer’s protocol. The assay was performed using the Eppendorf BioPhotometer D30 and aliquoted and was immediately used or stored at − 80 ◦C for later use for western blot assay for further analysis as earlier described.^25^

Western blotting and analysis were performed to identify the potential mechanisms and targeted pathways accounting for the observed biological responses for compounds **11** or **13**, following a previously described protocol.^25, 41^ Western blot analysis was performed; accordingly, lysates were denatured in 2X Laemmli sample buffer and loaded (to resolve the equal amount of protein-25g) per lane using a KD 4–12% polyacrylamide ready mini-PROTEAN TGX gels electrophoresis, which causes separation based on molecular weight. The resolved proteins were transferred into nitrocellulose filter membrane using a Trans-Blot Turbo Transfer Pack. The presence of proteins was confirmed by using a ponceau staining before proceeding to washes (2X) under tap water and TBS-T wash buffer. Unspecific epitopes in the membrane were blocked with 5% BSA (for phospho-protein targets) or 5% non-fat dry milk/1% Tween 20; in 20 mM Tris-buffered saline (TBS or wash buffer) at room temperature on a rotating shaker for 45 min. The membranes were incubated overnight at 4 °C or 2 h at room temperature in the corresponding monoclonal or polyclonal primary antibodies (1:250 to 1:1000 dilution). Blots were then washed three times (5 min each) with PBST and incubated in anti-mouse or anti-rabbit horseradish-peroxidase (HRP) conjugated secondary antibody (1:2000 dilution). Afterward, blots were re-washed three times (5 min each), exposed to enhanced chemiluminescence plus west pico for 5 min and autoradiographed in a BioRad Gel-Doc System (Bio-Rad Laboratories Inc., Hercules, CA, USA). The loading consistency was verified utilizing the house-keeping proteins (Rab11 and β-Actin) by stripping and re-probing blots with monoclonal β-actin primary antibody (Sigma, St Louis, MO) following the similar procedure as described earlier.^25, 41^ The bands were then visualized and imaged using the Bio-Rad ChemiDoc™ detection system. Densitometry was done using Quantity One, the BioRad digitized scientific imaging analysis software system program. The Rab11 and β-actin data were used as a normalization factor and the linear range of the band densities were confirmed through multiple exposures of blot, and the analysis was based on three different experiments. As described earlier, The final data were analyzed for statistical differences using one-way ANOVA by the Graphpad Prism software.^41^

### 4.3. *In silico* screening assays

#### 4.3.1. Target prediction and acquisition methodology

The SwissTargetPrediction database (http://www.swisstargetprediction.ch/) web server, which predicts the targets of bioactive molecules based on a combination of two-dimensional (2D) and 3D similarity measures with known ligands,^58, 109, 110^ was employed to screen and estimate the putative macromolecular targets of all the 90 derivatives synthesized. Briefly, these compounds’ respective canonical simplified molecular input-line entry system (SMILES) structure formats were individually converted using the ChemDraw tool prior to importing into the SwissTargetPrediction network database to be queried. After the SMILES structures were uploaded to the database, the species “Homo sapiens” was selected, and a report of probable matched targets data was obtained. During target prediction, derivatives that possessed the highest probability to regulate identified targets under the consideration of probability >0.112, and the specific targets were selected based on values on predicted scores. The structure of the targets was retrieved from Protein Data Bank (PDB), and all PDB codes are available in the master excel sheet as supplementary data.

#### 4.3.2. In Silico molecular docking simulation methodology

Molecular docking was used in this study to predict the binding interactions between ligands and their protein targets by calculating and predicting the conformation and directions of ligands at the active sites of proteins.^111^ The credibility of the connection between the identified targets and the molecules screened was evaluated by docking each compound with the predicted target proteins. The molecular docking data were evaluated by selecting the known natural and clinical ligands of the studied targets and docking them as the positive longitudinal control. First, the SMILES structures of the commercially available clinical ligands used in the study were retrieved from the PubChem network database (https://pubchem.ncbi.nlm.nih.gov/),^112^ together with those predicted for compounds to be screened, and the SMILES structures were converted into 3D conformation using OpenBabel. The suitable 3D crystallographic structure of the protein targets was retrieved from the RCSB Protein Data Bank (PDB) web (https://www.rcsb.org/),^113^ were predicted, and retrieved proteins (receptors) were in complex with other hetero-atoms and water molecules. After downloading the 3D PDB file of each of the structures of the targets, PyMOL was used to process it by removing the pre-complex water molecules, hetero-atoms and separating the structure of the protein and ligand (inhibitor) from the protein’s-natural ligand complex alone. The separated natural inhibitor was considered a natural ligand in the docking separately to analyze the data. Both were then converted to PDBQT files using AutoDock Tools (ADT) that comes from an open-source software suite (the MGL tools 1.5.6)^114^ from Molecular Graphics Laboratory package from The Scripps Research Institute (http://autodock.scripps.edu/). During processing (to prepare the file suitable for docking), polar hydrogen was added to the separated protein structure before converting to PDBQT file. There was no need to add polar hydrogen in any ligands, but the torsion angle of each ligand was defined. While OpenBabel software (v2.2.1) (https://pyrx.sourceforge.io/) was used for converting the SMILES format of all the ligands to generate their 3D conformations in PDB format, the program Raccoon^115^ was used to add the torsion angle that is important for docking, and once the torsion angle is found, was converted and saved in PDBQT file format like target, and was later used for docking as the ligand.

The grid box was set centered in the cartesian coordinate x-, y-, and z-directions of the docking site defined according to the pre-complexed ligand (natural ligand) of PDB files of each target after removing water molecules and hetero atoms for each of the natural ligand in its receptor, with a default grid point spacing of 0.375 Å. After preparing all the ligands and targets, the binding efficiency was predicted using the free accessible graphic user interface (GUI) AutoDock Vina software (v.1.2.0.)^116^ in Linux OS. After docking simulation, 10 other conformations of the docked ligands were obtained, each ligand with the minimum binding energy and RMSD value was chosen. The ligand and the target complexes (ligand-protein interactions) were then visualized using PyMOL,^117^ and processed using Discovery Studio (version 3.5, Dassault Systèmes BIOVIA, San Diego). During the docking process, the PDBQT files of these structures were used in AutoDock Vina software, as these files contained the information of potential ligands and receptors required for the process. After obtaining docking results and the binding energy of each ligand and receptors using AutoDock Vina, the results were output in PDBQT files and were uploaded to PyMOL software to establish the 3D data binding models. Binding energy was used as a docking score to evaluate the protein-ligand binding potential of the derivatives to the identified targets. Those results with value ≤ −8 were selected and considered to have a moderate binding potential and a stable combination irrespective of the cutoff reads of the natural and clinical ligands. The lower the binding energy (binding energy < -8.00 kcal/mol) indicates good binding strength, sufficient to move forward with the predicted target for each of the derivative.

#### 4.3.3. In silico drug-likeness, pharmacokinetics and toxicity predictions

SwissADME was used to predict the ADME of each compound, where gastrointestinal absorption (GA), drug-likeness (DL), and other parameters were considered to define the bioactive compounds.^58, 110^ The SwissADME online server predictor estimate information on the *in silico* pharmacokinetic properties, druglike nature, the logarithm of the partition coefficient (milog P), number of hydrogen-bond donors (HBDs), number of hydrogen-bond acceptors (HBAs), total topological polar surface area of the compound (TPSA), number of rotatable bonds (Nrotbs), and Lipinski’s rule of five for drug likeliness.^58, 110^

#### 4.3.4. Statistical analysis

As previously described, all statistical analyses were performed using GraphPad Prism Software version 9.1 (GraphPad Prism Inc., San Diego, CA, U.S.A.).^25^ All quantitative and grouped data are expressed and presented as means ± standard deviation (SD) or ±, standard error of the mean (SEM) from at least three independent experiments. The significance was analyzed by one-way ANOVA, Student t-test or ANOVA with Dunn’s multiple comparison or Bonnferoni’s post hoc tests used to determine the difference between two or more groups. The following symbols represent the levels of statistical significance within each analysis: *\*p-value <0.05, **p-value<0.01, ***p-value <0.001* and ****p-value <0.001.

## Supporting information

Supplemental data on structure and characterization of synthesized compounds

## Acknowledgments

The authors acknowledge the financial support of NSF CHE-1954734 (awarded to JF). The following grants and funding partially supported this work: a Start-up fund from the University of Louisiana at Monroe (ULM) College of Pharmacy (COP), and a Faculty Research Seed grant #5CALHN-260615 awards from the ULMCOP; a Pilot and a full research project awards from a Louisiana Biomedical Research Network (LBRN)-IDeA Networks of Biomedical Research Excellence (INBRE) award from the National Institute of General Medical Sciences of the National Institutes of Health (NIGMS/NIH) grant number P2O GM103424-18, an LBRN-INBRE-COBRE Administrative Supplement Award from NIGMS/NIH grant 3P20GM103424-18S1, and a Louisiana Board of Regents Support Fund grant LEQSF (2021-24)-RD-A-22 (awarded to JCC). The content is solely the responsibility of the authors and does not necessarily represent the official views of the funding agencies.

## Appendix A

Supplementary data to this article can be found online at

## Abbreviations used

ADMET: absorption, distribution, metabolism, excretion, and toxicity
BCC: basal cell carcinoma
*CDK*: cyclin-dependent kinases
*COX*: cyclooxygenase
CDK16: cyclin-dependent kinase 16
DMEM: Dulbecco’s modified eagle medium
DMSO: dimethyl sulfoxide
EGCG: epigallocatechin-3-gallate
FBS: fetal bovine serum
HPLC: high performance liquid chromatograph
HRMS: high-resolution mass spectrometry
IC_50_: half-maximum inhibitory concentration
IHC: immunohistochemical
JAK: janus kinase
LC-MS/MS: liquid chromatography coupled to tandem mass spectrometry
MAPK: mitogen-activated protein kinase
*MDM2*: *m*ouse double minute 2 homolog
*MMP9*: matrix metallopeptidase 9
MS: mass spectrometry
*mTOR*: mammalian target of rapamycin
MTT: 3- (4,5- dimethyl-2-thiazolyl)-2,5-diphenyl-2-H-tetrazolium bromide
*NAD*: nicotinamide adenine dinucleotide
NAMPT: nicotinamide phosphoribosyltransferase
NF-κB: nuclear factor κ-light-chain-enhancer of activated B cell
NMR: nuclear magnetic resonance
*PARP*: poly (ADP-ribose) polymerase
PBS: phosphate buffered saline
PI3K: phosphoinositide 3-kinase
SAR: structure-activity relationship
SCC: squamous cell carcinoma
SD: standard deviation
TGF-β: transforming growth factor β
TLC: thin layer chromatography
STAT: signal transducer and activator of transcription.

Ref 107: Stern RS. Prevalence of a history of skin cancer in 2007: results of an incidence-based model. Arch Dermatol. 2010 Mar;146(3):279-82.

