## Supplemental data on structure and characterization of synthesized compounds for "Identification of potential inhibitors of cutaneous Melanoma and Non-Melanoma skin cancer cells through in-vitro and in-silico screening of a small library of Phenolic compounds"

*Methyl 2,6-dibromo-3,4,5-trimethoxybenzoate (**1**)* – $^{1}$H NMR ($CDCl_3$, 400 MHZ)

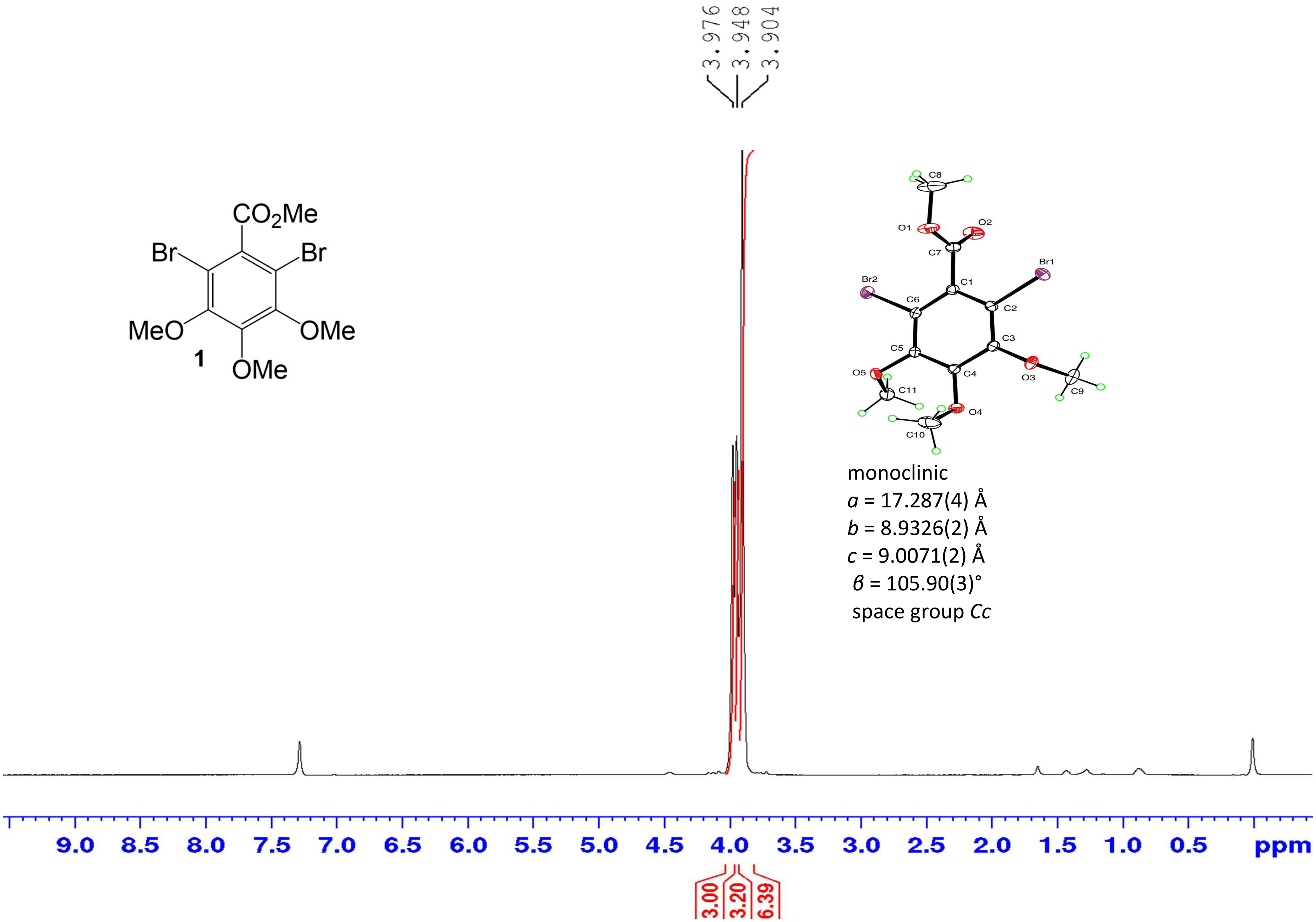

*Methyl 2,6-dibromo-3,4,5-trimethoxybenzoate (**1**)* – $^{13}$C NMR ($CDCl_3$, 100 MHZ)

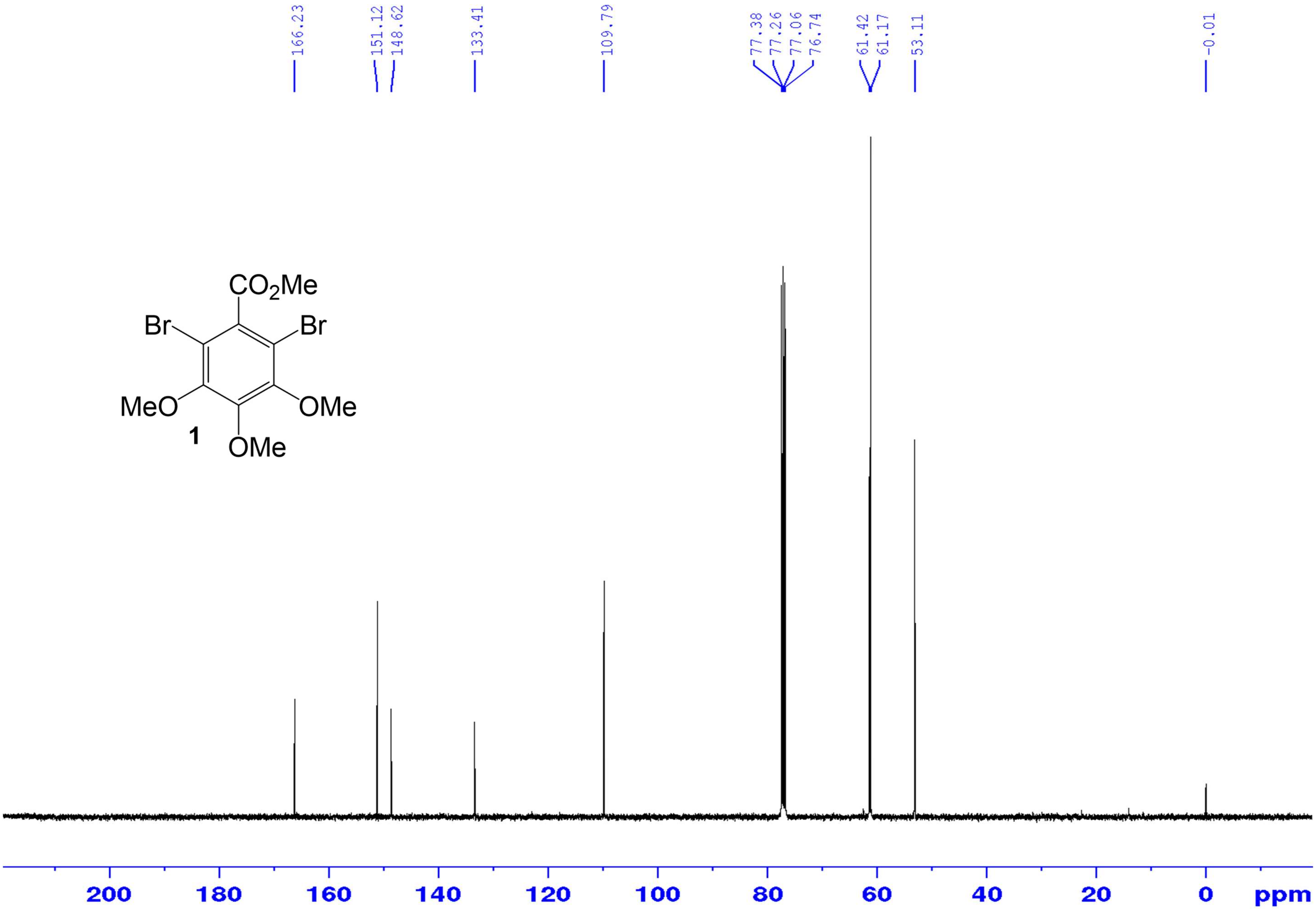

*Methyl 3,4,5-trimethoxy-2-nitrobenzoate* (***2***) – $^{1}$H NMR ($CDCl_3$, 400 MHZ)

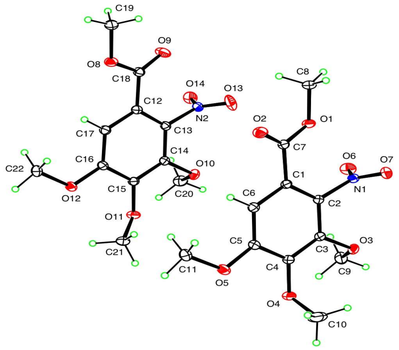

monoclinic
*a* = 20.520 (4) Å
*b* = 9.7438 (2) Å
*c* = 24.434 (5) Å
*β* = 99.46 (1)°
space group *C*2/*c*

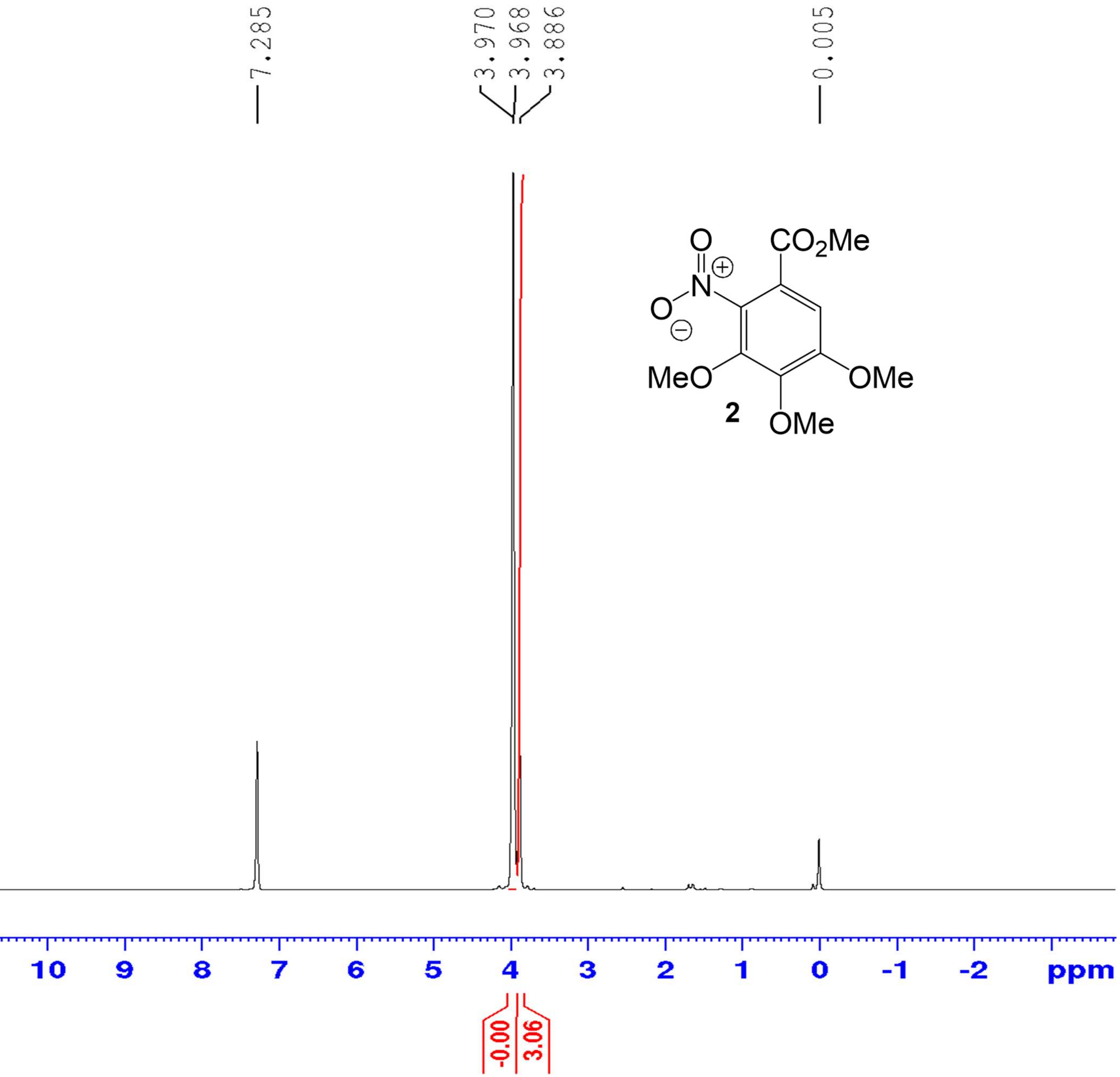

*Methyl 3,4,5-trimethoxy-2-nitrobenzoate (**2**)* – $^{13}$C NMR ($CDCl_3$, 100 MHZ)

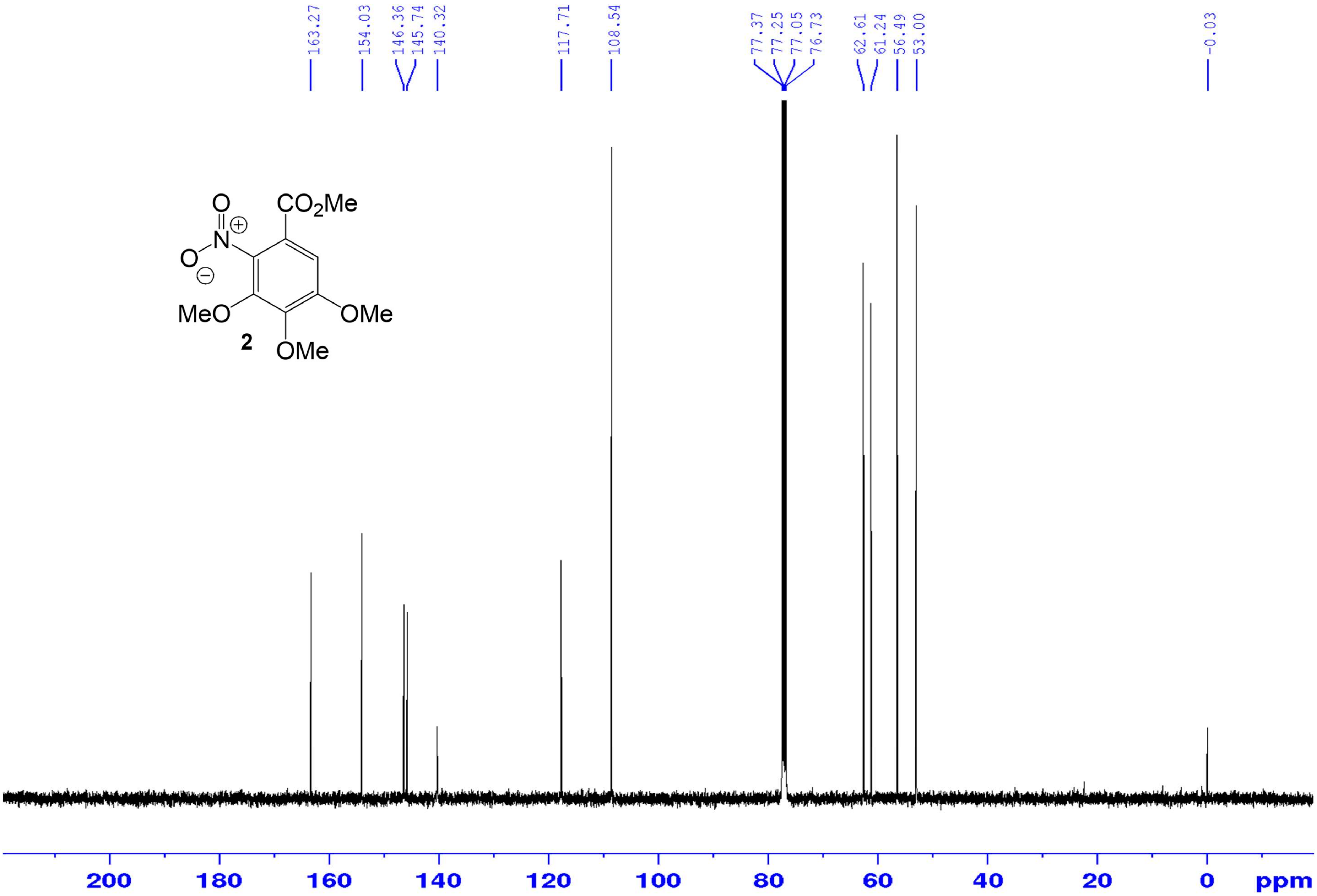

*4-Cyclohexenyl-N,N-dimethylaniline (**3**)* – $^{1}$H NMR ($CDCl_3$, 400 MHZ)

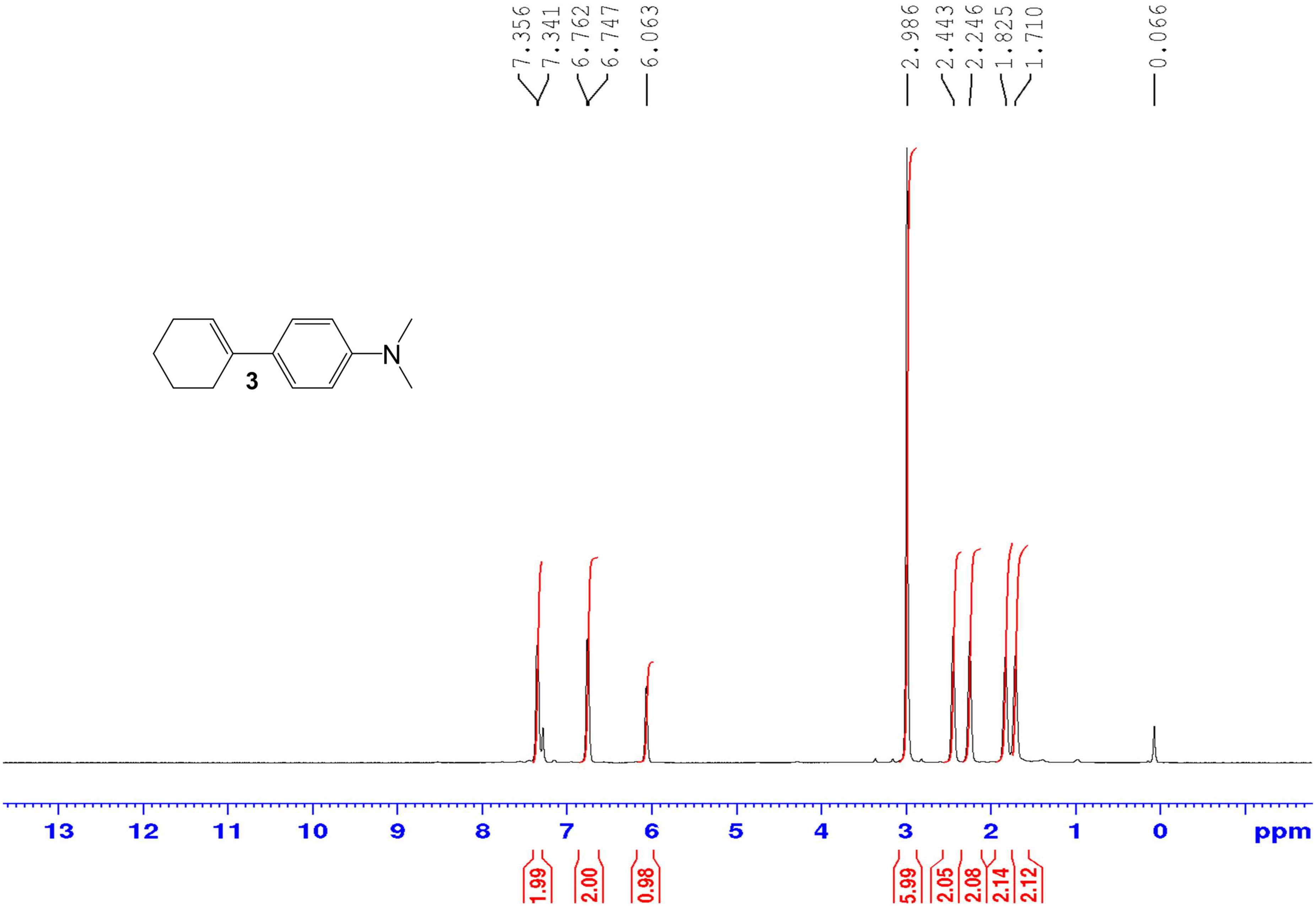

*4-Cyclohexenyl-N,N-dimethylaniline* (**3**) – $^{13}$C NMR (100  MHz)

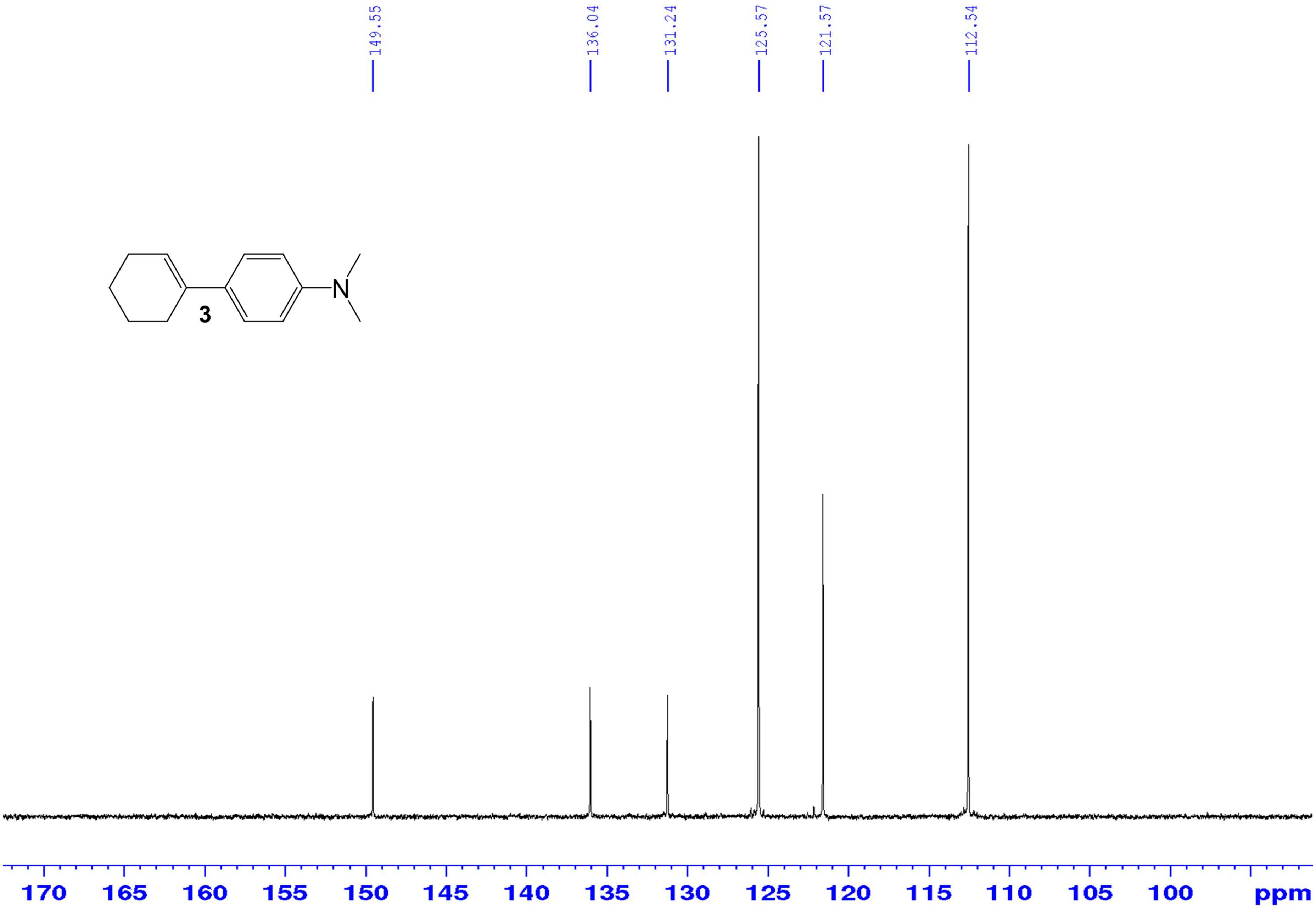

*4-Cyclohexenyl-N,N-dimethylaniline (**3**)* – HR-ESIMS ($[M+H]^+$ Calculated: 202.159026, Observed: 202.1621)

A) Mass spectrum (m/z range 160 to 350)

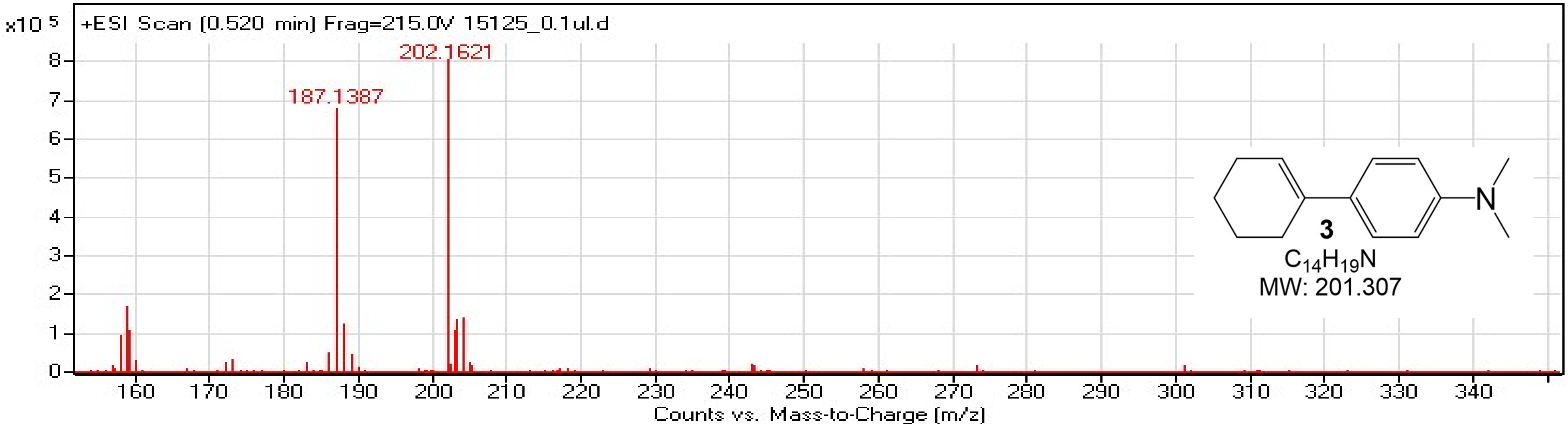


B) Mass spectrum (m/z range 200 to 207)

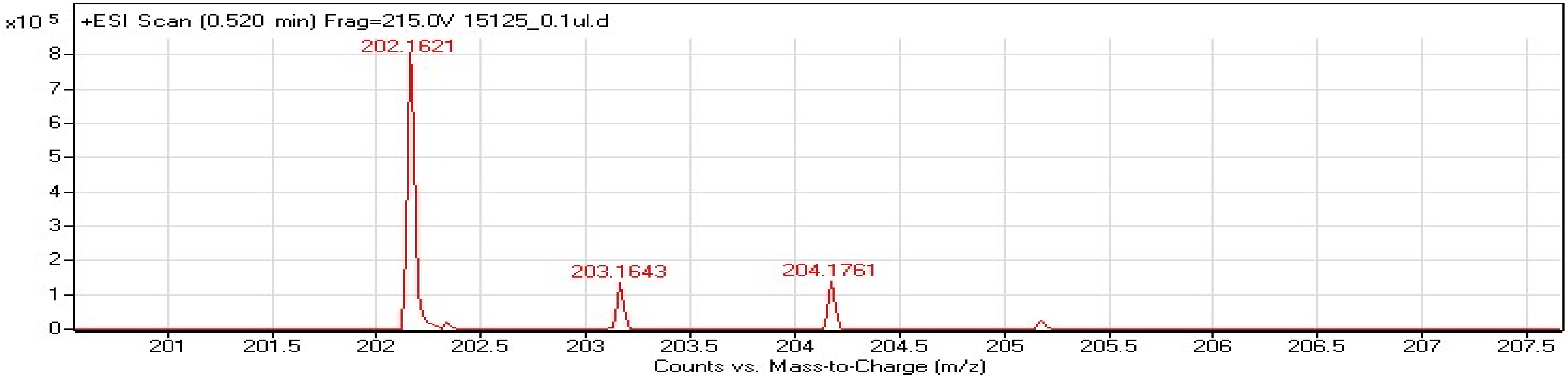


| Formula | Calculated Mass | mDa Error | ppm Error | RDB |
|---|---|---|---|---|
| C14 H20 N | 202.159026 | 3.073772 | 15.204451 | 5.5 |

*2-Chloro-4-cyclohexenyl-N-ethylaniline (**4**)* – $^{1}$H NMR ($CDCl_3$, 400 MHz)

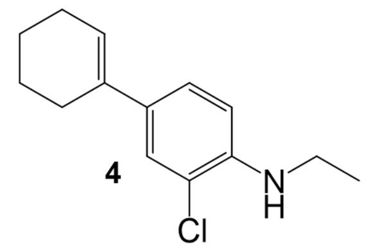


**4**

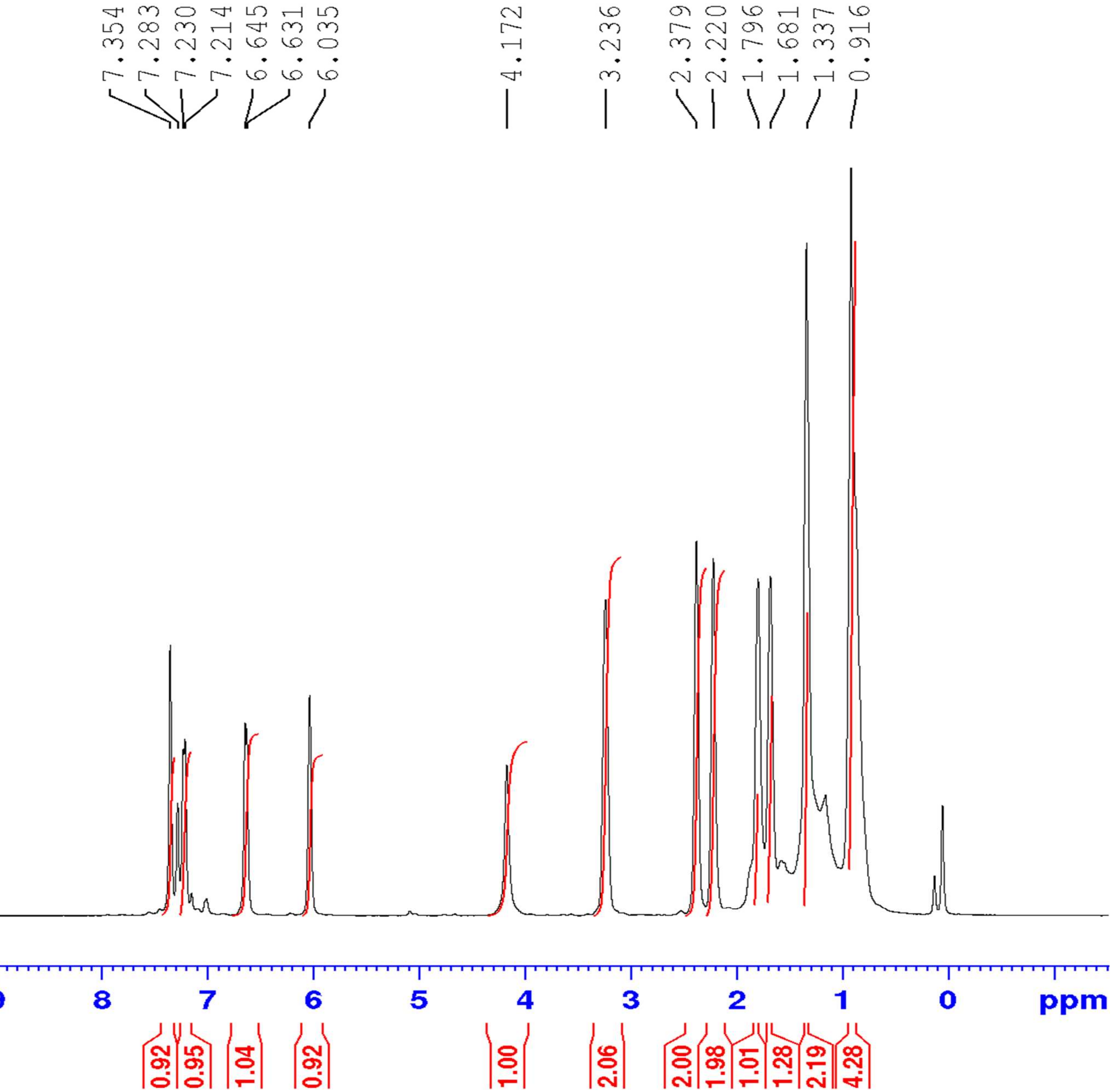

*2-Chloro-4-cyclohexenyl-N-ethylaniline (**4**)* – $^{13}$C NMR ($CDCl_3$, 100 MHz)

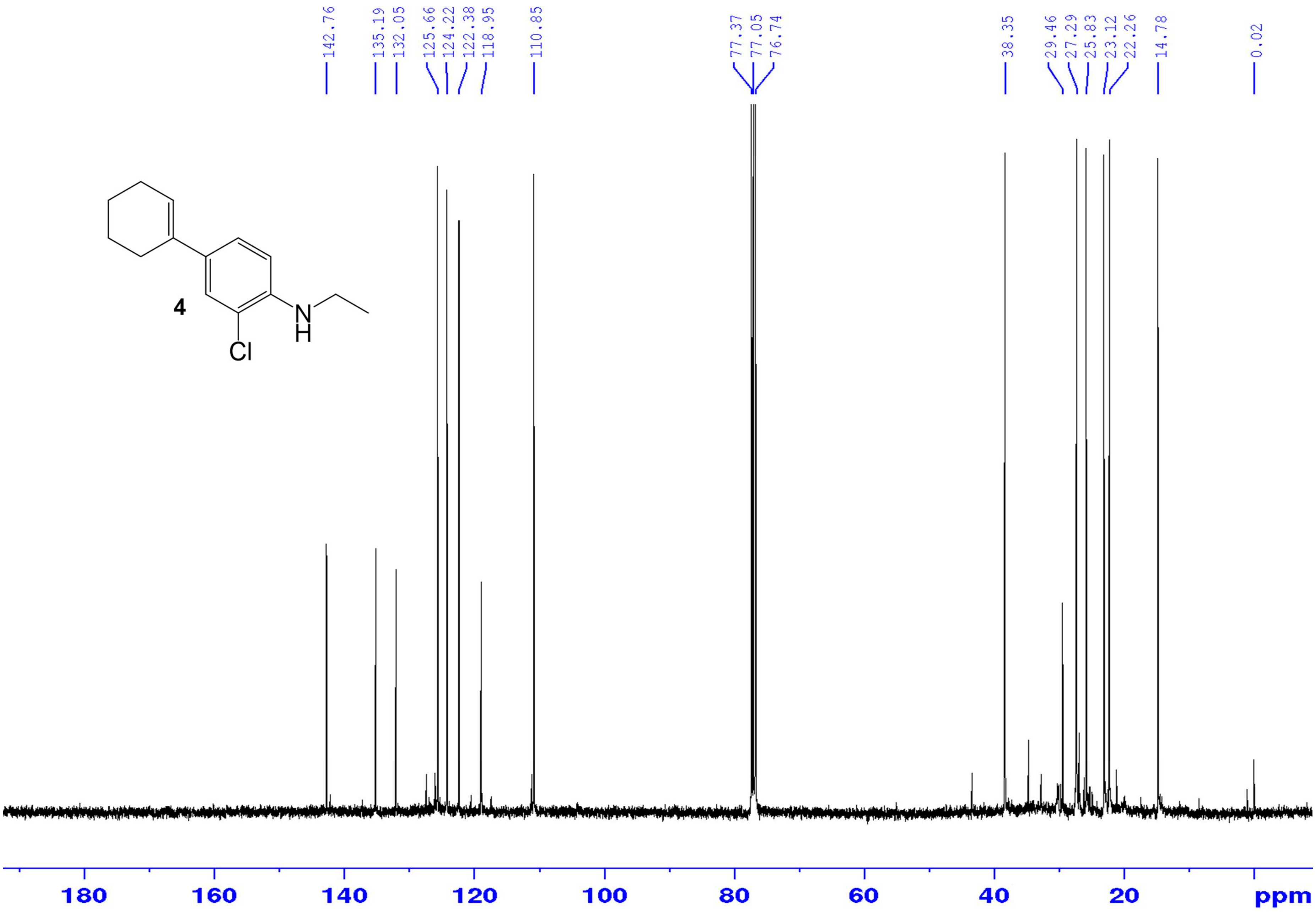

*2-Chloro-4-cyclohexenyl-N-ethylaniline (**4**)* – HR-ESIMS ([M]$^+$ Calculated: 235.1123, Observed: 235.1121)

### Single Mass Analysis

#### Tolerance = 5.0 PPM / DBE: min = -1.5, max = 50.0
#### Isotope cluster parameters: Separation = 1.0 Abundance = 1.0%

Monoisotopic Mass, Odd and Even Electron Ions
92 formula(e) evaluated with 1 results within limits (up to 50 closest results for each mass)

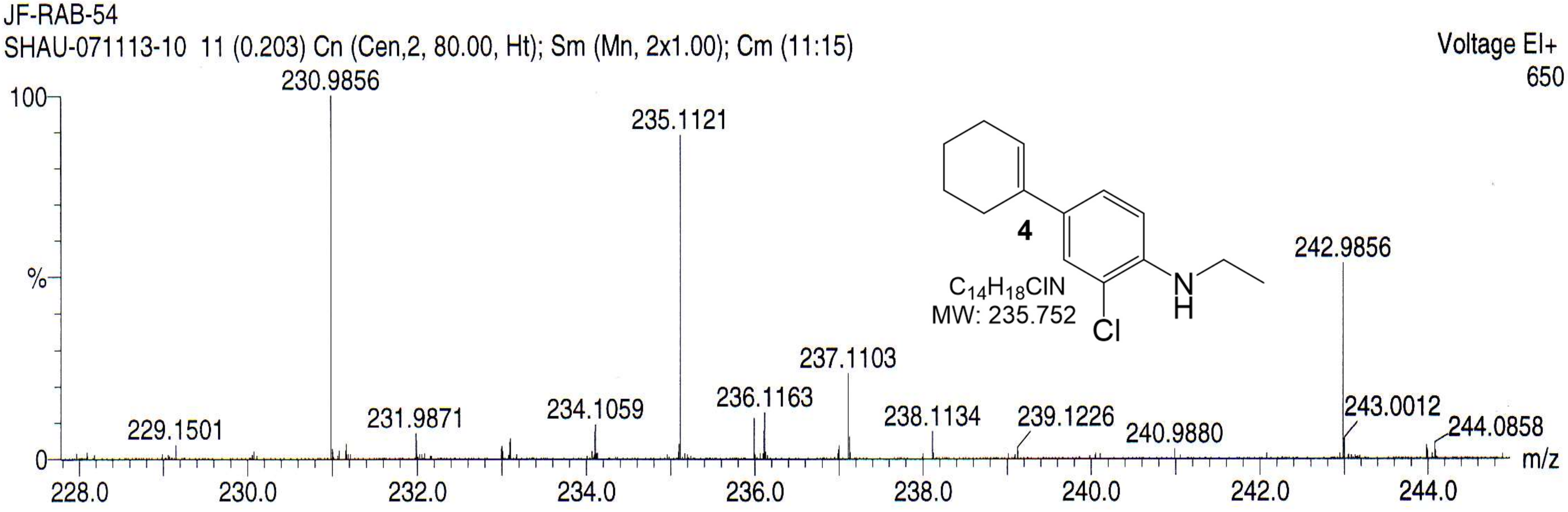


| | | | | | | |
|---|---|---|---|---|---|---|
| Minimum: | | | | -1.5 | | |
| Maximum: | | 3.0 | 5.0 | 50.0 | | |
| Mass | Calc. Mass | mDa | PPM | DBE | Score | Formula |
| 235.1121 | 235.1123 | -0.2 | -0.8 | 10.5 | 1 | C17 H15 O |

*6-Ethoxy-2,2,4-trimethyl-1,2-dihydroquinoline (**5**)* – $^1$H NMR ($CDCl_3$, 400 MHZ)

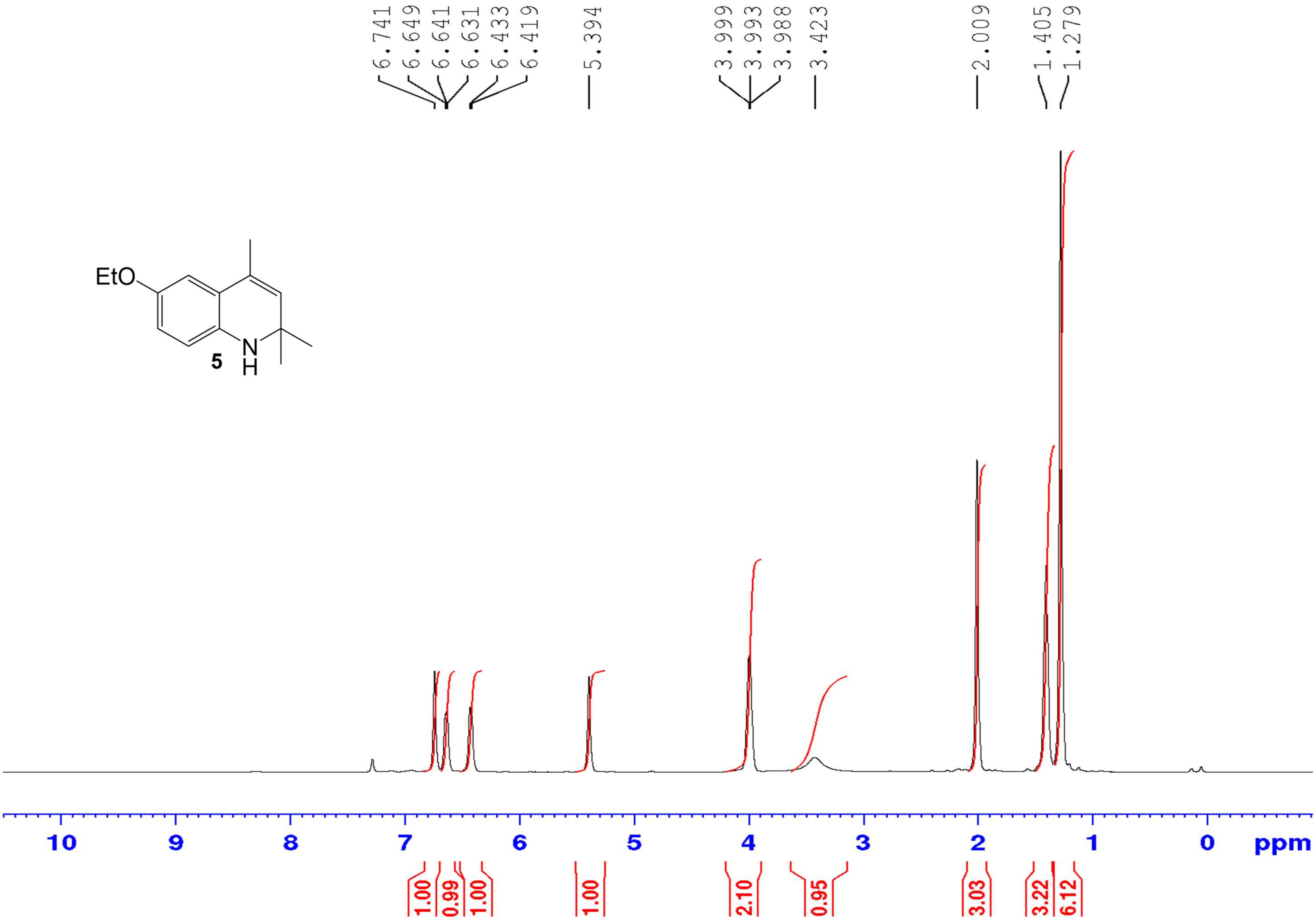

*6-Ethoxy-2,2,4-trimethyl-1,2-dihydroquinoline (**5**)* – $^{13}$C NMR (CDCl$_3$, 100 MHZ)

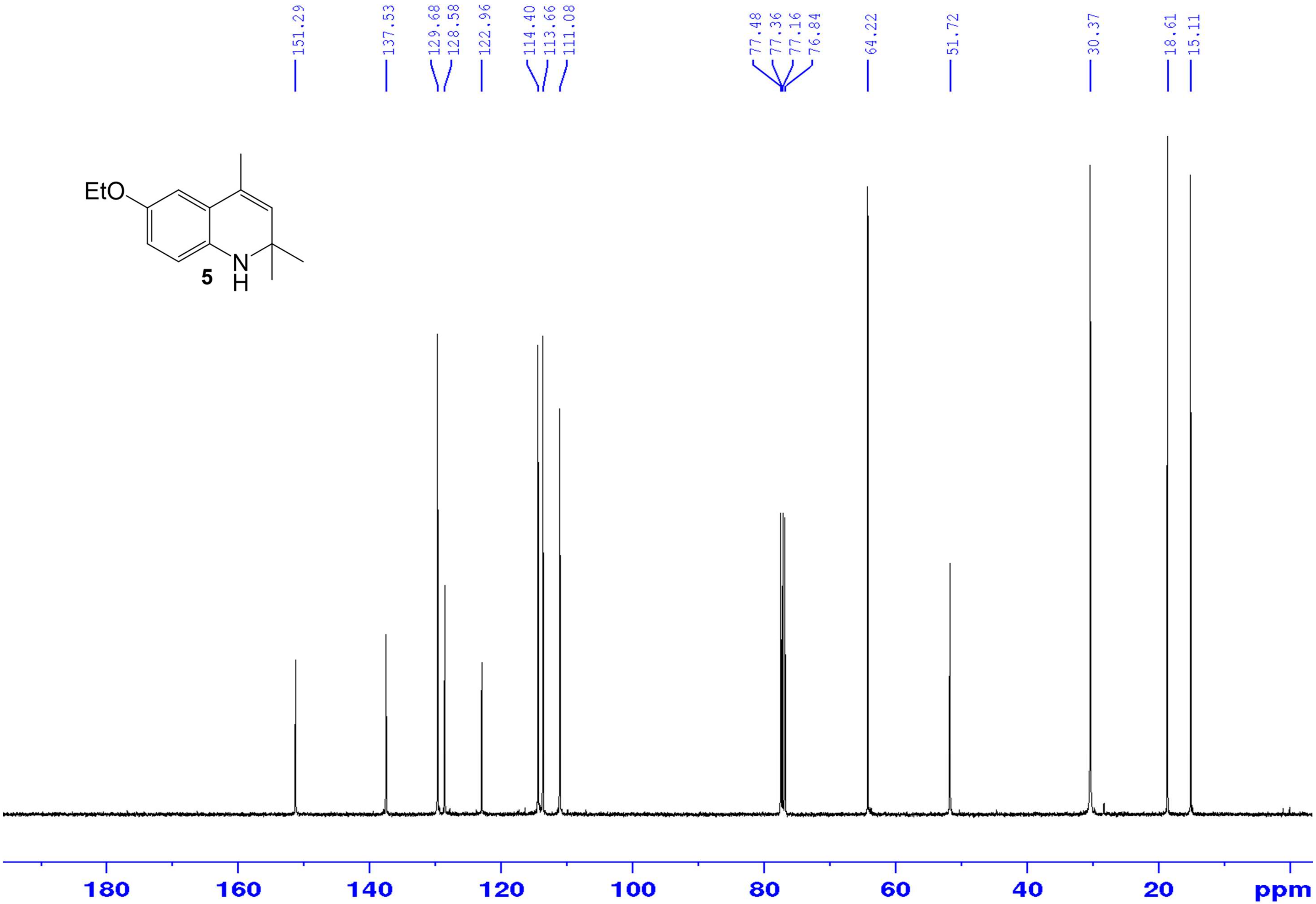

*8-Ethoxy-2,2,4-trimethyl-1,2-dihydroquinoline (**6**)* – $^{1}$H NMR ($CDCl_3$, 400 MHZ)

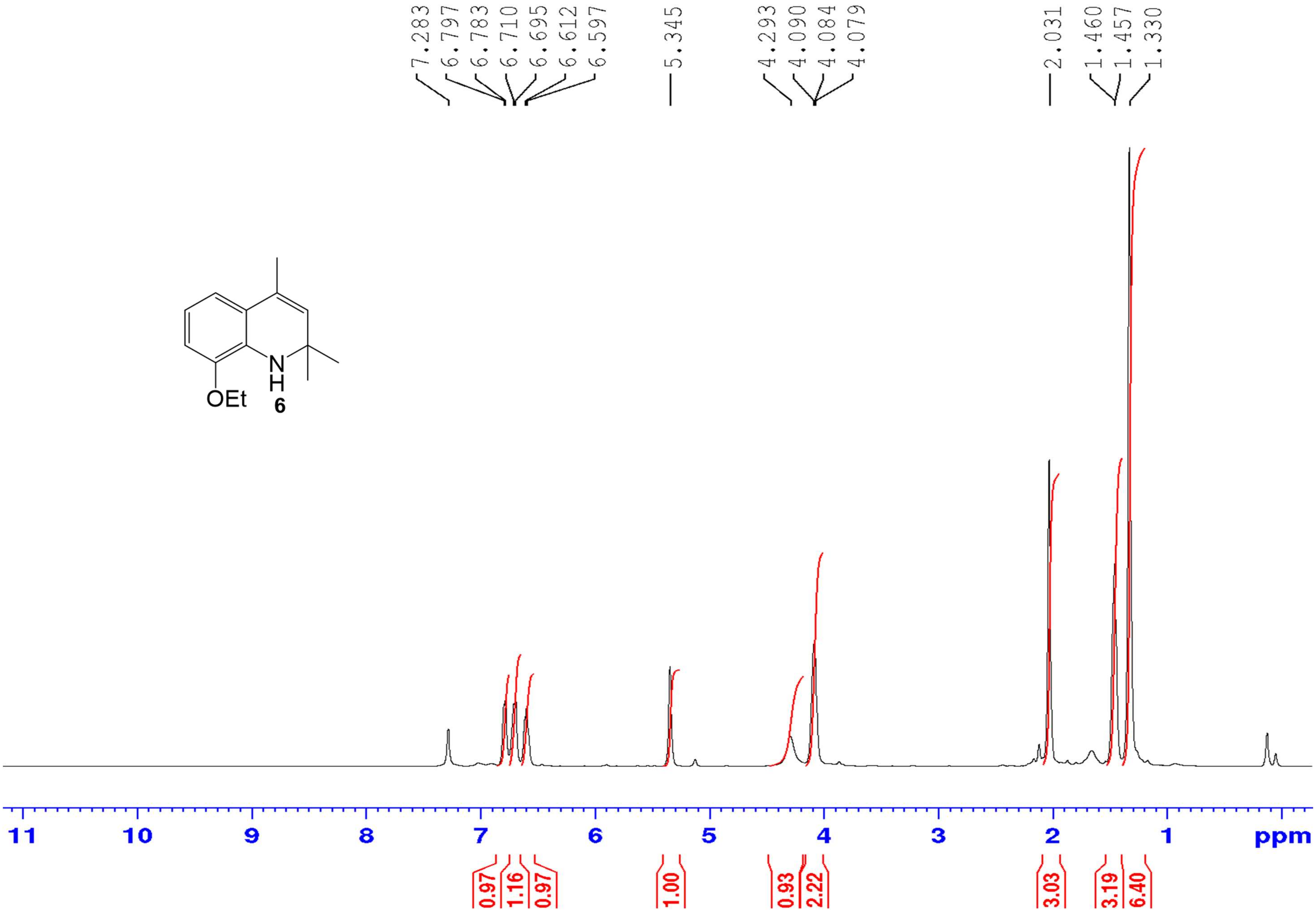

*8-Ethoxy-2,2,4-trimethyl-1,2-dihydroquinoline (**6**)* – $^{13}C$ NMR ($CDCl_3$, 100 MHZ)

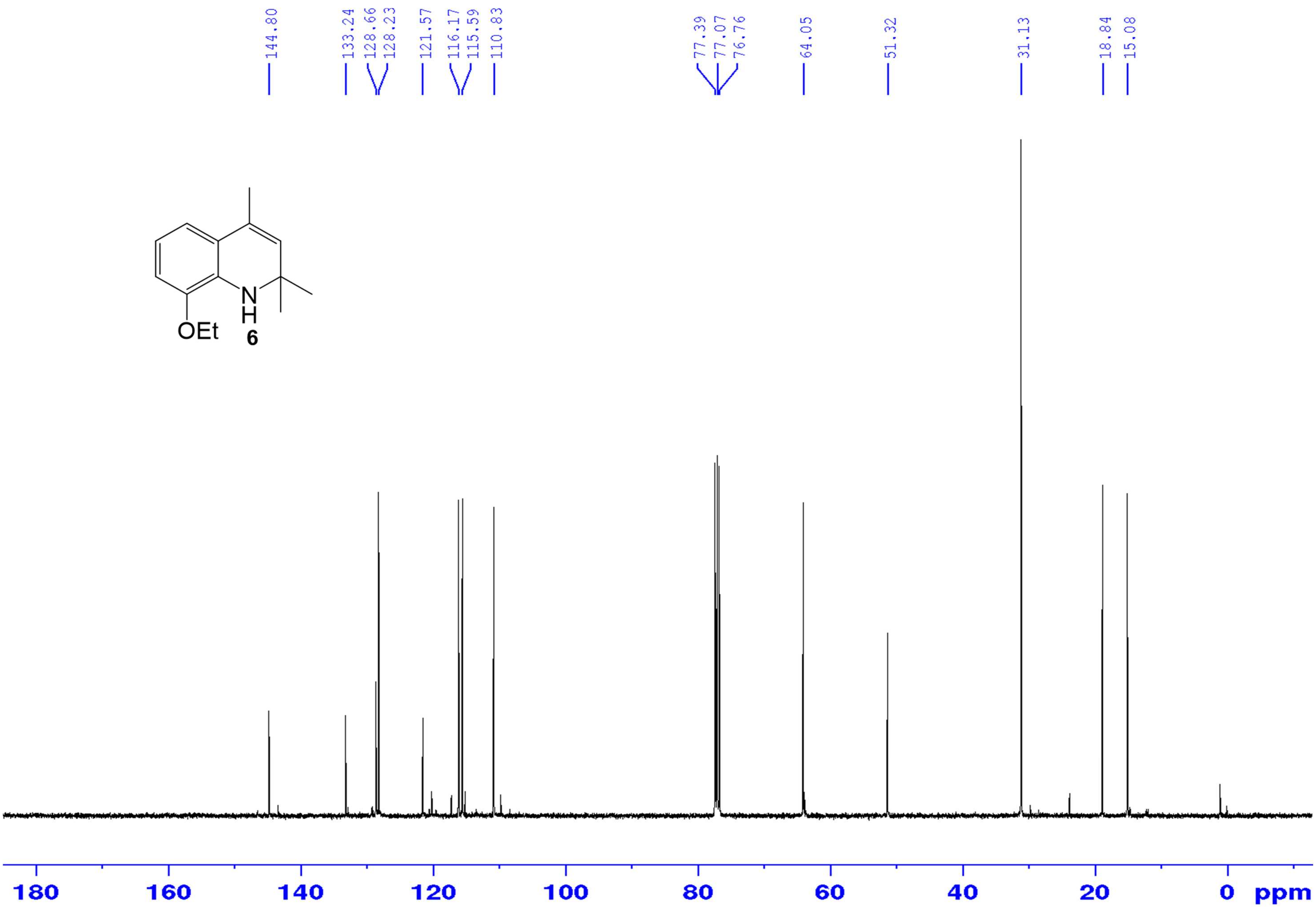

*8-Chloro-6-cyclopentenyl-2,3,4,5-tetrahydro-4,4-tetramethylene-1H-cyclopenta[c]quinoline (**7**)* – [1]H NMR ($CDCl_3$, 400 MHz)

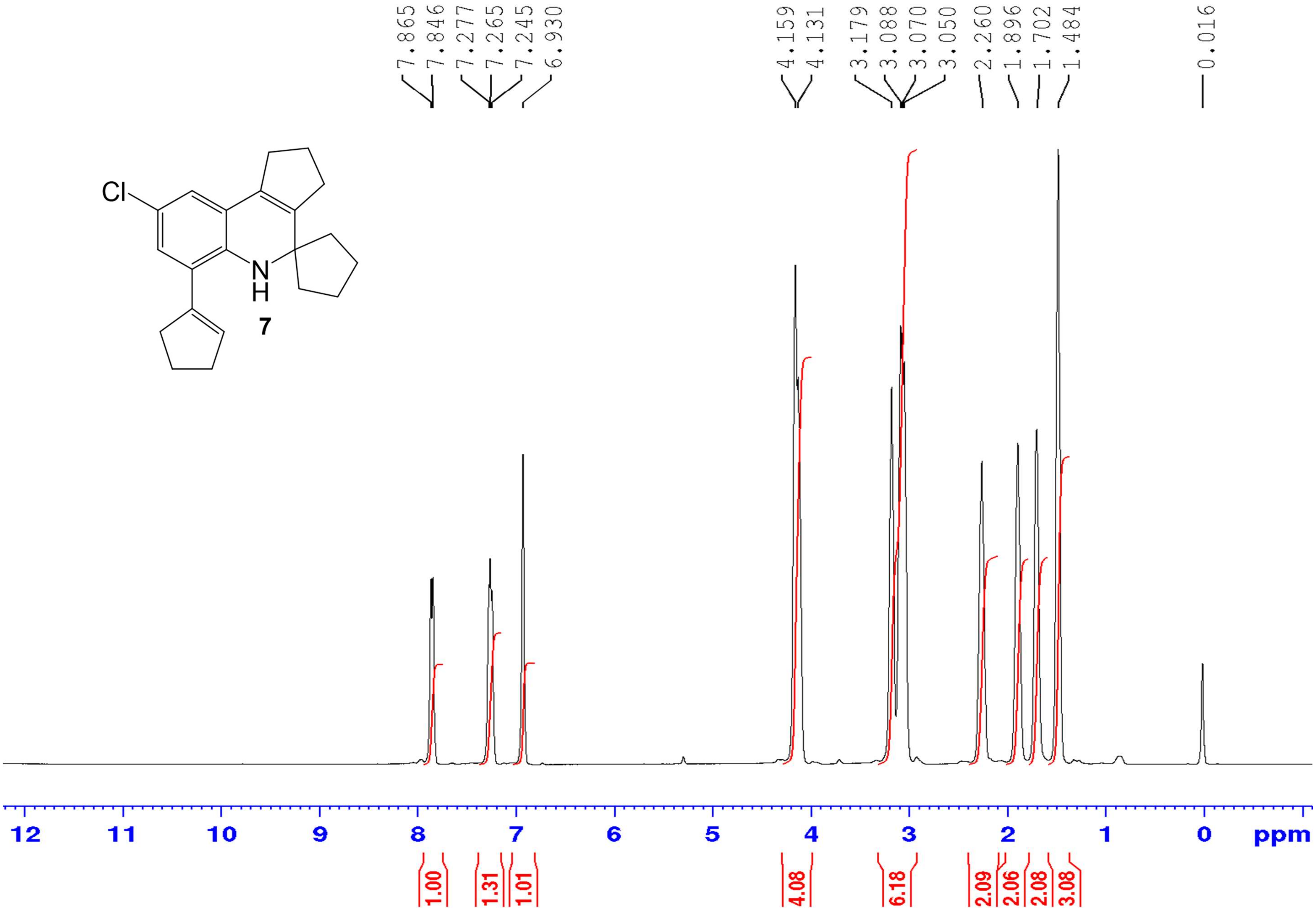

*8-Chloro-6-cyclopentenyl-2,3,4,5-tetrahydro-4,4-tetramethylene-1H-cyclopenta[c]quinoline (**7**)* – $^{13}$C NMR ($CDCl_3$, 100 MHz)

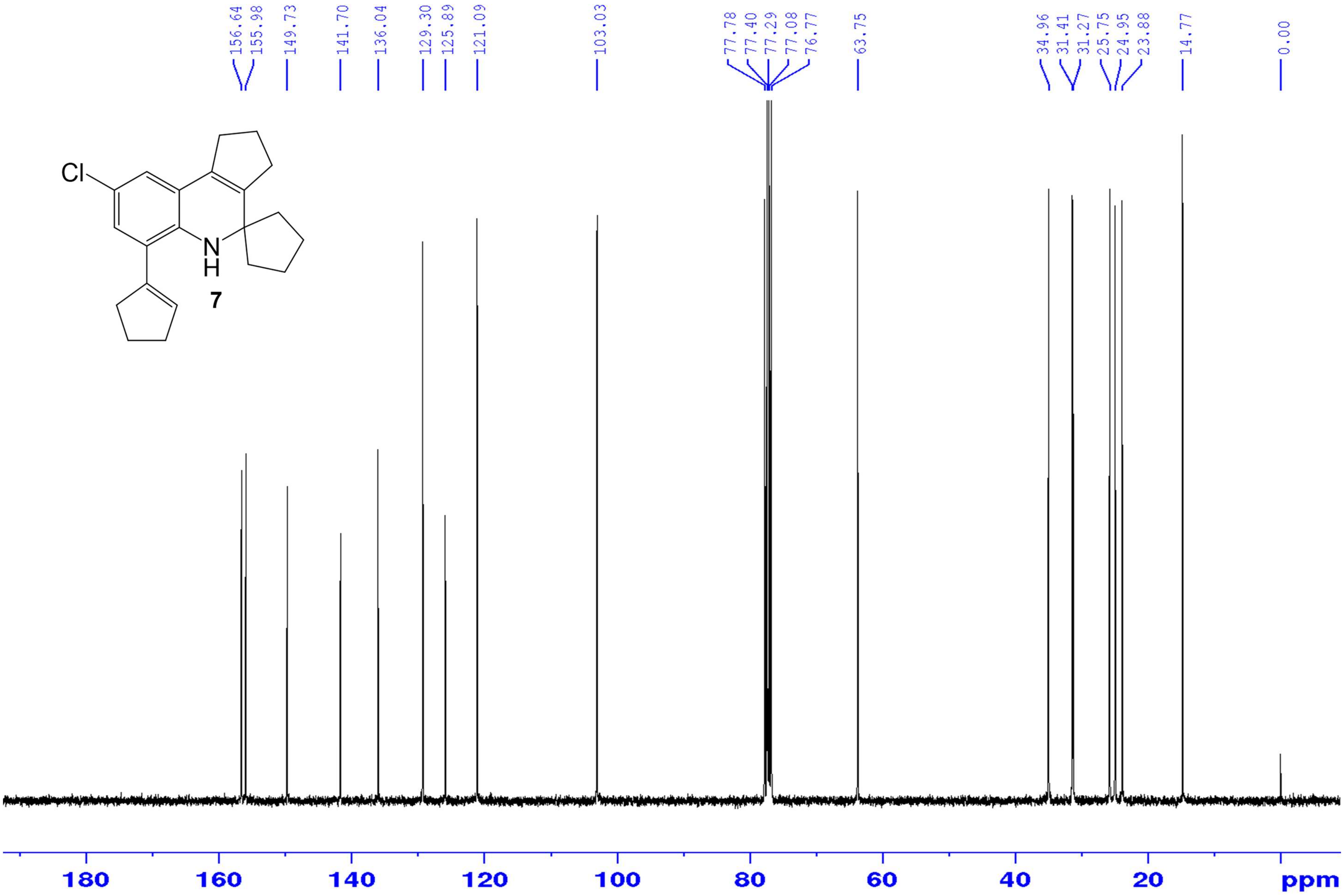

*8-Chloro-6-cyclopentenyl-2,3,4,5-tetrahydro-4,4-tetramethylene-1H-cyclopenta[c]quinoline* (**7**) – HR-ESIMS ($[M+H]^+$ Calculated: 326.167004, Observed: 326.1669)

A) Mass spectrum (m/z range 150 to 1100)

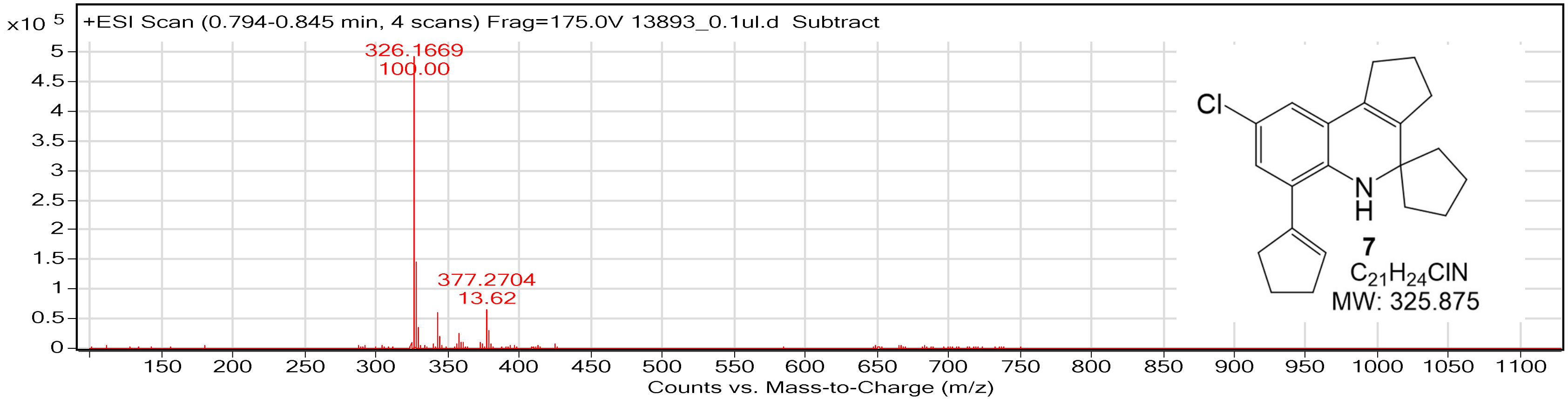


B) Mass spectrum (m/z range 325 to 332)

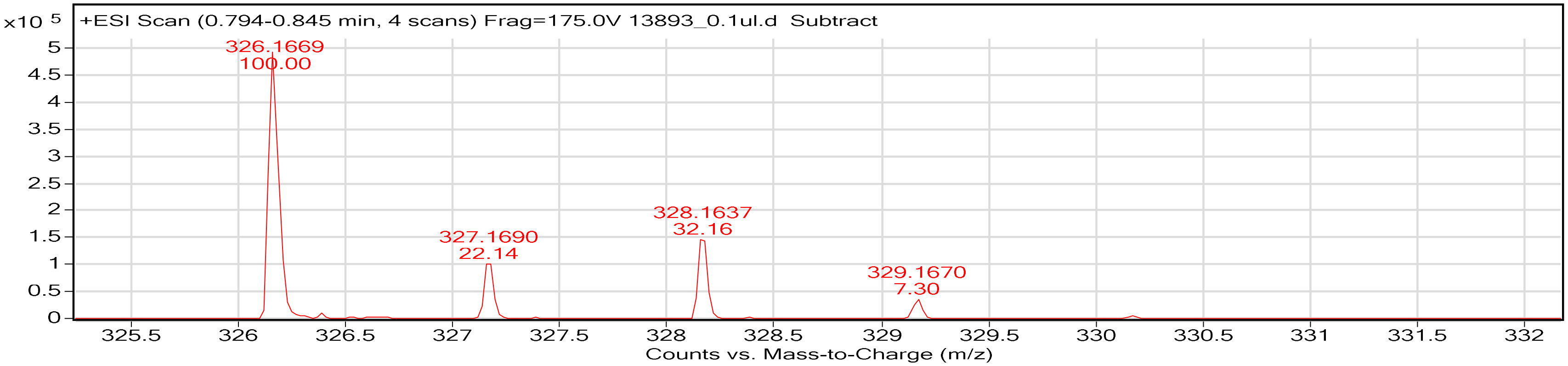


Empirical Formula Confirmation Report

| Formula | Calculated Mass | mDa Error | ppm Error | RDB |
|---|---|---|---|---|
| C21 H25 N Cl | 326.167004 | -0.104157 | -0.31934 | 9.5 |

*8-Cyclohexenyl-6-ethoxy-1,2-dihydro-2,2,4-trimethylquinoline* (***8***) – [1]H NMR ($CDCl_3$, MHz)

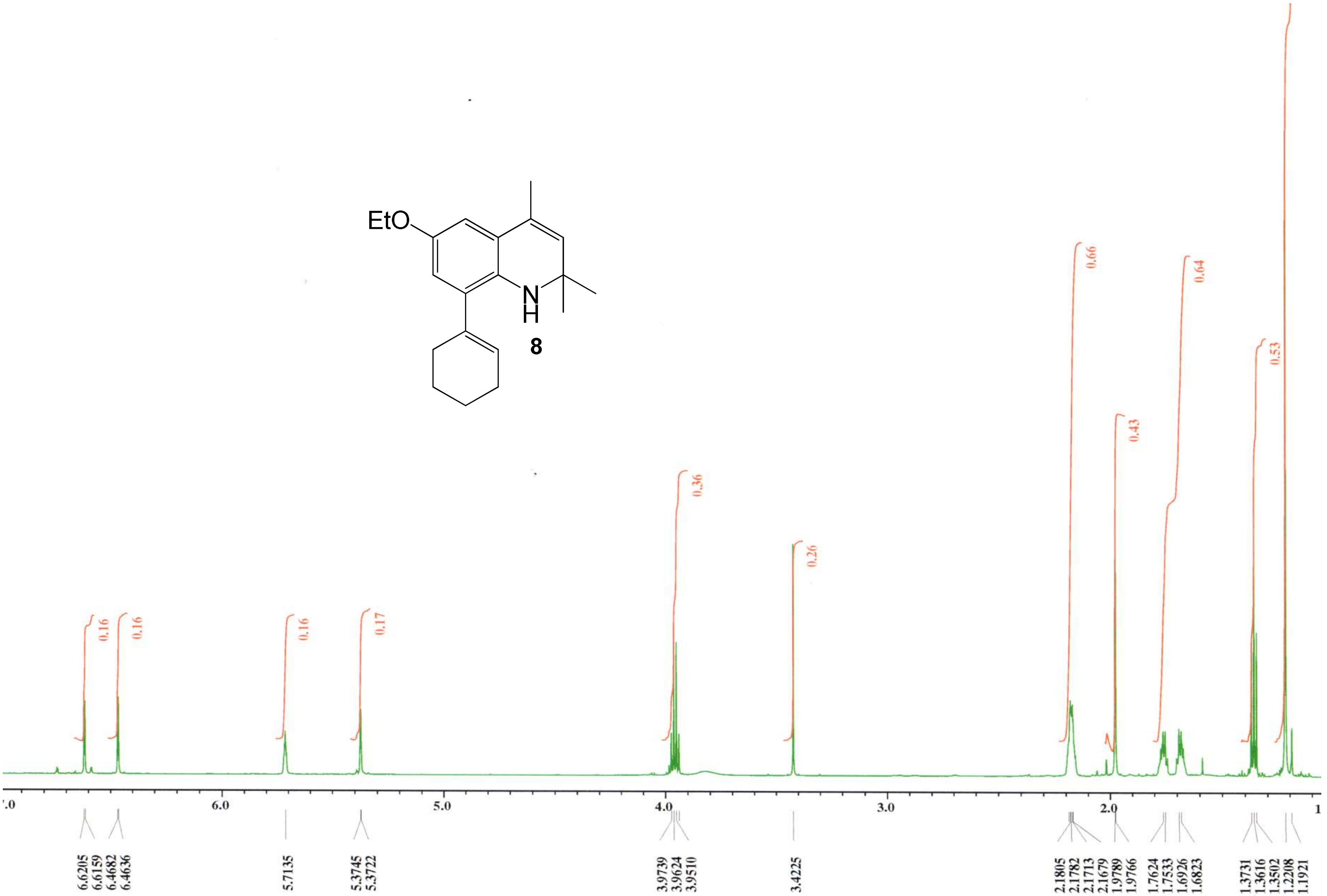

*8-Cyclohexenyl-6-ethoxy-1,2-dihydro-2,2,4-trimethylquinoline (**8**)* – $^{13}$C NMR ($CDCl_3$, 100 MHz)

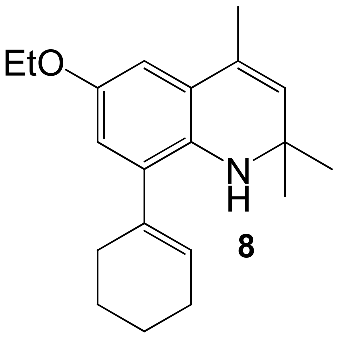


**8**

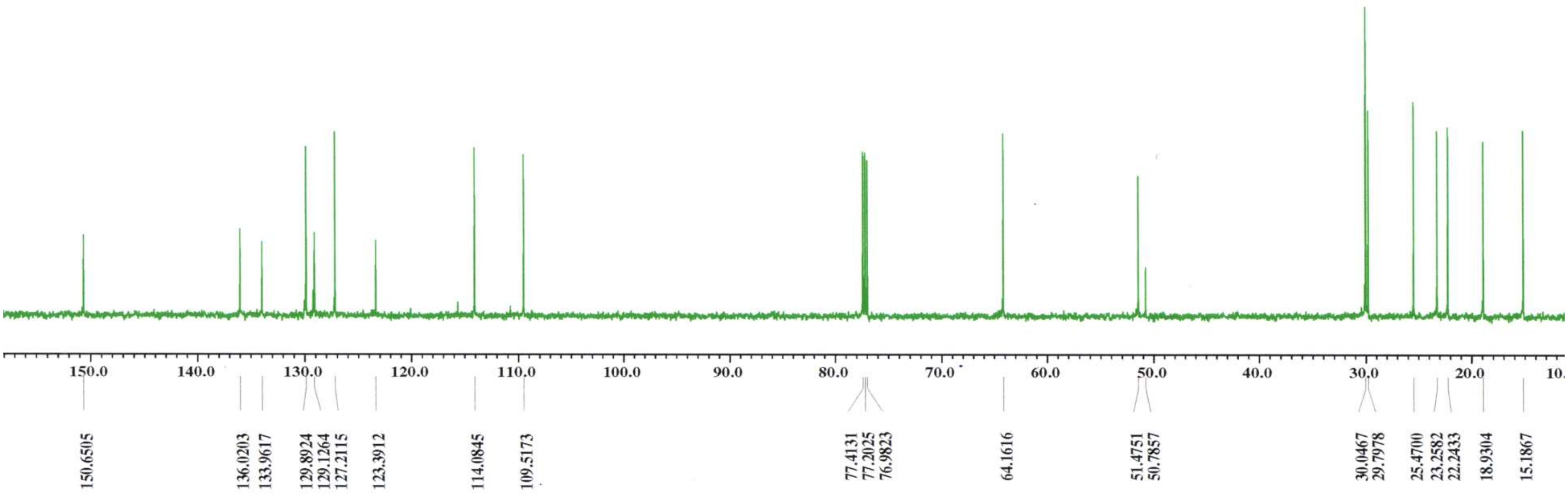

*8-Cyclohexenyl-6-ethoxy-1,2-dihydro-2,2,4-trimethylquinoline (**8**)* – HR-ESIMS ([M+H]$^+$ Calculated: 298.216541, Observed: 298.2158)

#### A) Mass spectrum (m/z range 50 to 1600)

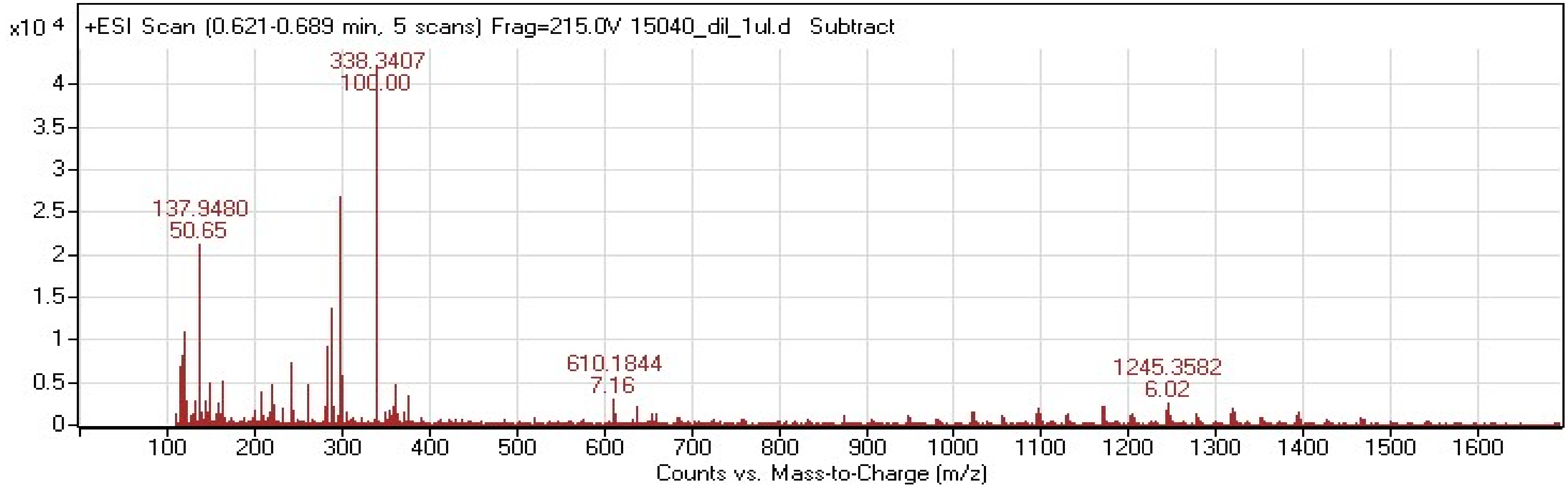


#### B) Mass spectrum (m/z range 297 to 302)

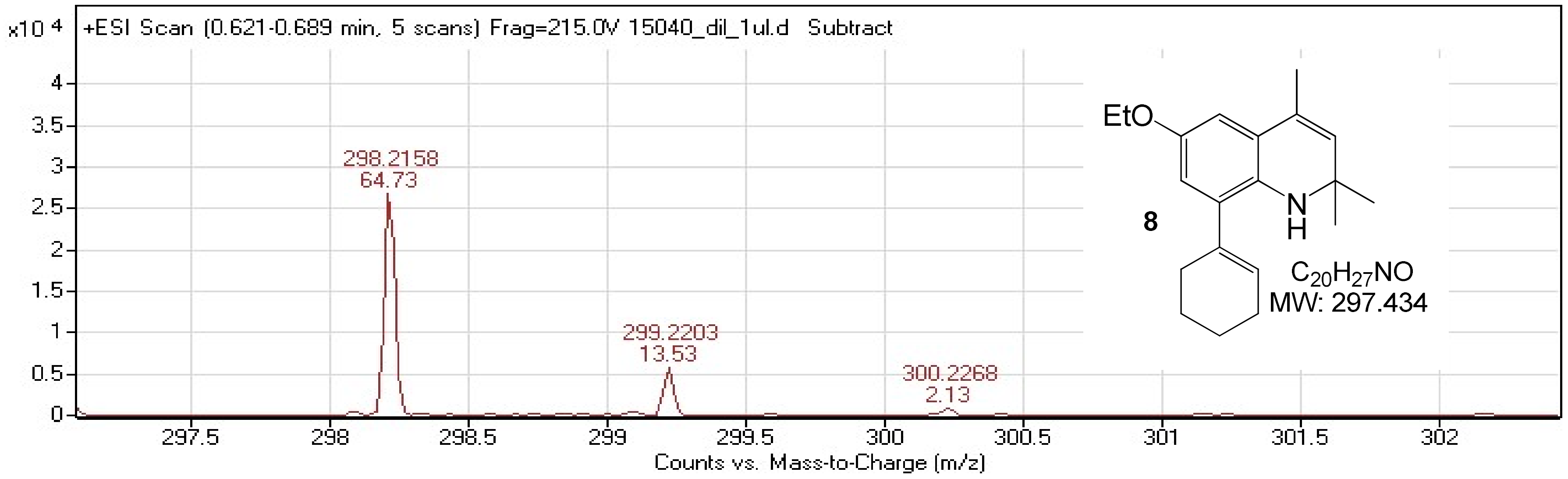

*6,8-Dicyclopentenyl-1,2-dihydro-2,2,4-trimethylquinoline (**9**)* – [1]H NMR ($CDCl_3$, 400 MHz)

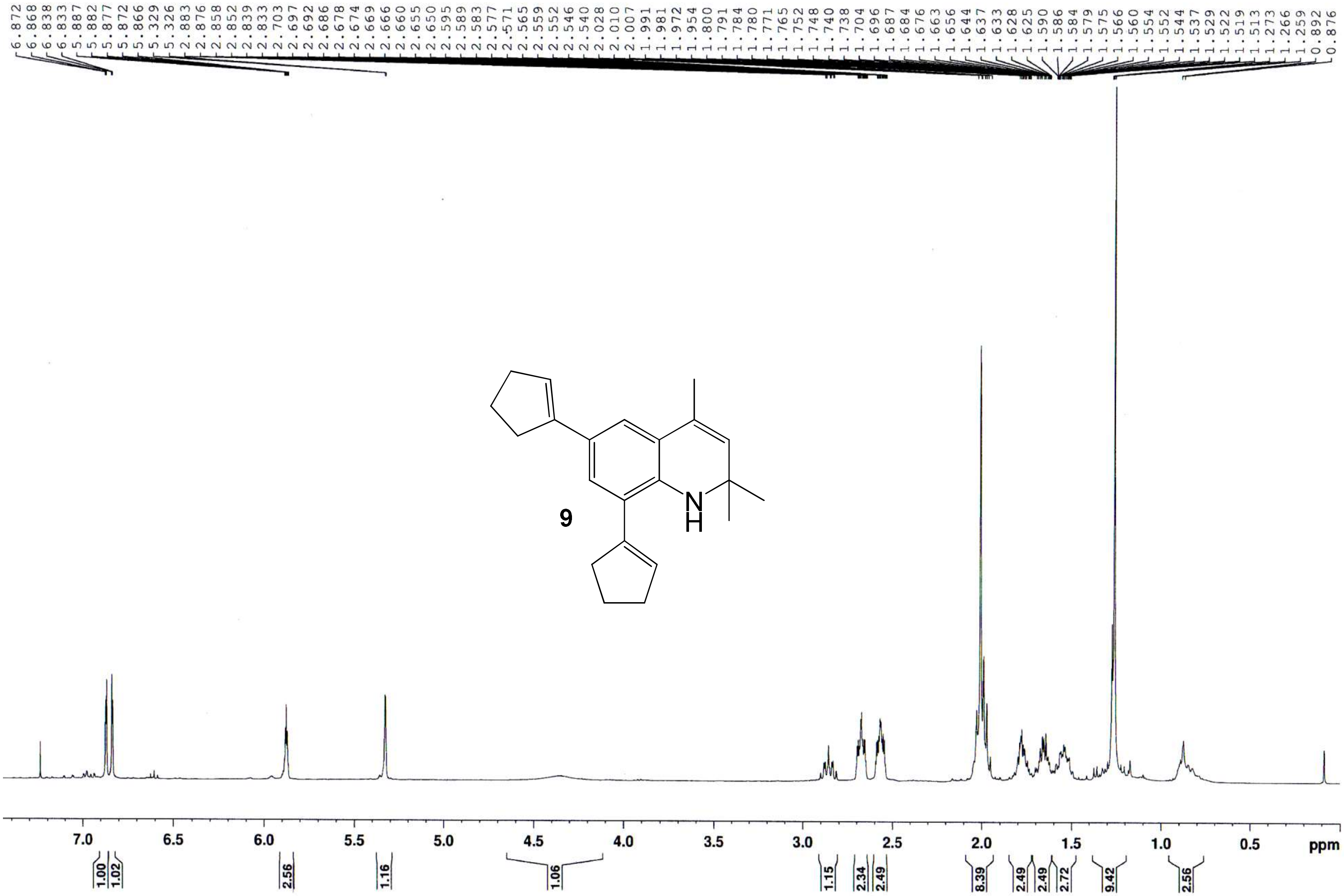

*6,8-Dicyclopentenyl-1,2-dihydro-2,2,4-trimethylquinoline (**9**)* – $^{13}$C NMR ($CDCl_3$, 100 MHz)

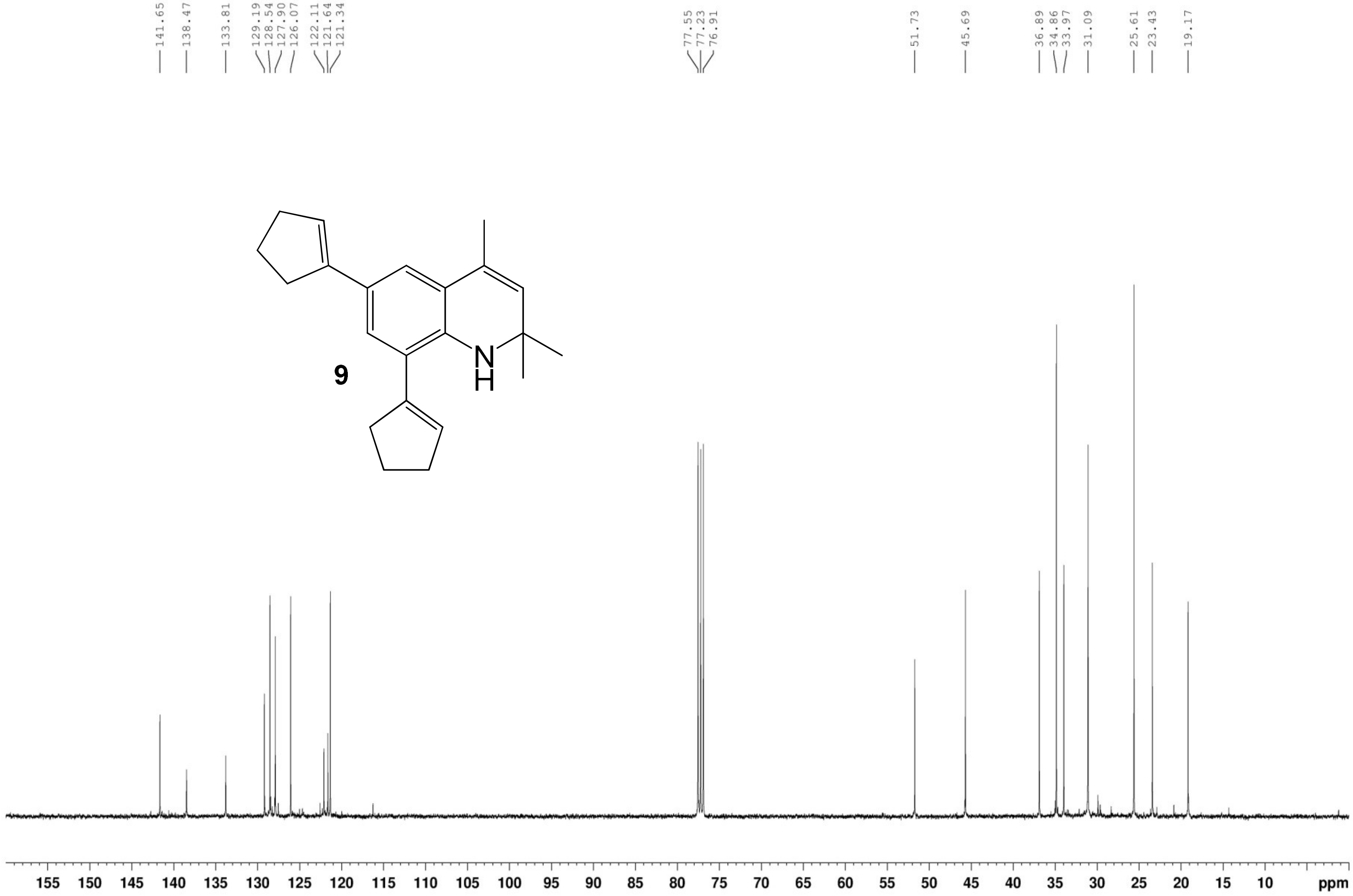


**9**

*6,8-Dicyclopentenyl-1,2-dihydro-2,2,4-trimethylquinoline (**9**)* – HR-ESIMS ($[M]^+$ Calculated: 305.2144, Observed: 305.2152)

### Elemental Composition Report

**Page 1**

#### Single Mass Analysis

Tolerance = 5.0 PPM / DBE: min = -1.5, max = 50.0
Isotope cluster parameters: Separation = 1.0 Abundance = 1.0%

Monoisotopic Mass, Odd and Even Electron Ions
99 formula(e) evaluated with 1 results within limits (up to 50 closest results for each mass)

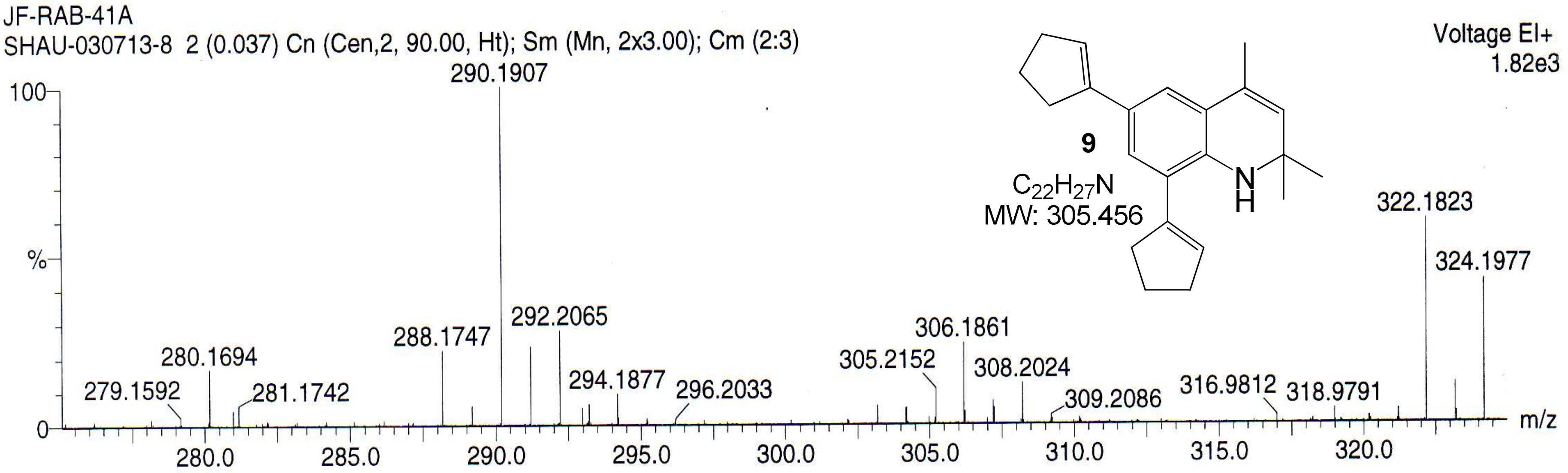


| | | | | | | |
|---|---|---|---|---|---|---|
| Minimum: | | | | -1.5 | | |
| Maximum: | | 2.0 | 5.0 | 50.0 | | |
| Mass | Calc. Mass | mDa | PPM | DBE | Score | Formula |
| 305.2152 | 305.2144 | 0.8 | 2.8 | 10.0 | 1 | C22 H27 N |

*N1,N2-diethyl-4-methoxy-N2-(4-methoxyphenyl)benzene-1,2-diamine (**11**)* – $^{1}$H NMR ($CDCl_3$, 400 MHZ)

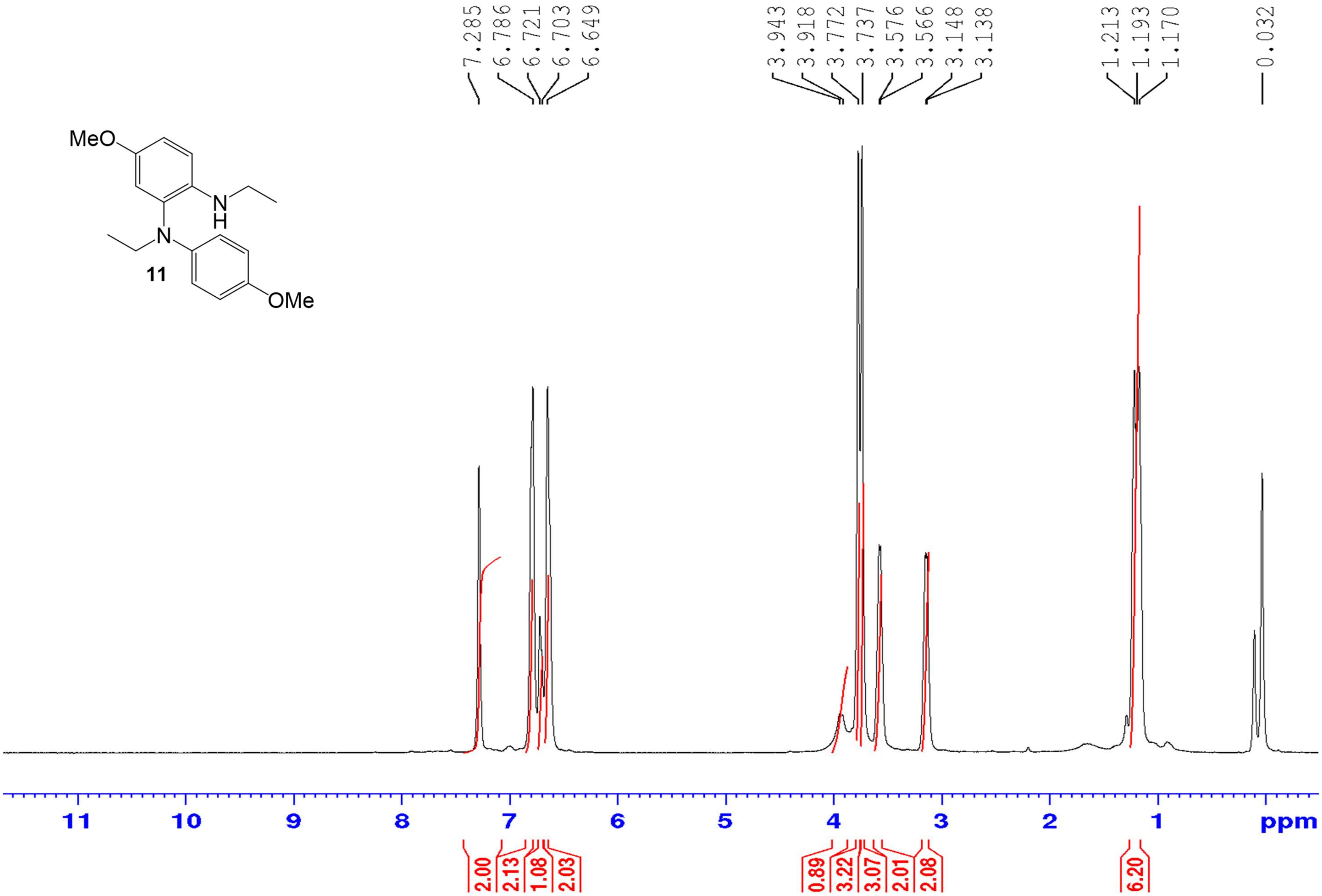

*N1,N2-diethyl-4-methoxy-N2-(4-methoxyphenyl)benzene-1,2-diamine (**11**)* – $^{13}$C NMR ($CDCl_3$, 100 MHZ)

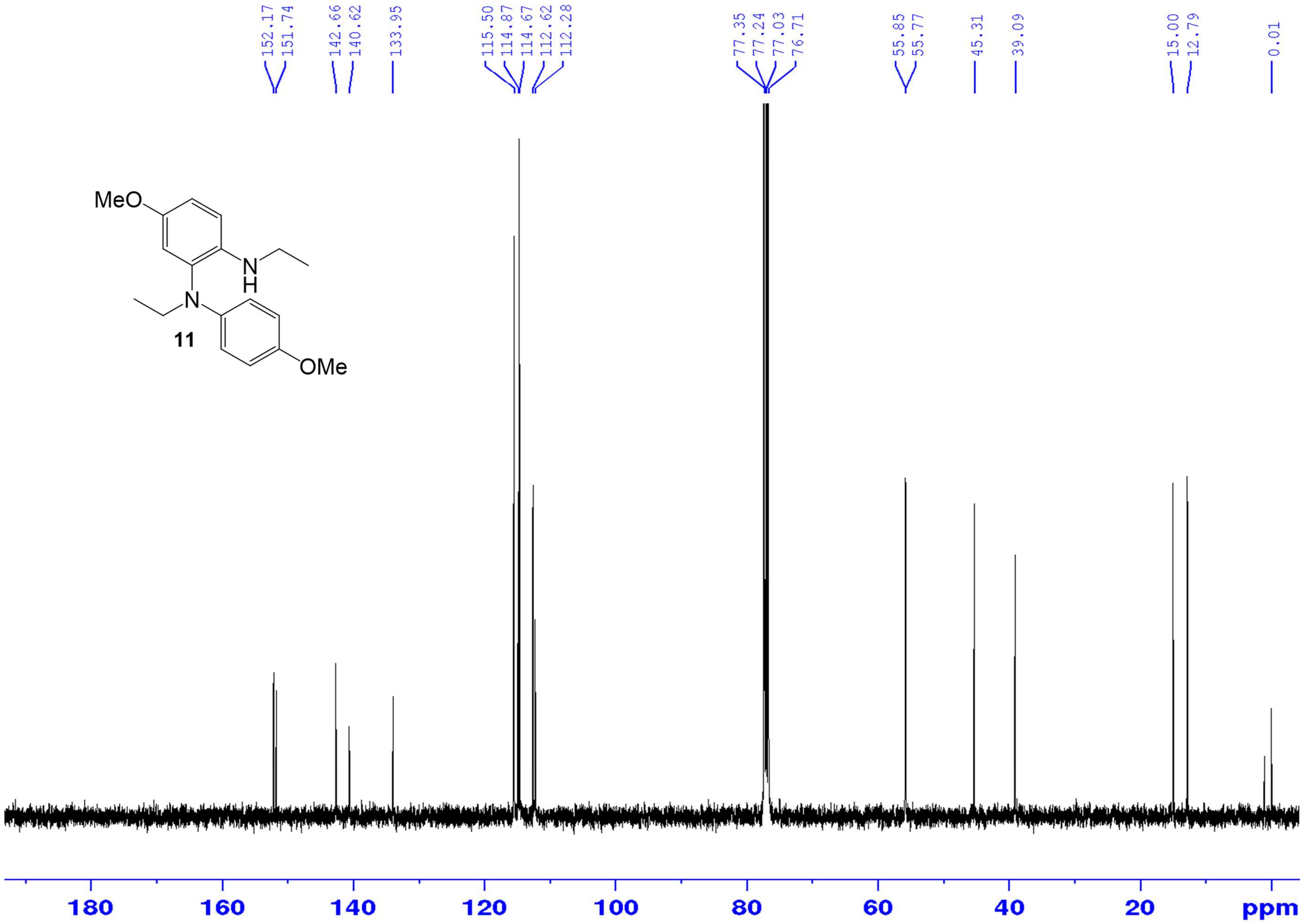

*N1,N2-diethyl-4-methoxy-N2-(4-methoxyphenyl)benzene-1,2-diamine (**11**)* – HR-ESIMS ($[M+H]^+$ Calculated: 301.19105, Observed: 301.19224)

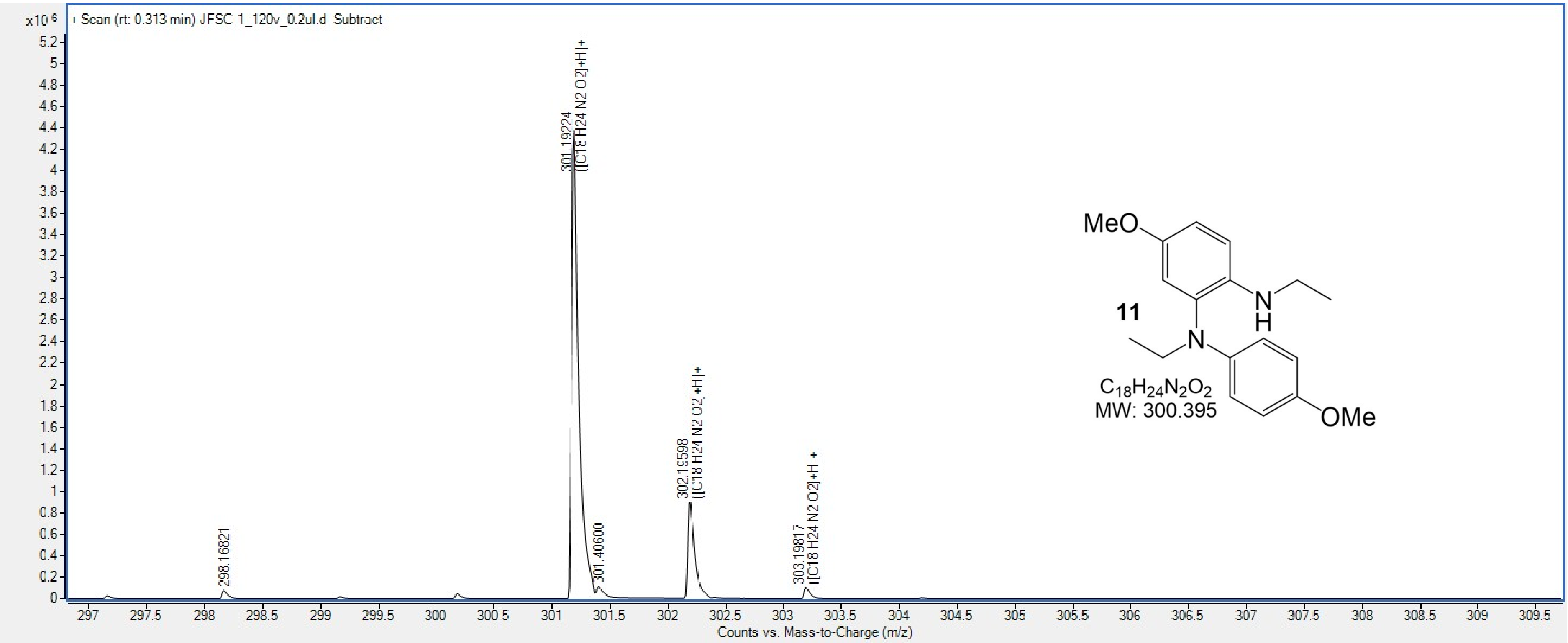


| Formula | Species | m/z, | Score | Diff (abs. ppm), | Mass |
|---|---|---|---|---|---|
| $C_{18}H_{24}N_2O_2$ | $[M+H]^+$ | 301.19224 | 94.1 | 4.27 | 300.18378 |

*1-(2-Amino-N,N,5-trimethylphenyl)naphthalen-2-ol* ***(12)*** – $^{1}$H NMR ($CDCl_3$, 400 MHz)

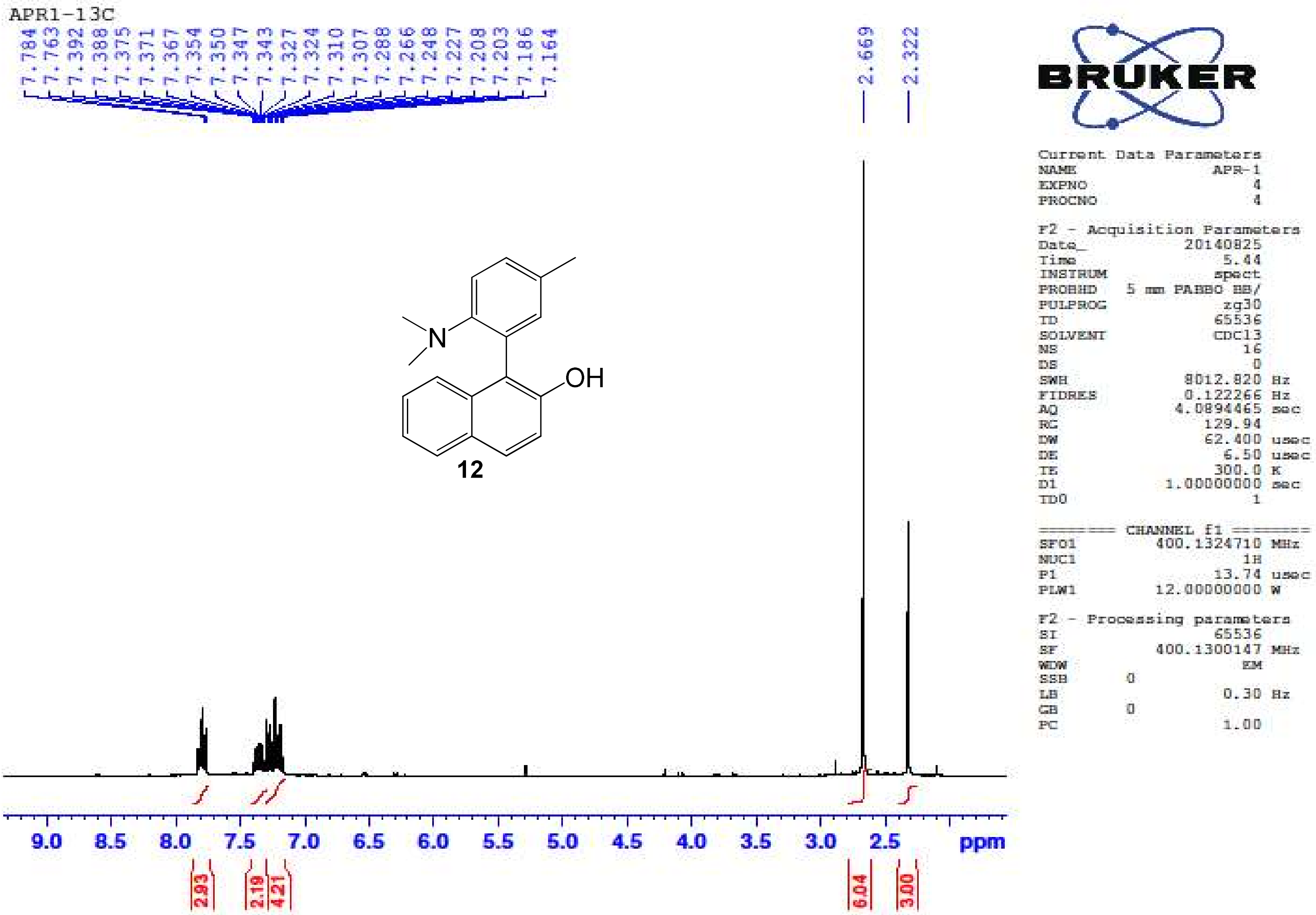

*1-(2-Amino-N,N,5-trimethylphenyl)naphthalen-2-ol* ***(12)*** – $^{13}$C NMR ($CDCl_3$, 100 MHz)

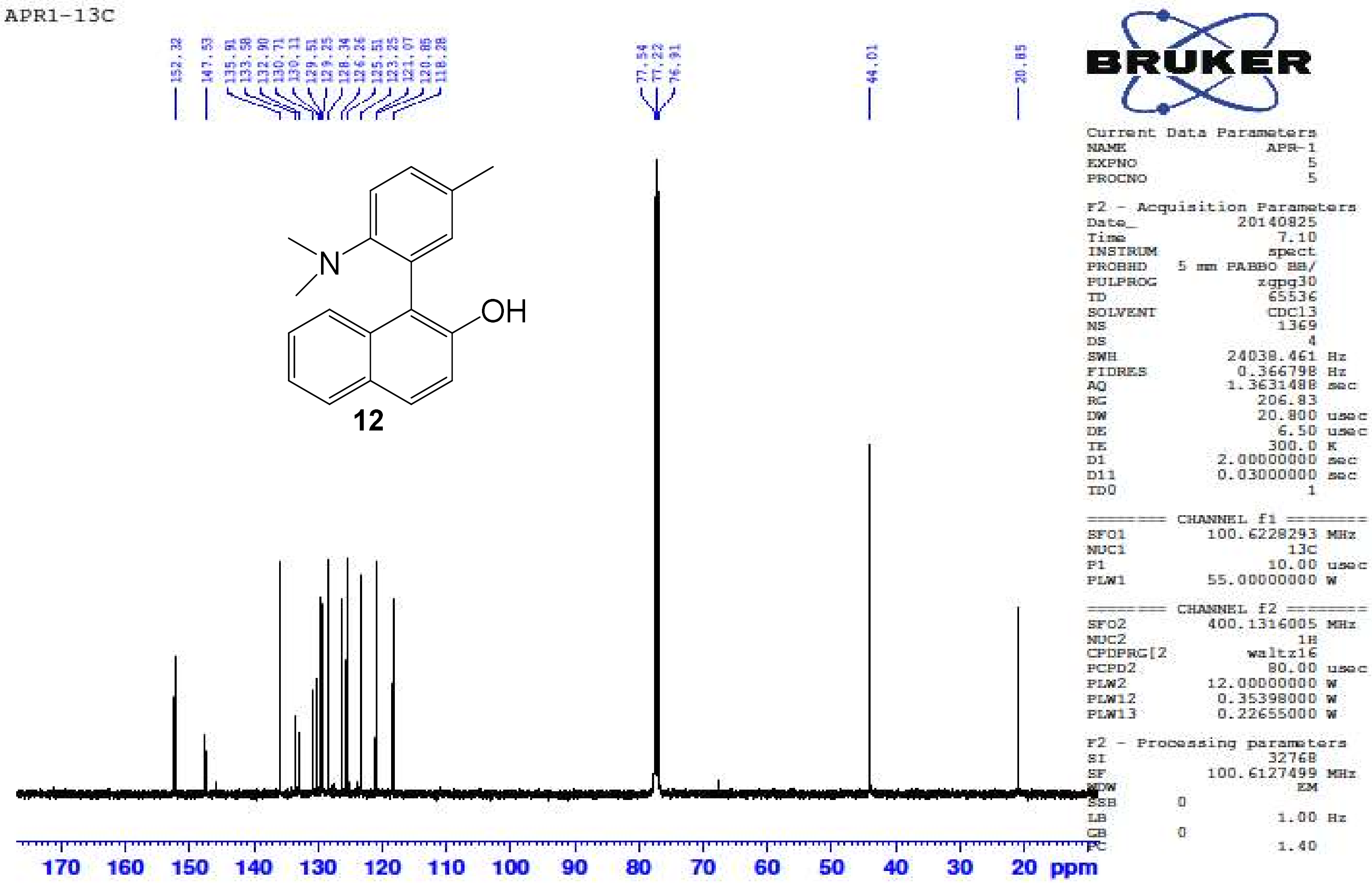

*1-(2-Amino-N,N,5-trimethylphenyl)naphthalen-2-ol* ***(12)*** – HR-ESIMS ($[M+H]^+$ Calculated: 278.1539, Observed 278.1538)

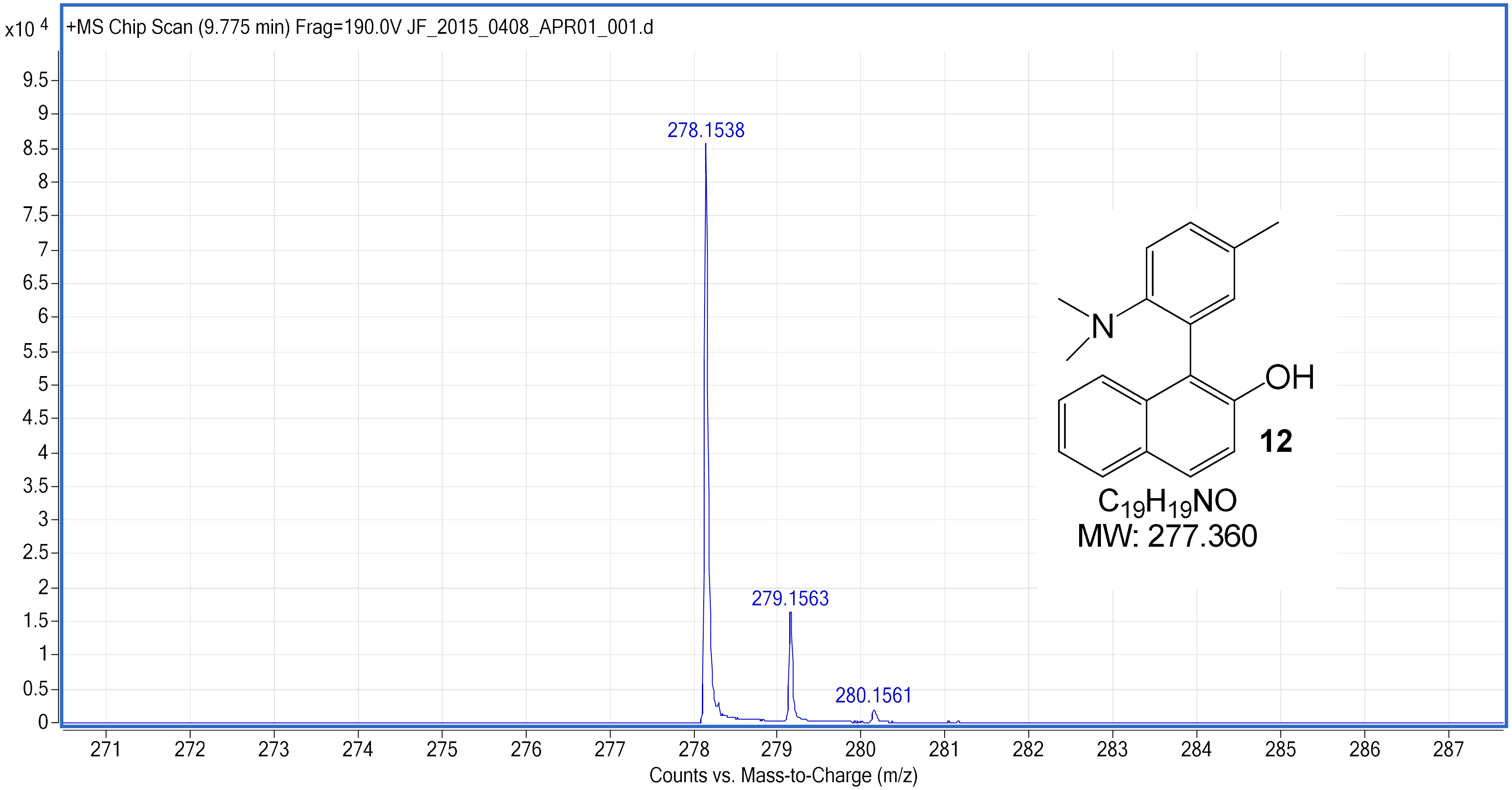

*1-(2-Amino-5-ethoxy-N,N-dimethyl-phenyl)naphthalen-2-ol* ***(13)*** – [1]H NMR (CDCl$_3$, 400 MHz)

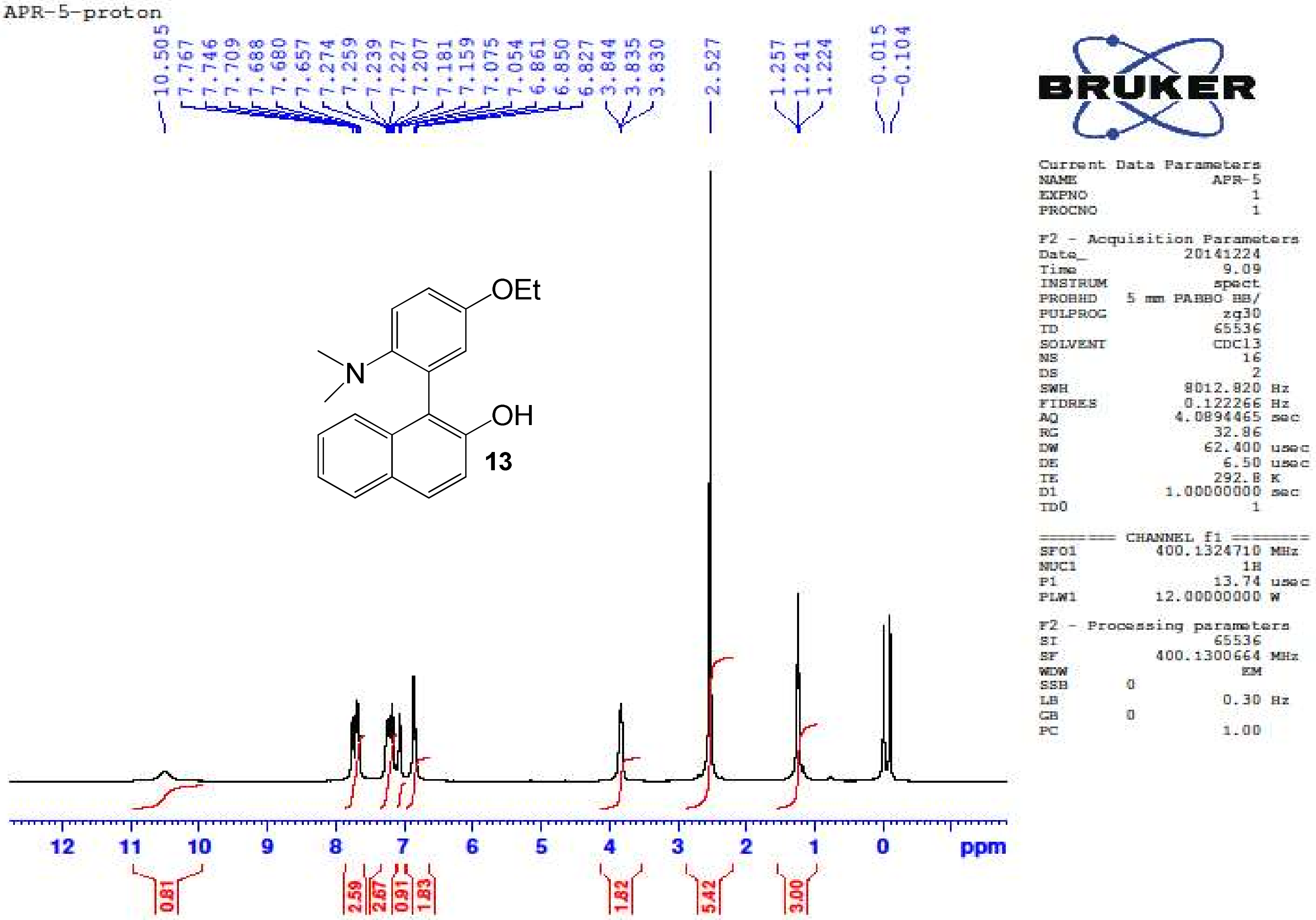

*1-(2-Amino-5-ethoxy-N,N-dimethyl-phenyl)naphthalen-2-ol* ***(13)*** – $^{13}$C NMR ($CDCl_3$, 100 MHz)

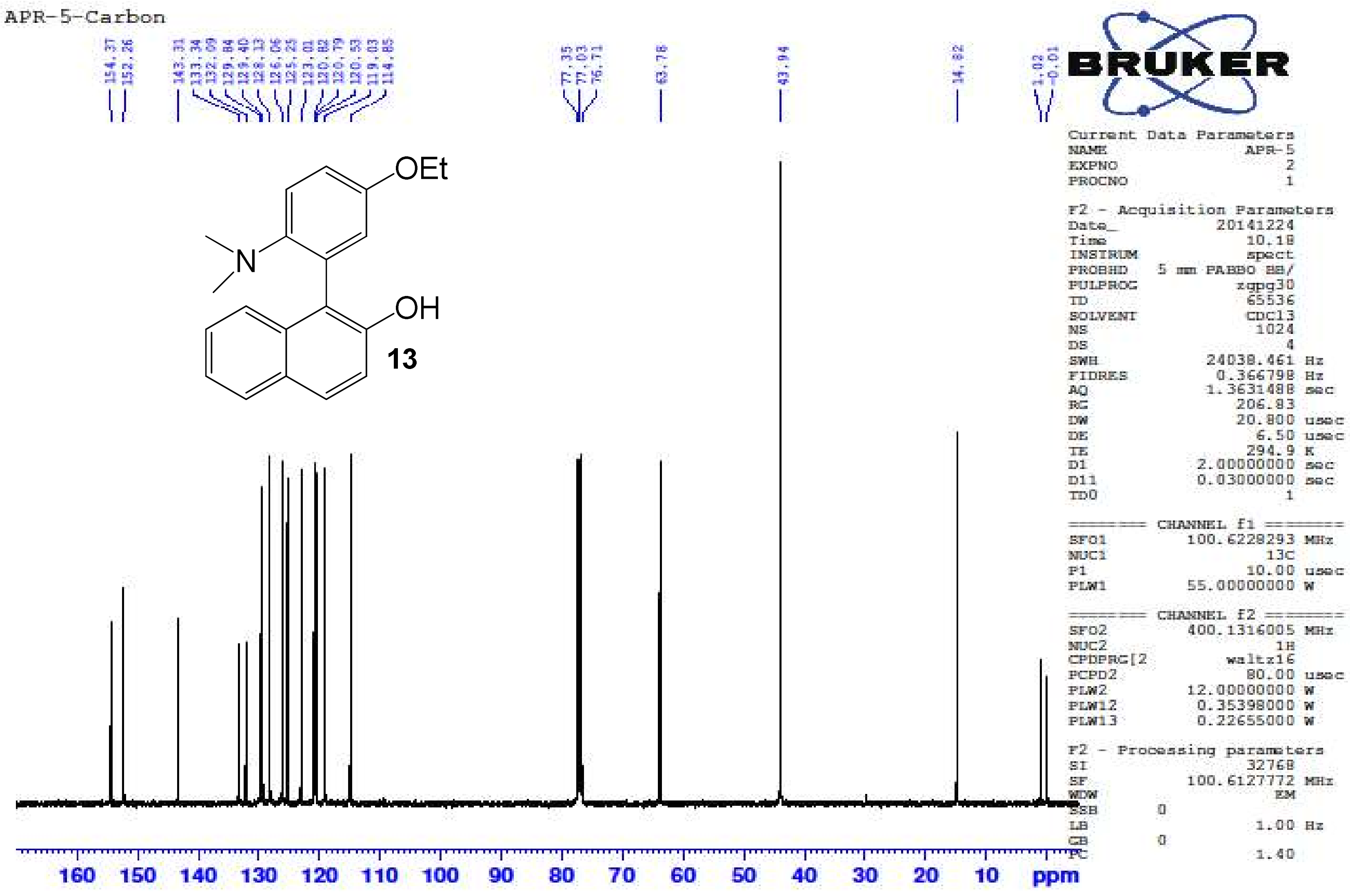

*1-(2-Amino-5-ethoxy-N,N-dimethyl-phenyl)naphthalen-2-ol **(13)*** – HR-ESIMS ($[M+H]^+$ Calculated: 308.1645, Observed 308.1655)

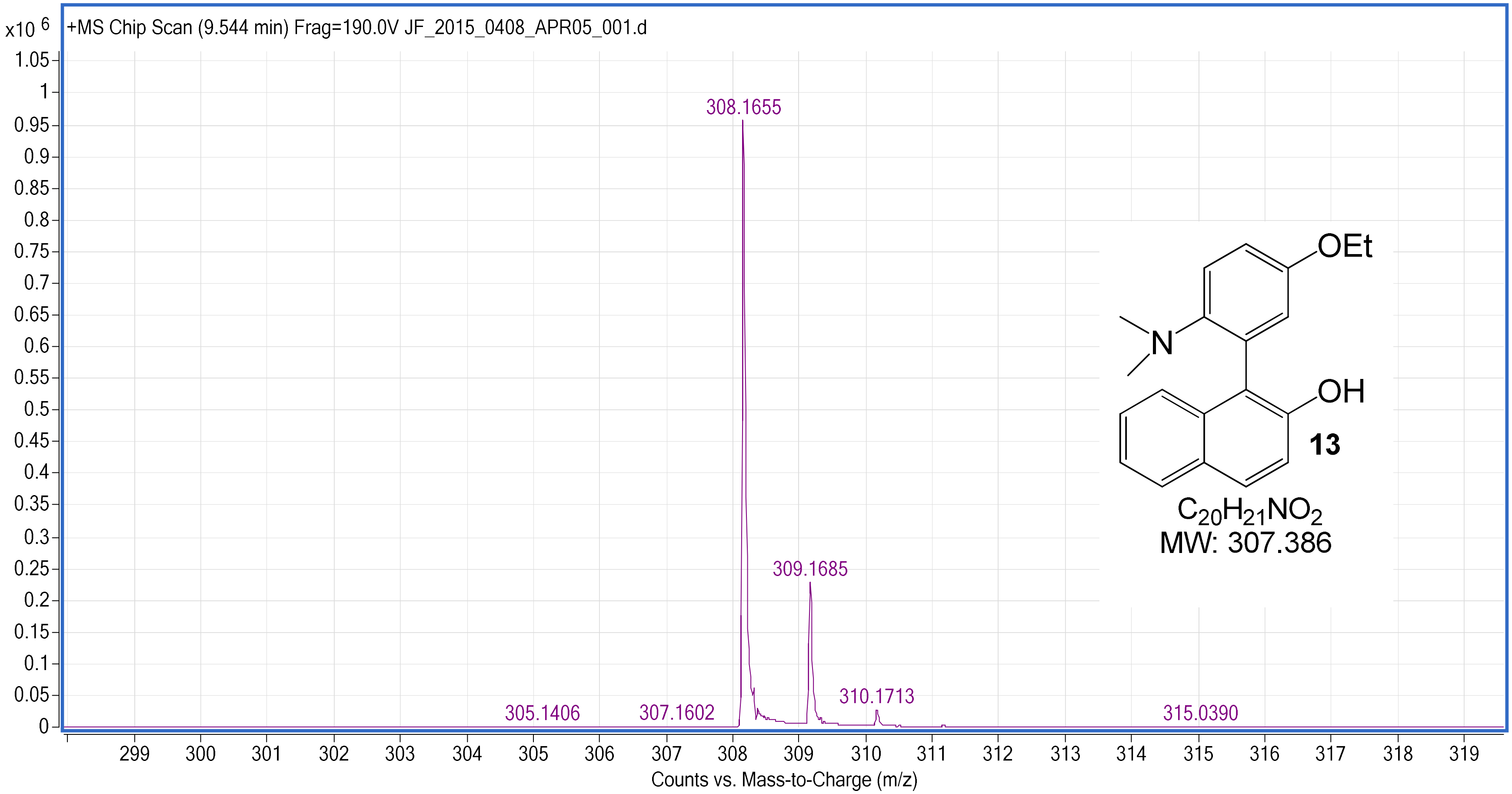

*1-(2-Amino-N-benzyl-5-ethoxy-N-methylphenyl)naphthalen-2-ol* ***(14)*** – [1]H NMR ($CDCl_3$, 400 MHz)

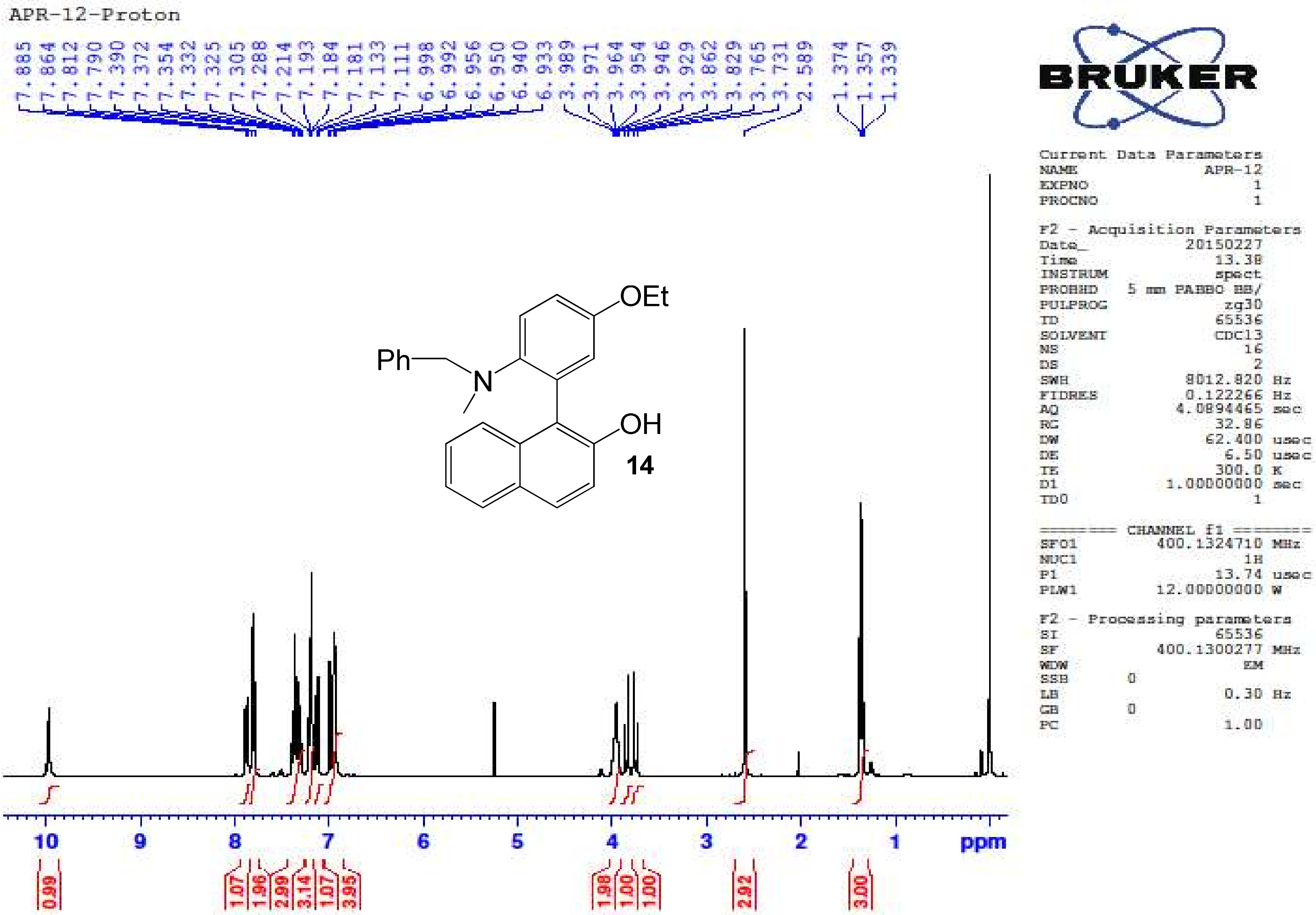

*1-(2-Amino-N-benzyl-5-ethoxy-N-methylphenyl)naphthalen-2-ol* ***(14)*** – [13]C NMR ($CDCl_3$, 100 MHz)

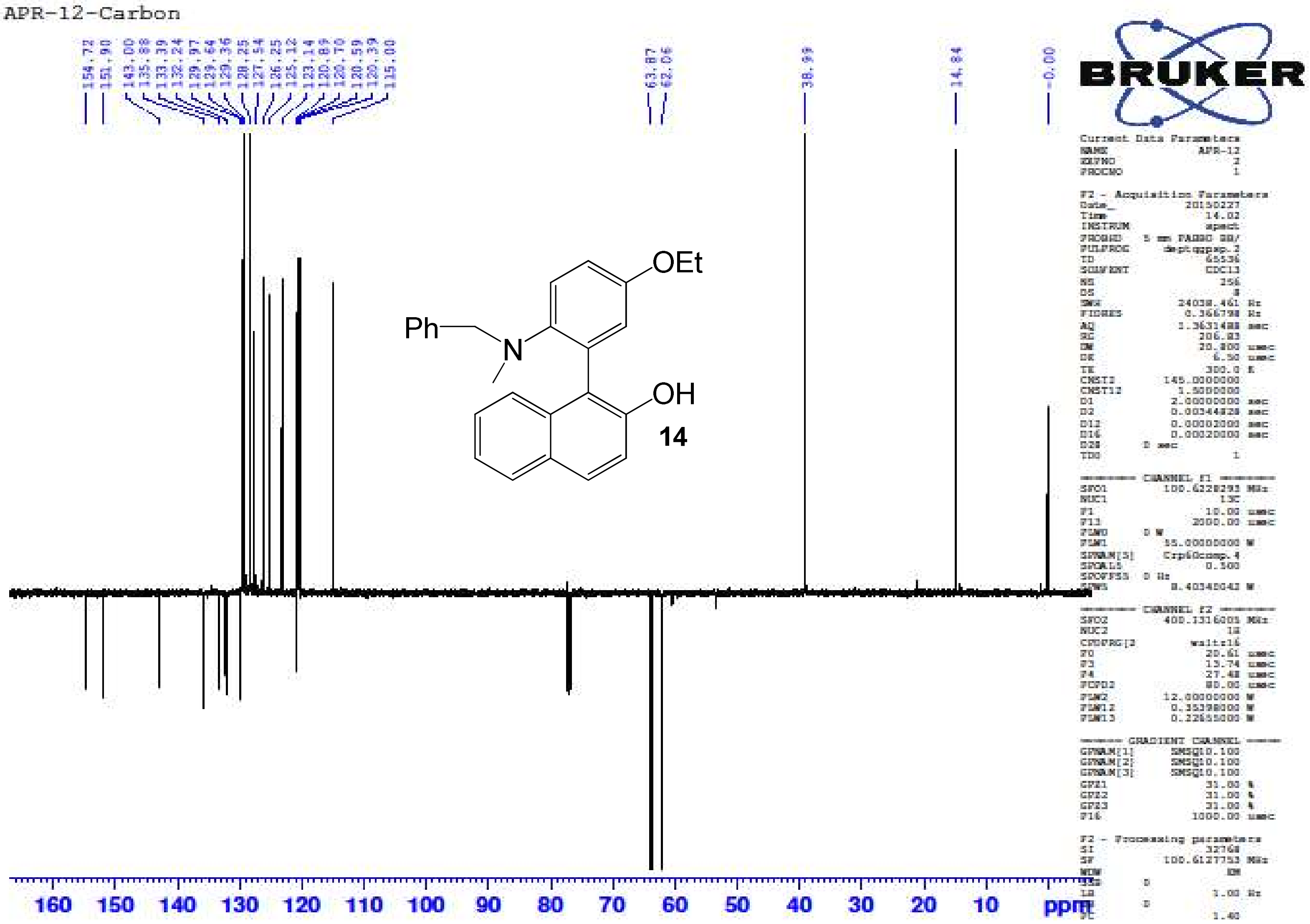

*1-(2-Amino-N-benzyl-5-ethoxy-N-methylphenyl)naphthalen-2-ol* ***(14)*** – HR-ESIMS ($[M+H]^+$ Calculated: 384.1958, Observed 384.1973)

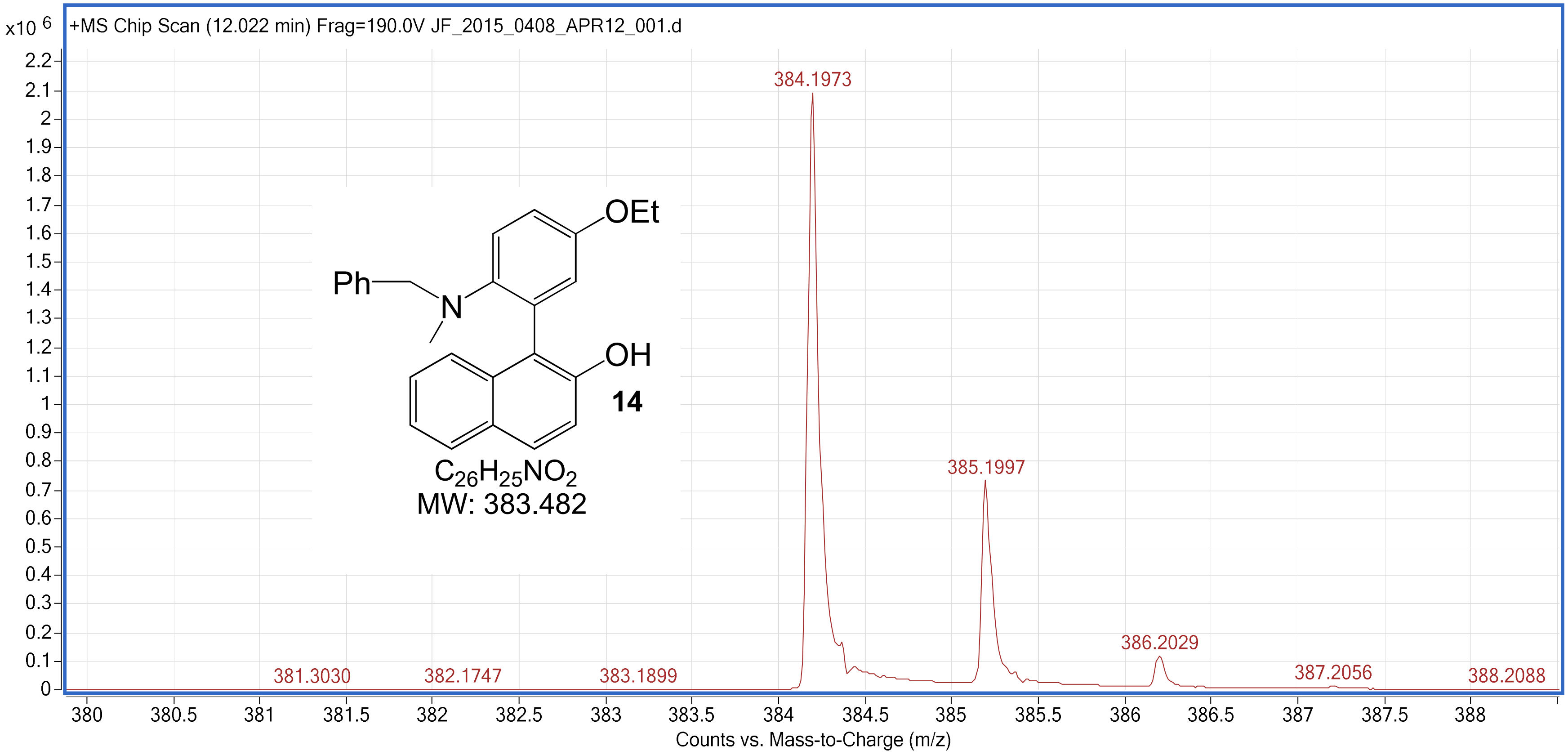

*1-[2-Amino-N-((2E)but-2-enyl)-5-ethoxy-N-methylphenyl]naphthalen-2-ol* ***(15)*** – [1]H NMR ($CDCl_3$, 400 MHz)

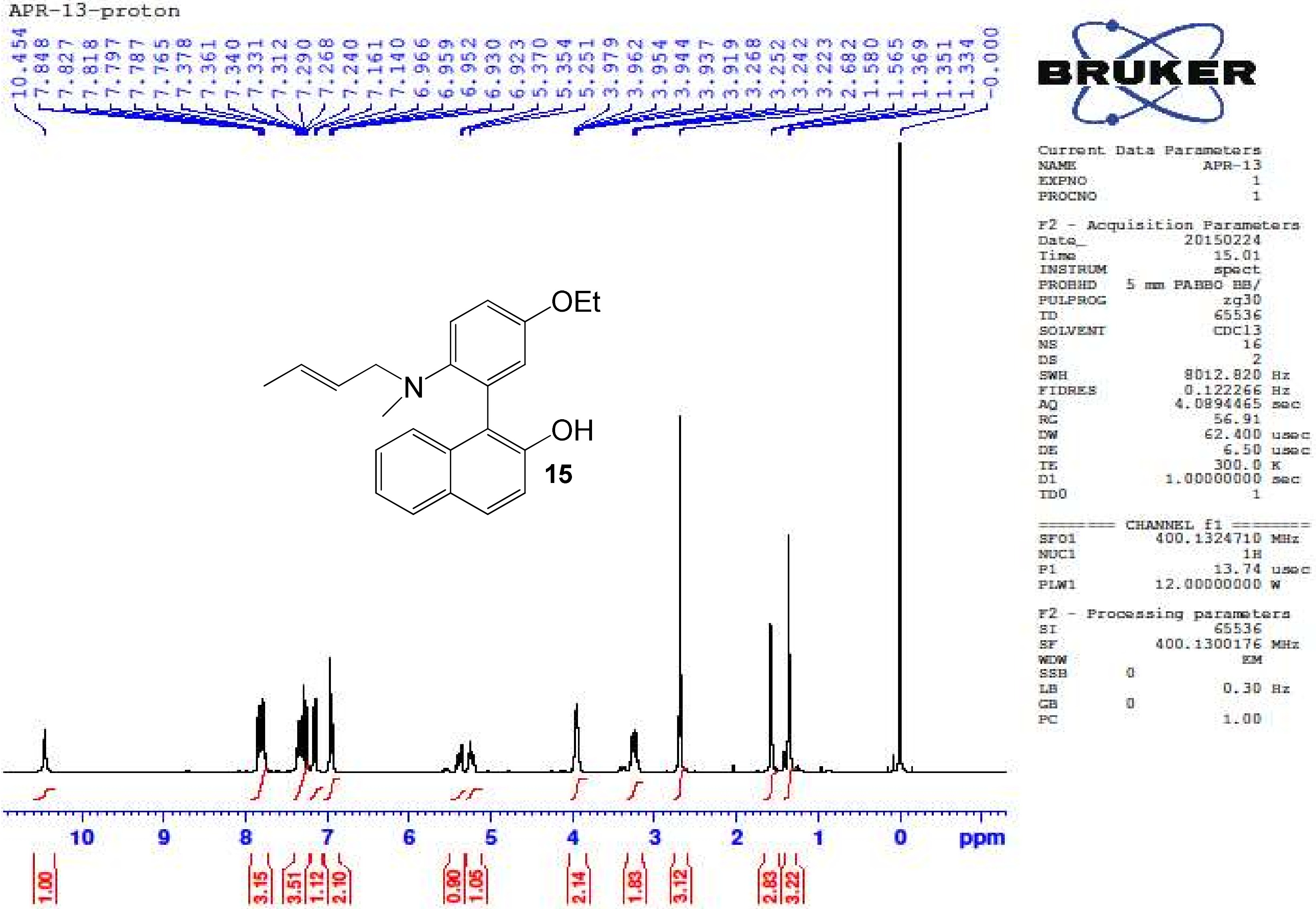

*1-[2-Amino-N-((2E)but-2-enyl)-5-ethoxy-N-methylphenyl]naphthalen-2-ol* ***(15)*** – $^{13}$C NMR ($CDCl_3$, 100 MHz)

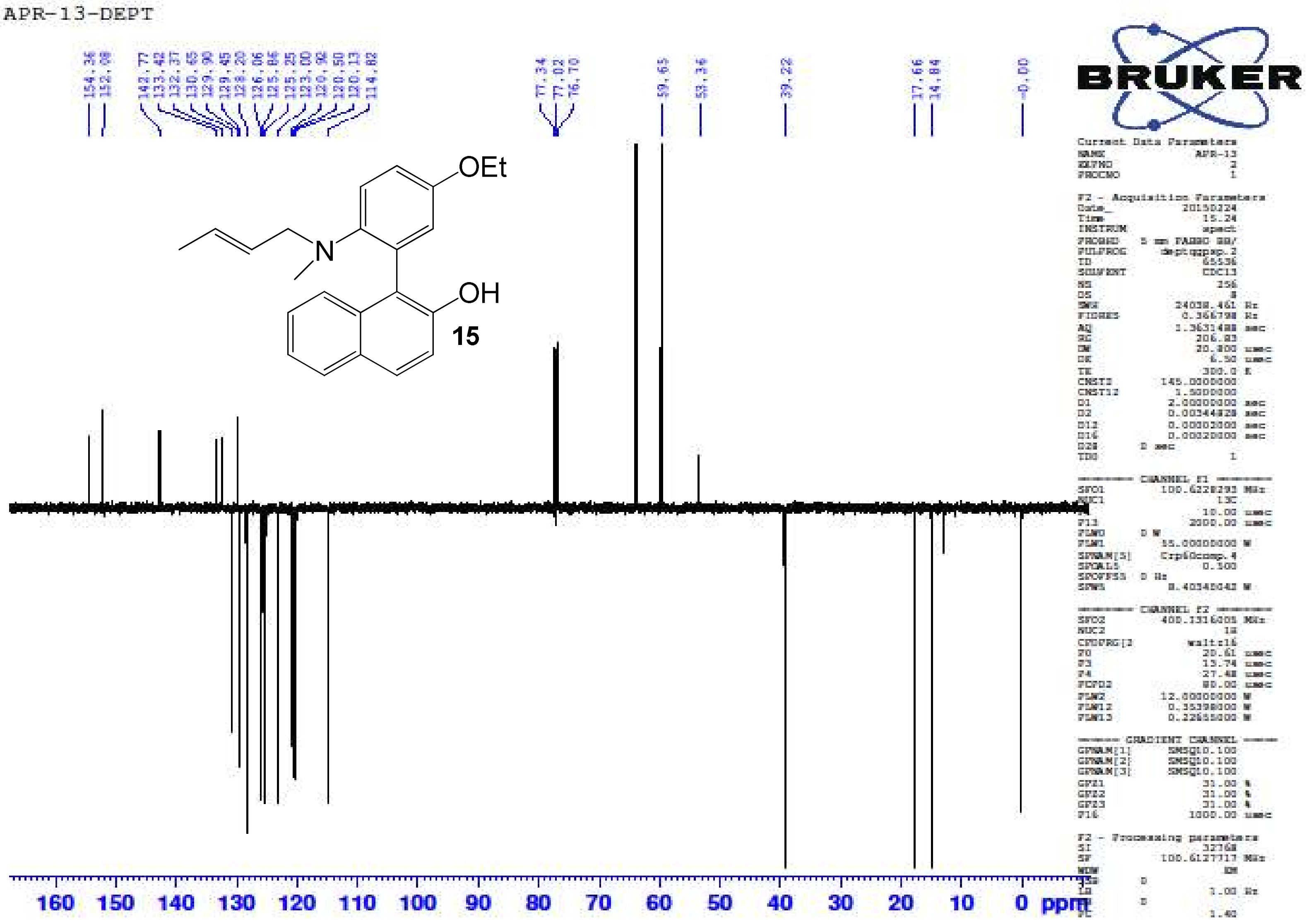

*1-[2-Amino-N-((2E)but-2-enyl)-5-ethoxy-N-methylphenyl]naphthalen-2-ol* ***(15)*** – HR-ESIMS ($[M+H]^+$ Calculated: 348.1958, Observed 348.1962)

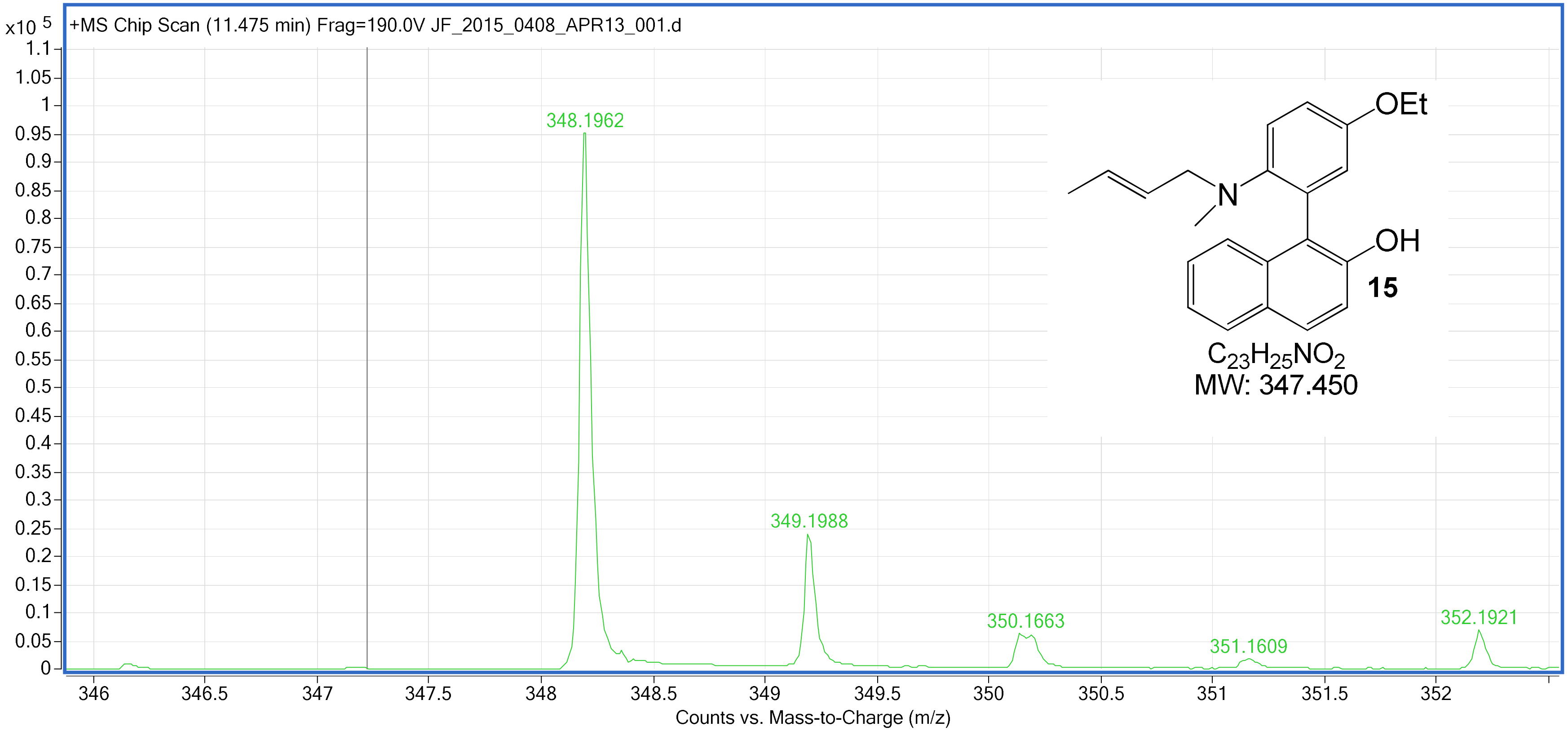

*2-(2-Amino-5,N,N-trimethylphenyl)naphthalen-1-ol* ***(16)*** – [1]H NMR ($CDCl_3$, 400 MHz)

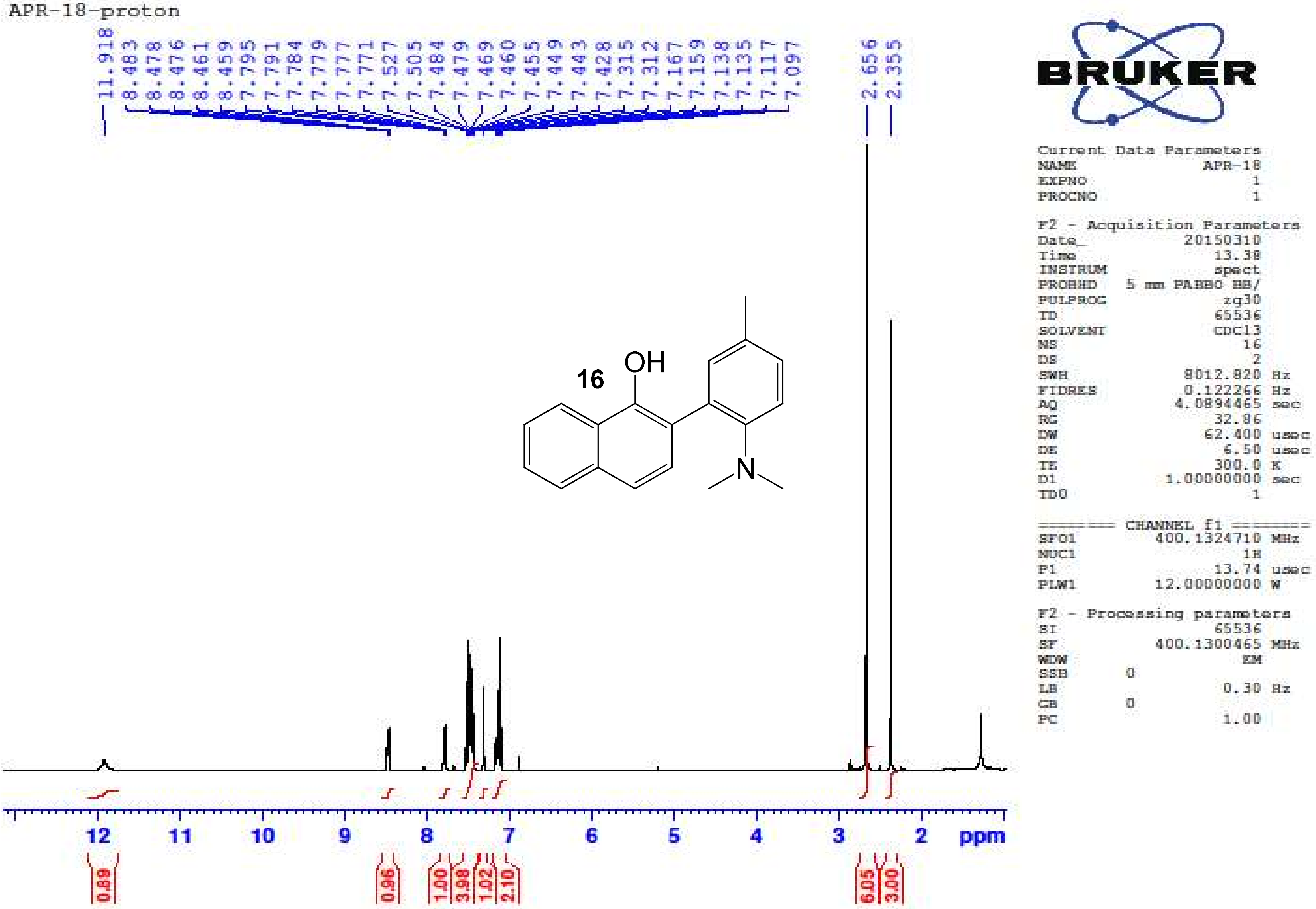

*2-(2-Amino-5,N,N-trimethylphenyl)naphthalen-1-ol* ***(16)*** – $^{13}$C NMR ($CDCl_3$, 100 MHz)

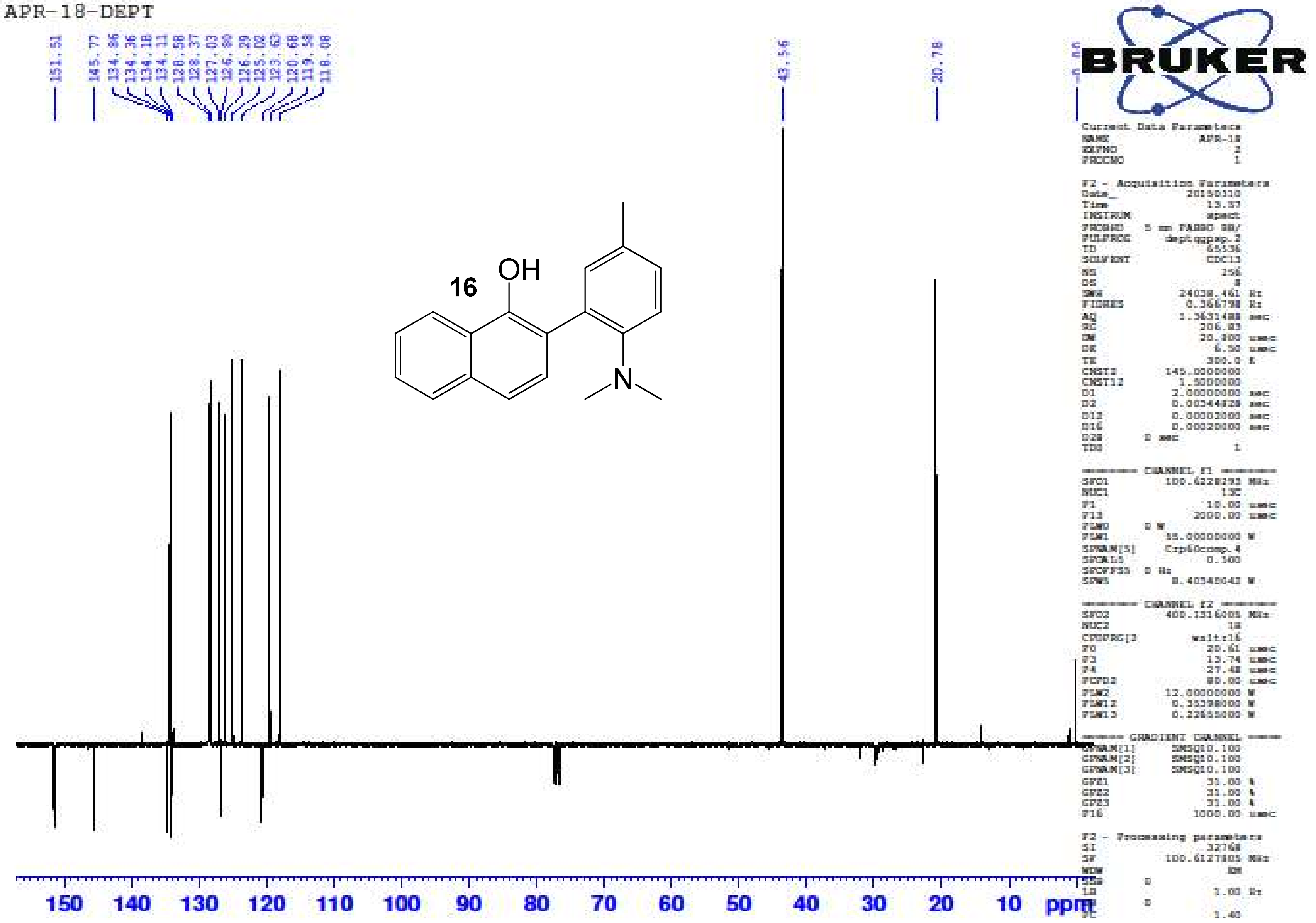

*2-(2-Amino-5,N,N-trimethylphenyl)naphthalen-1-ol* ***(16)*** – HR-ESIMS ($[M+H]^+$ Calculated: 278.1539, Observed 278.1544)

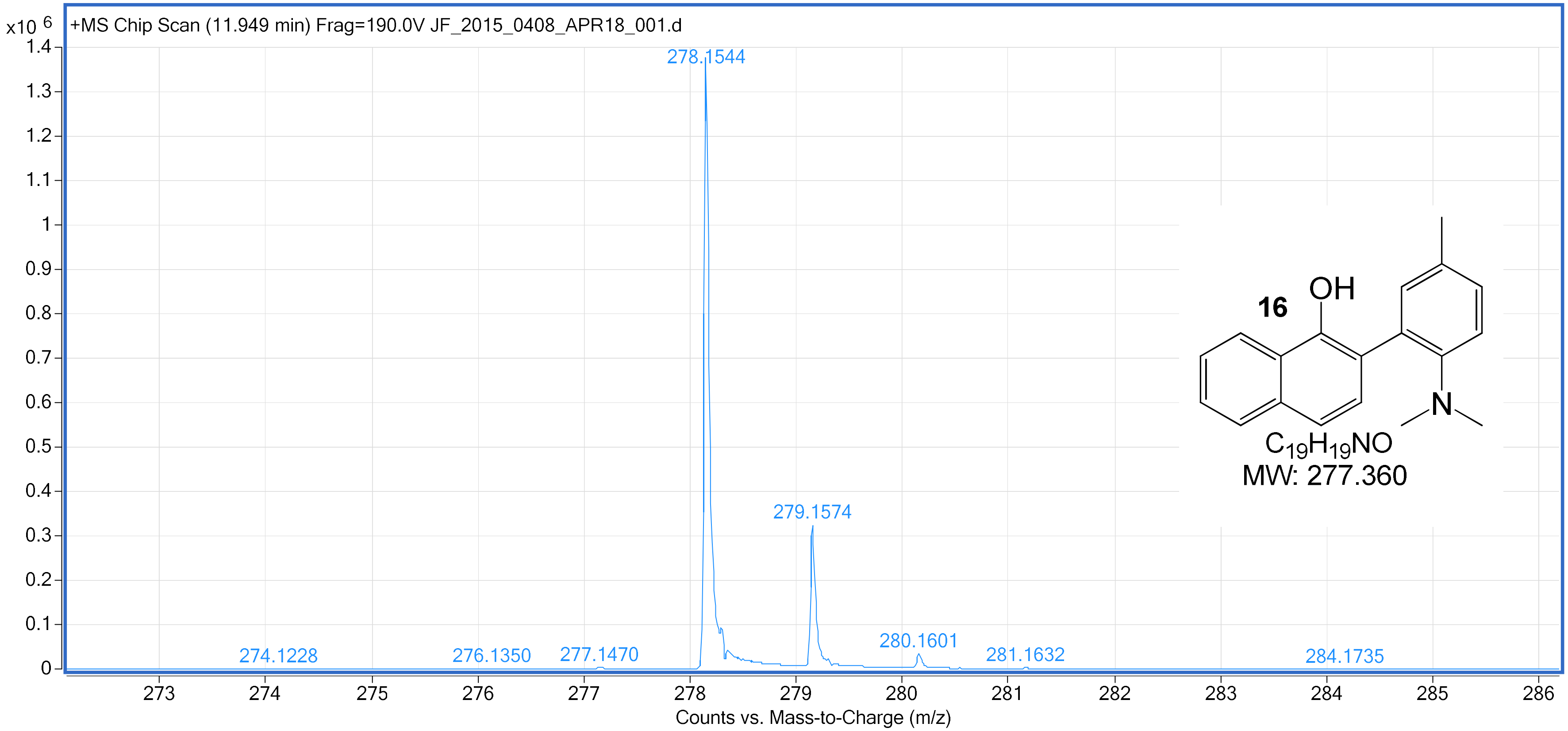

2'-(Dimethylamino)-5'-ethoxybiphenyl-2-ol **(17)** – $^{1}H$ NMR ($CDCl_3$, 400 MHz)

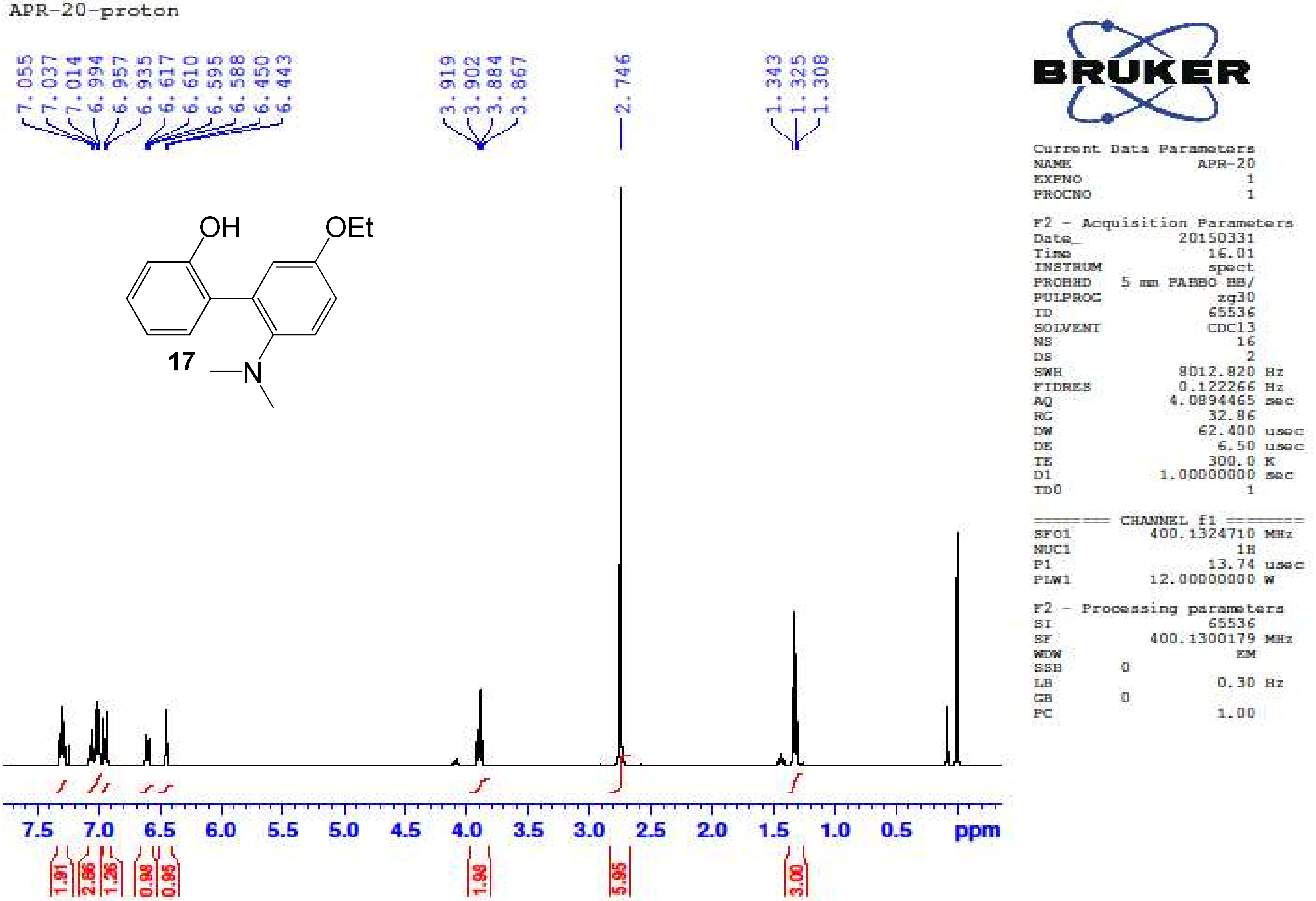

2'-(Dimethylamino)-5'-ethoxybiphenyl-2-ol **(17)** – $^{13}$C NMR ($CDCl_3$, 100 MHz)

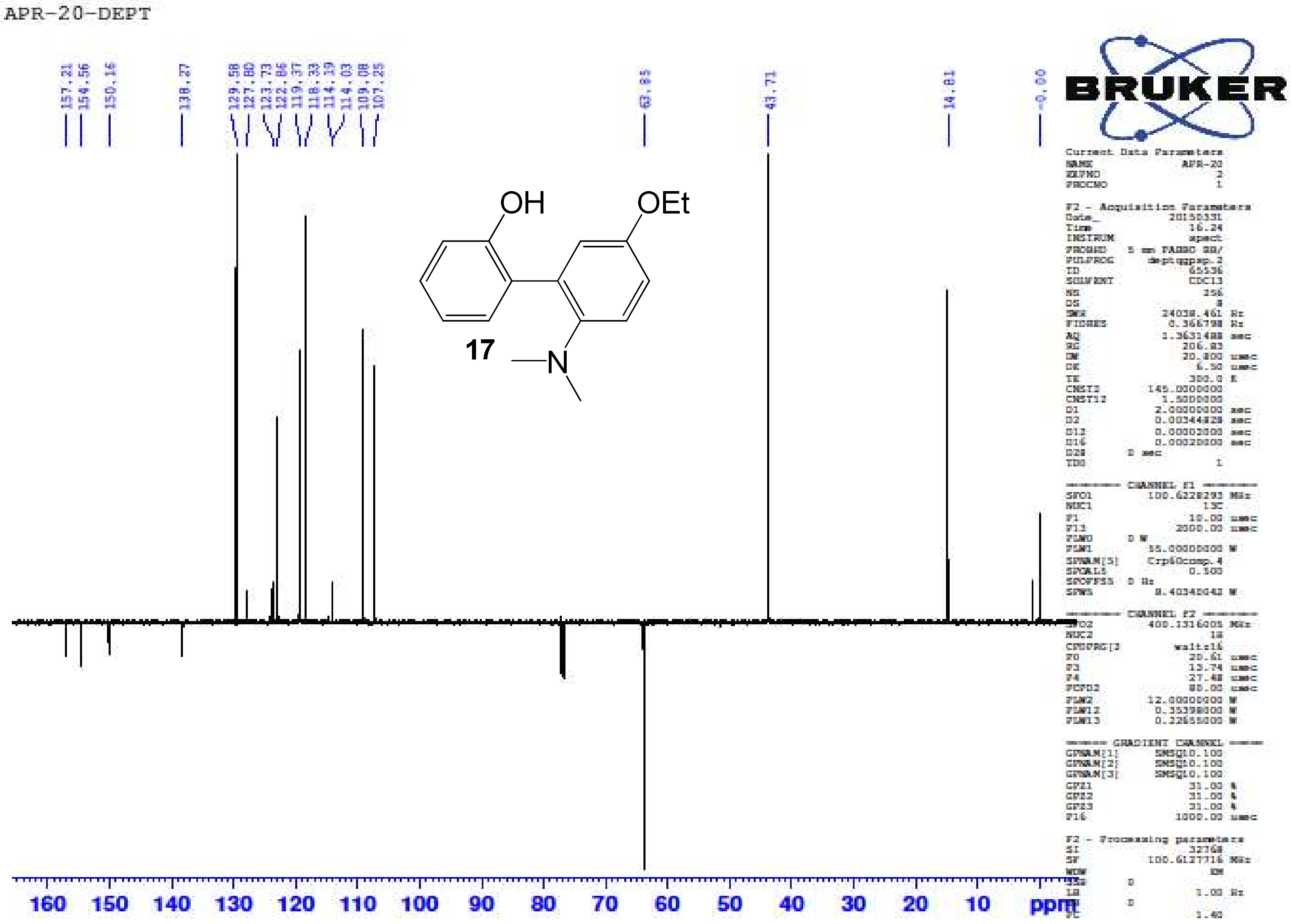

2'-(Dimethylamino)-5'-ethoxybiphenyl-2-ol **(17)** – HR-ESIMS ($[M+H]^+$ Calculated: 258.1489, Observed 258.1498)

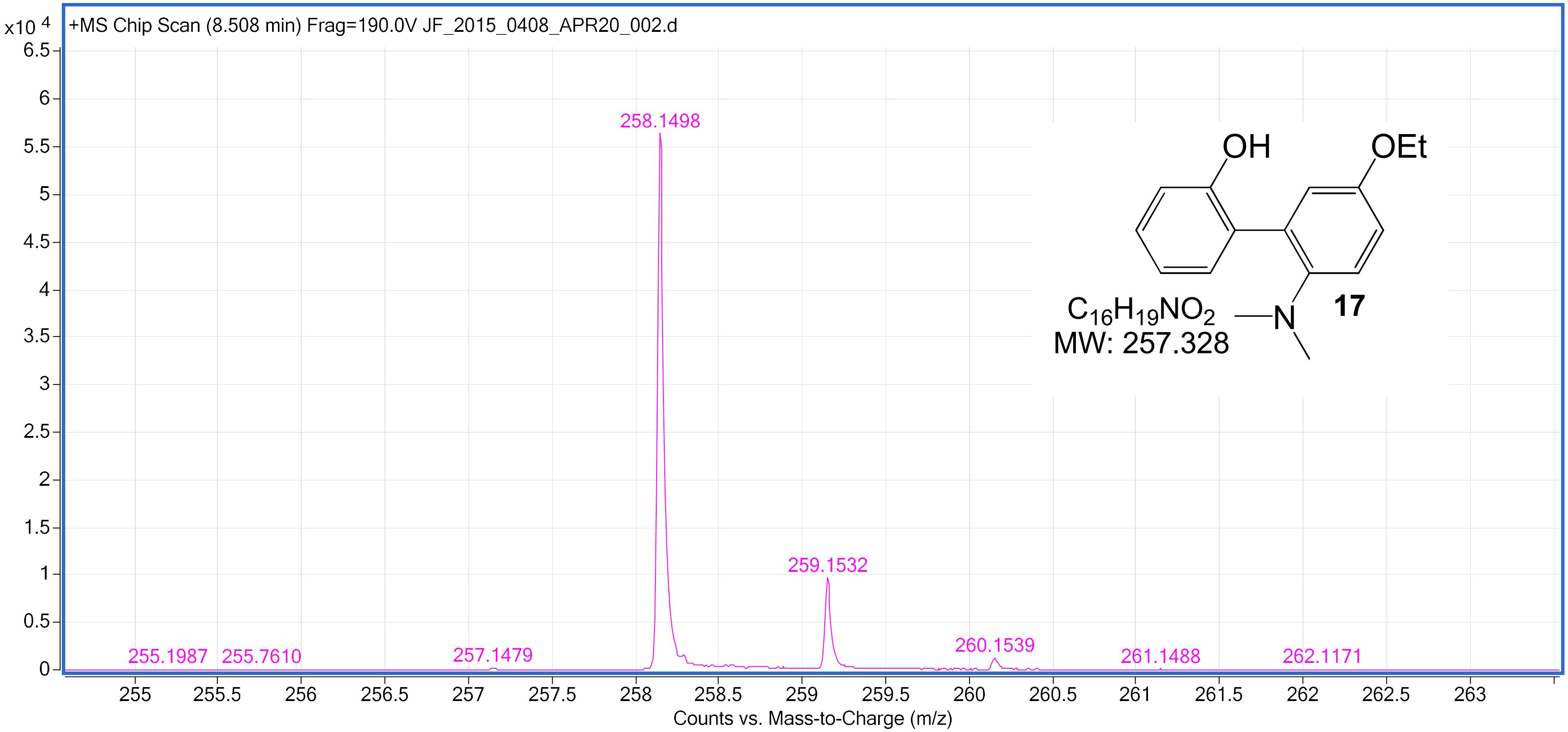

5-Bromo-2'-(dimethylamino)-5'-ethoxybiphenyl-2-ol **(18)** – $^{1}H$ NMR ($CDCl_3$, 400 MHz)

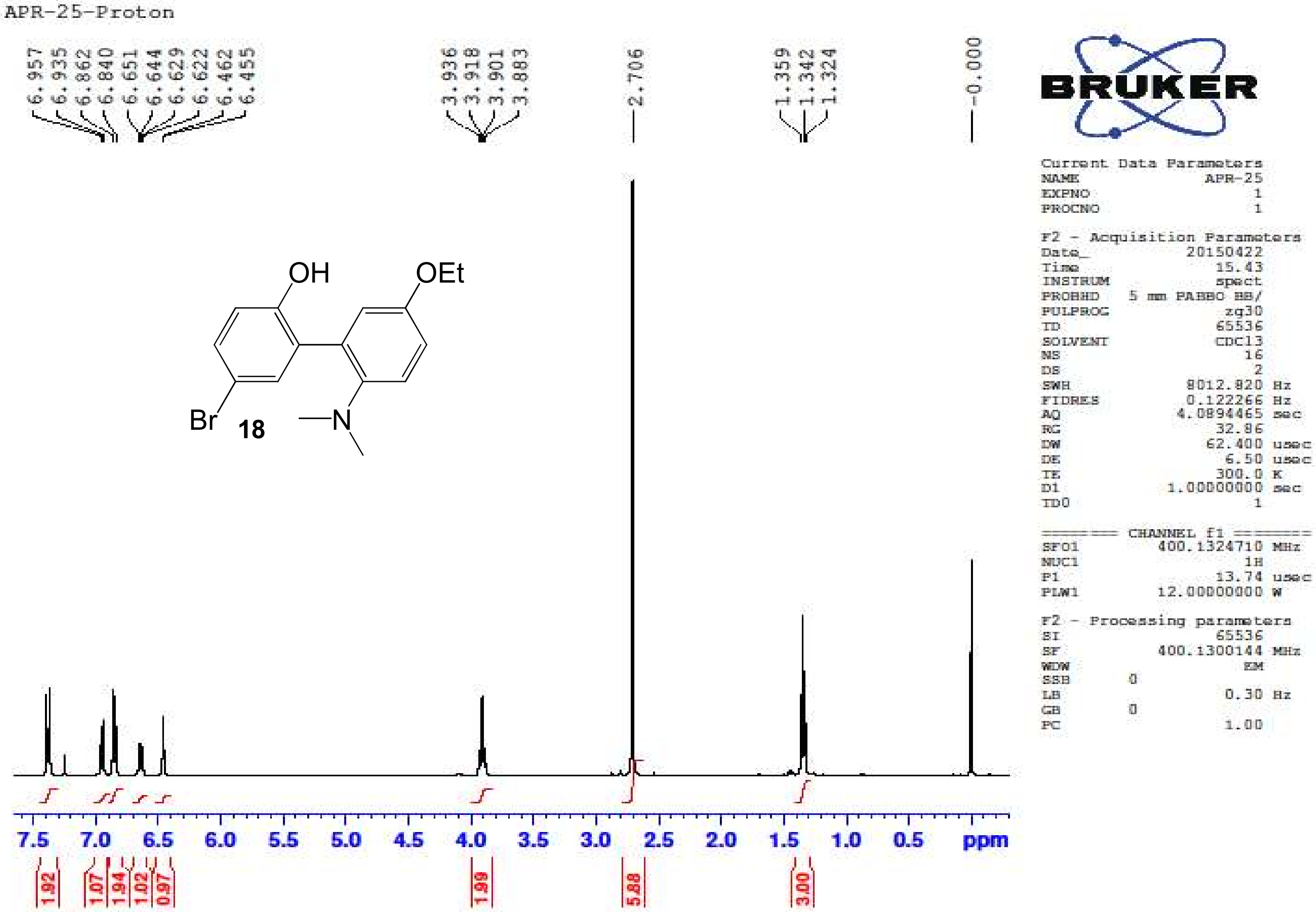

5-Bromo-2'-(dimethylamino)-5'-ethoxybiphenyl-2-ol **(18)** – $^{13}C$ NMR ($CDCl_3$, 100 MHz)

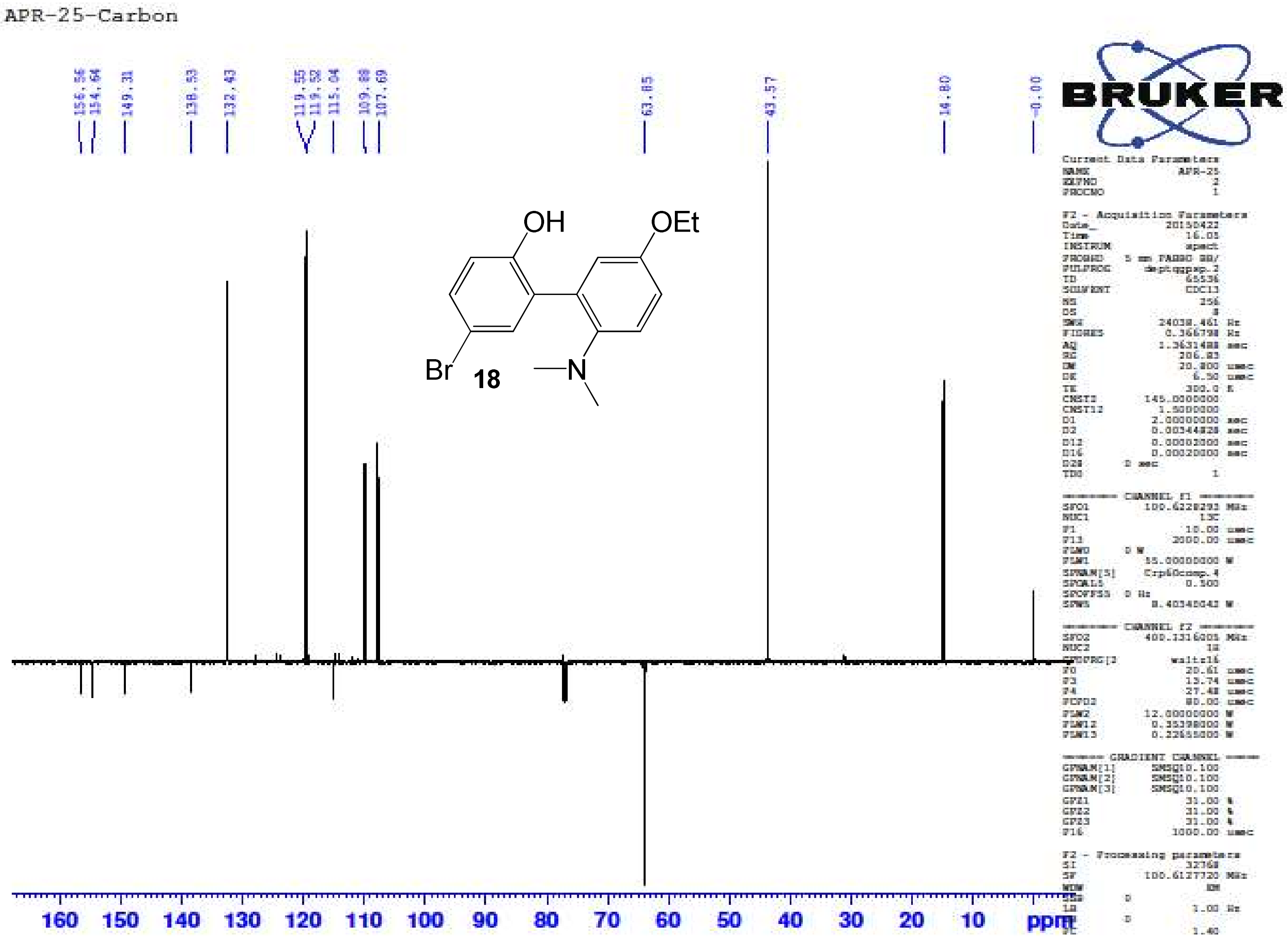

5-Bromo-2'-(dimethylamino)-5'-ethoxybiphenyl-2-ol **(18)** – HR-ESIMS ([M+] Calculated: 336.0594, Observed: 336.0615)

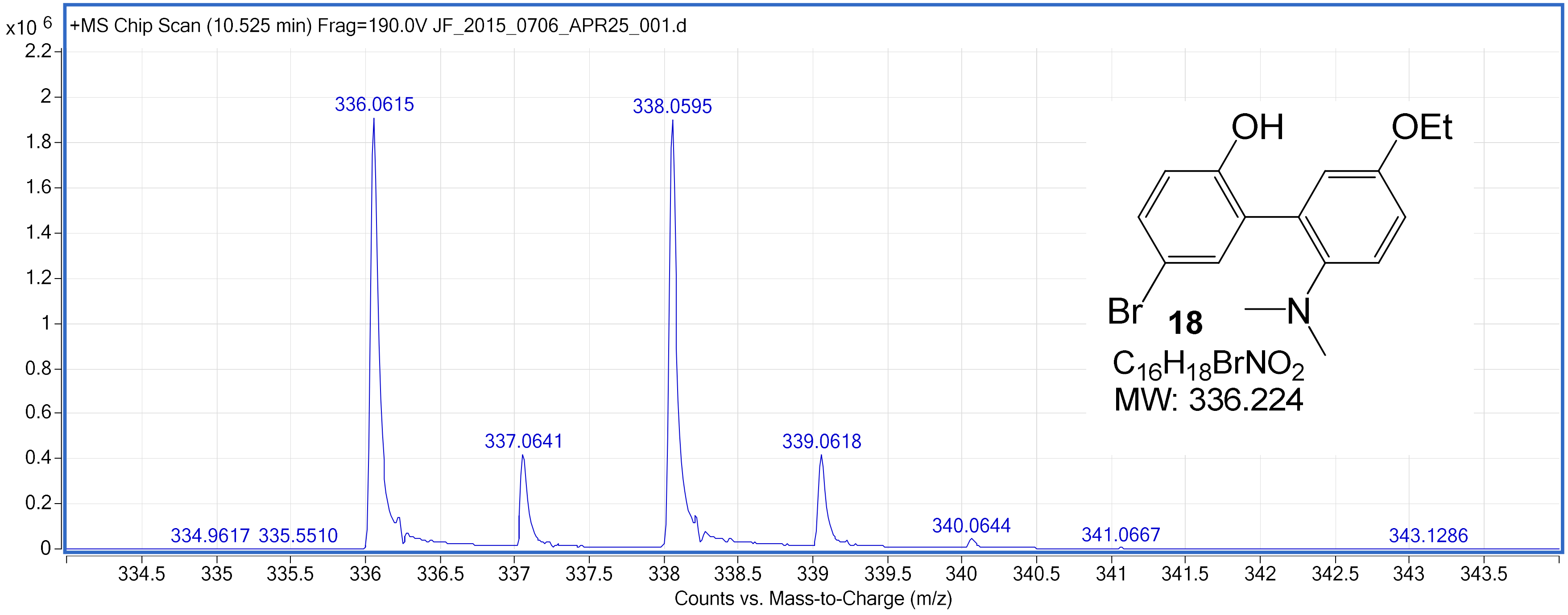

5-Chloro-2'-(dimethylamino)-5'-ethoxybiphenyl-2-ol **(19)** – $^{1}$H NMR ($CDCl_3$, 400 MHz)

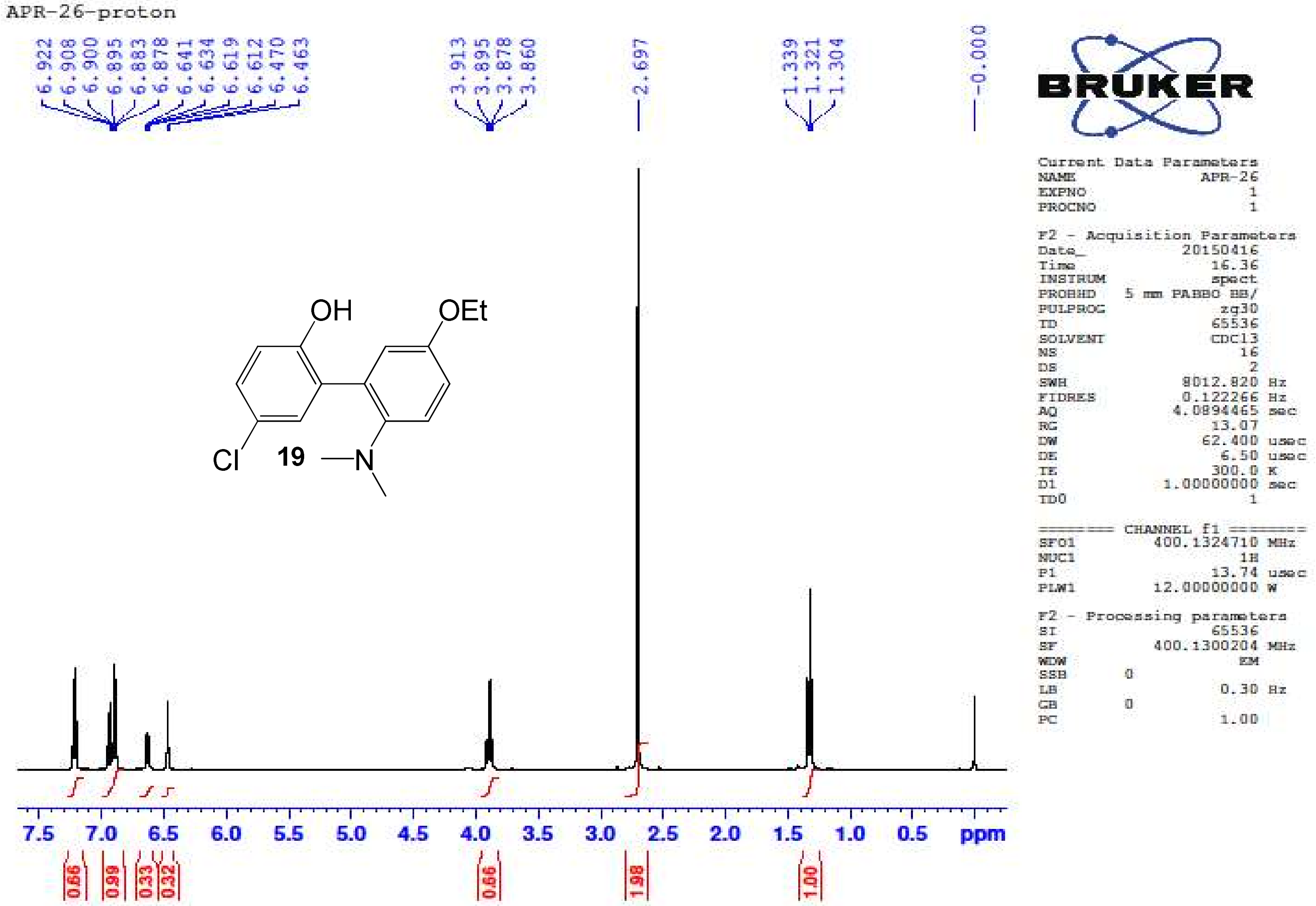

5-Chloro-2'-(dimethylamino)-5'-ethoxybiphenyl-2-ol **(19)** – $^{13}$C NMR ($CDCl_3$, 100 MHz)

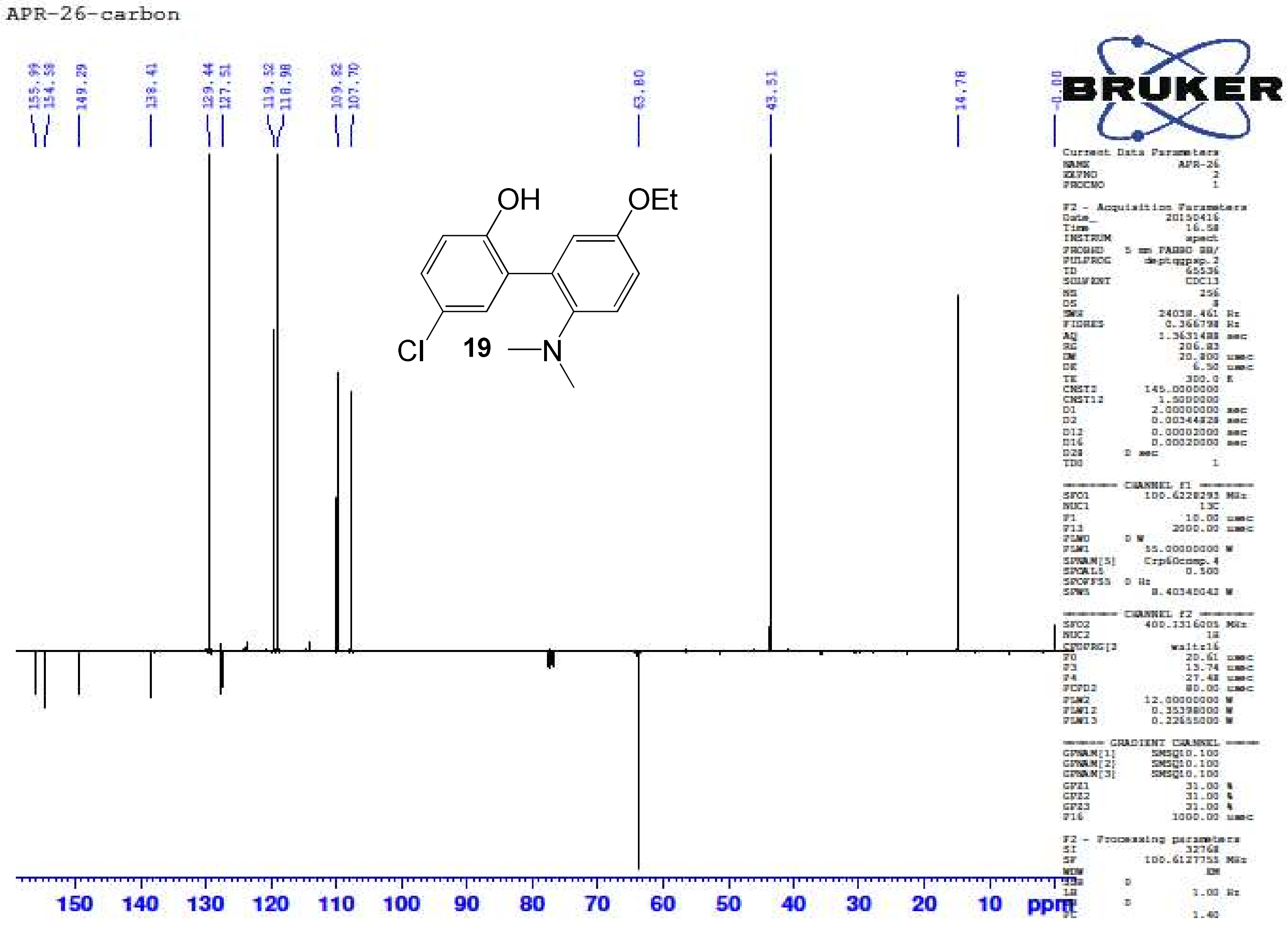

5-Chloro-2'-(dimethylamino)-5'-ethoxybiphenyl-2-ol **(19)** – HR-ESIMS ($[M+H]^+$ Calculated: 292.1099, Observed 292.1116)

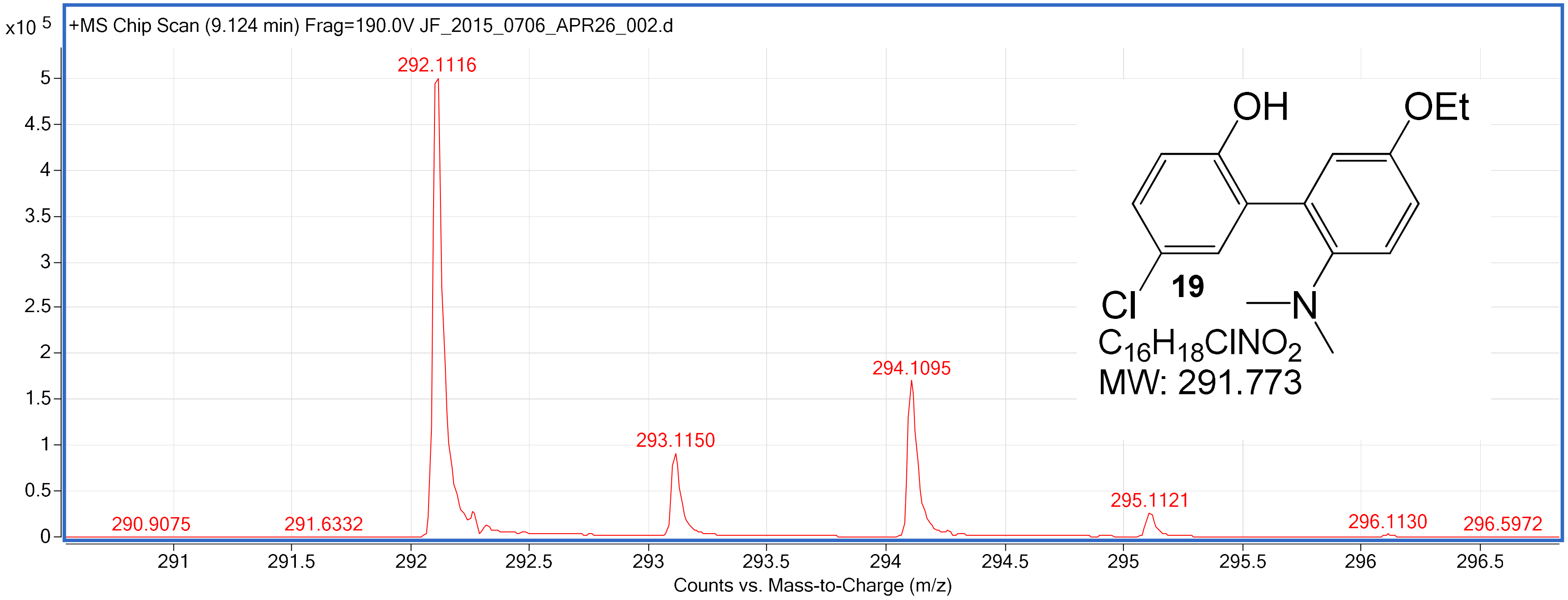

2'-(Dimethylamino)-5'-ethoxy-5-methoxybiphenyl-2-ol **(20)** – $^{1}H$ NMR ($CDCl_3$, 400 MHz)

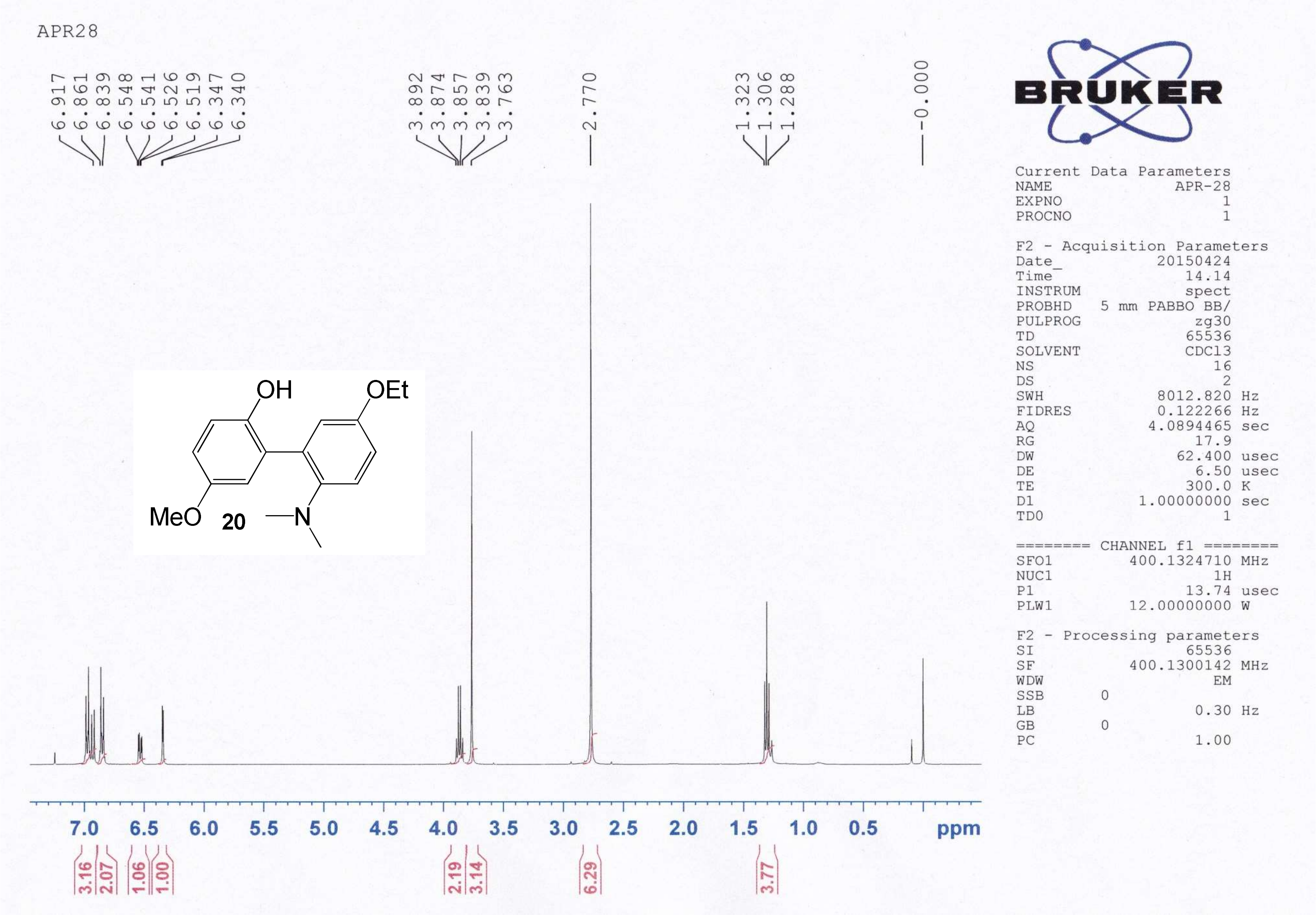

2'-(Dimethylamino)-5'-ethoxy-5-methoxybiphenyl-2-ol **(20)** – $^{13}$C NMR ($CDCl_3$, 100 MHz)

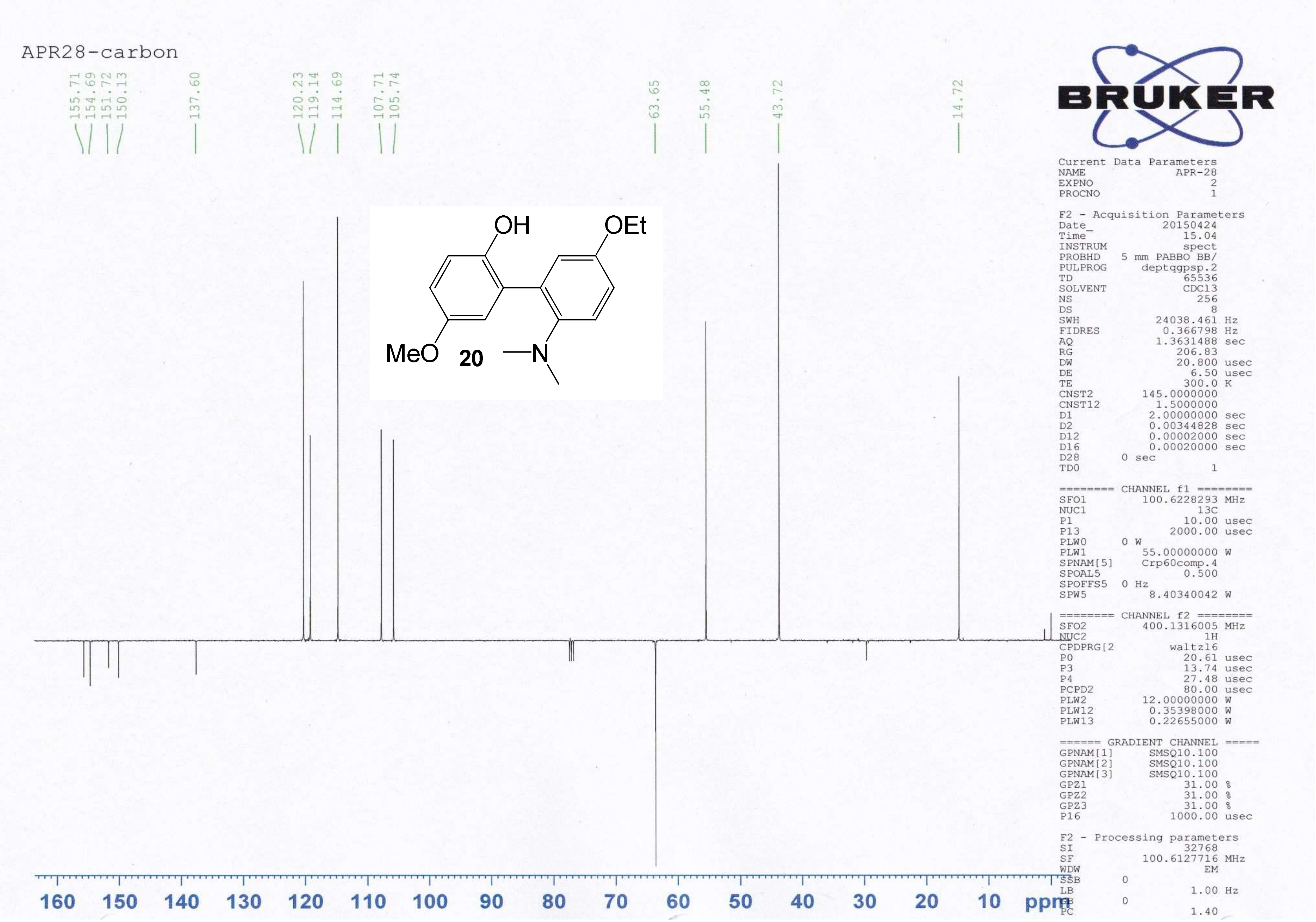

2'-(Dimethylamino)-5'-ethoxy-5-methoxybiphenyl-2-ol **(20)** – HR-ESIMS ($[M+H]^+$ Calculated: 288.1594, Observed: 288.1596)

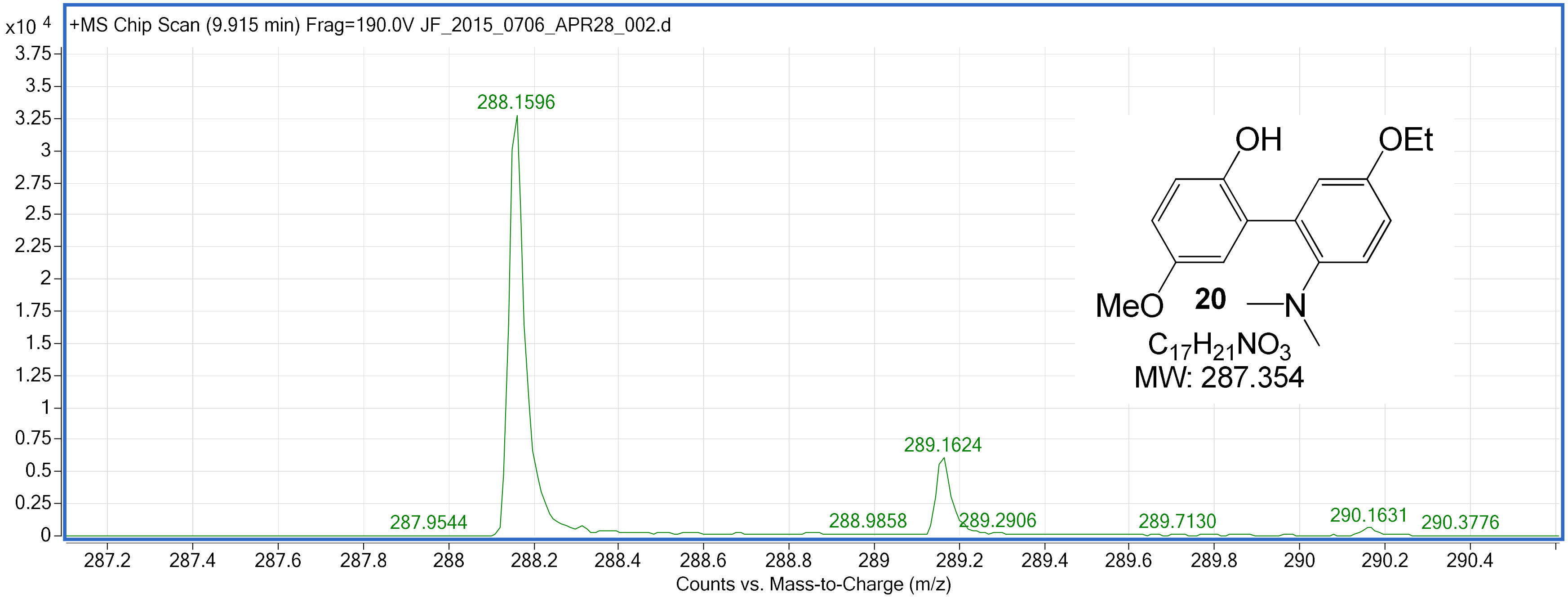

2'-(Dimethylamino)-5'-ethoxy-5-methylbiphenyl-2-ol **(21)** – $^{1}H$ NMR ($CDCl_3$, 400 MHz)

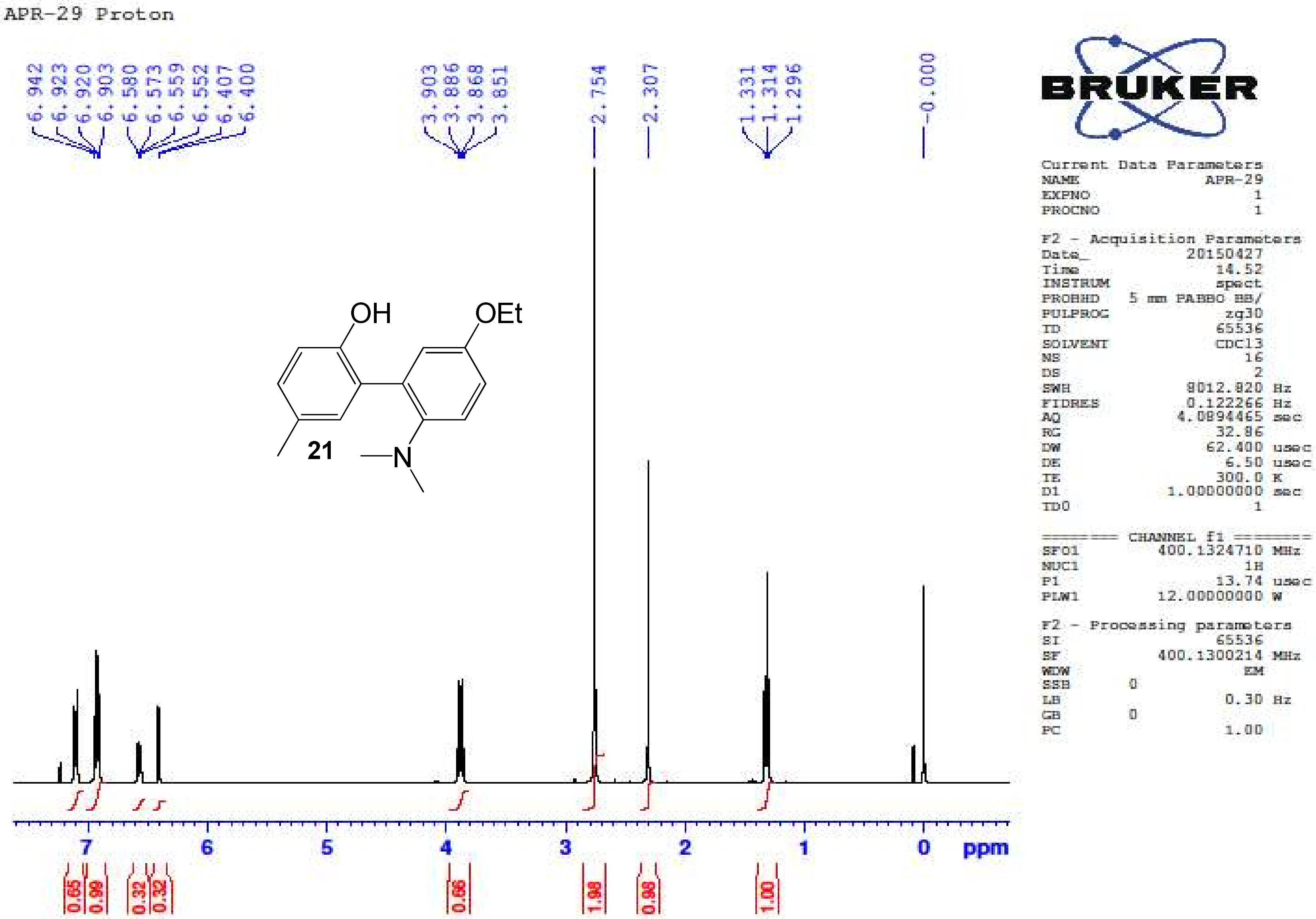

2'-(Dimethylamino)-5'-ethoxy-5-methylbiphenyl-2-ol **(21)** – $^{13}$C NMR ($CDCl_3$, 100 MHz)

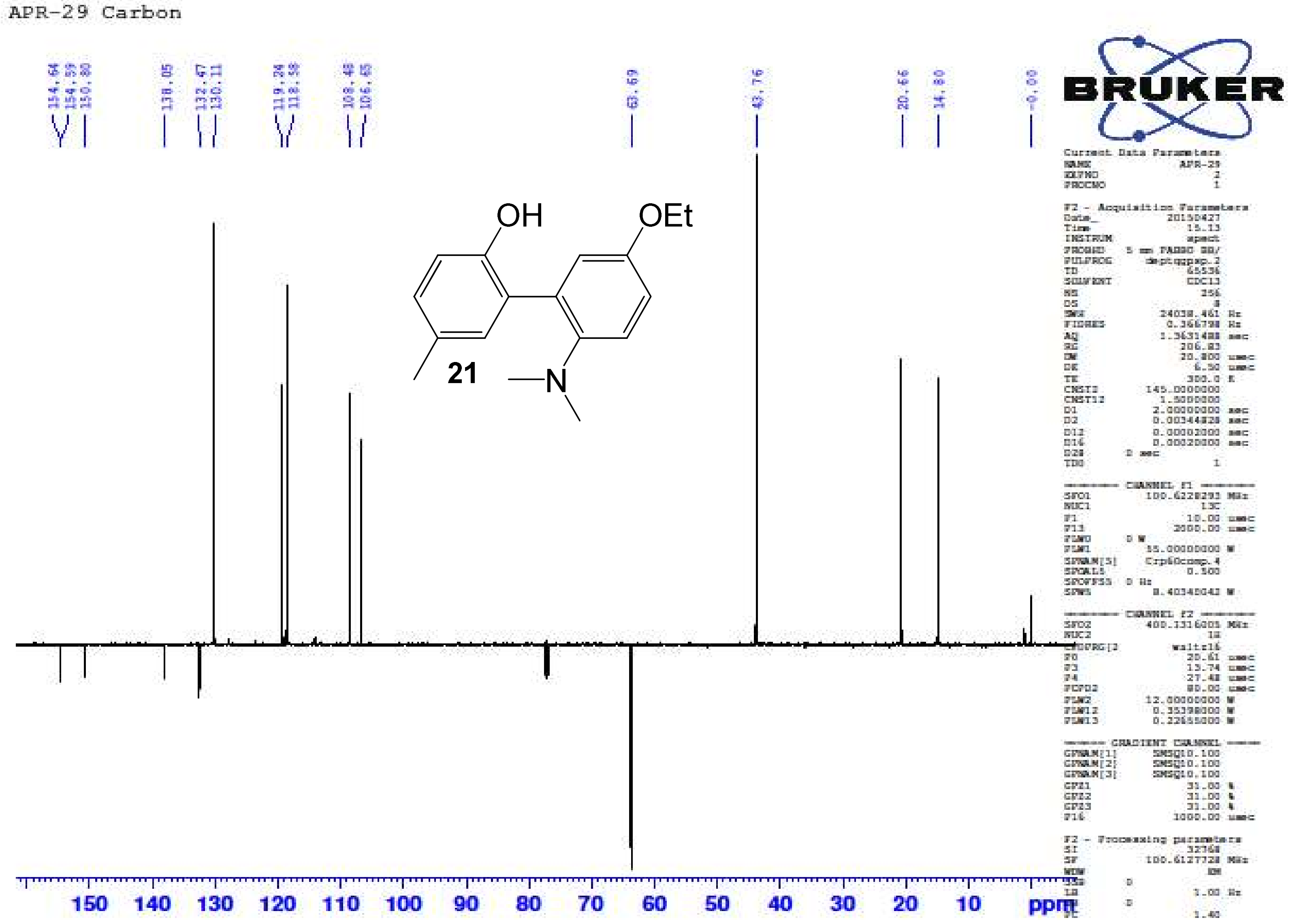

2'-(Dimethylamino)-5'-ethoxy-5-methylbiphenyl-2-ol **(21)** – HR-ESIMS ($[M+H]^+$ Calculated: 272.1645, Observed 272.1656)

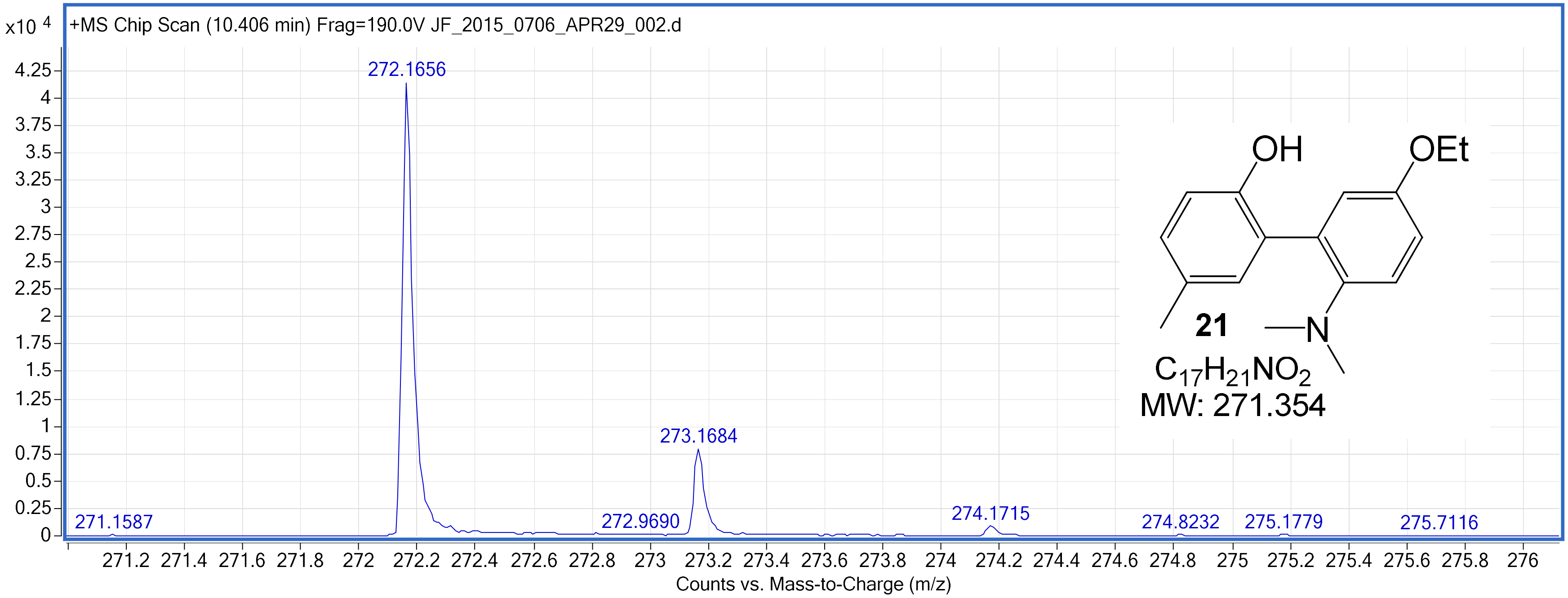

2'-(Dimethylamino)-5'-ethoxy-5-isopropylbiphenyl-2-ol **(22)** – $^{1}H$ NMR ($CDCl_3$, 400 MHz)

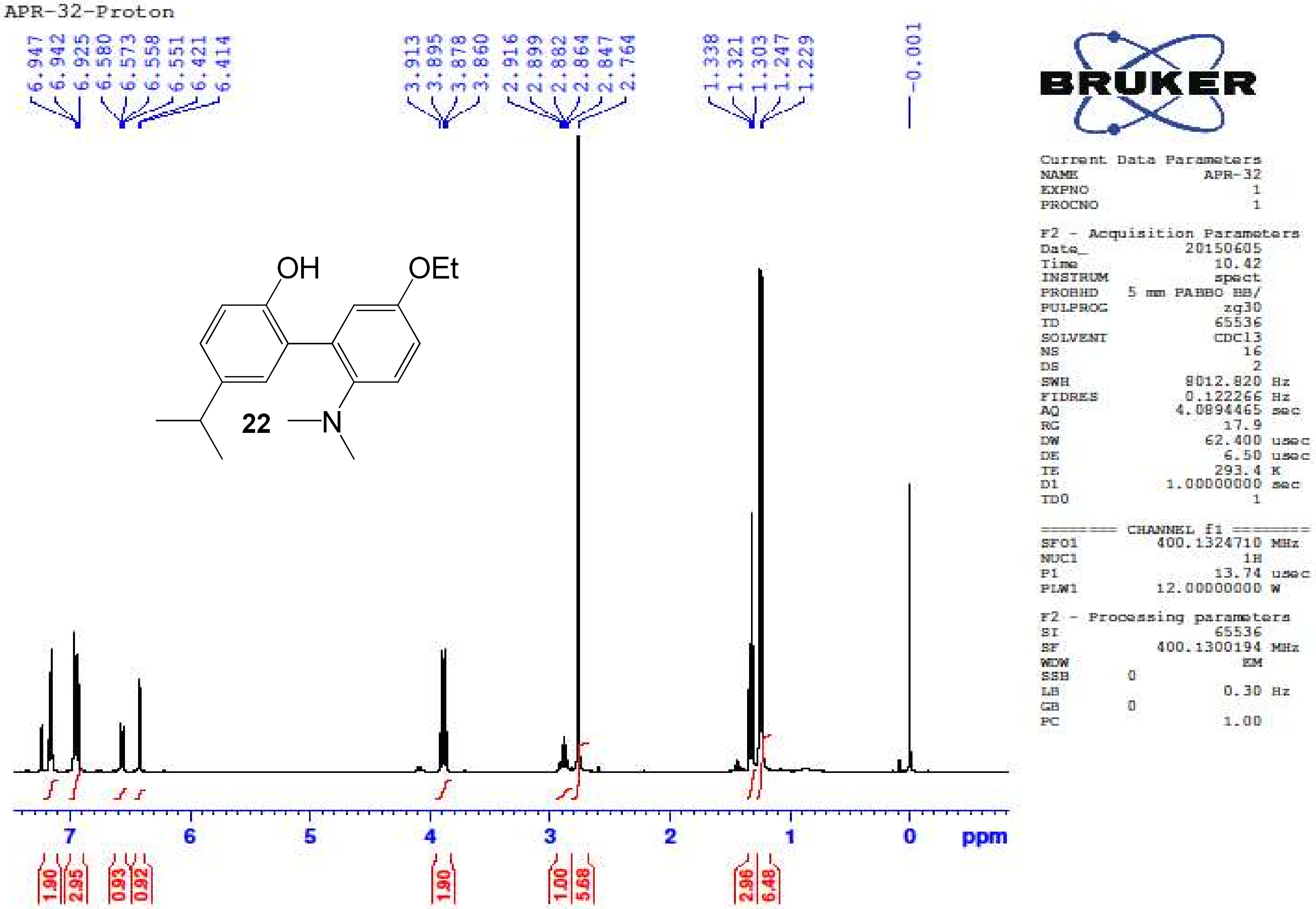

2'-(Dimethylamino)-5'-ethoxy-5-isopropylbiphenyl-2-ol **(22)** – $^{13}$C NMR ($CDCl_3$, 100 MHz)

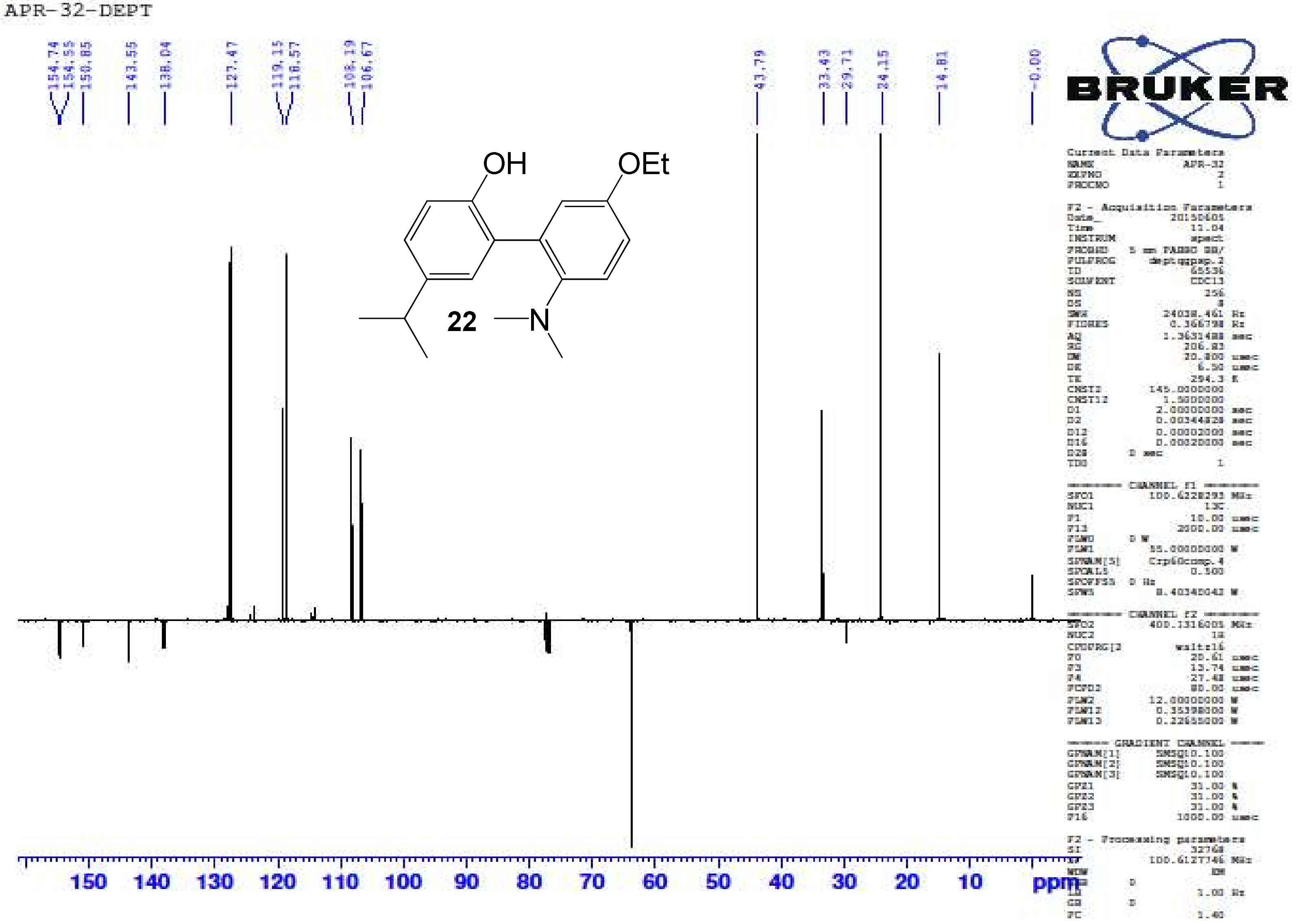

2'-(Dimethylamino)-5'-ethoxy-5-isopropylbiphenyl-2-ol **(22)** – HR-ESIMS ([M+H]$^+$ Calculated: 300.1958, Observed: 300.1953)

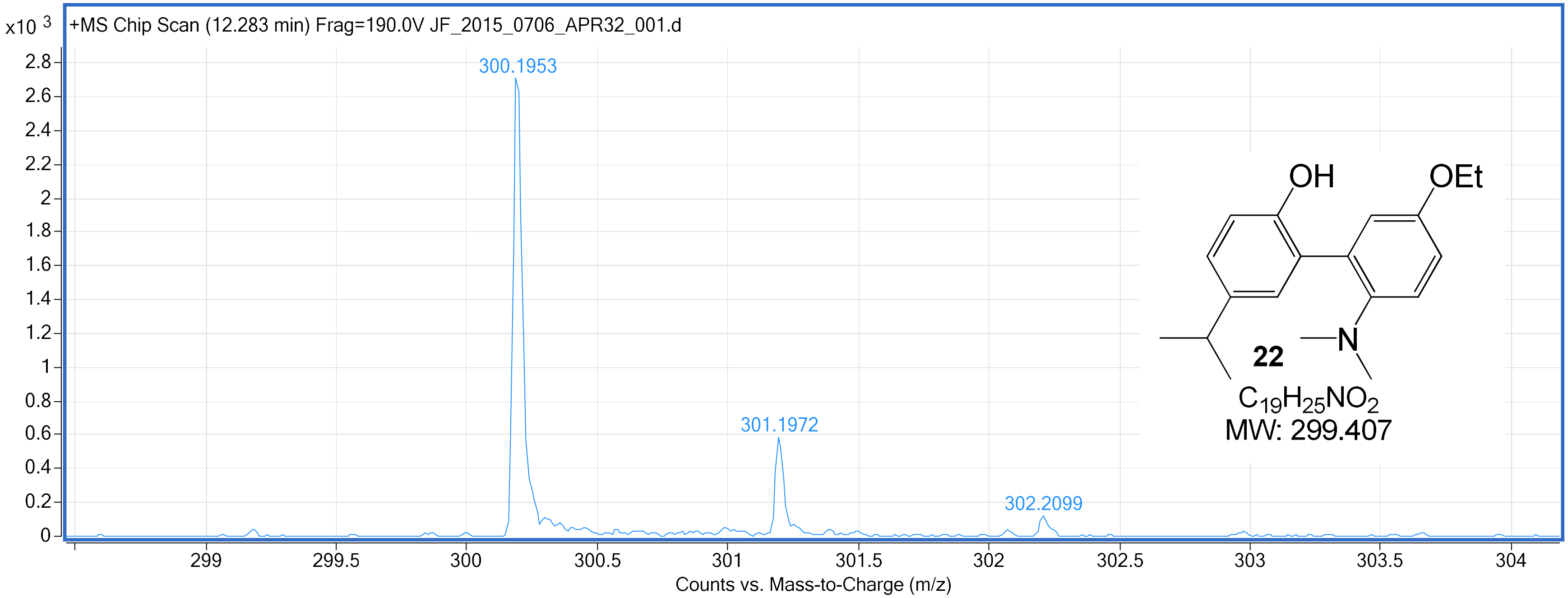

4-Chloro-2'-(dimethylamino)-5'-ethoxybiphenyl-2-ol **(23)** – [1]H NMR ($CDCl_3$, 400 MHz)

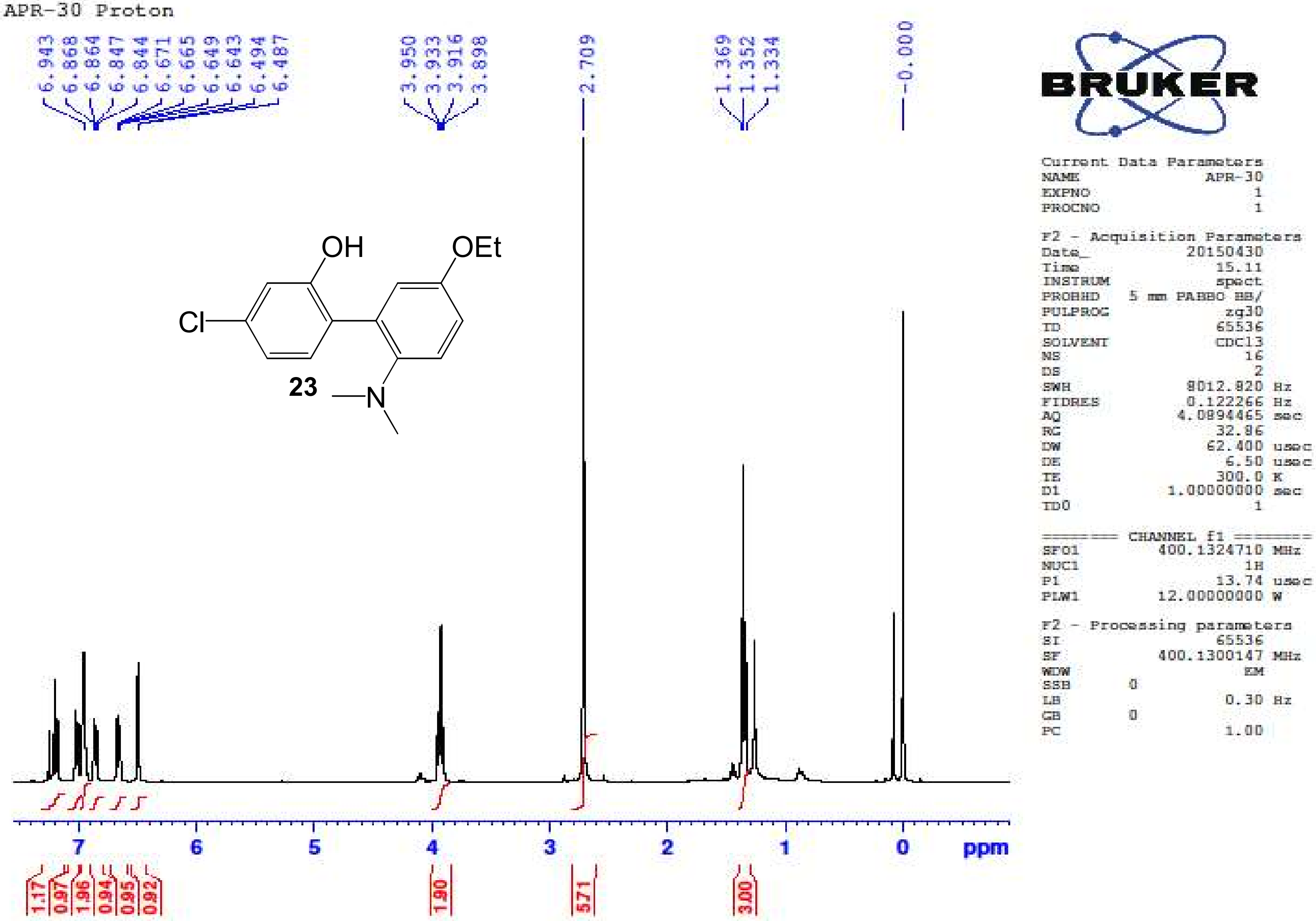

4-Chloro-2'-(dimethylamino)-5'-ethoxybiphenyl-2-ol **(23)** – $^{13}$C NMR (CDCl$_3$, 100 MHz)

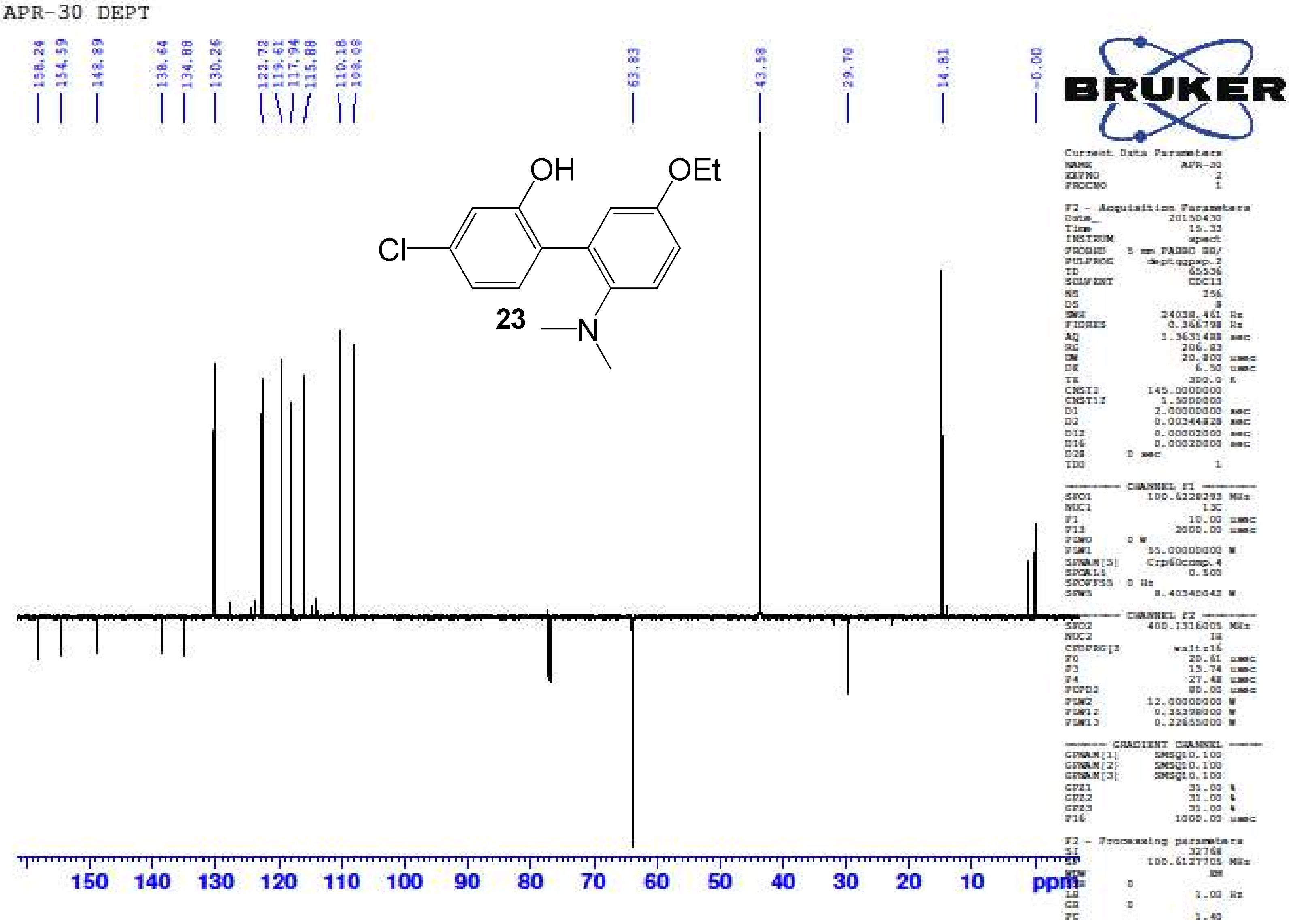

4-Chloro-2'-(dimethylamino)-5'-ethoxybiphenyl-2-ol **(23)** – HR-ESIMS ($[M+H]^+$ Calculated: 292.1099, Observed: 292.1107)

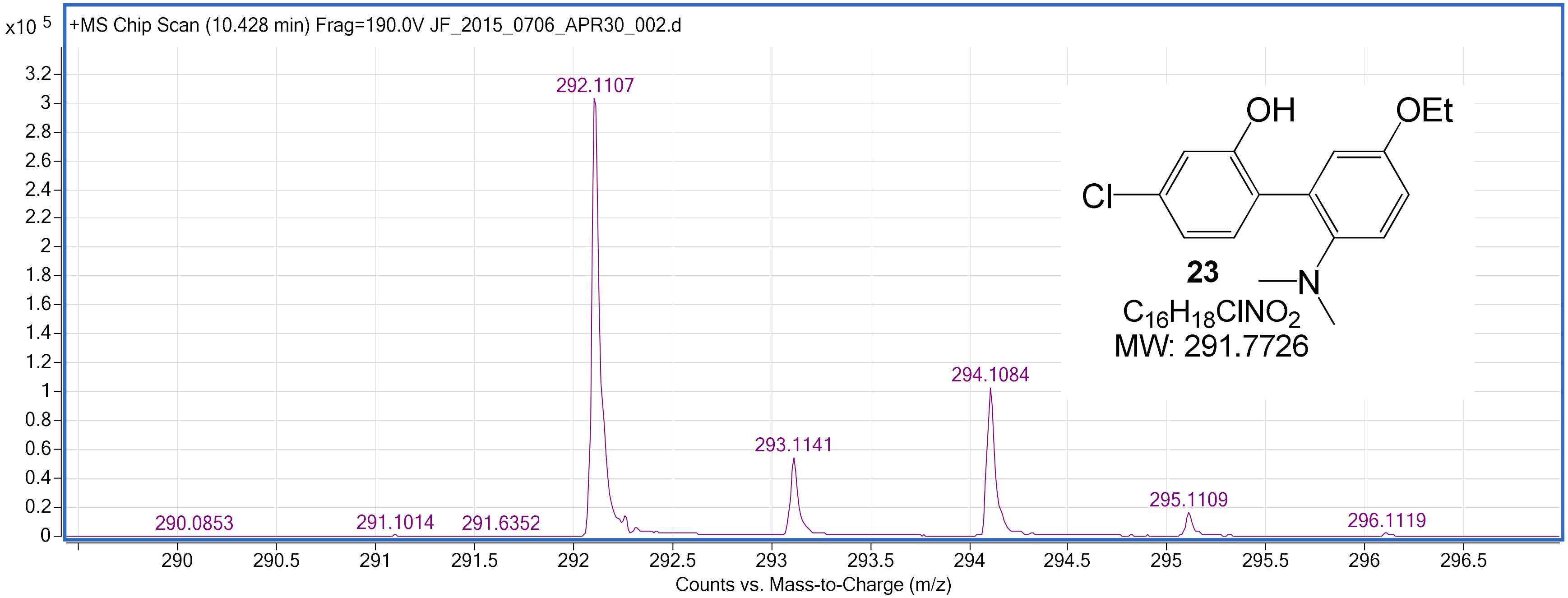

4,5-Dichloro-2'-(dimethylamino)-5'-ethoxybiphenyl-2-ol **(24)** – [1]H NMR ($CDCl_3$, 400 MHz)

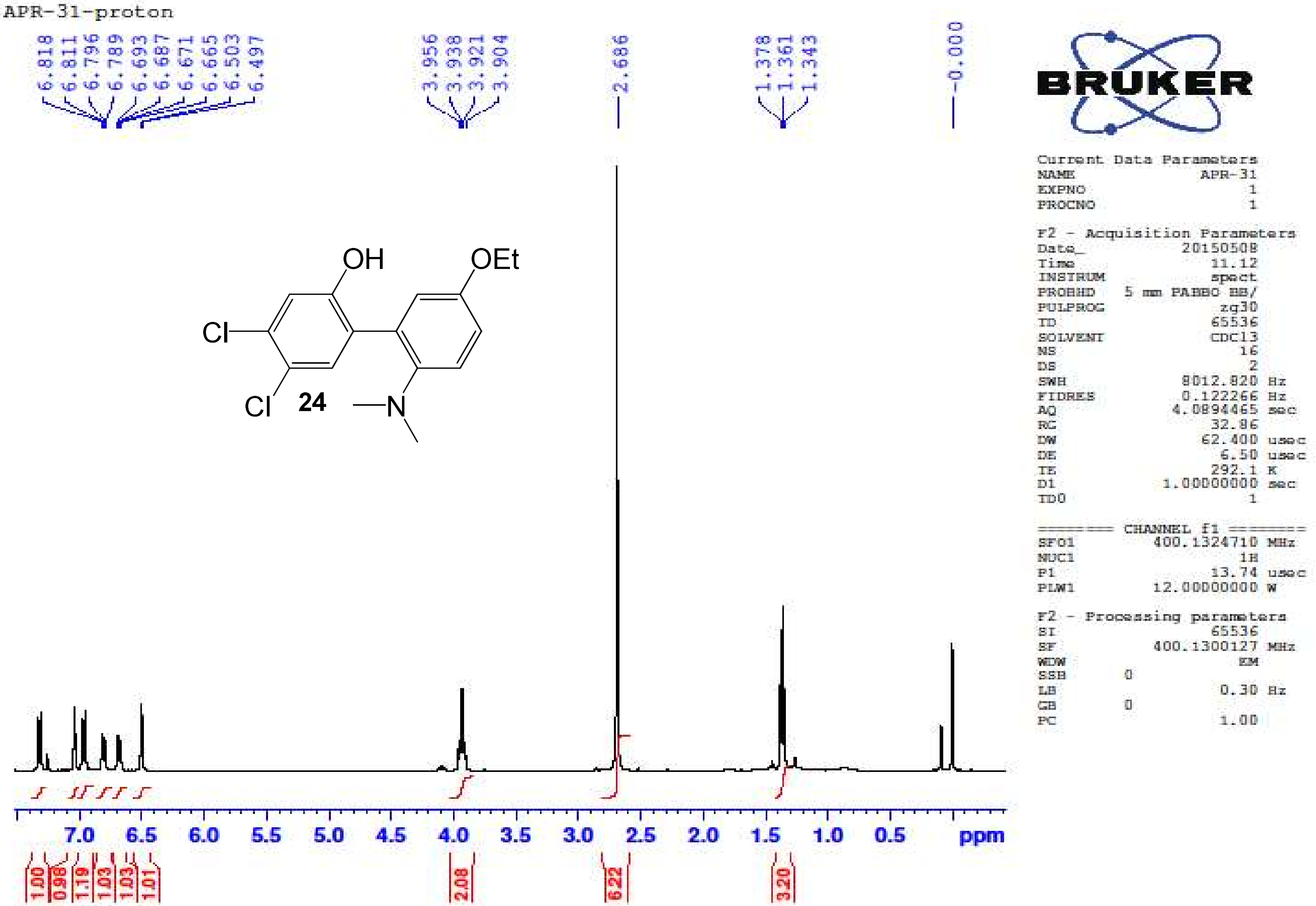

4,5-Dichloro-2'-(dimethylamino)-5'-ethoxybiphenyl-2-ol **(24)** – $^{13}$C NMR ($CDCl_3$, 100 MHz)

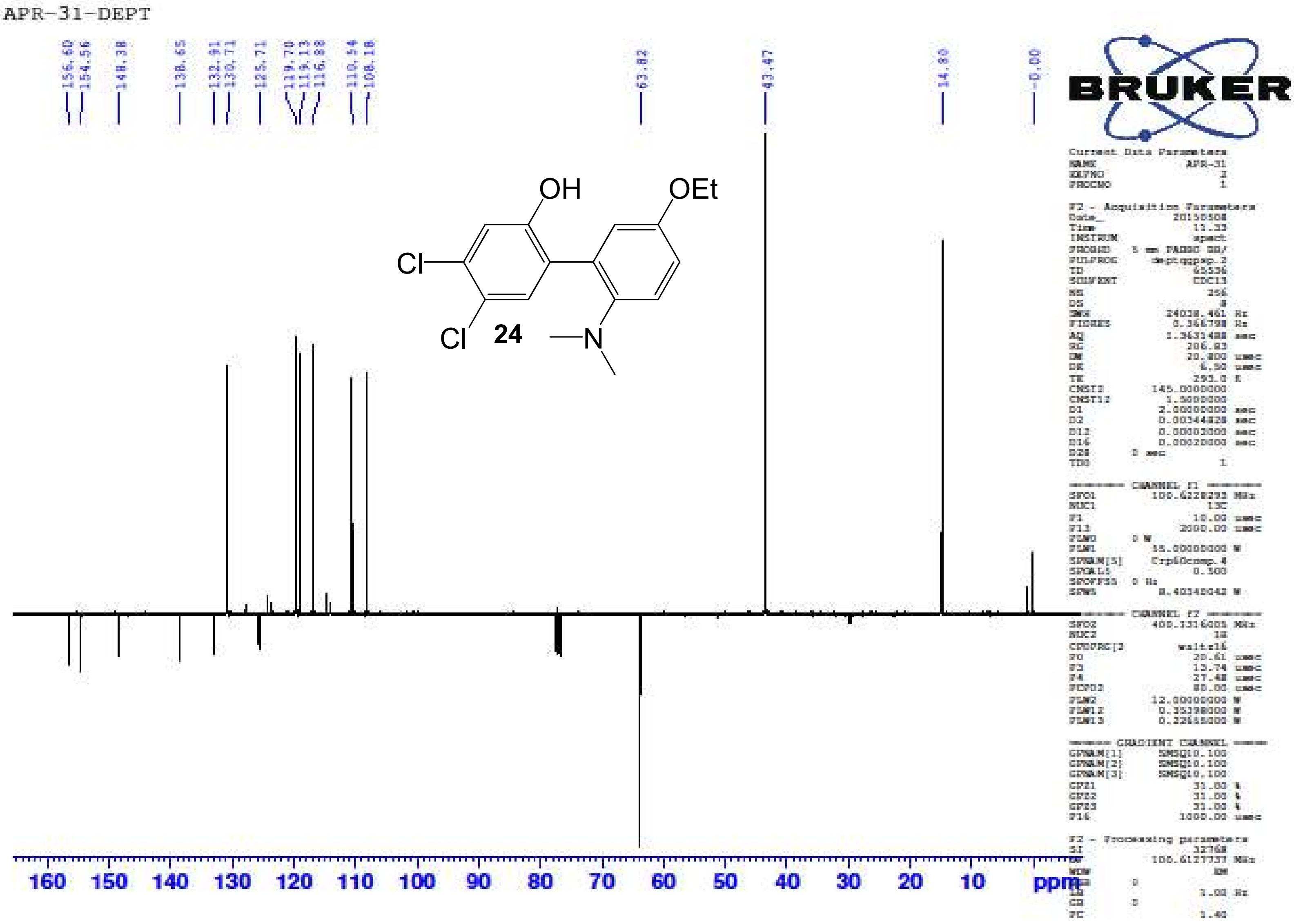

4,5-Dichloro-2'-(dimethylamino)-5'-ethoxybiphenyl-2-ol **(24)** – HR-ESIMS ($[M]^+$ Calculated: 326.0709, Observed: 326.0715)

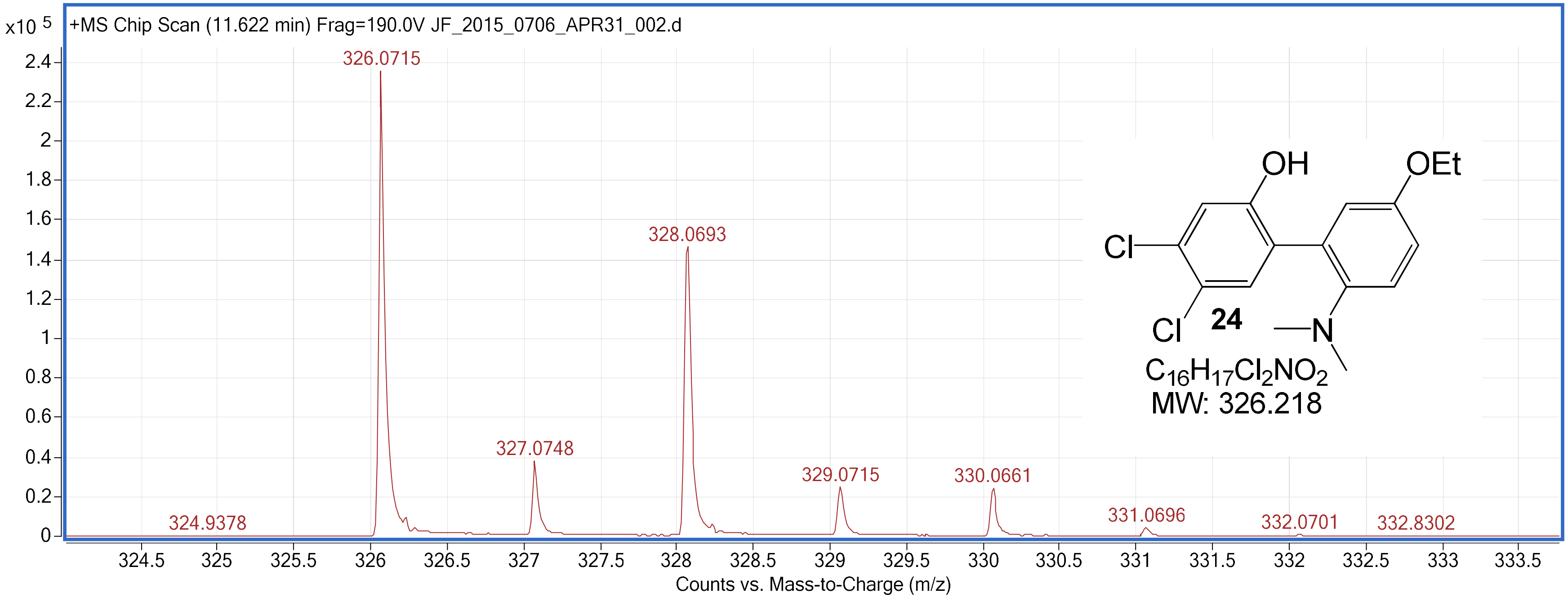

2'-(Dimethylamino)-5'-ethoxy-4,5-dimethylbiphenyl-2-ol **(25)** – $^{1}$H NMR ($CDCl_3$, 400 MHz)

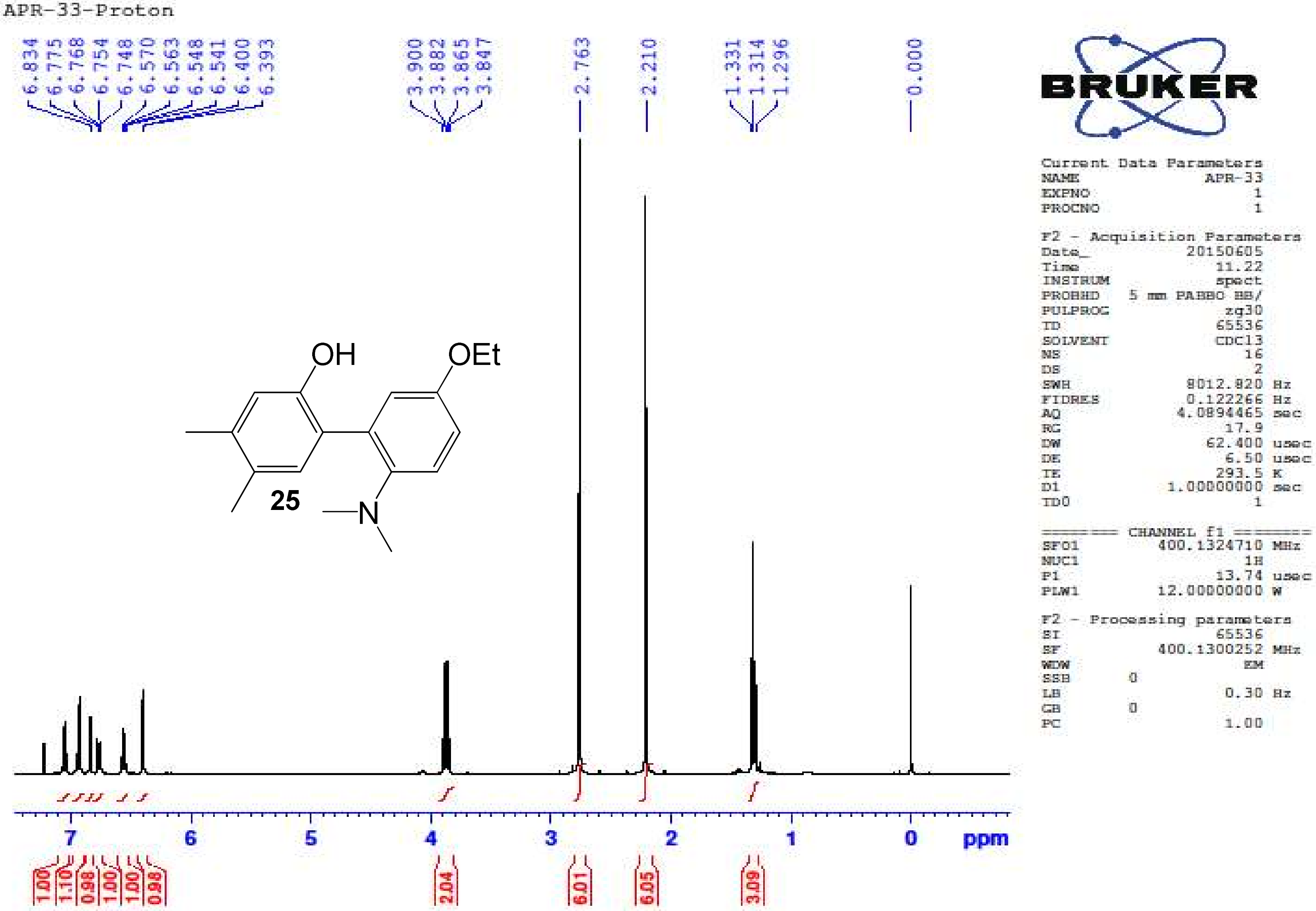

2'-(Dimethylamino)-5'-ethoxy-4,5-dimethylbiphenyl-2-ol **(25)** – $^{13}$C NMR (CDCl$_3$, 100 MHz)

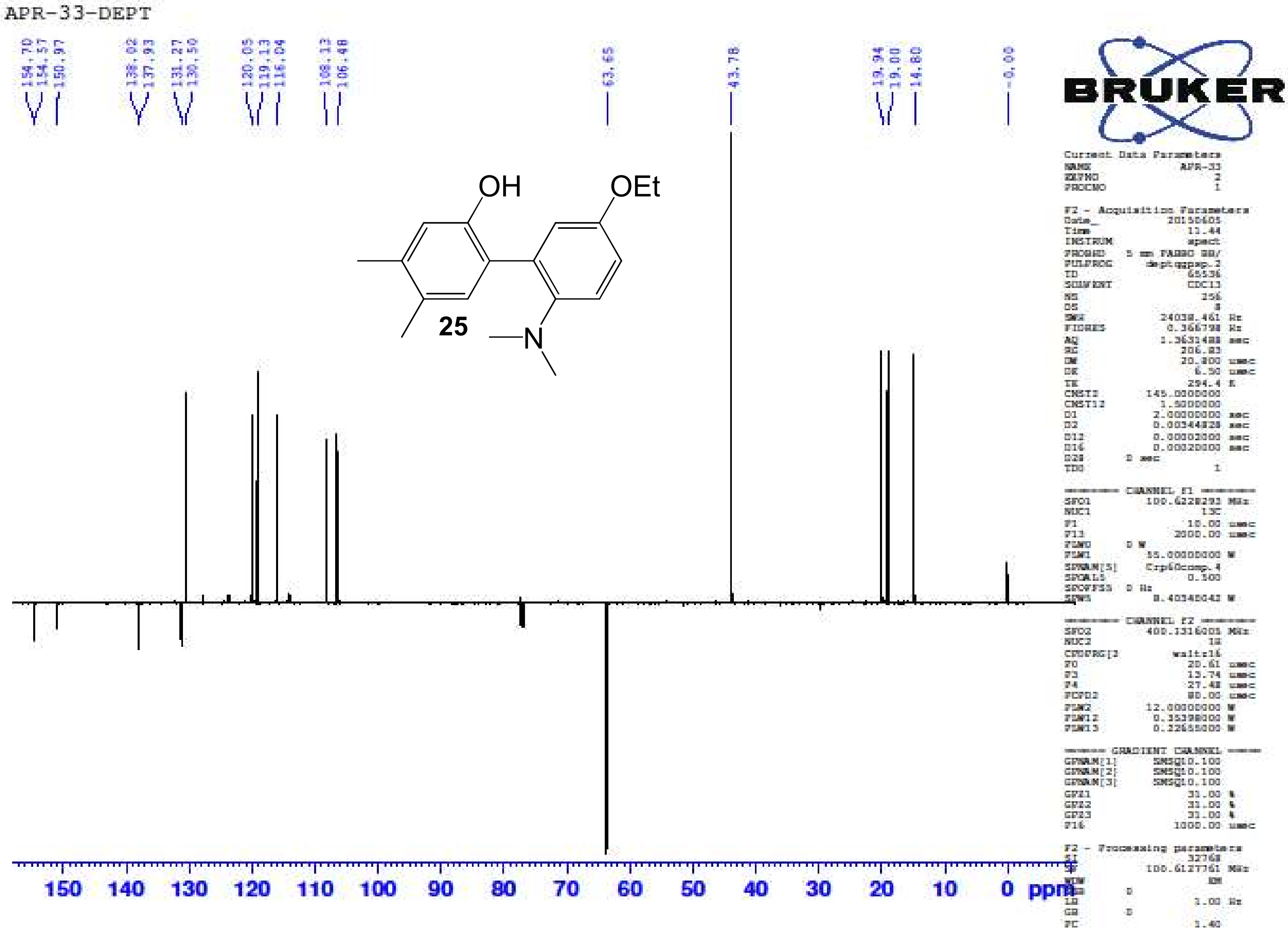

2'-(Dimethylamino)-5'-ethoxy-4,5-dimethylbiphenyl-2-ol **(25)** – HR-ESIMS ([M+H]+ Calculated: 286.1802, Observed: 286.1800)

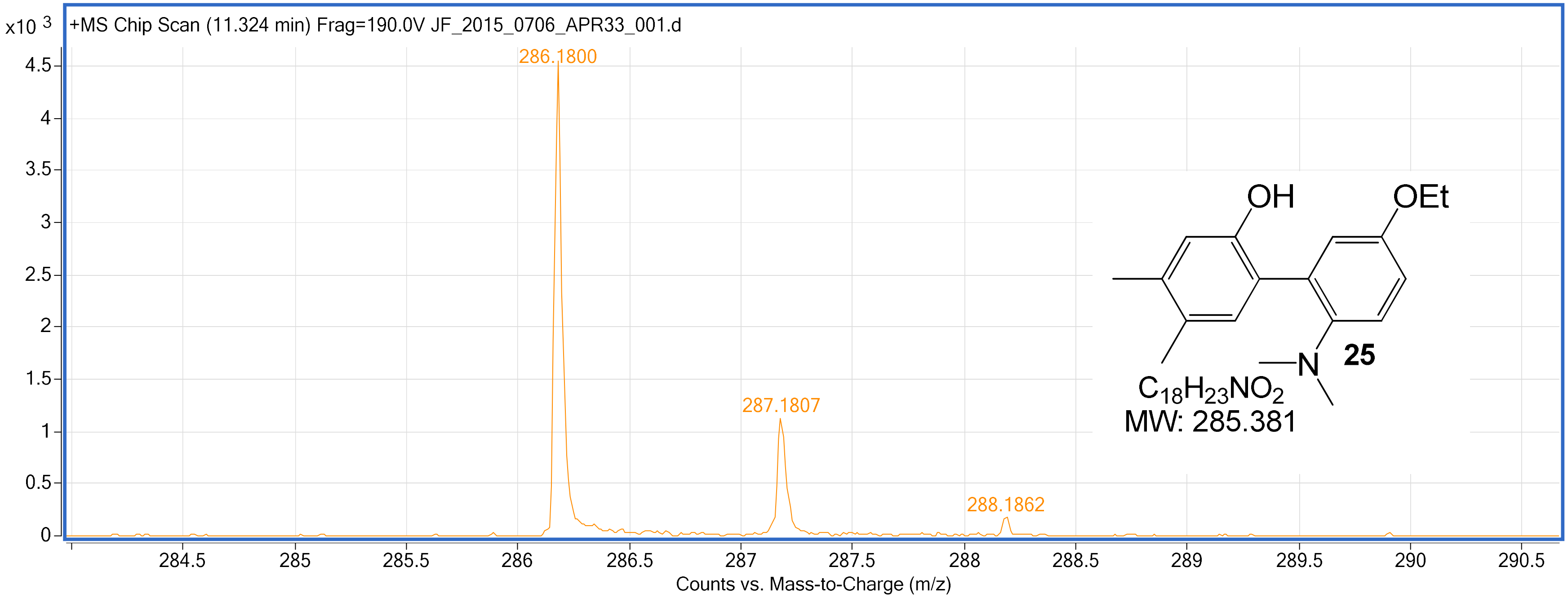

5'-Ethoxy-2'-(methyl(propyl)amino)biphenyl-2-ol **(26)** – $^{1}H$ NMR ($CDCl_3$, 400 MHz)

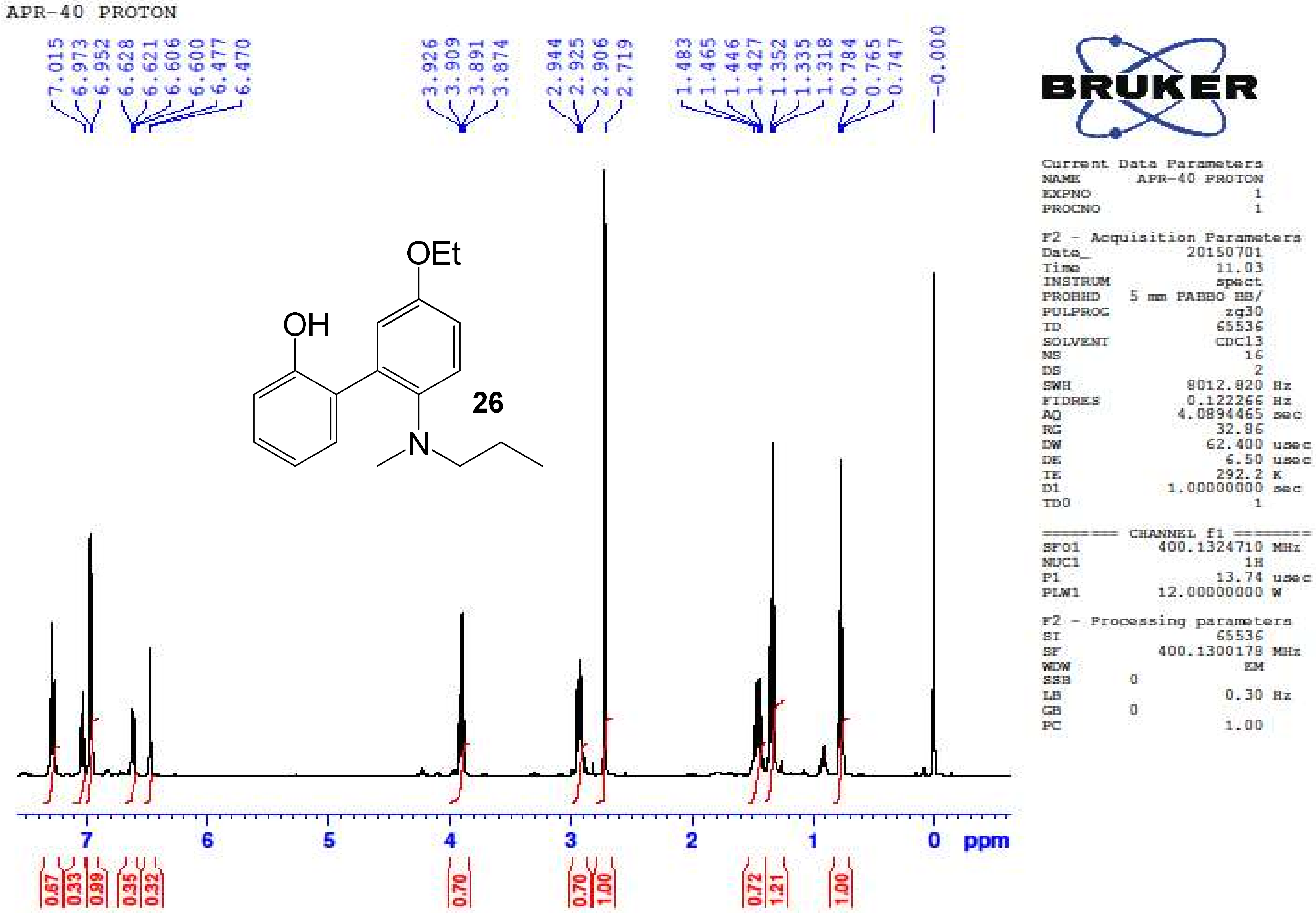

5'-Ethoxy-2'-(methyl(propyl)amino)biphenyl-2-ol **(26)** – $^{13}$C NMR ($CDCl_3$, 100 MHz)

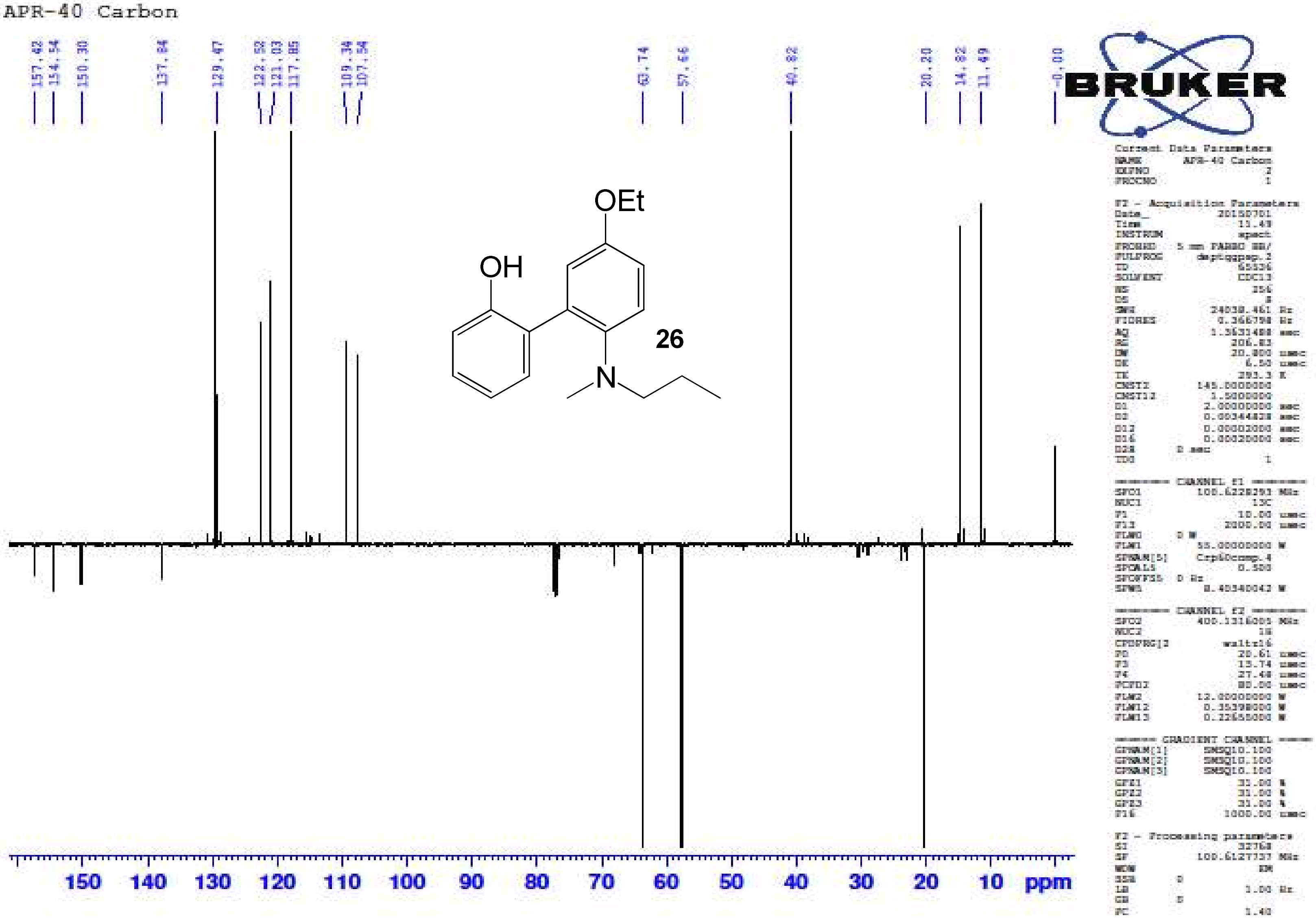

5'-Ethoxy-2'-(methyl(propyl)amino)biphenyl-2-ol **(26)** – HR-ESIMS ($[M+H]^+$ Calculated: 286.1802, observed: 286.1801)

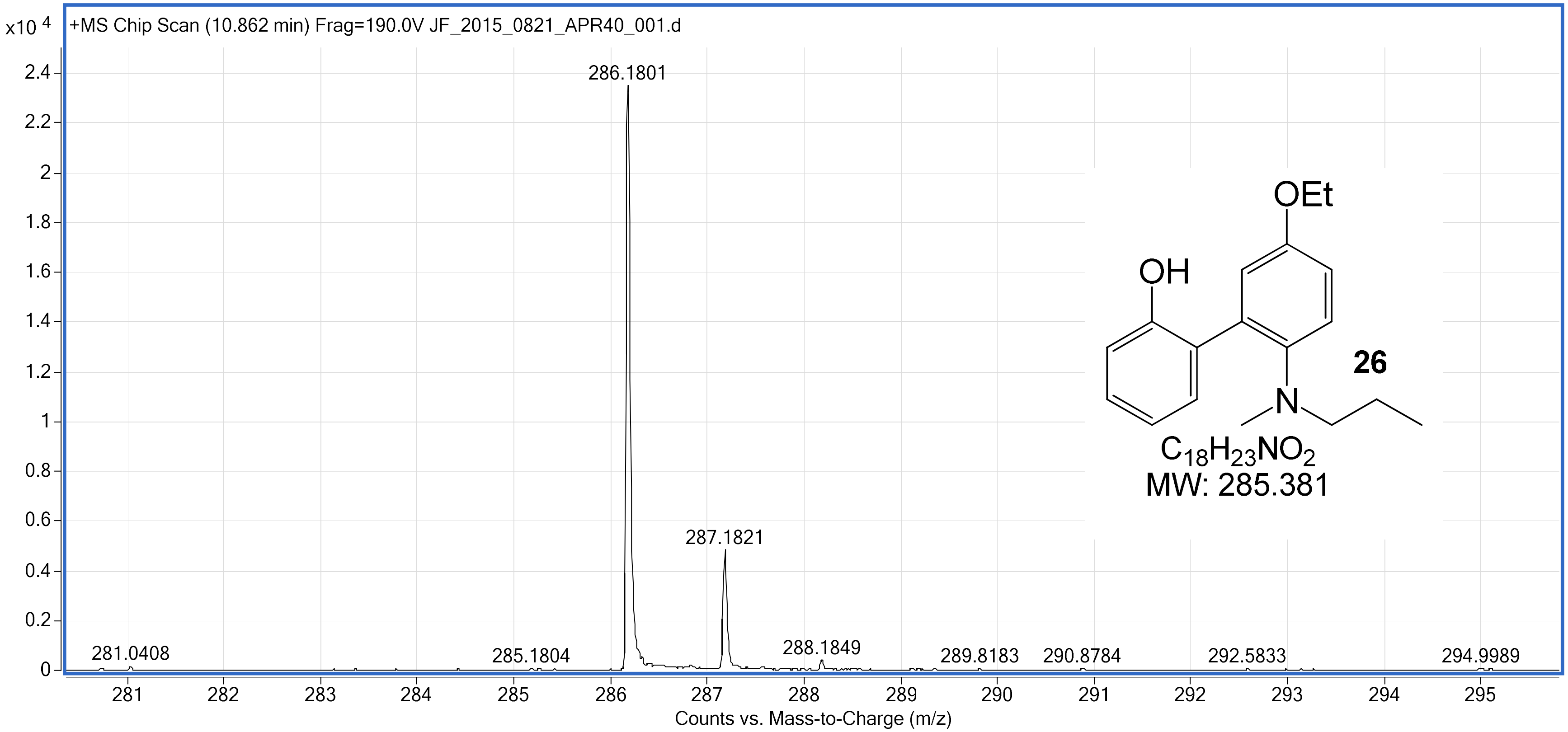

5-Bromo-5'-ethoxy-2'-(methyl(propyl)amino)biphenyl-2-ol **(27)** – $^{1}H$ NMR ($CDCl_3$, 400 MHz)

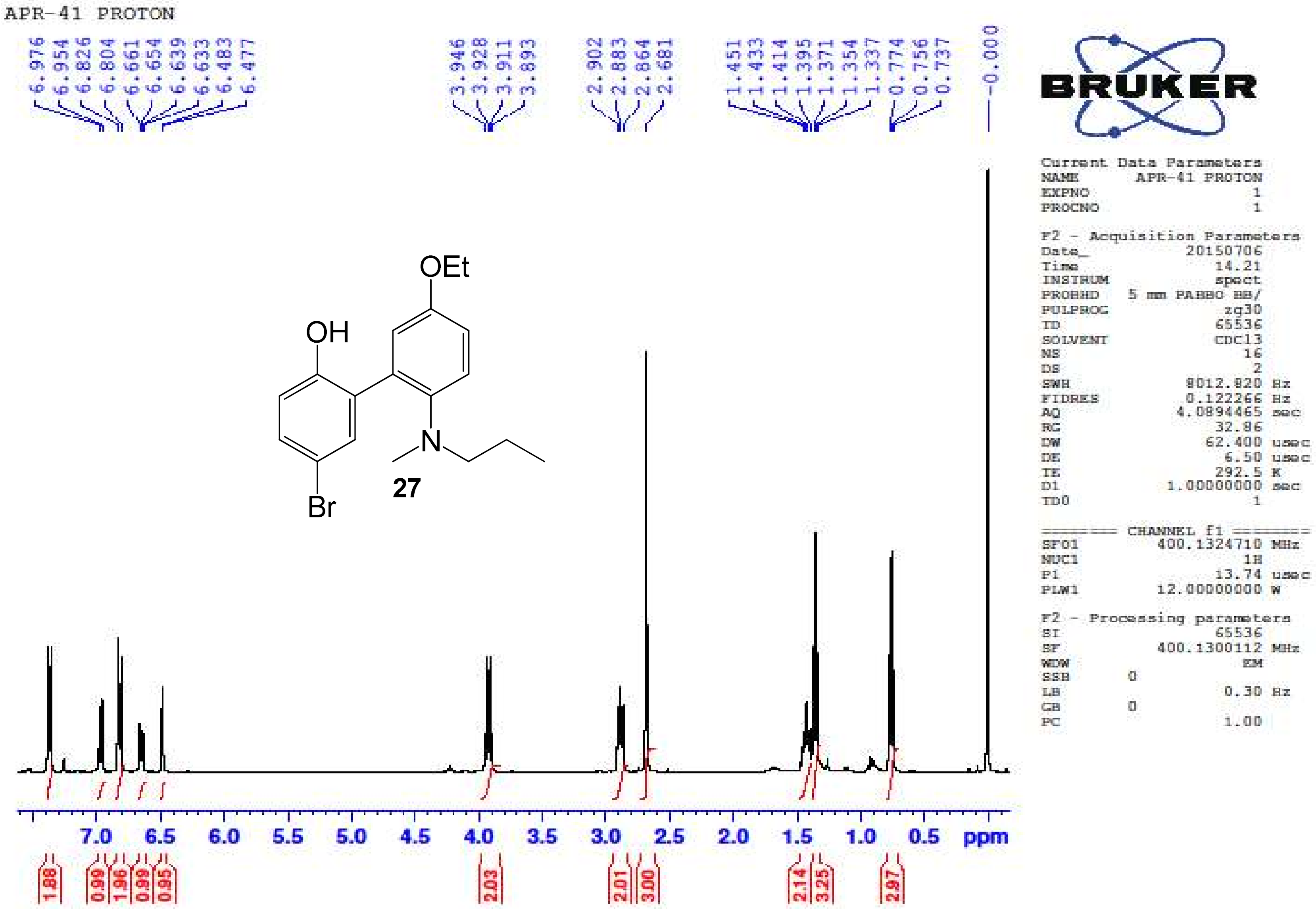

5-Bromo-5'-ethoxy-2'-(methyl(propyl)amino)biphenyl-2-ol **(27)** – $^{13}$C NMR ($CDCl_3$, 100 MHz)

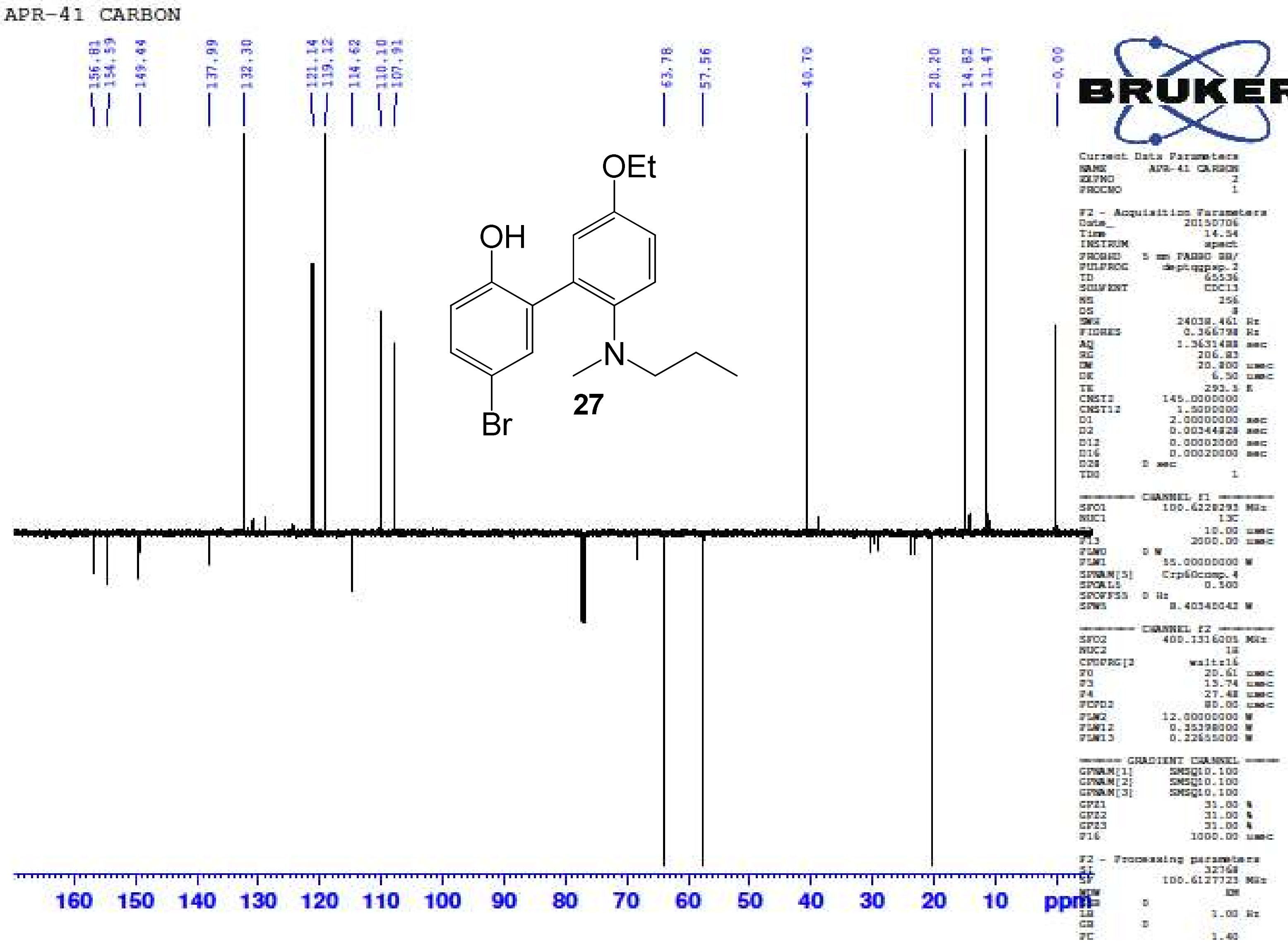

5-Bromo-5'-ethoxy-2'-(methyl(propyl)amino)biphenyl-2-ol **(27)** – HR-ESIMS ($[M]^+$ Calculated: 364.0907, Observed 364.0927)

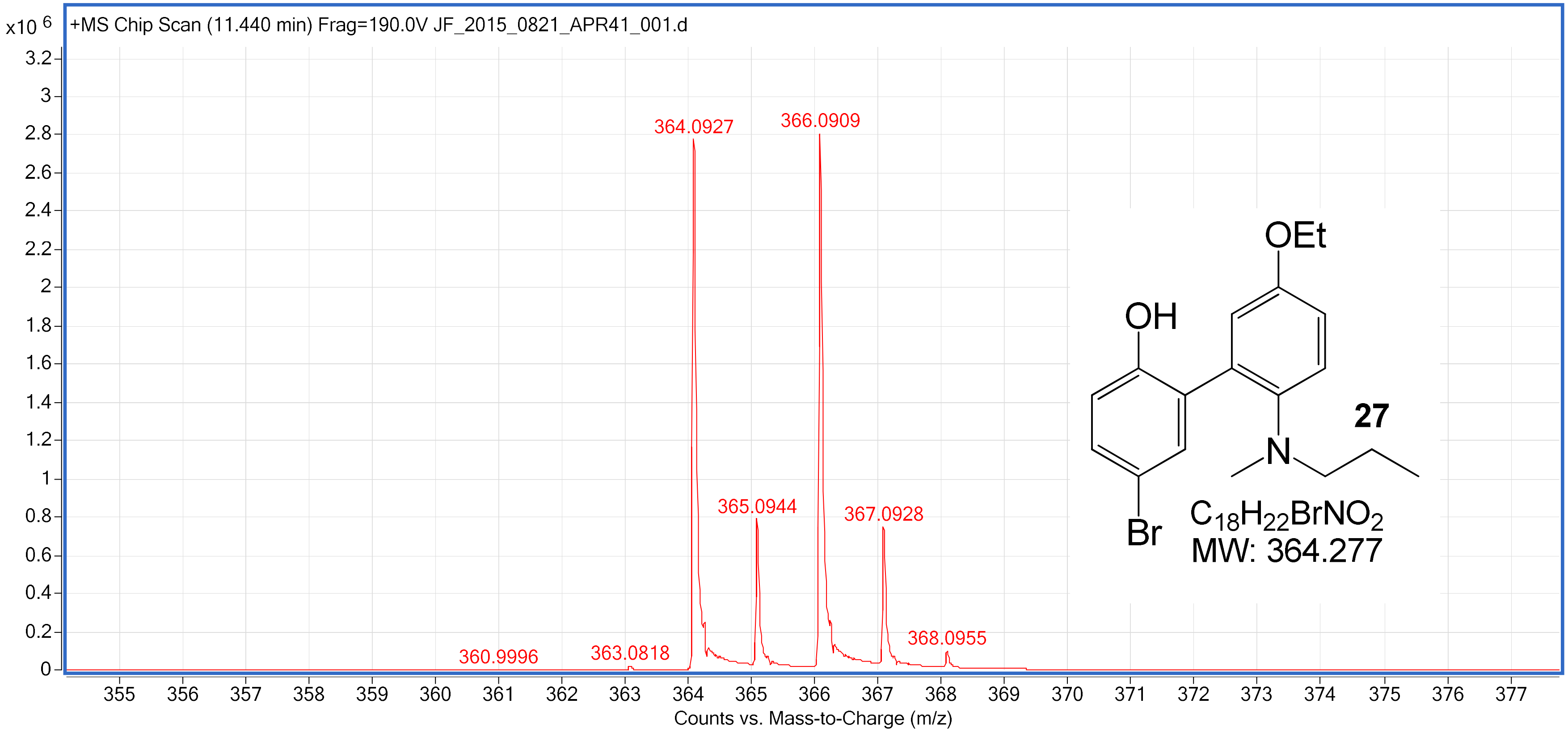

5'-Ethoxy-2'-(isobutyl(methyl)amino)biphenyl-2-ol **(28)** – $^1$H NMR ($CDCl_3$, 400 MHz)

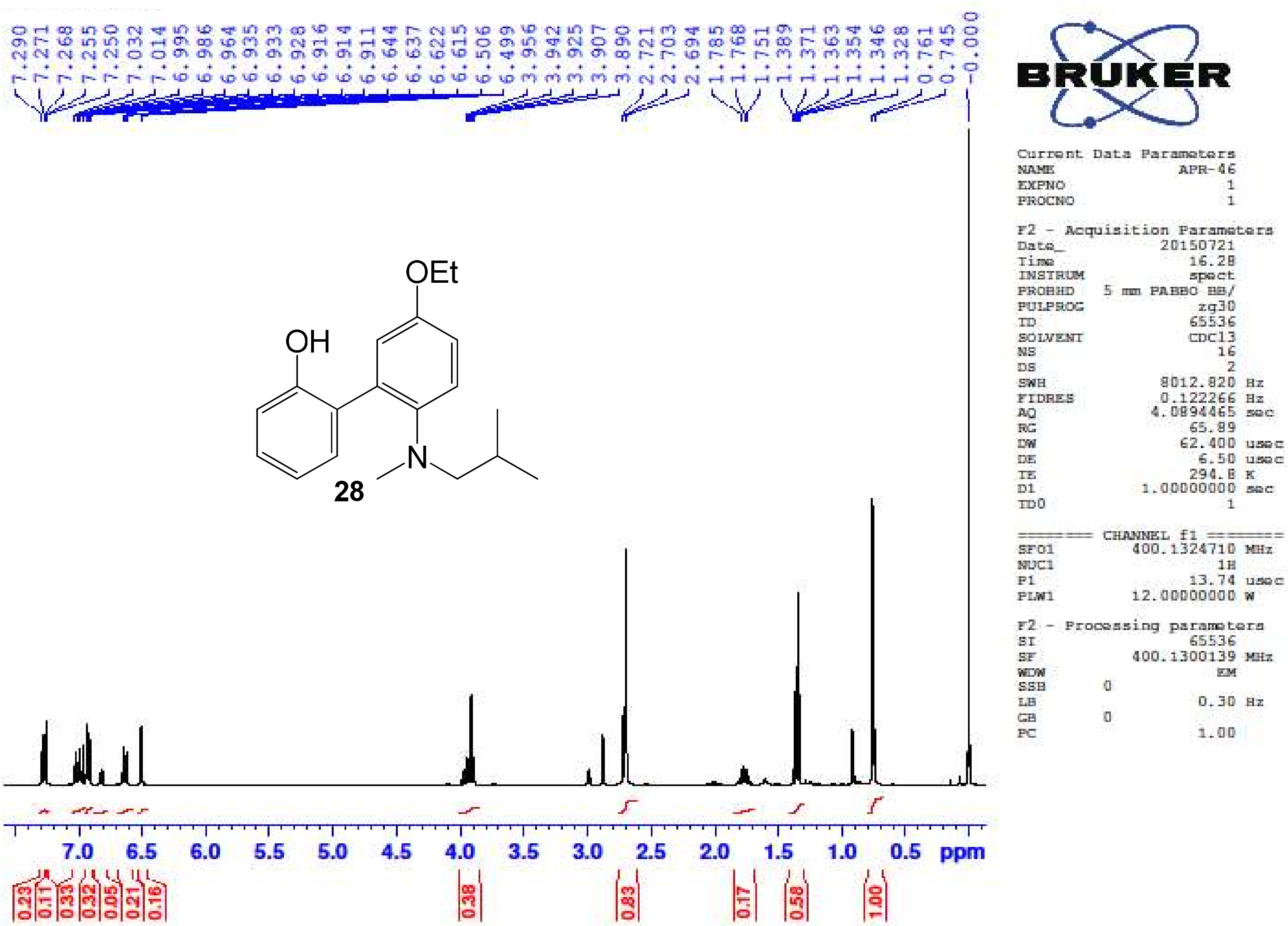

5'-Ethoxy-2'-(isobutyl(methyl)amino)biphenyl-2-ol **(28)** – $^{13}$C NMR ($CDCl_3$, 100 MHz)

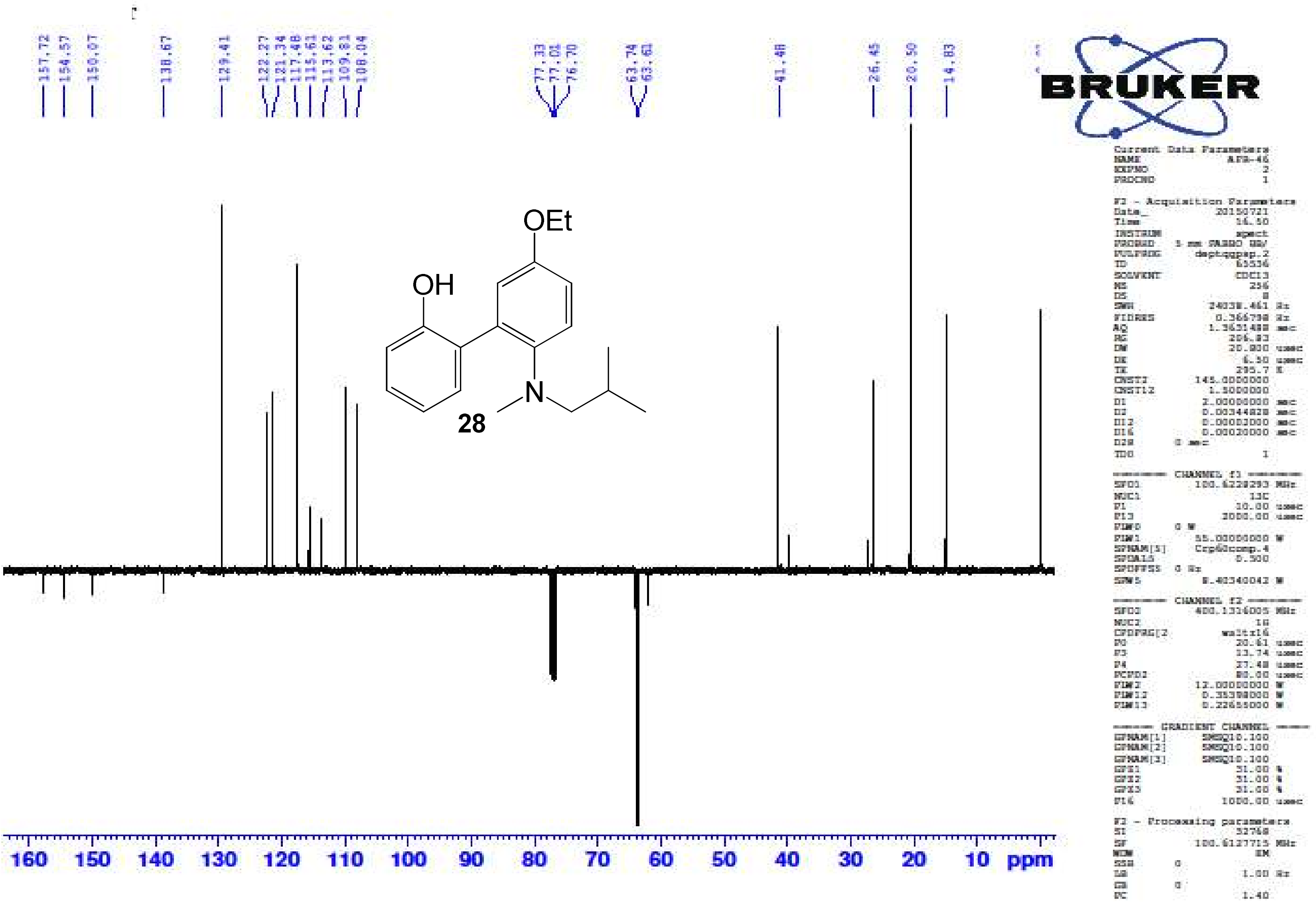

5-Bromo-5'-ethoxy-2'-(isobutyl(methyl)amino)biphenyl-2-ol **(29)** – $^{1}H$ NMR ($CDCl_3$, 400 MHz)

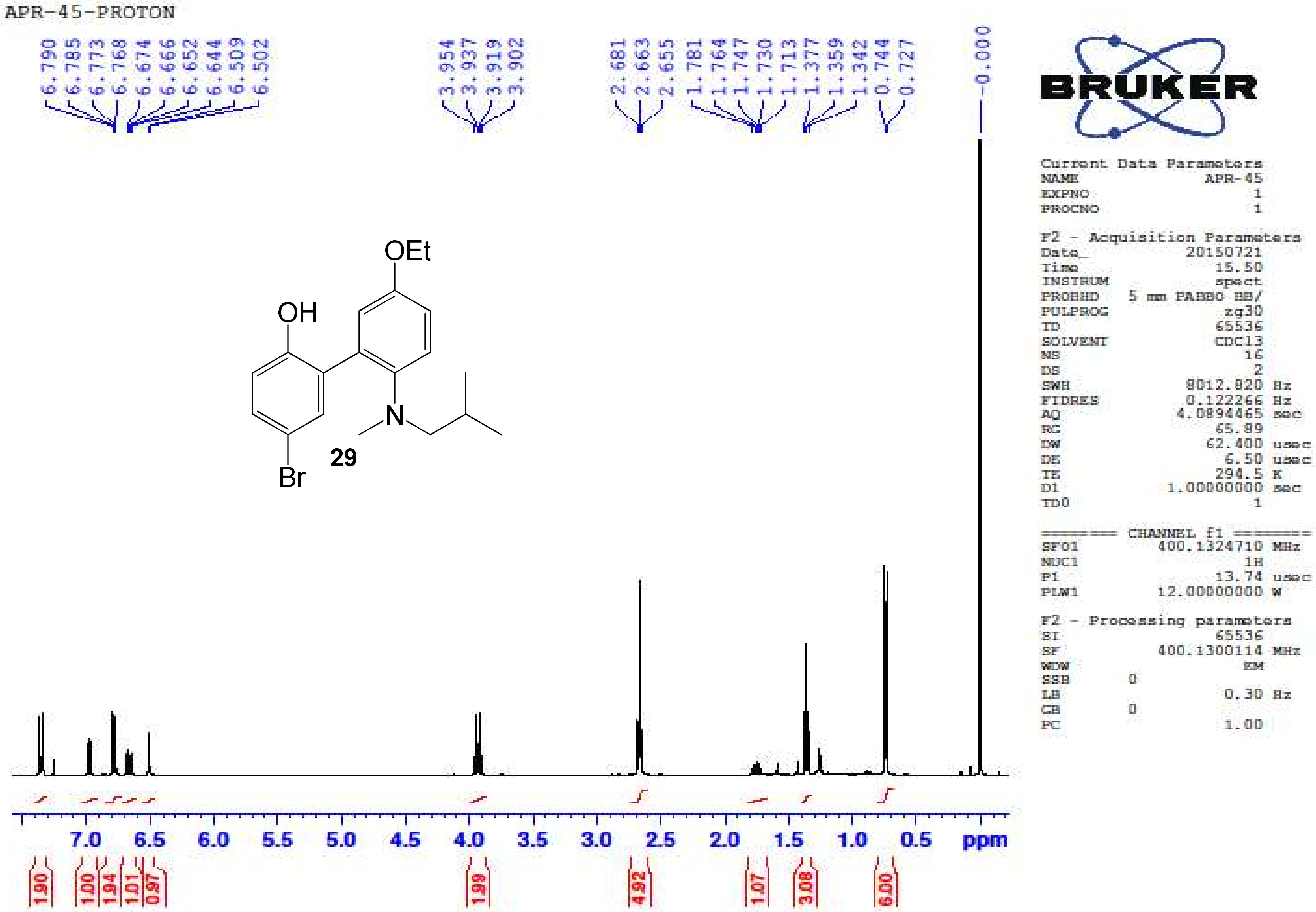

5-Bromo-5'-ethoxy-2'-(isobutyl(methyl)amino)biphenyl-2-ol **(29)** – $^{13}$C NMR (CDCl$_3$, 100 MHz)

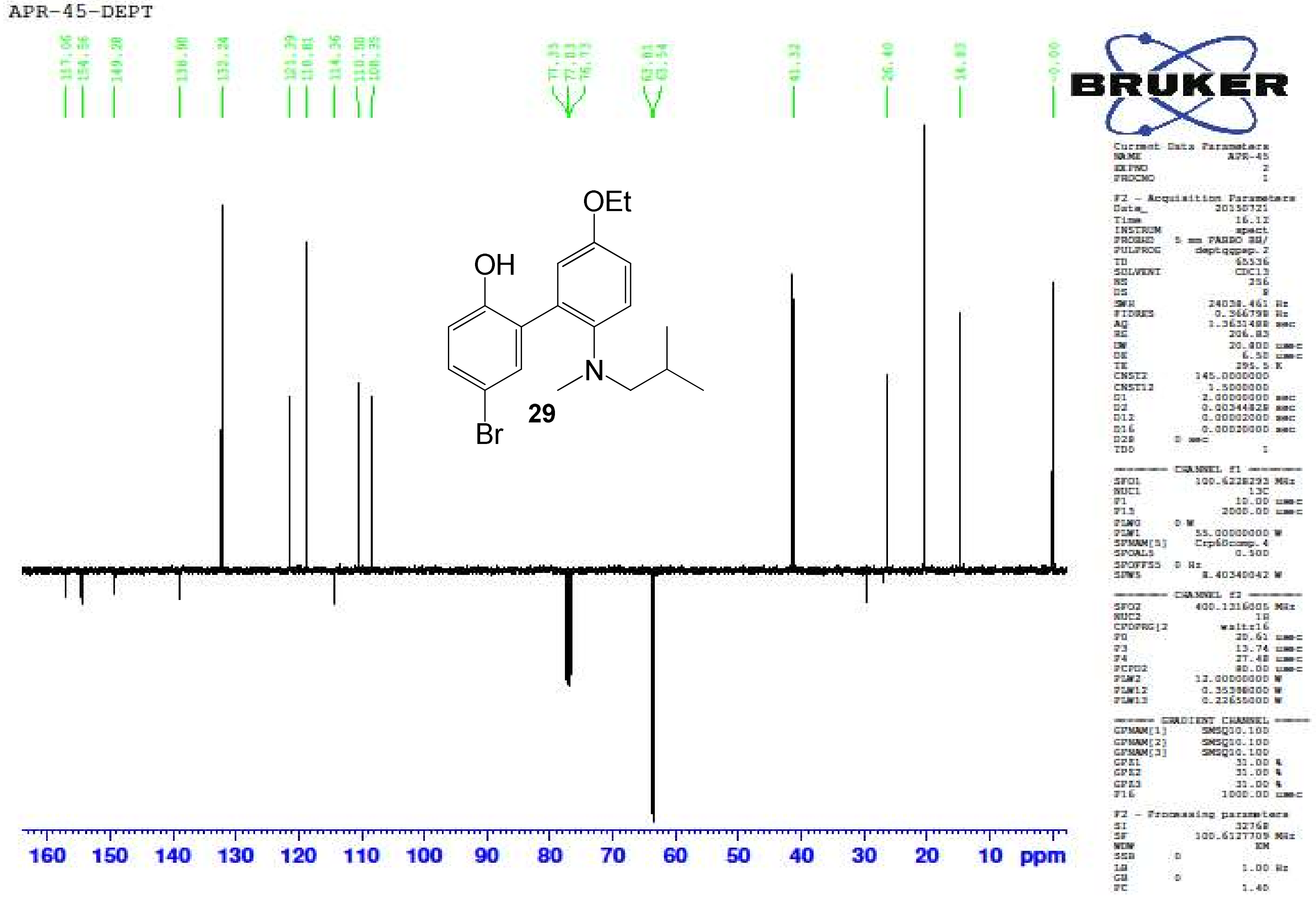

5-Bromo-5'-ethoxy-2'-(isobutyl(methyl)amino)biphenyl-2-ol **(29)** – HR-ESIMS ($[M]^+$ Calculated: 378.1063, Observed: 378.1087)

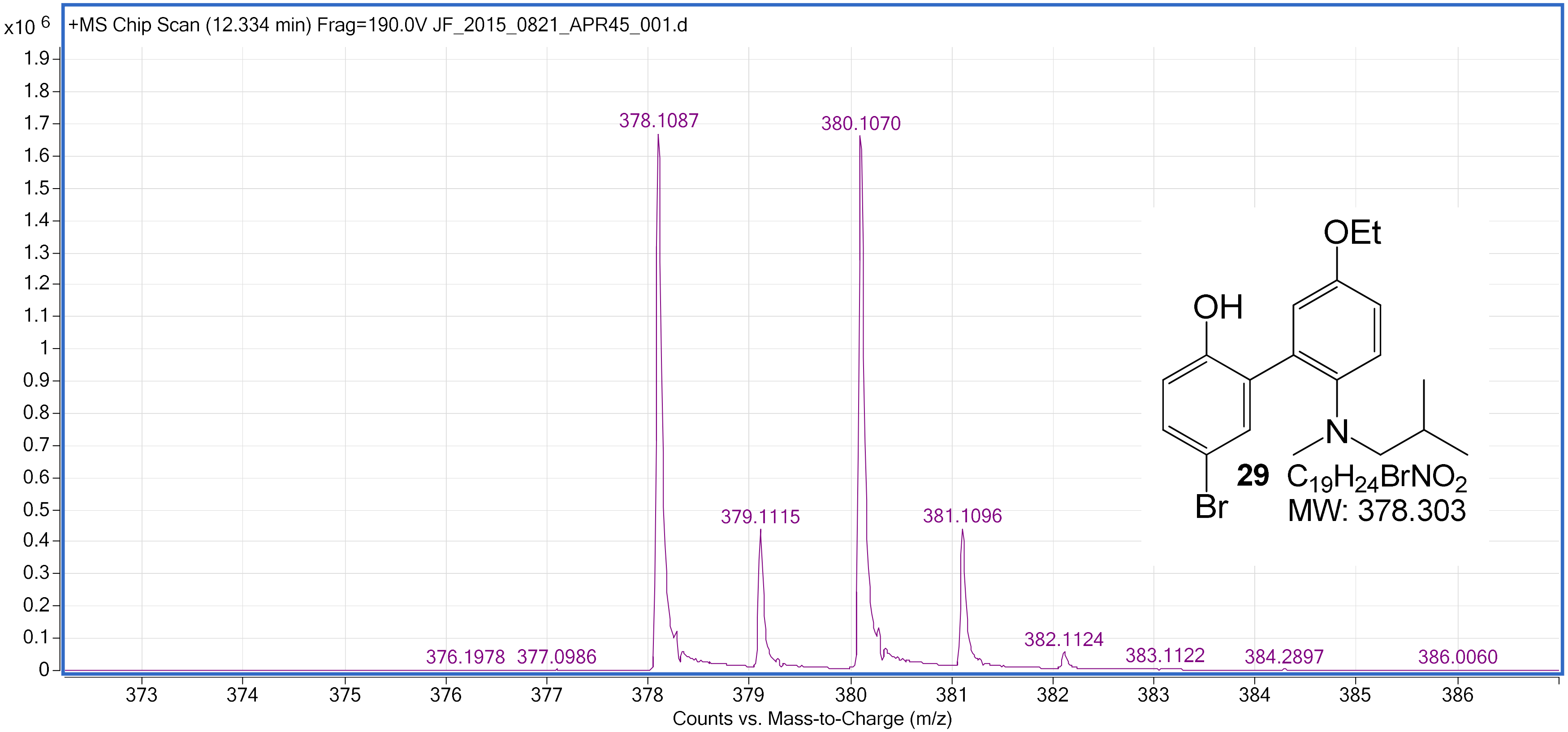

5-Bromo-5'-ethoxy-2'-(isopropyl(methyl)amino)biphenyl-2-ol **(30)** – $^1$H NMR ($CDCl_3$, 400 MHz)

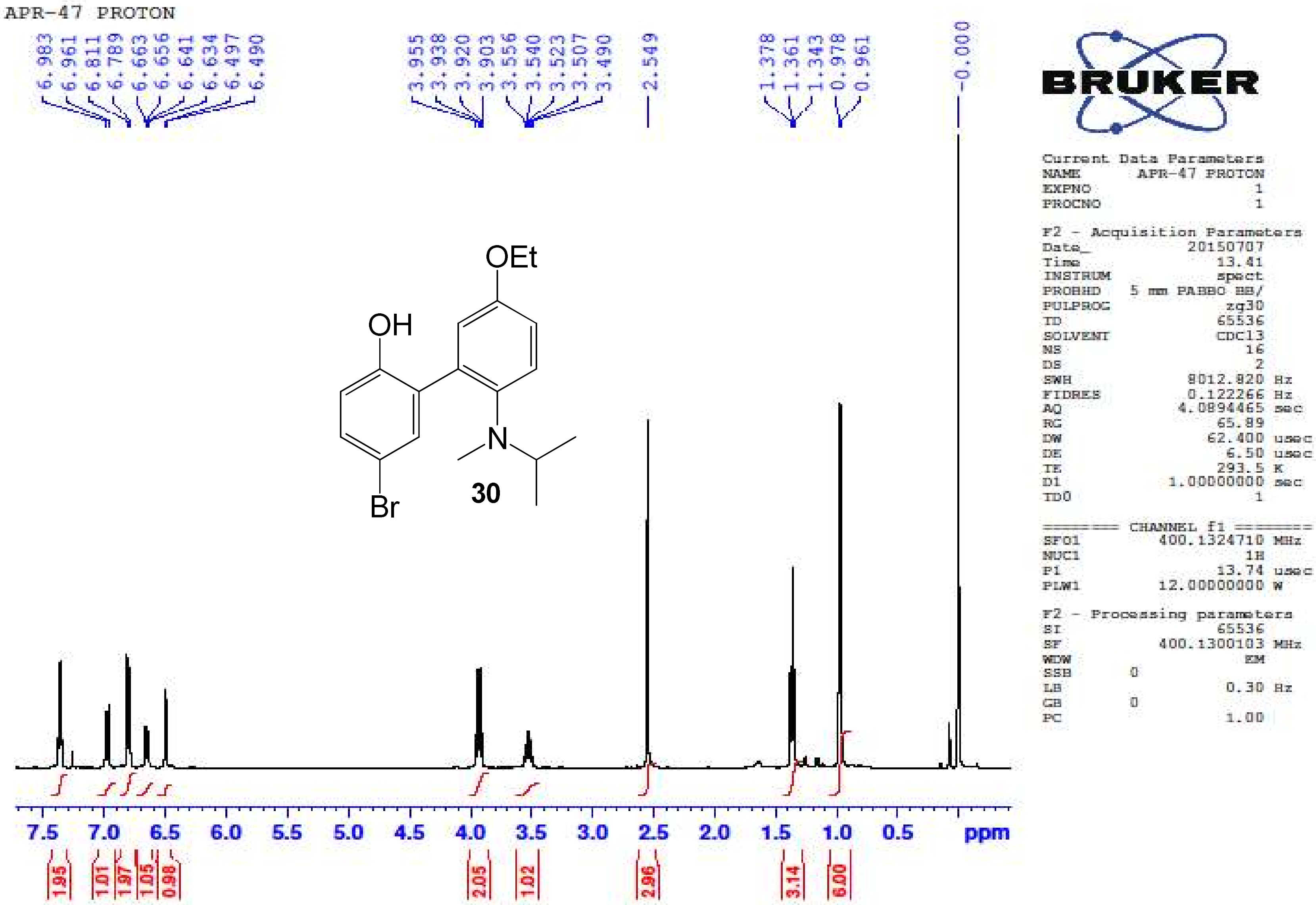

5-Bromo-5'-ethoxy-2'-(isopropyl(methyl)amino)biphenyl-2-ol **(30)** – $^{13}$C NMR ($CDCl_3$, 100 MHz)

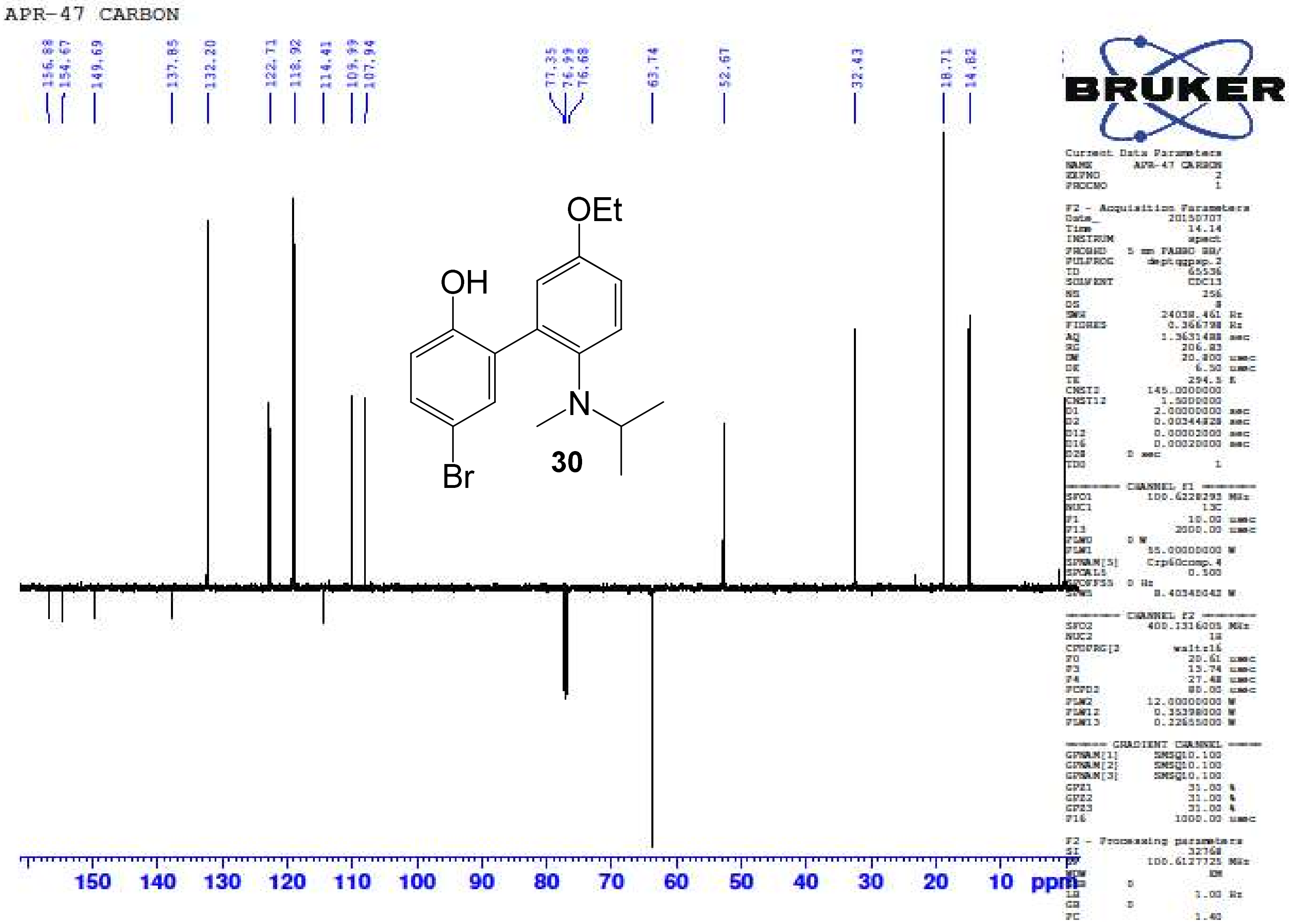

5-Bromo-5'-ethoxy-2'-(isopropyl(methyl)amino)biphenyl-2-ol **(30)** – HR-ESIMS ($[M]^+$ Calculated: 364.0907, Observed: 364.0909)

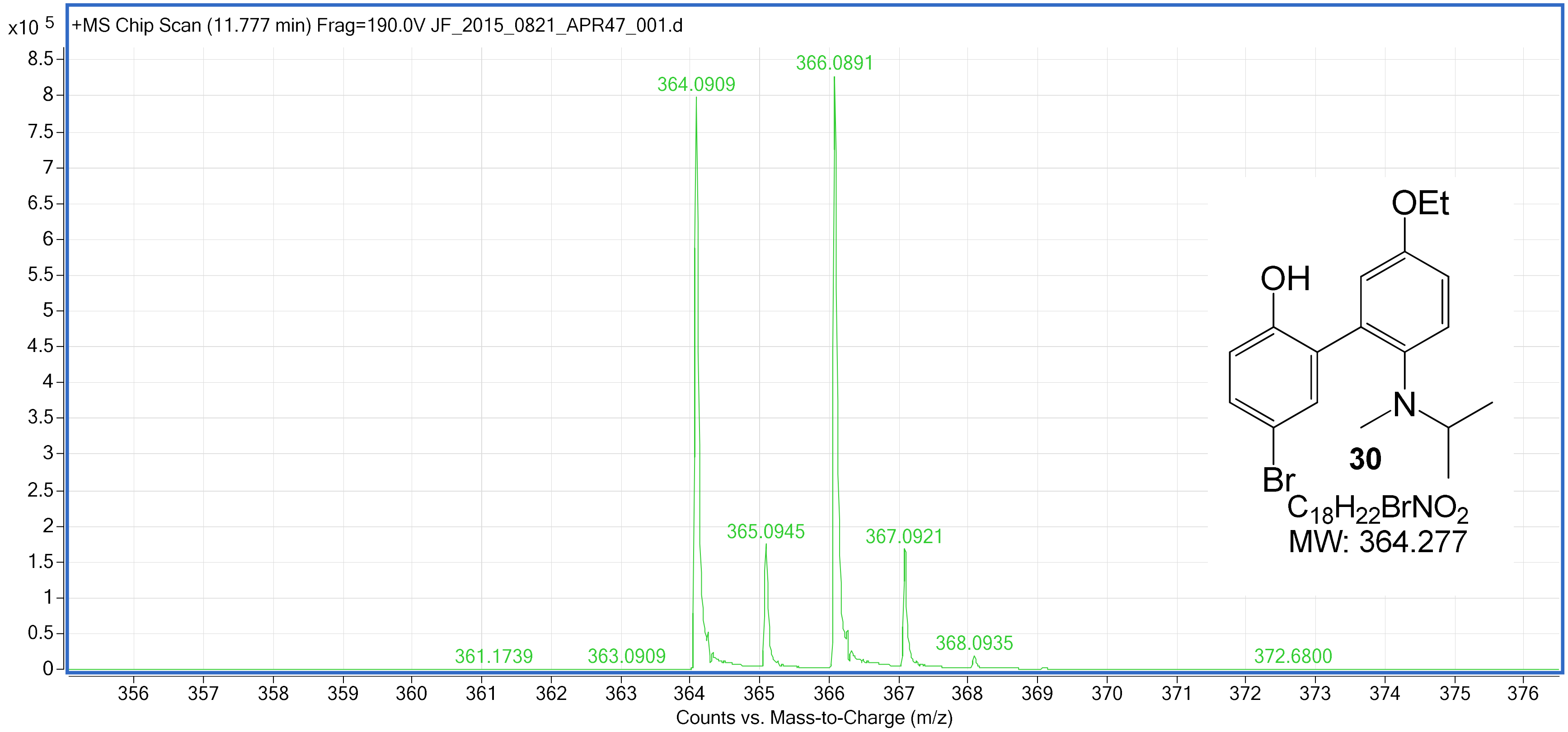

2'-(Benzyl(methyl)amino)-5'-ethoxybiphenyl-2-ol **(31)** – $^{1}$H NMR ($CDCl_3$, 400 MHz)

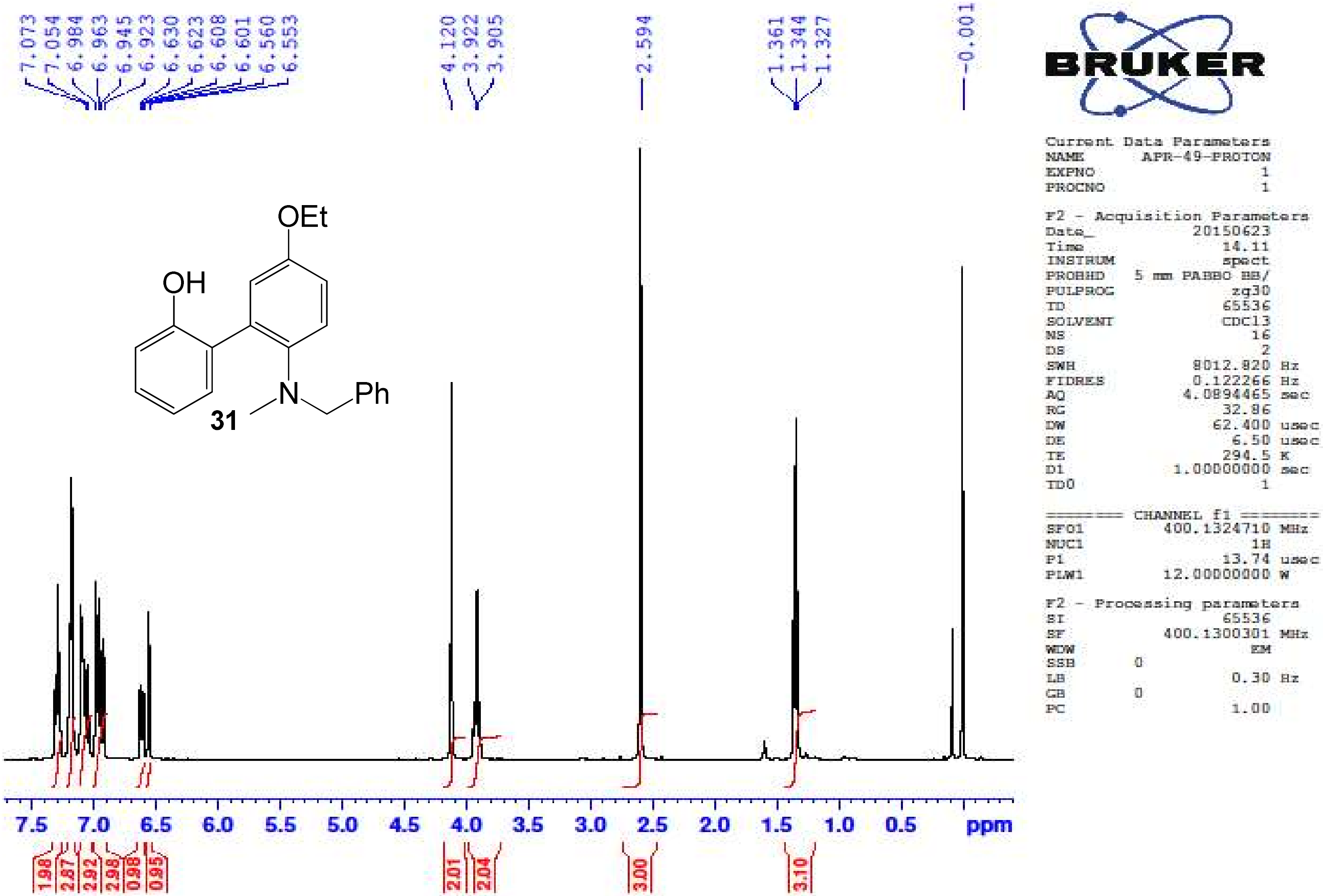

2'-(Benzyl(methyl)amino)-5'-ethoxybiphenyl-2-ol **(31)** – $^{13}$C NMR ($CDCl_3$, 100 MHz)

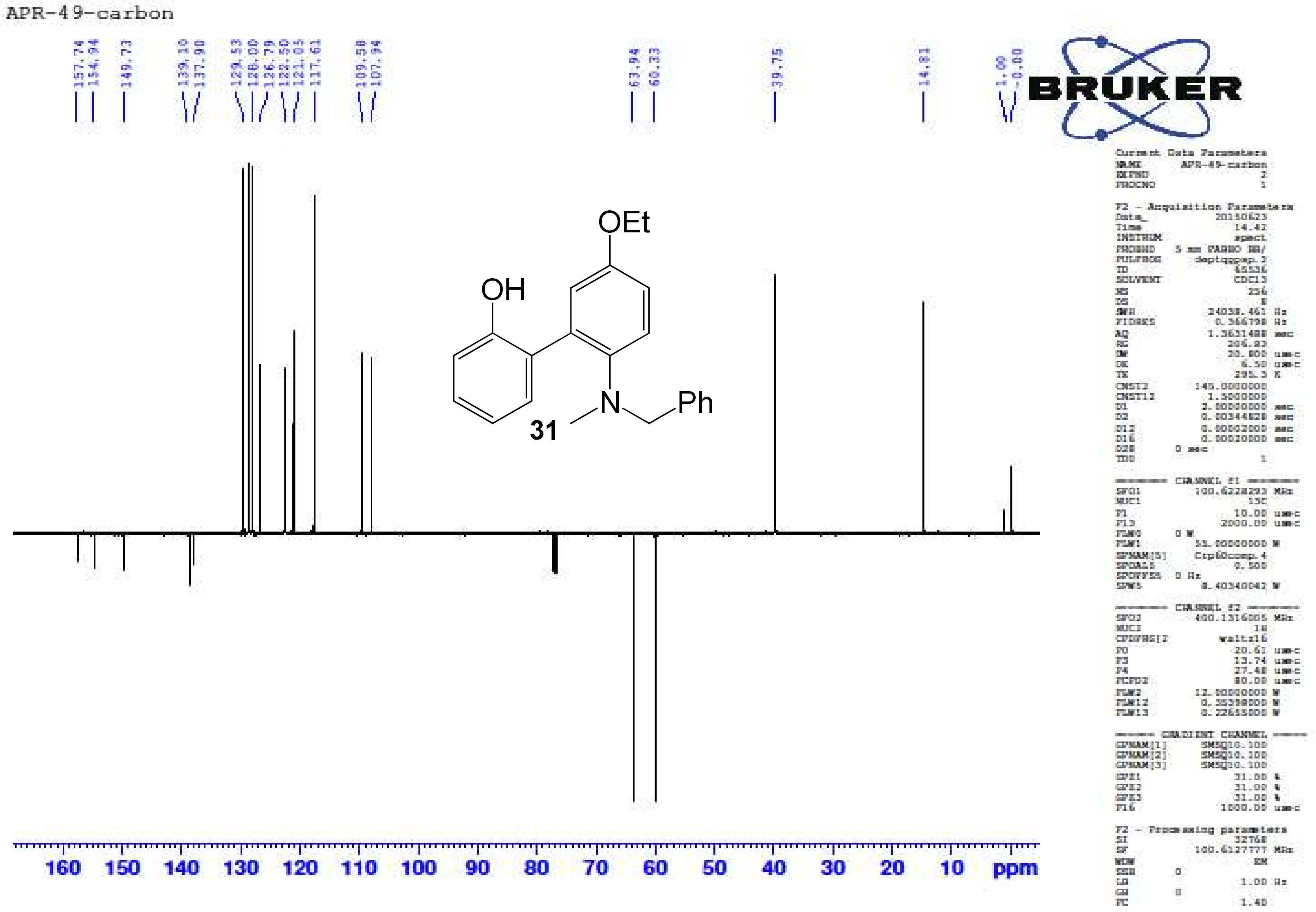

2'-(Benzyl(methyl)amino)-5'-ethoxybiphenyl-2-ol **(31)** – HR-ESIMS ($[M+H]^+$ Calculated: 334.1802, Observed: 334.1814)

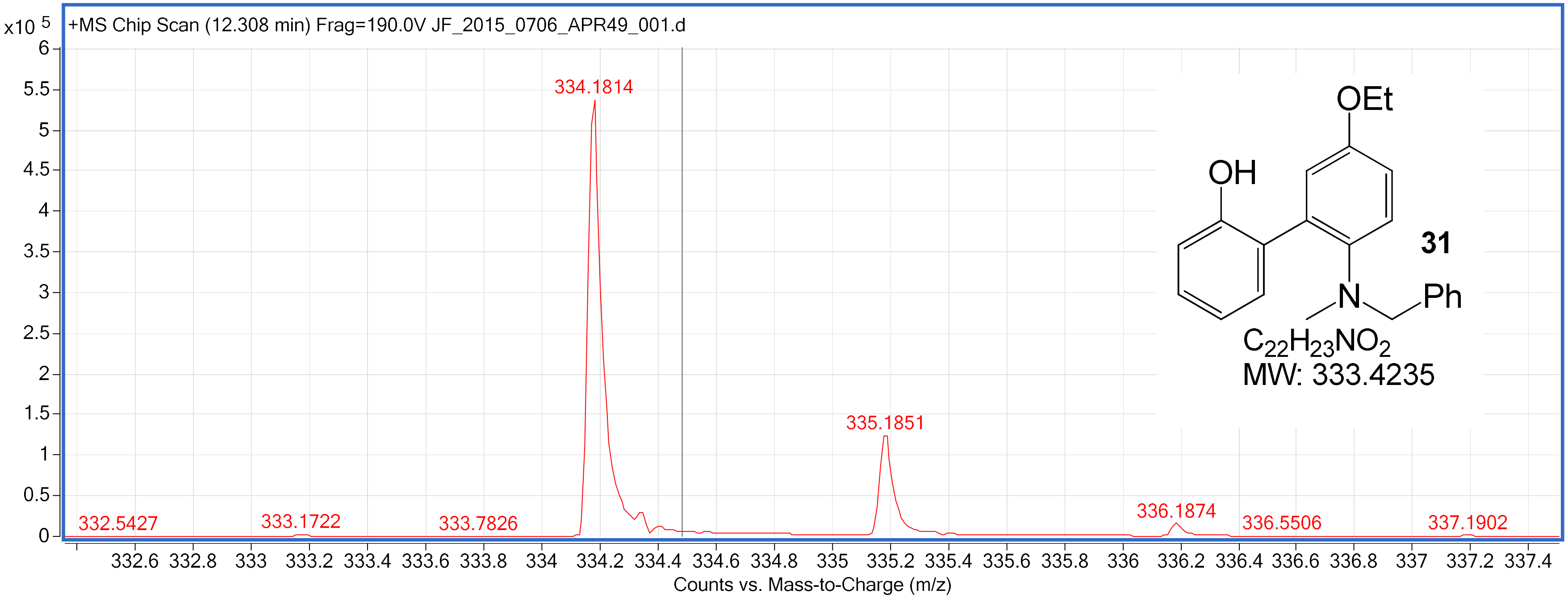

2'-(Benzyl(methyl)amino)-5-bromo-5'-ethoxybiphenyl-2-ol **(32)** – $^{1}$H NMR ($CDCl_3$, 400 MHz)

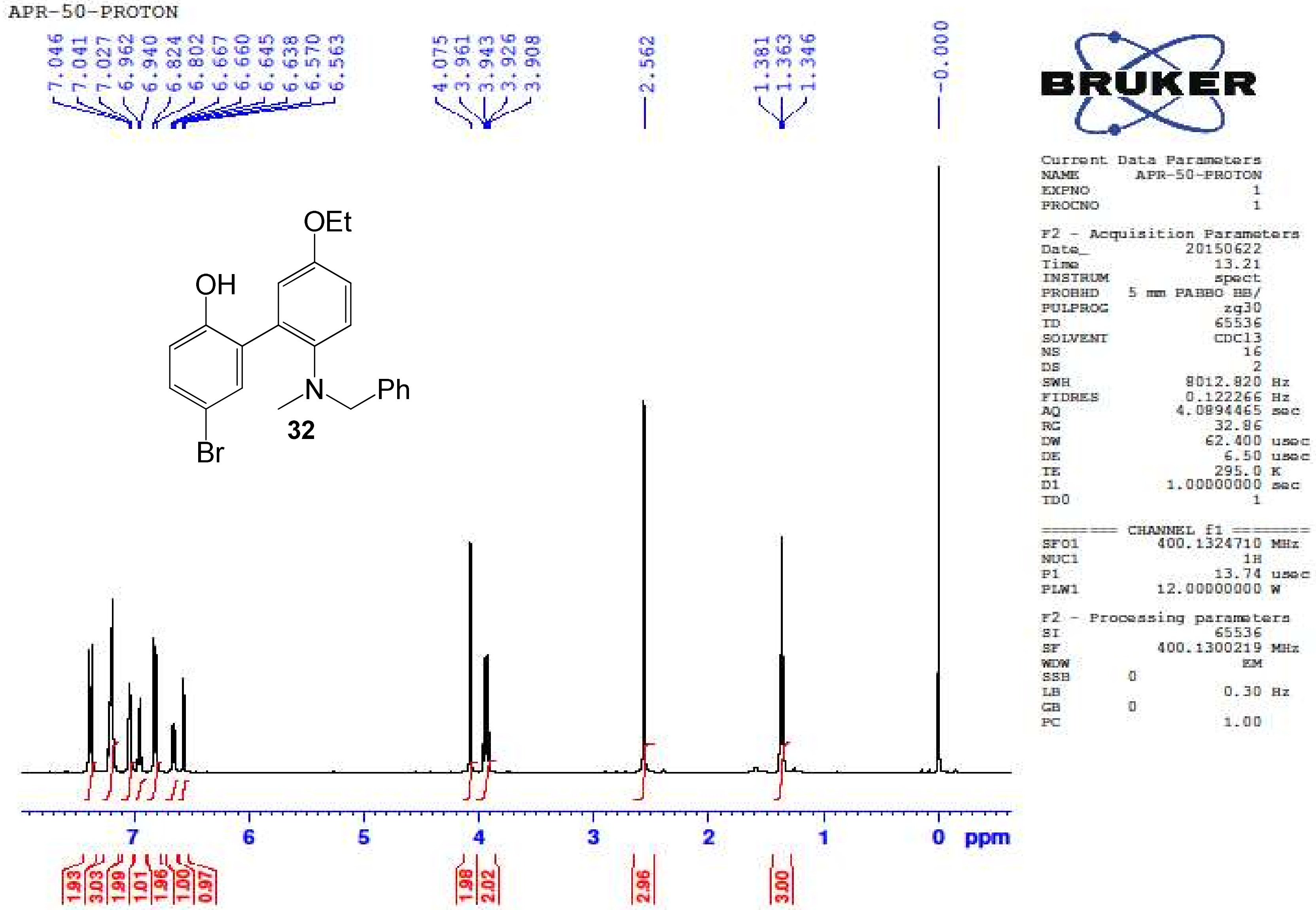

2'-(Benzyl(methyl)amino)-5-bromo-5'-ethoxybiphenyl-2-ol **(32)** – $^{13}C$ NMR ($CDCl_3$, 100 MHz)

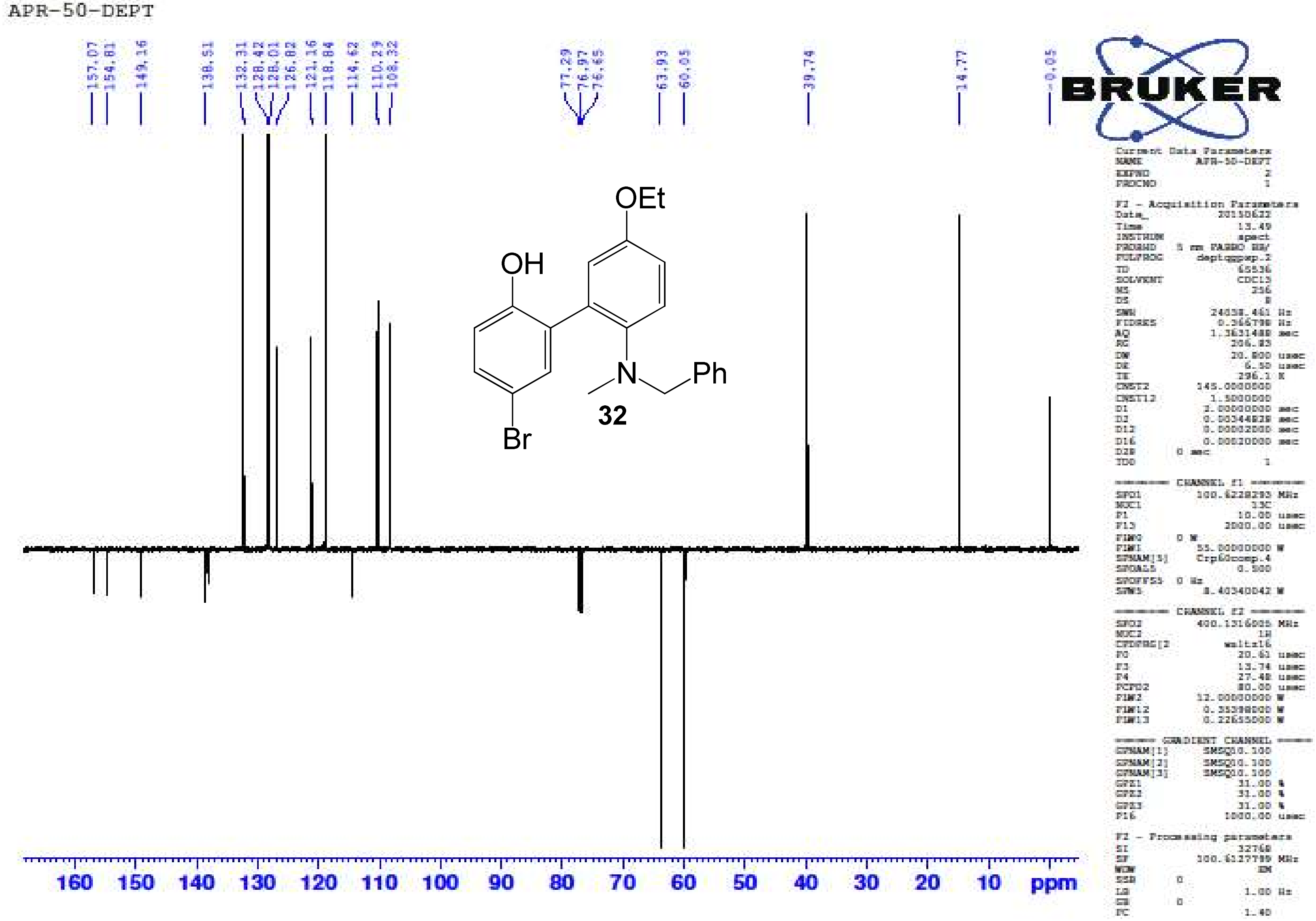

2'-(Benzyl(methyl)amino)-5-bromo-5'-ethoxybiphenyl-2-ol **(32)** – HR-ESIMS ($[M]^+$ Calculated: 412.0907, Observed: 412.0908)

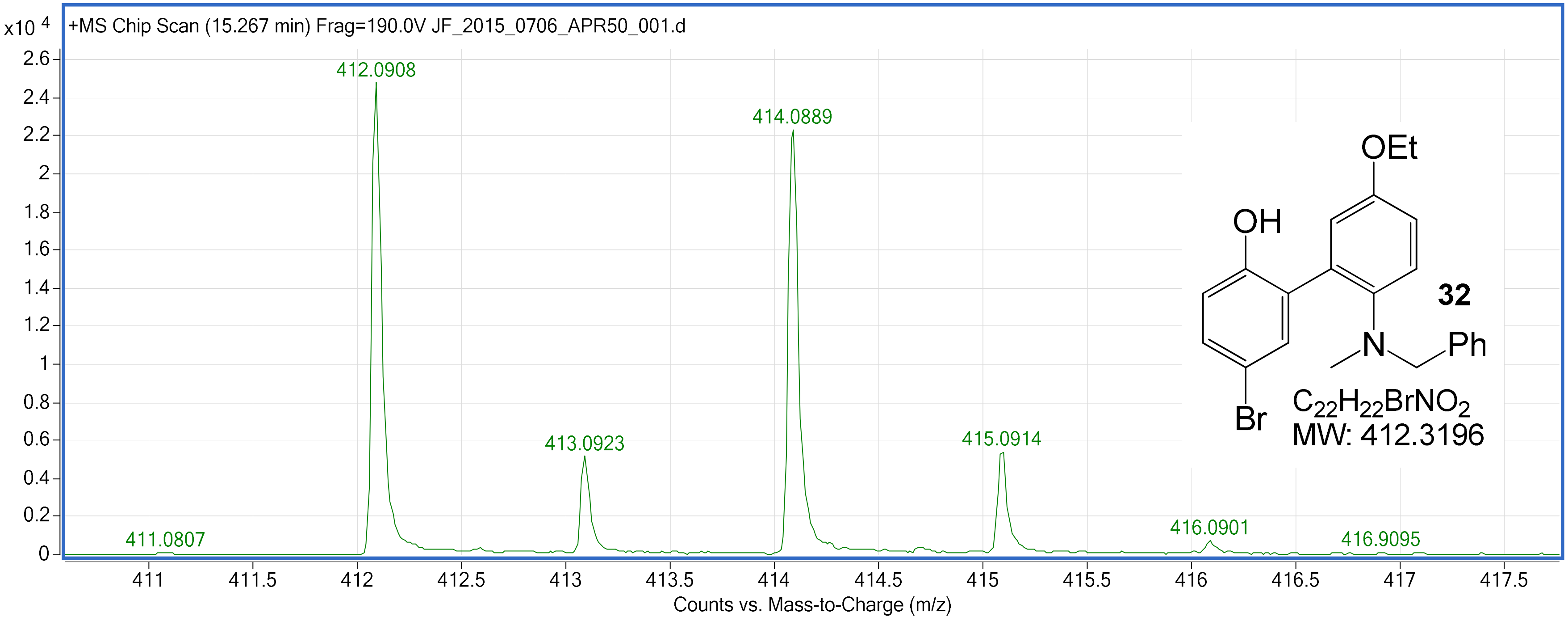

5'-Ethoxy-2'-(ethyl(propyl)amino)biphenyl-2-ol **(33)** – $^{1}$H NMR ($CDCl_3$, 400 MHz)

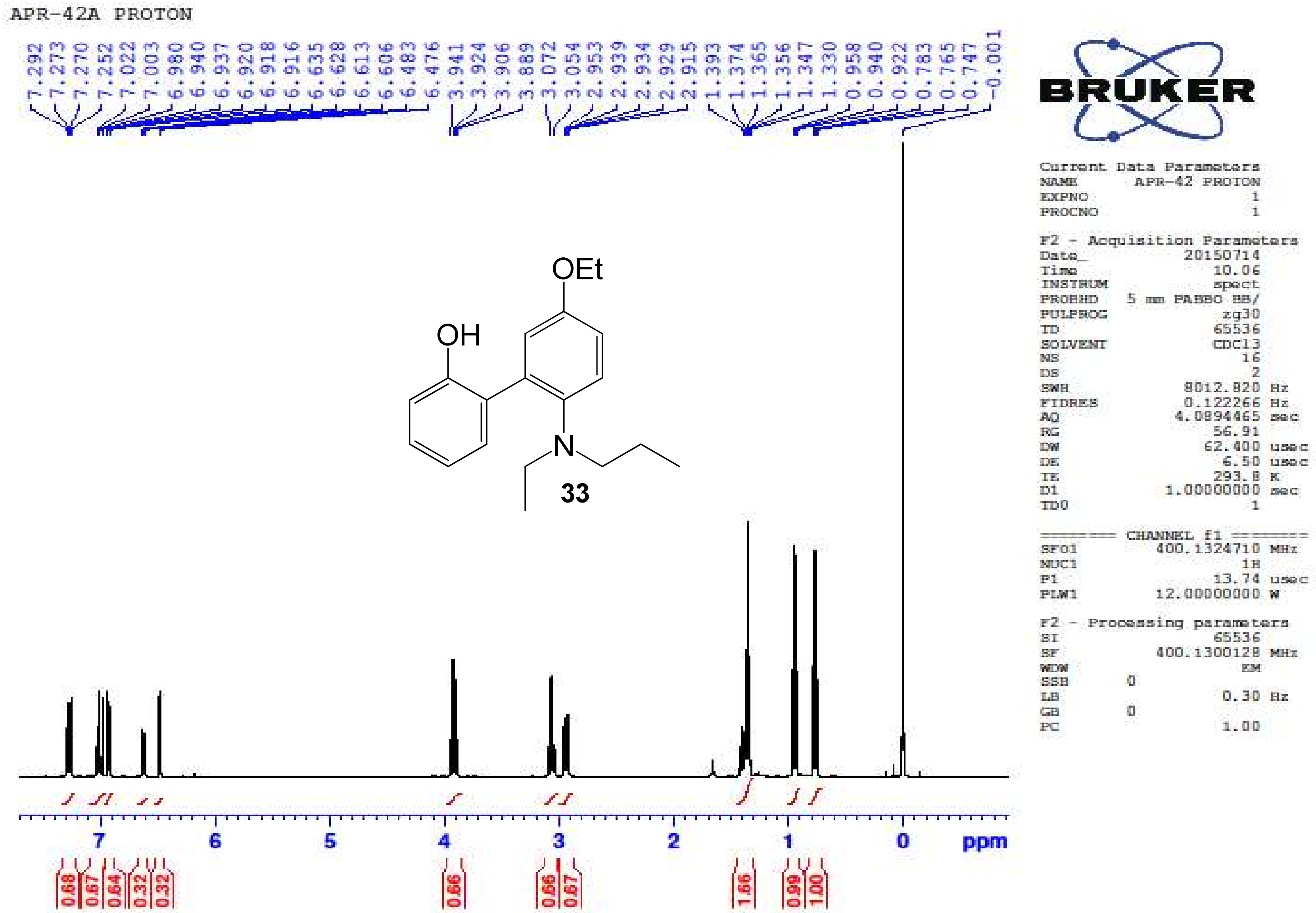

5'-Ethoxy-2'-(ethyl(propyl)amino)biphenyl-2-ol **(33)** – $^{13}$C NMR ($CDCl_3$, 100 MHz)

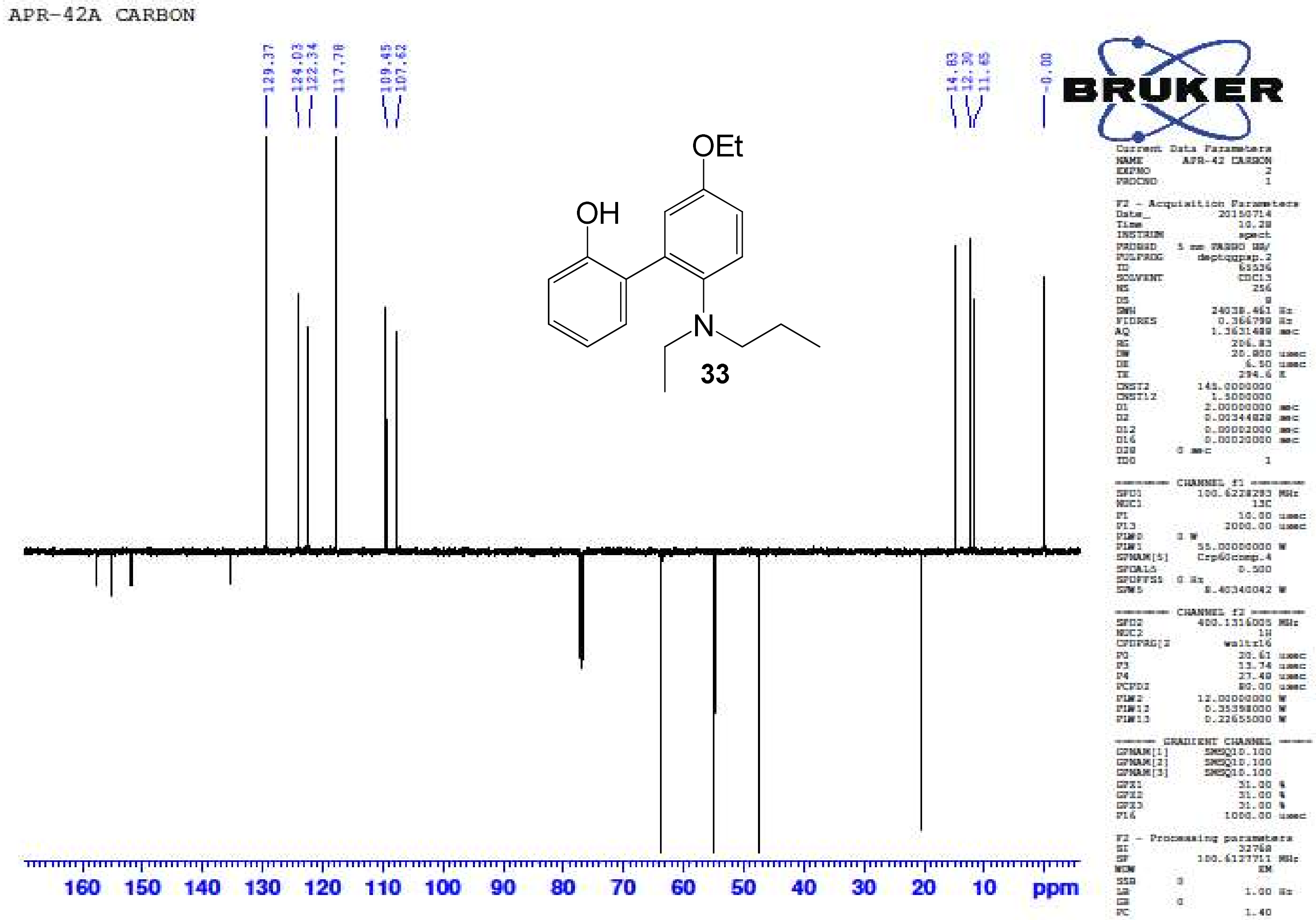

5'-Ethoxy-2'-(ethyl(propyl)amino)biphenyl-2-ol **(33)** – HR-ESIMS ($[M+H]^+$ Calculated: 300.1958, Observed: 300.1976)

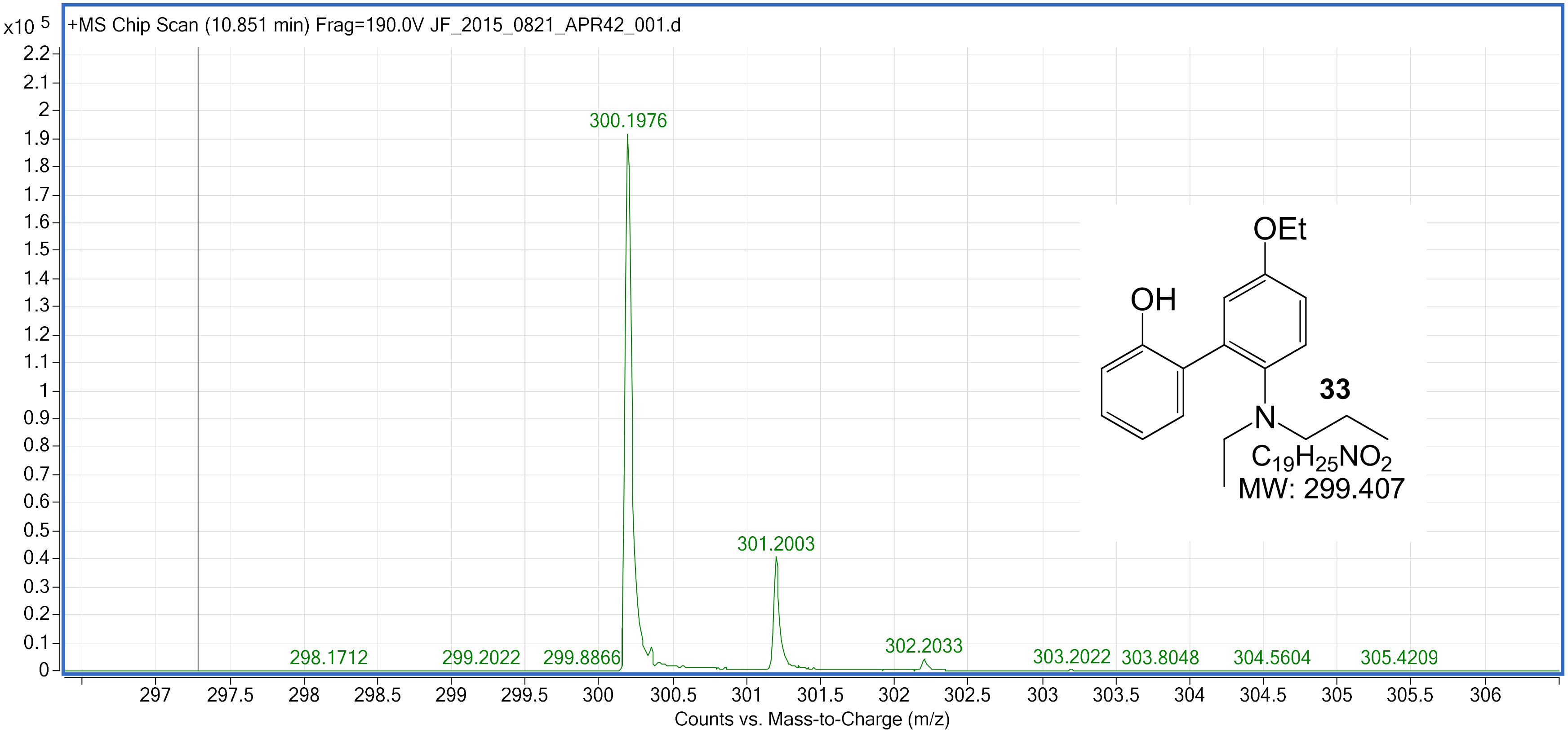

5-Bromo-5'-ethoxy-2'-(ethyl(propyl)amino)biphenyl-2-ol **(34)** – $^{1}H$ NMR ($CDCl_3$, 400 MHz)

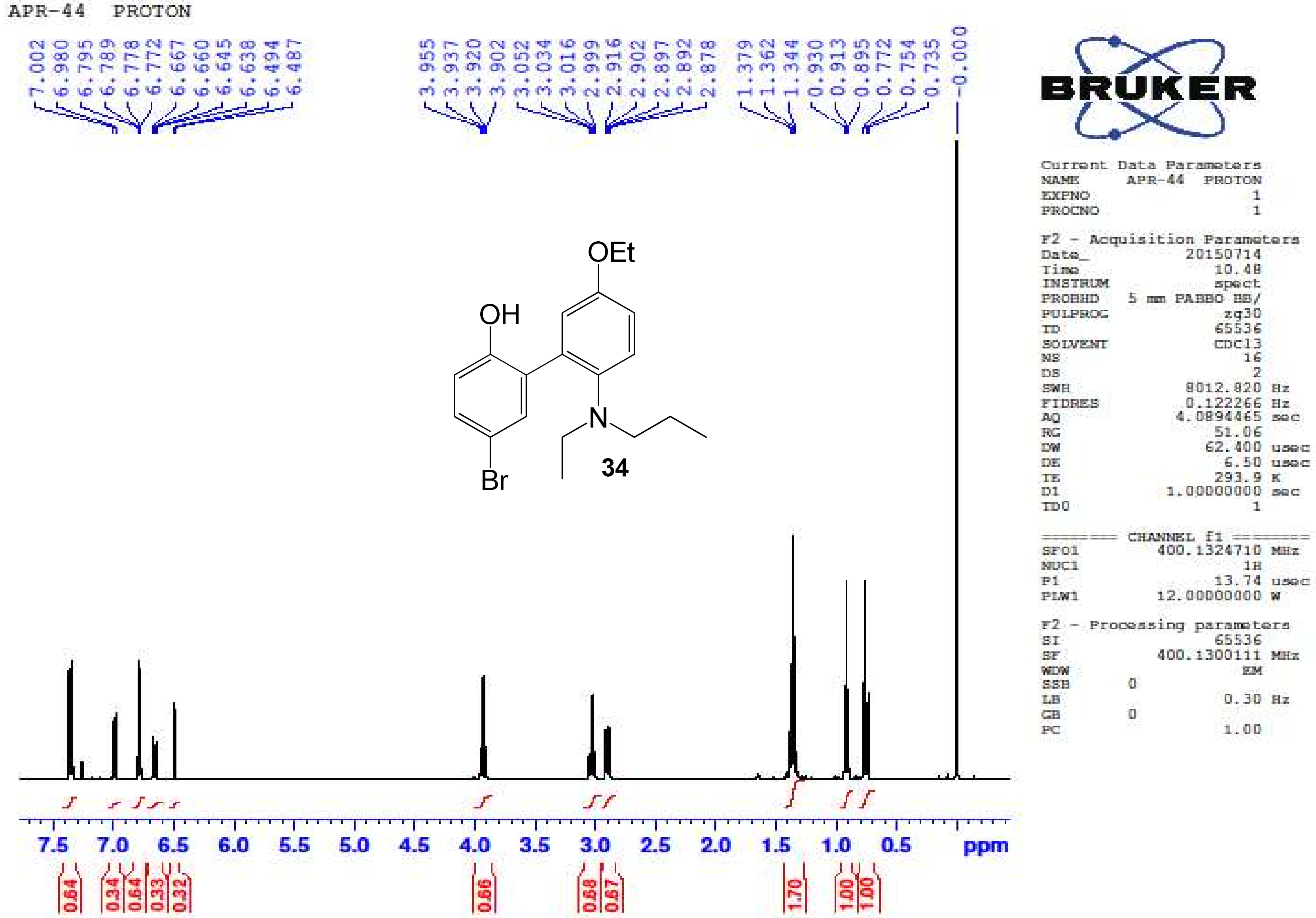

5-Bromo-5'-ethoxy-2'-(ethyl(propyl)amino)biphenyl-2-ol **(34)** – $^{13}C$ NMR ($CDCl_3$, 100 MHz)

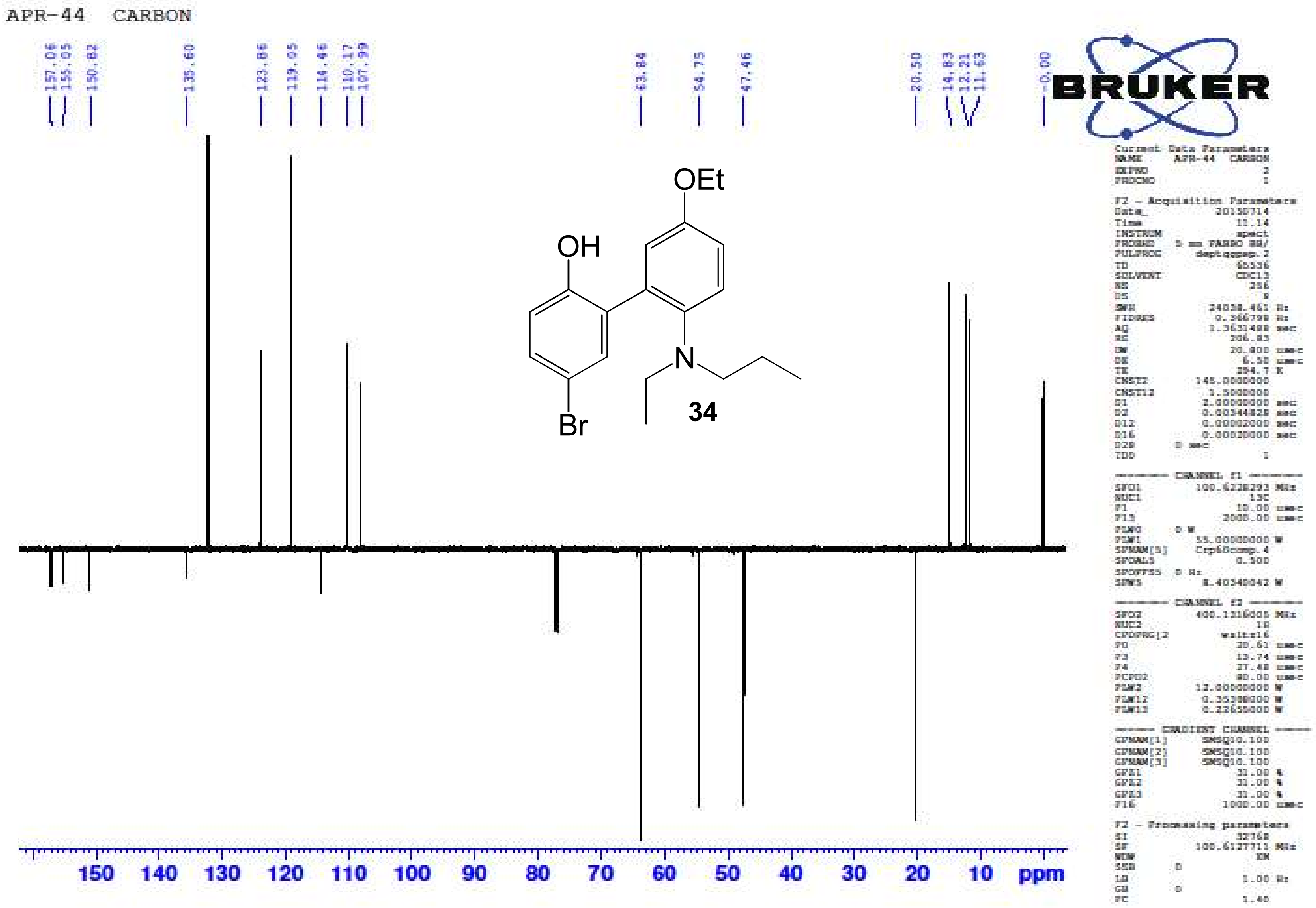

5-Bromo-5'-ethoxy-2'-(ethyl(propyl)amino)biphenyl-2-ol **(34)** – HR-ESIMS ($[M]^+$ Calculated: 378.1063, Observed: 378.1069)

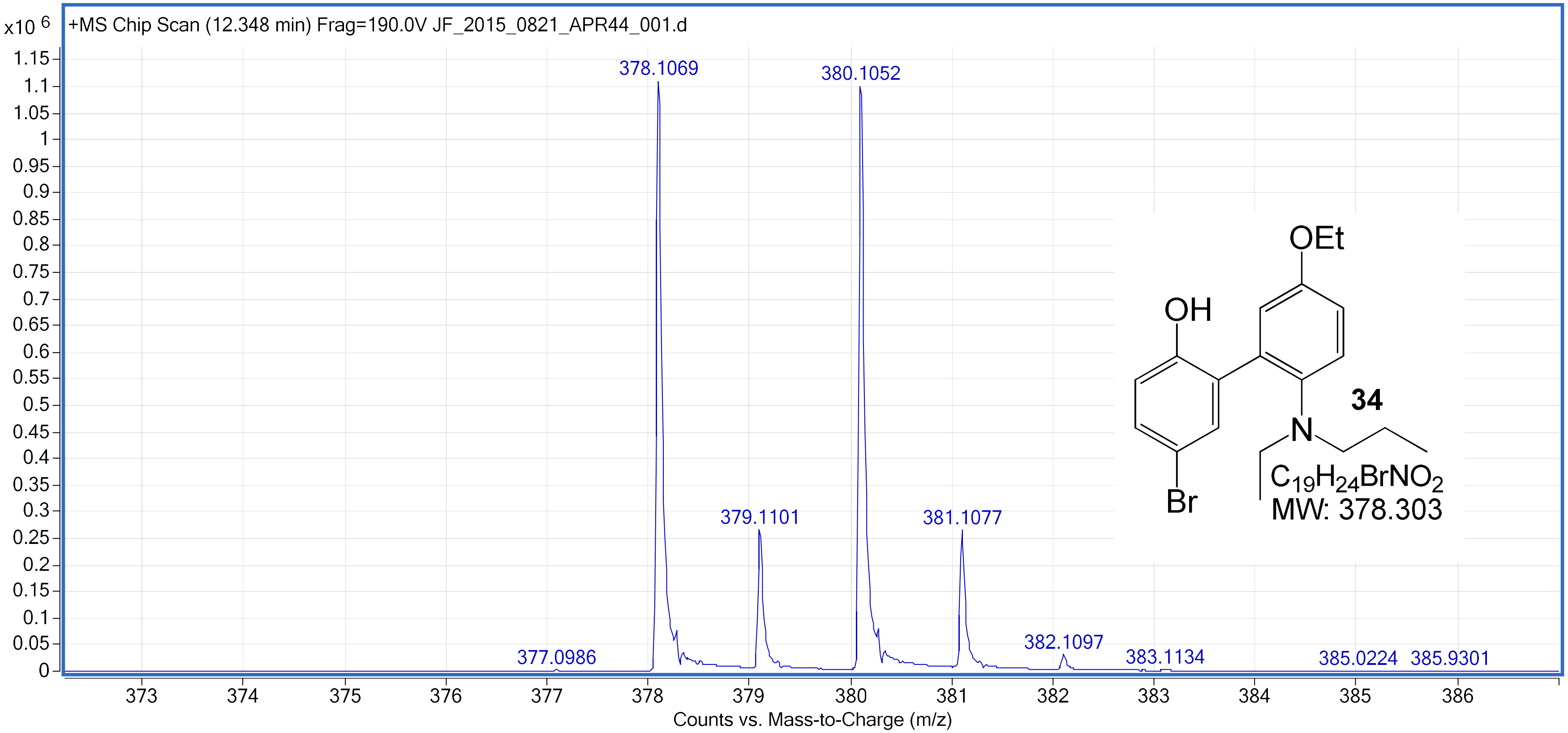

5-Bromo-2'-(diethylamino)-5'-ethoxybiphenyl-2-ol **(35)** – $^{1}H$ NMR ($CDCl_3$, 400 MHz)

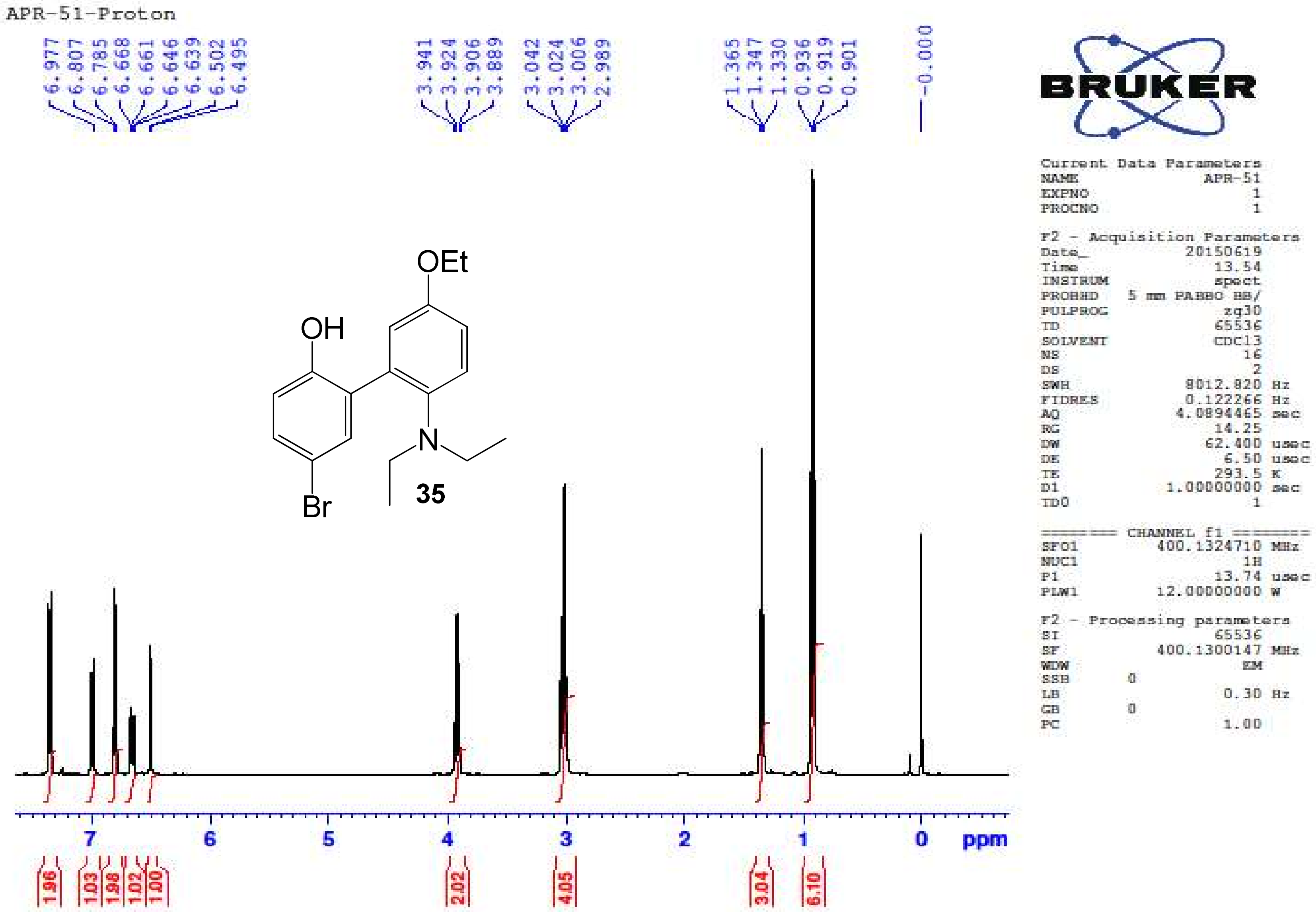

5-Bromo-2'-(diethylamino)-5'-ethoxybiphenyl-2-ol **(35)** – $^{13}C$ NMR ($CDCl_3$, 100 MHz)

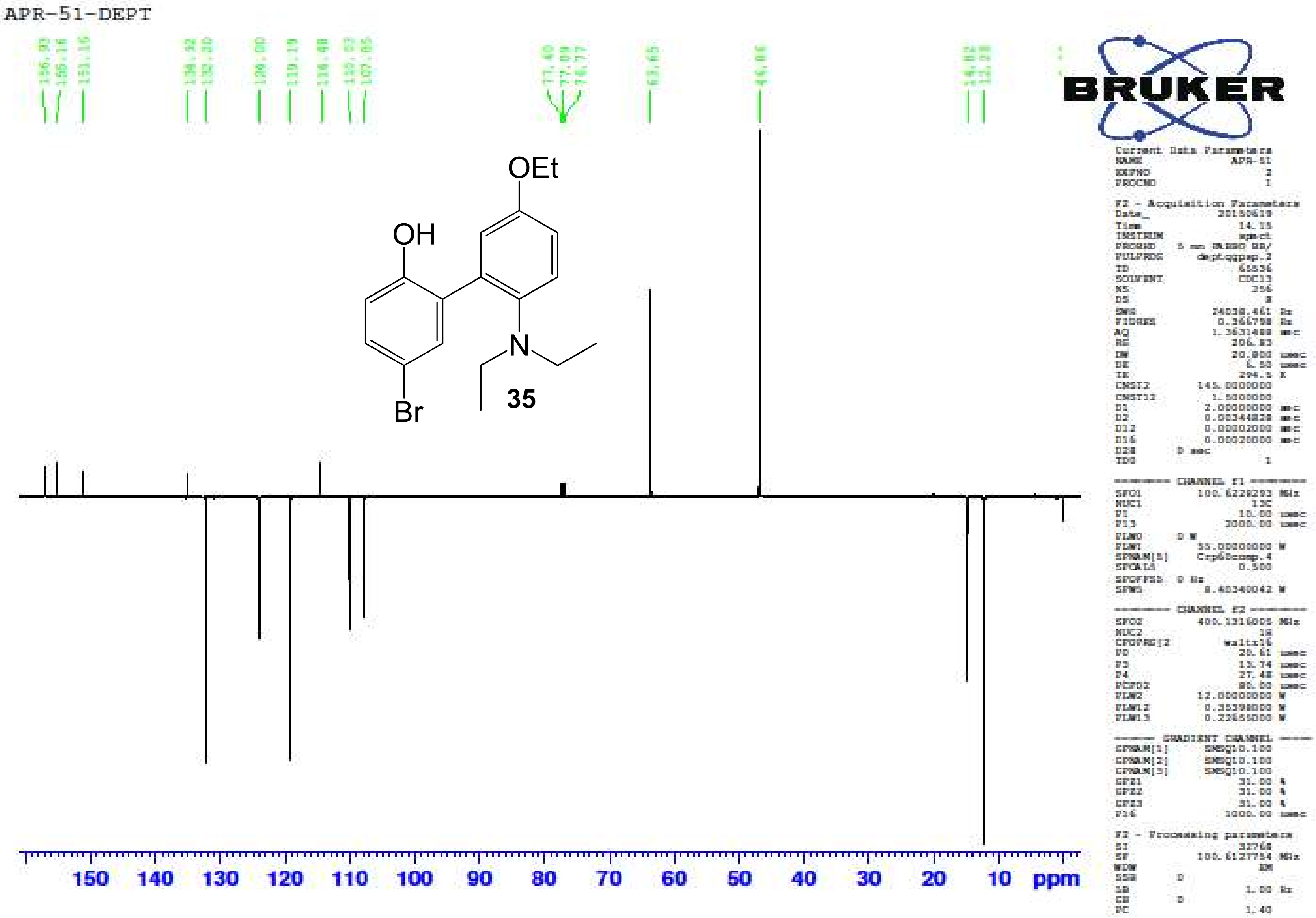

5-Bromo-2'-(diethylamino)-5'-ethoxybiphenyl-2-ol **(35)** – HR-ESIMS ($[M]^+$ Calculated: 364.0907, Observed: 364.0910)

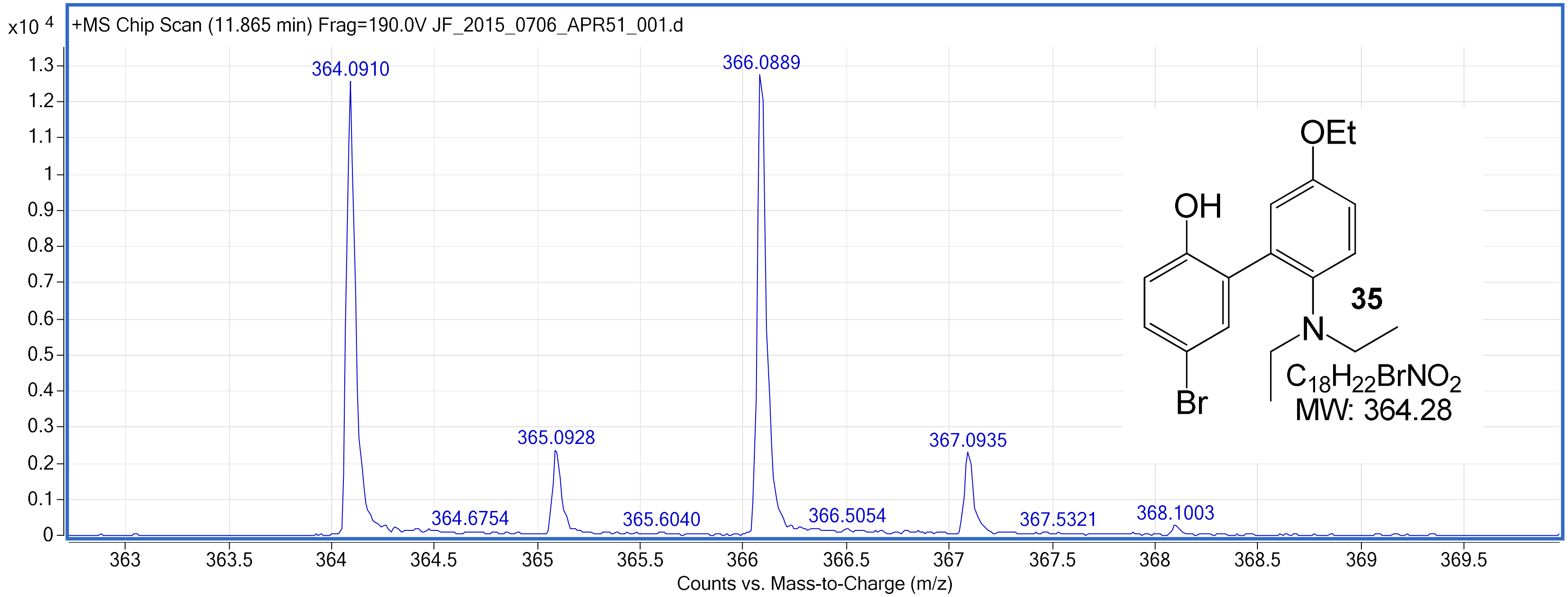
